# Integrated Proteomics analysis of baseline protein expression in pig tissues

**DOI:** 10.1101/2023.11.07.566009

**Authors:** Shengbo Wang, Andrew Collins, Ananth Prakash, Silvie Fexova, Irene Papatheodorou, Andrew R. Jones, Juan Antonio Vizcaíno

## Abstract

The availability of an increasingly large amount of public proteomics datasets presents an opportunity for performing combined analyses to generate comprehensive organism-wide protein expression maps across different organisms and biological conditions. *Sus scrofa*, the domestic pig, is a model organism relevant for food production and for human biomedical research. Here we reanalyzed 14 public proteomics datasets from the PRIDE database coming from pig tissues to assess baseline (without any biological perturbation) protein abundance in 14 organs, encompassing a total of 20 healthy tissues from 128 samples. The analysis involved the quantification of protein abundance in 599 mass spectrometry runs.

We compared protein expression patterns among different pig organs and examined the distribution of proteins across these organs. Then, we studied how protein abundances compared across different datasets and studied the tissue specificity of the detected proteins. Of particular interest, we conducted a comparative analysis of protein expression between pig and human tissues, revealing a high degree of correlation in protein expression among orthologs, particularly in brain, kidney, heart, and liver samples.

We have integrated the protein expression results into the Expression Atlas resource for easy access and visualisation of the protein expression data individually or alongside gene expression data.

## Introduction

In recent years, high-throughput mass spectrometry (MS)-based proteomics methods have made significant advances and have become essential tools in biological research ^1^. These improvements are the result of significant developments in MS instrumentation, chromatographic methods, sample preparation automation and computational analysis ^2^. The dominant experimental technique for MS-based proteomics has historically been data-dependent acquisition (DDA) bottom-up proteomics ^3^. Among the quantitative techniques, label-free DDA approaches are well accepted. However, Data Independent Acquisition (DIA) approaches are currently becoming increasingly popular.

In parallel to the technical developments, in recent years, the proteomics community has embraced open data practices, leading to a substantial increase in the availability of shared datasets in the public domain. The field has mirrored the progress witnessed in genomics and transcriptomics. The PRIDE database ^4^, as part of the ProteomeXchange consortium ^5^, is the most used proteomics data repository worldwide. The availability of extensive public proteomics datasets has paved the way for various applications, including meta-analysis studies involving the reanalysis and integration of quantitative proteomics datasets ^6–9^. By systematically reanalysing these datasets, original findings can be updated, confirmed and/or strengthened. Moreover, novel insights beyond the scope of the original studies can be obtained through alternative reanalysis strategies to those used in the original studies ^10^.

To enable the access to proteomics data by the wider scientific community, PRIDE is developing data dissemination and integration pipelines with existing popular resources at the European Bioinformatics Institute (EBI). Expression Atlas ^11^ (https://www.ebi.ac.uk/gxa/home) is a well-established database for gene expression data, and has more recently incorporated protein expression information derived from reanalysed datasets into its ’bulk’ section. As a result, the integration of proteomics expression/protein abundance data with transcriptomics information, primarily from RNA-Seq experiments, enhances the comprehensive understanding of molecular expression across various biological contexts. This approach ensures the long-term accessibility and integration of proteomics data, benefiting researchers, including those without expertise in proteomics, in their exploration of multi-omics information.

We have already performed combined analyses of baseline (without any perturbation) protein expression for human ^7^, mouse and rat tissues ^6^. Here we are reporting an analogous study of baseline protein expression in the model organism *Sus scrofa,* the domestic pig. The study of pig proteomics datasets is crucial for advancing food production, animal welfare, and human biomedical research, as it offers insights into genetic and environmental factors affecting farm animal production and leverages the close genetic and proteomic similarities between pigs and humans ^12, 13^. The resource PeptideAtlas provided a few years ago a build for pig, including extensive peptide and protein identification data, but is not providing protein abundance information ^8^. Additionally, PaxDB recently released a new version 5.0 ^14^, providing expression data coming from different vertebrates including *Sus scrofa,* but quantitative data is based on spectral counting, a semi-quantitative technique. Also, no tissue specific information is provided there, apart from liver. To the best of our knowledge, we are providing the first combined quantitative analysis of label-free DDA datasets in pig.

Here, we report the reanalysis and integration of 14 public label-free pig baseline tissue datasets including 14 organs and a total of 20 healthy tissues from 128 samples. The results were incorporated into Expression Atlas as baseline studies. Additionally, we report a comparative analysis of protein expression across pig and human tissues, among other analyses.

## Methods

### 1. Datasets

The PRIDE database hosted 165 publicly available MS proteomics datasets of *Sus scrofa* as of October 2022. For this study, we manually selected datasets based on several predefined criteria, which included: (i) label-free DDA studies from baseline tissues (without any perturbation), and without enrichment for post-translational modifications; (ii) datasets generated using Thermo Fisher Scientific instruments to avoid the heterogeneity introduced by data generated by other platforms; and (iii) datasets with sufficient sample metadata, manually curated from the original publication. This resulted in the identification of 14 pig datasets for further analysis.

Sample and experimental metadata were manually curated using Annotare ^15^, and adhering to the Investigation Description Format (IDF) and Sample-Data Relationship Format (SDRF) files ^16^, which are needed for integration of the data into Expression Atlas. The IDF file contains an overview of the experimental design, including details on experimental factors, protocols, publication information, and contact information. The SDRF file contains complementary information - the sample metadata that describes the relationships between various sample characteristics and the associated data files within the dataset.

### 2. Proteomics raw data processing

All datasets underwent analysis using MaxQuant version 2.0.3.0 ^17^ in multi-threaded mode on a Linux high-performance computing cluster for peptide/protein identification and protein quantification. Input parameters for each dataset, including MS1 and MS2 tolerances, digesting enzymes, fixed and variable modifications, were set according to the specifications provided in their respective publications and accounting for two missed cleavage sites. The false discovery rate (FDR) at both the peptide spectrum match (PSM) and protein levels was set to 1%. The remaining parameters of MaxQuant were set to the default values: a maximum of 5 modifications per peptide, a minimum peptide length of 7 amino acids, and a maximum peptide mass of 4,600 Da. For the “match between runs” option, a minimum match time window of 0.7 seconds and a minimum retention time alignment window of 20 seconds were applied. MaxQuant parameter files can be downloaded from Expression Atlas. The *Sus scrofa* UniProt ^18^ Reference proteome release-2021_04 (including isoforms, 49,865 sequences) was used as the target sequence database for the pig datasets. MaxQuant uses a built-in database of contaminants, and a decoy database was generated by reversing the input database sequences following the respective enzymatic digestion.

### 3. Post-processing

The post-processing of MaxQuant results followed the methodology detailed in previous publications ^6^. In short, after removing the protein groups labelled as potential contaminants, decoys, and those with less than 2 PSMs, the protein intensities in each sample were normalised by scaling the iBAQ intensity values with the total signal in each MS run and converting to parts per billion (ppb).

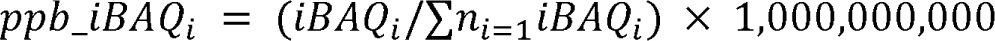

UniProt protein accessions, from the MaxQuant output-proteinGroups.txt file, were mapped to their Ensembl gene identifiers (ENSSSCG) using the ID mapping dataset (Release 2022/03) at the UniProt website (https://www.uniprot.org/id-mapping) ^19^. The resulting id mapping data file (idmapping_selected.tab), by default maps UniProt protein accessions to Ensembl gene identifiers of all pig breeds (i.e., Landrace, Pietrain, etc.) rather than to the reference breed, thus leading to multiple Ensembl gene identifiers being returned per UniProt protein identifier. To resolve this, we downloaded the *Sus scrofa* model Ensembl fasta peptide dump (Release 11.1 gene set, https://ftp.ensembl.org/pub/release-110/fasta/sus_scrofa/pep/Sus_scrofa.Sscrofa11.1.pep.all.fa.gz) and used this as a filter to keep only gene-mappings specific to the reference pig breed. We used the reference pig Ensembl gene identifiers for further downstream analysis.

During downstream post-processing we removed protein groups which mapped to more than one gene identifier and for cases where two or more protein groups mapped to the same gene identifier, protein intensities were aggregated using the median value. The parent genes to which the different protein groups were mapped to are equivalent to ‘canonical proteins’ in UniProt (https://www.uniprot.org/help/canonical_and_isoforms) and therefore the term protein abundance is used to describe the protein abundance of the canonical protein throughout the manuscript.

### 4. Integration into Expression Atlas

The normalised protein abundances along with the validated SDRF files, summary files detailing the post-processing quality assessment and MaxQuant parameter files (mqpar.xml) are available to download from Expression Atlas. Table 1 describes the datasets and their corresponding E-PROT identifiers.

**Table 1.** List of pig proteomics datasets and their main characteristics.

| Expression Atlas accession number | Proteomics dataset identifier* | Breed | Tissues | Organs | Mass spectrometer | Number of MS runs | Number of samples | Fraction | Number of protein groups† | Number of peptides† | Number of unique peptides† | Number of unique genes (canonical proteins) mapped† |
| --- | --- | --- | --- | --- | --- | --- | --- | --- | --- | --- | --- | --- |
| E-PROT-111 | PXD001800<sup>27</sup> | Landrace | retina | eye | Q Exactive Plus | 48 | 16 | yes | 4,579 | 36,992 | 28,055 | 3,643 |
| E-PROT-113 | PXD002918<sup>28</sup> | NA | biceps femoris | muscle | LTQ Orbitrap XL | 6 | 6 | no | 1,272 | 9,148 | 6,163 | 841 |
| E-PROT-114 | PXD003204<sup>29</sup> | Landrace | bile duct, heart, spleen, lung, brain, liver, diaphragm, pancreas, kidney | biliary system, heart, spleen, lung, brain, liver, diaphragm, pancreas, kidney | LTQ Orbitrap Elite | 108 | 9 | Yes | 5,969 | 85,655 | 57,338 | 5,065 |
| E-PROT-115 | PXD009577<sup>30</sup> | NA | heart | heart | Q Exactive HF | 30 | 6 | yes | 4,258 | 47,975 | 32,956 | 3,582 |
| E-PROT-117 | PXD011360<sup>31</sup> | Landrace x Piétrain | cecum, colon, ileum, Jejunum | intestine | Q Exactive Plus | 16 | 16 | no | 4,716 | 55,392 | 41,184 | 3,981 |
| E-PROT-118 | PXD011536<sup>32</sup> | NA | liver | liver | Q Exactive HF-X | 5 | 5 | no | 2,356 | 18,816 | 15,078 | 1,773 |
| E-PROT-119 | PXD011755<sup>33</sup> | House swine | eye | eye | LTQ Orbitrap XL | 204 | 12 | yes | 1,337 | 7,039 | 5,660 | 888 |
| E-PROT-120 | PXD012636<sup>34</sup> | NA | heart | heart | Q Exactive HF | 120 | 4 | yes | 7,747 | 151,177 | 94,341 | 6,982 |
| E-PROT-122 | PXD014893<sup>35</sup> | NA | biceps femoris, triceps muscles | muscle | Q Exactive HF-X | 8 | 4 | yes | 1,287 | 13,065 | 8,527 | 1,004 |
| E-PROT-124 | PXD016003<sup>36</sup> | NA | biceps femoris, triceps muscles | muscle | Q Exactive HF-X | 8 | 4 | yes | 2,379 | 22,409 | 15,650 | 1,885 |
| E-PROT-125 | PXD017671<sup>37</sup> | NA | liver | liver | Q Exactive HF-X | 8 | 8 | no | 3,128 | 32,215 | 24,585 | 2,576 |
| E-PROT-126 | PXD019852<sup>38</sup> | NA | heart | heart | Q Exactive HF-X | 8 | 8 | no | 1,527 | 13,511 | 10,085 | 1,143 |
| E-PROT-130 | PXD026910<sup>39</sup> | NA | subcutaneous adipose tissue, mesenteric adipose tissue | adipose tissue | Q Exactive HF | 10 | 10 | No | 2,586 | 23,718 | 17,676 | 2,107 |
| E-PROT-131 | PXD027772<sup>40</sup> | Landrace x Swabian-Hall | skeletal muscle, heart | skeletal muscle, heart | Q Exactive HF | 20 | 20 | no | 3,242 | 39,140 | 25,993 | 2,797 |
| TOTAL | 14 datasets |  | 20 tissues | 14 organs |  | 599 MS runs | 128 samples |  |  |  |  |  |

### 5. Protein abundance comparison across datasets

The normalised protein abundances (in ppb values) within each dataset were transformed into ranked bins as described in ^7^. Briefly, normalised protein abundance (ppb) of each MS run was sorted from lowest to highest and binned into 5 equal length bins. Proteins ranked in the lowest bin (bin 1) represent the lowest abundance, and correspondingly proteins ranked in bin 5 have the highest abundance. To analyse and compare the data effectively, protein abundances from ’tissues’ were grouped into ’organs’. For example, the Ileum and jejunum ‘tissues’ were grouped as ‘small intestine’. Similarly, triceps, biceps femoris, diaphragm and skeletal muscle were grouped as ‘muscle’. When combining tissues into organs, median bin values were used.

Proteins of all the samples were selected for UMAP (Uniform Manifold Approximation and Projection) ^20^ representation and analysed for binned abundance values using the R programming language (https://www.R-project.org/). Pearson correlation coefficients (rp) were calculated for all samples based on paired complete observations and used to generate a heatmap. Missing values were marked as NA (not available). For each organ, the median rp was calculated from all paired rp values of its respective sample. Columns and rows of the samples were clustered hierarchically using Euclidean distances.

### 6. Organ-specific expression profile analysis

To make comparisons of protein expression across organs based on organ specificity, we grouped the proteins into three categories based on the classification scheme of Uhlen *et al.*^21^: (1) “Organ-enriched”: present in one unique organ with bin values 2-fold higher than the mean bin value across all organs; (2) “Group enriched”: present in at least 7 organs, with bin values 2-fold higher than the mean bin value across all organs; and (3) “Mixed”: the remaining canonical proteins that are not part of the above two categories.

We then performed Gene Ontology (GO) term enrichment analysis through an over-representation test on the “Organ-enriched” and “Group enriched” using the mapped gene lists for each organ. The computational analysis was carried out in the R programming language with the package clusterProfiler ^22^ version 3.16.1, by using the function enrichGO() for the GO term over-representation test. The pvalueCutoff was set to 0.05 and the qvalue Cutoff to 0.05.

### 7. Comparison of protein expression values between pig and human tissues

The orthologous genes for pig and human were obtained following the procedure described in the Ensembl BioMart ^23^. Briefly, we first selected the “Ensembl genes 110”, then chose “Human genes (GRCh38.p14)”, clicked on “Filters” in the left menu, then unfold the “MULTI SPECIES COMPARISONS” box, ticked the “Homolog filters” option and chose “Orthologous Pig Genes” from the drop-down menu. Then, we clicked on “Attributes” in the left menu, unfold the “Pig ORTHOLOGS” box and selected the pig gene ID and pig gene name. Finally, we clicked on the “Results” button (top left) to download the list of orthologous genes between human and pig. The orthologous gene list was filtered to include only parent gene identifiers from pig samples in this study and the parent genes of human samples described in our previous study using human baseline tissue samples ^7^.

We used the calculation of “edit distance” ^7^ of a protein, which was computed as the difference between two pairs of protein abundance bins in pigs and humans. The following categories were used to classify their groups of protein expression samples: (1) “Group A“: protein abundance is similar between human and pig tissues; (2) “Group B“: protein abundance is higher in human tissues when compared to pig tissues; and (3) “Group C “: Protein abundance is higher in pig tissues when compared to human tissues.

A GO term enrichment analysis was performed using the mapped gene lists for each organ in each group (“Group A“, “Group B“, or “Group C“) as foreground, and the gene list of all three groups as the background. The settings were the same used for organ-specific expression profile analysis in the previous section.

The one-to-one mapped ortholog identifiers were used to compare pig and human protein intensities. Additionally, their normalised protein abundance (using ppb values) in 10 organs (adipose tissue, brain, colon, heart, kidney, liver, lung, pancreas, small intestine, and spleen) was used to assess pairwise correlations. Linear regression was calculated using the linear fit ‘lm’ method in the R programming language.

### 7. Correlation between gene (RNA-seq) and protein expression

One pig RNA-seq experimental tissue baseline dataset (the only one available) was obtained from Expression Atlas (dataset E-MTAB-5895). The dataset was composed of pig samples from a Duroc breed ^24^. Transcriptomics data had been previously collated in Expression Atlas, and FPKMS (Fragments Per Kilobase of transcript per Million mapped reads) data were computed by iRAP (https://github.com/nunofonseca/irap) based on the raw data, which were first averaged based on technical replicates, then quantile normalised within each set of biological replicates using limma ^25^ and finally averaged again over all biological replicates. Biological metadata were collected in the SDRF format, consistent with the proteomics data.

### 8. Comparison of protein abundance with spectral counting values from PaxDB

We compared the protein abundances generated in our study with the protein abundance data from PaxDB version 5.0 (https://www.pax-db.org/) ^26^ available for *Sus scrofa*. Normalised iBAQ abundances were compared with the spectral counting abundances for liver, the only matching organ available. This comparison was not possible for other pig organs as other data in PaxDB are labelled as ‘whole organism’. Ensembl gene ids (ENSSSCG) were mapped to protein ids (ENSSSCP) in PaxDB using the Ensembl BioMart, as described in this tutorial ^23^.

## Results

### 1. Pig proteomics datasets

In summary, we obtained protein expression data from 20 healthy tissues in 14 organs, coming from 14 public datasets. The analyses covered a total of 599 MS runs from 128 samples that were annotated as healthy/control/non-treated samples, thus representing baseline protein expression. Non-control/disease samples associated with these datasets were also analysed but are not discussed here. Normalised protein abundances values (as ppb) from both control/healthy/non-treated and disease/treated tissue samples are available to view as heatmaps in Expression Atlas. The protein abundances along with sample annotations, the sample quality assessment summary and experimental parameter inputs for MaxQuant can be downloaded from Expression Atlas as text files. The total number of proteins and peptides identified in these datasets are shown in Table 1.

### 2. Protein coverage across organs and datasets

A total of 7,767 protein groups were identified from the reanalysis of the 14 pig datasets, among which 2,164 protein groups (27.9%) were uniquely present in only one organ and 523 protein groups (6.7%) were ubiquitously observed (Table S1 in Supporting File 2). However, it should be emphasised that a specific list of typical proteins detected in only one organ should be treated with caution, as the FDR of this list will be amplified due to the accumulation of false positives when the datasets were analysed individually. For proteins detected in four or more datasets, this should not be a problem, as from the common number of decoy protein hits across datasets, a protein FDR of less than 1% could be inferred for those proteins (Figure S1 in Supporting File 1).

Protein groups were mapped to 7,780 genes (which are equivalent to canonical proteins, the term that we will be using from now on in the manuscript). The largest number of canonical proteins was detected in samples from heart (6,264, 80.5% of the total) and the lowest number in samples from adipose tissue (1,913, 24.6%) and from the biliary system (1,983, 25.5%) (Figure 1A). The lower number of proteins identified in the biliary system could be attributed to the smallest sample size (only one sample out of 128, 0.08%). Dataset PXD012636, a dataset containing pig heart samples, which was fractionated, provided the highest number of detected canonical proteins (6,062, 77.9%), whereas the smallest number of proteins were detected in dataset PXD002918 (biceps femoris, 789, 10.1%, non-fractionated dataset) (Figure 1B).

**Figure 1.**
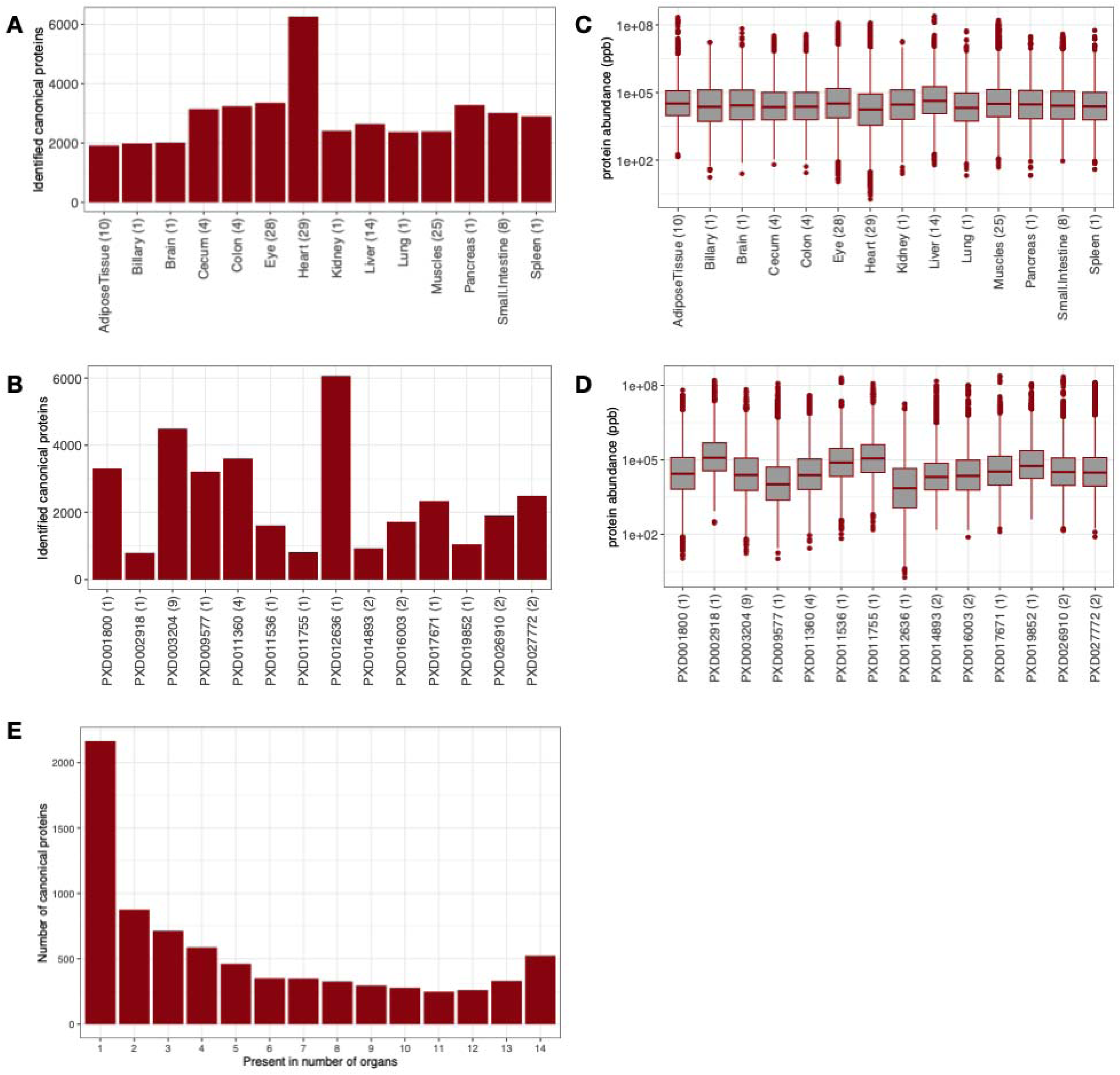
Distribution of canonical proteins detected per organ and dataset. (**A**) Number of canonical proteins identified across different pig organs. The number in brackets denotes the number of samples. (**B**) Number of canonical proteins identified in different datasets. The number in the brackets indicates the number of unique tissues in the dataset. (**C**) Range of normalised iBAQ protein abundances across different organs. The number in brackets indicates the number of samples. (**D**) Range of normalised iBAQ protein abundances across different datasets. The number in brackets indicates the number of unique tissues in the dataset. (**E**) Distribution of canonical proteins identified across different organs.

We studied the normalised protein abundance distribution in organs (Figure 1C) and found that all organs had similar median abundances. However, one cannot attribute biological meaning to these observations, since the method of normalisation by definition fixes each sample to have the same “total abundance”, which then gets shared out amongst all proteins. The normalised protein abundance distribution in datasets indicated a lower than median abundance detected in the dataset PXD012636 (heart), as a direct result of more proteins being detected overall in this dataset (Figure 1D). In terms of distribution of proteins detected per organ, most proteins were found in just one organ (Figure 1E).

### 3. Protein abundance comparison across organs and datasets

Next, we studied how protein abundances compared across different datasets and organs. To make protein abundance values more comparable between datasets, we transformed the normalised iBAQ intensities into ranked bins as explained in ‘Methods’: i.e., proteins included in bin 5 are highly abundant whereas proteins included in bin 1 are expressed in the lowest abundances (among the detected proteins). We found that 494 (6.3%) proteins were expressed in at least 3 organs, with a median bin value greater than 4 (not including 4). At the other end of the scale, 337 (4.3%) canonical proteins were expressed in at least 3 organs at a median bin value less than 2 (not including 2). The bin transformed abundances in all organs and in all datasets are provided in the Table S2 and S3 in Supporting File 2.

To compare protein expression across all pig organs, we calculated pairwise Pearson correlation coefficients (rp) for the 128 samples (Figure 2A). We observed a good correlation of protein expression within the liver (median rp = 0.77) and muscle (median rp = 0.65) samples. We then performed a cluster analysis using UMAP ^20^ on all samples to test the effectiveness of the bin-transformed method in reducing batch effects (Figure 2B). We observed that samples from different datasets belonging to the same organ were generally clustered together. For example, liver samples from datasets PXD003204, PXD011536 and PXD017671 clustered together (colour blue in Figure 2B). Additionally, muscle samples from PXD002918 (tissue biceps femoris), PXD003204 (tissue diaphragm), PXD014893 (tissue biceps femoris and triceps muscle) and PXD016003 (tissue biceps femoris, triceps muscle) clustered together too (colour purple). Similarly, heart samples from datasets PXD003204, PXD009577, PXD012636 and PXD019852 clustered together as well (colour dark green).

**Figure 2.**
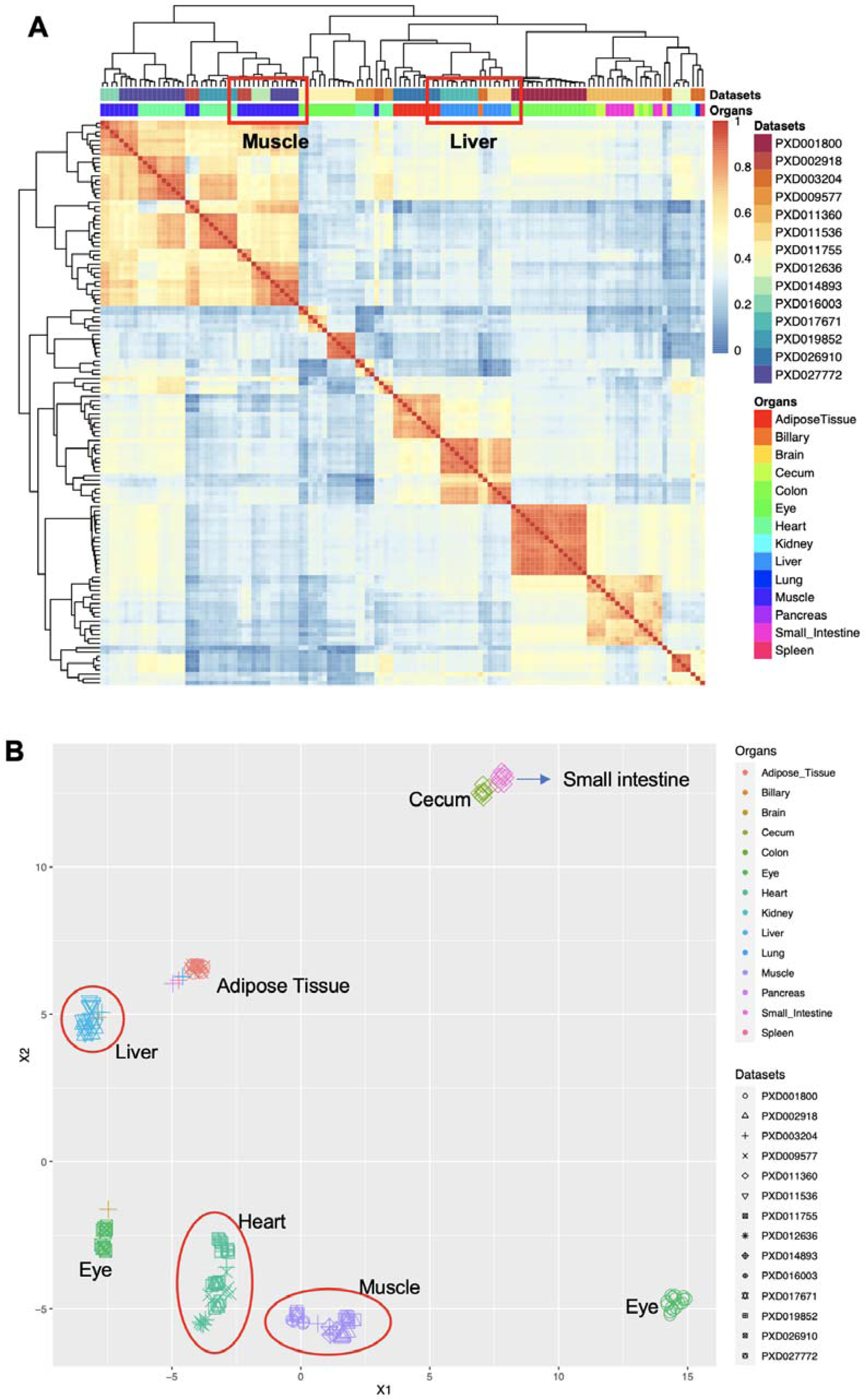
**A.** Heatmap of pairwise Pearson correlation coefficients across all pig samples among datasets and organs. The colours on the heatmap represent the correlation coefficients calculated using the bin transformed values. Hierarchical clustering of the columns and rows of the samples was performed using Euclidean distances. **B.** UMAP representation among different datasets and organs. Groups of datasets coming from the same organ are highlighted.

### 4. The organ elevated proteome and the over-representative biological processes

To get more insights about organ expression specificity, proteins were classified into three different groups: ‘group-enriched’, “organ-enriched” and “mixed” (see ‘Methods’ for details, Table S4 in Supporting File 2). The analysis (Figure S2 in Supporting File 1) showed that on average, 26.8% of the total elevated canonical proteins were organ group-specific in pig. In addition, 4.2% were unique organ-enriched in pig. The highest ratio of group-specific was found in pancreas (36.8%), and the highest ratio of organ-enriched proteins was found in heart (23.9%).

A GO enrichment analysis (see ‘Methods’) was performed on those proteins that were ‘organ-enriched’ and ‘group-enriched’, Overall, 310 GO terms were found statistically significant in all organs. The most two significant GO terms were ‘organic acid metabolic process’ (GO:0006082) and ‘small molecule catabolic process’ (GO:0044282), both in liver and in the biliary system. These terms were followed by ‘RNA processing’ (GO:0006396), in pancreas. For the whole list of GO terms enriched for each organ, see the Table S5 in Supporting File 2.

### 5. Comparative analysis of pig and human protein expression

We also performed a comparative analysis of protein abundances (in bins) between the pig baseline tissue datasets with the results obtained in our previous analogous study involving human datasets coming from baseline tissues, which was performed using the same overall methodology ^7^ (Table S6 in Supporting File 2). Pig is often used as a model organism for human biomedical research and then it is interesting to compare protein expression in both organisms. We calculated metrics to study the differences between the protein abundances for all organs found in the two studies (see ’Material and Methods’ for full details), as shown in Figure 3 (also see Table S7 in Supporting File 2). Three groups of proteins were found according to their protein expression levels: (i) “Group A“: protein expression is similar between human and pig tissues; (ii) “Group B“: protein expression is higher in human tissues; and (iii) “Group C “: protein expression is higher in pig tissues.

**Figure 3.**
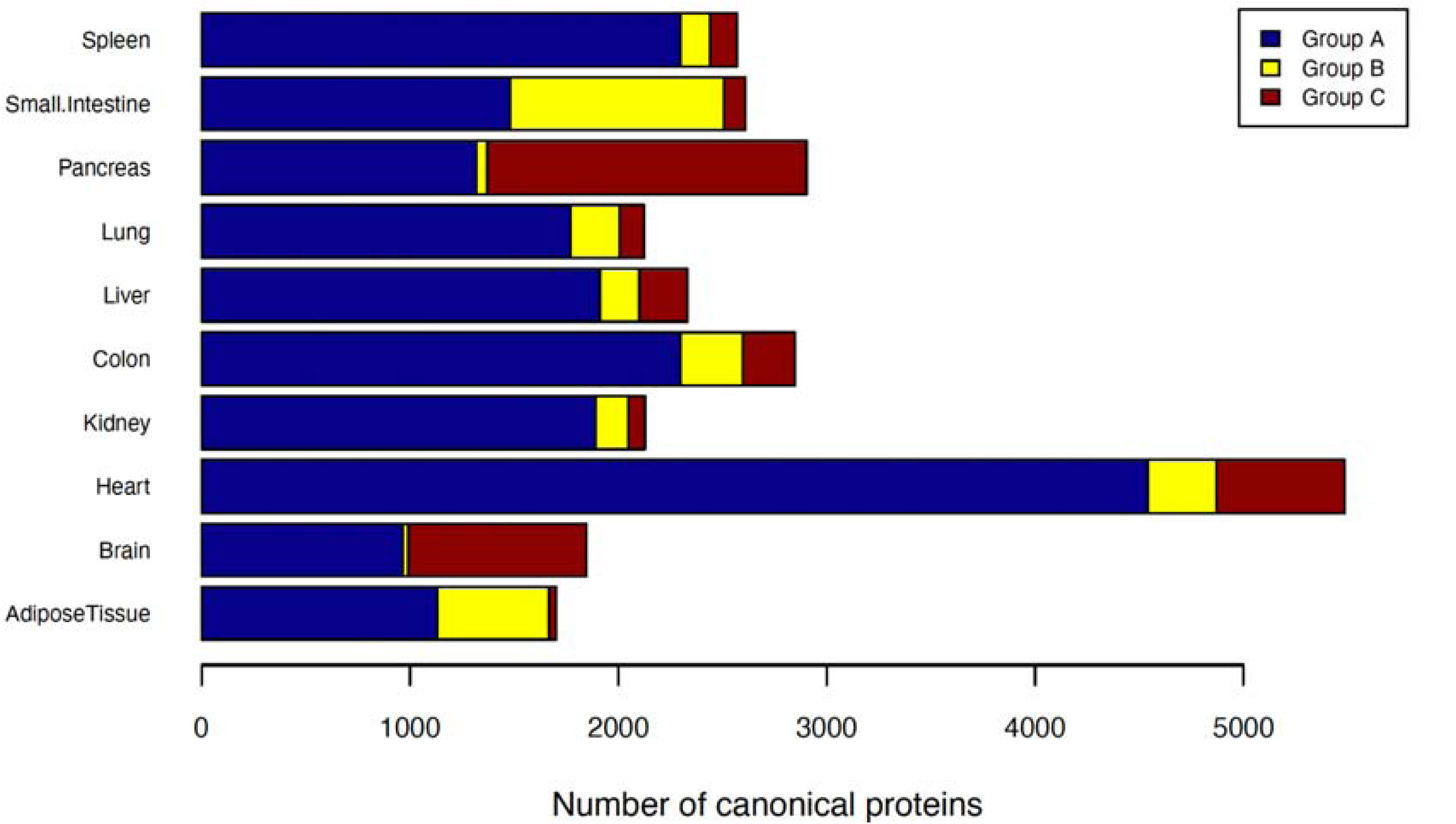
Organ specificity of canonical proteins based on edit distances between pig and human. The canonical proteins detected both in pig and human samples, were classified into three groups: “Group A” (similar protein expression levels between human and pig), “Group B” (higher protein expression in human tissues) and “Group C” (higher protein expression in pig tissues).

We found that for pig, protein expression levels were higher in pancreas, brain, and heart than in the corresponding human tissues, whereas protein expression levels were higher in the human’s small intestine and adipose tissue, when compared to the corresponding pig tissues. For other organs, the number of proteins in “Group B” and “Group C” were quite similar. Since different sizes and counts of datasets have been used for the different organs in both species, the organ-level trends reflect the results found in our studies. In our view, they are not necessarily meaningful for understanding species level differences. Instead, for individual proteins, or groups of related proteins, the comparison gives a potentially useful guide to relative protein abundance between orthologous pairs in individual organs.

We then performed a GO enrichment analysis ^41^ per organ of the proteins included in the three groups using GO terms related to biological processes (see ‘Methods’). We found 740 GO terms to be enriched in all organs overall (see all enriched GO terms in Table S8 in Supporting File 2), and in particular in heart and kidney. For instance, in “Group A”, we found enrichment for ‘intracellular transport’ (GO:0046907) in 9 organs (brain, colon, heart, kidney, liver, lung, pancreas, small intestine and spleen). In “Group B”, we observed ‘ribonucleoside monophosphate biosynthetic process’ (GO:0009156) enriched in adipose tissue and ‘peptide biosynthetic process’ (GO:0043043) in small intestine. Also, in “Group C”, we observed ‘vesicle-mediated transport’ (GO: 0016192) enriched in brain and ‘carboxylic acid metabolic process’ (GO:0019752) in pancreas. For the whole list, also including the GO terms enriched for other organs, see Table S8 in Supporting File 2.

### 6. Comparison of protein abundances across orthologs between pig and human datasets

In a previous study we compared the expression of canonical proteins found in three different species: human, mouse and rat ^6^. Here, we used the same approach to compare canonical protein expression between human ^7^ and pig organs. Overall, 13,248 detected human canonical proteins were compared with 7,800 detected pig canonical proteins (Table S9 in Supporting File 2). When comparing protein abundance (in ppb), we only considered the corresponding orthologous genes with unambiguous (one-to-one) mappings, which resulted in 6,811 common protein orthologs.

When comparing the protein expression of orthologs in human and pig, we observed a relatively overall high correlation in protein abundance in heart (R^2^ = 0.62) and liver (R^2^ = 0.53), medium correlation for brain (R^2^ = 0.44), colon (R^2^ = 0.42), kidney (R^2^ = 0.40) and spleen (R^2^ = 0.39), and low correlation in small intestine (R^2^ = 0.18), lung (R^2^ = 0.21), pancreas (R^2^ = 0.26), and adipose tissue (R^2^ = 0.27) (Figure 4).

**Figure 4.**
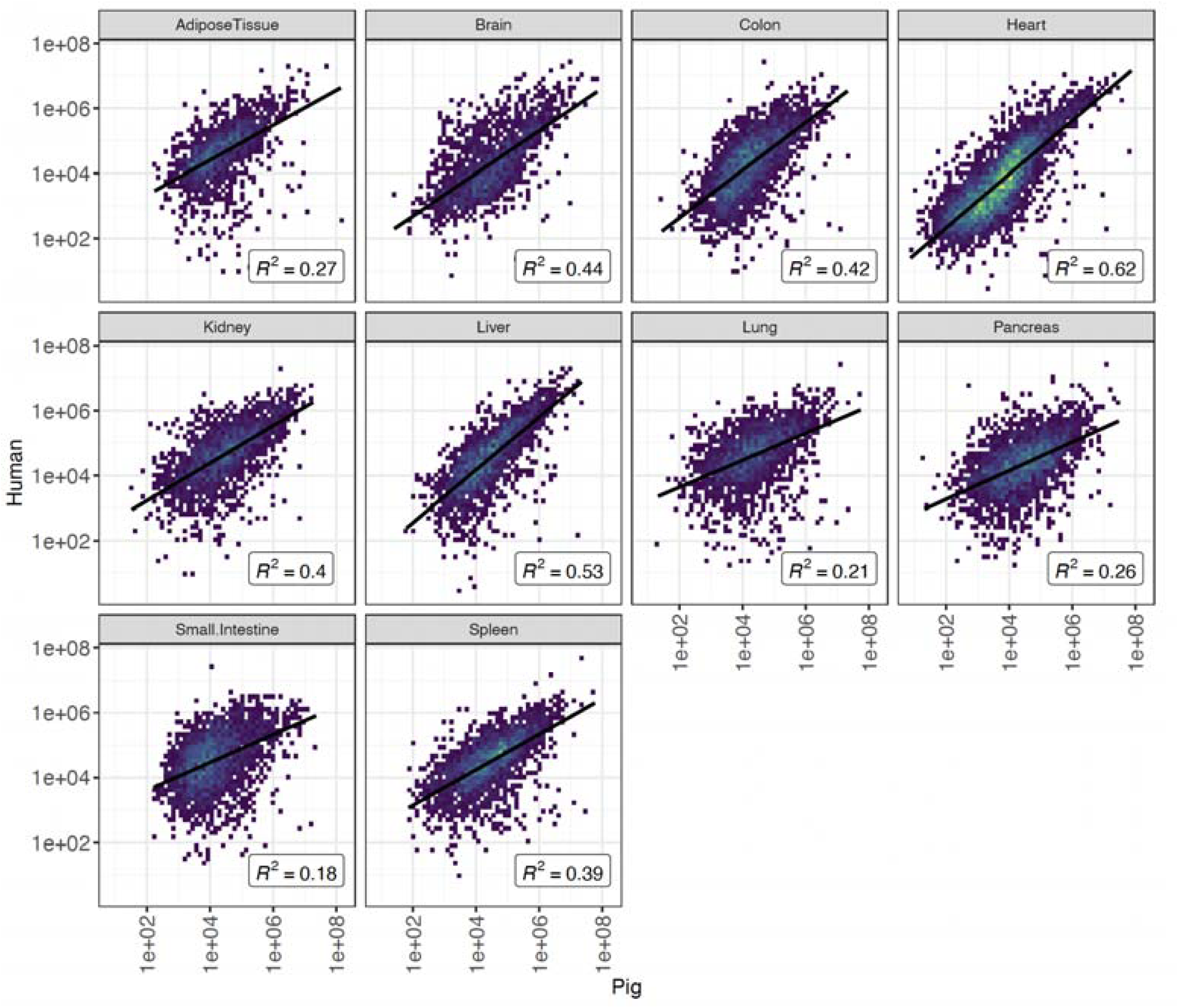
Comparison of protein abundance between orthologs of pig and human in various organs.

We also investigated the correlation of protein expression across different organs in pigs across the datasets used in this study (Figure 5). The data sometimes followed expected distributions whereby tissues predicted to be more similar to each other in terms of biological function have higher pairwise correlations e.g., small intestine and colon (R^2^ = 0.87) and spleen and pancreas (R^2^ = 0.78). However, other high correlations were found between lung and spleen (R^2^ = 0.83) and lung and pancreas (R^2^ = 0.71) - likely due to these data originating from a single study. The correlation between the brain and the remaining organs was generally low, as it would be expected. A plotted representation of the abundance of all sorted proteins for both species is provided in Figure S1 in Supporting File 3.

**Figure 5.**
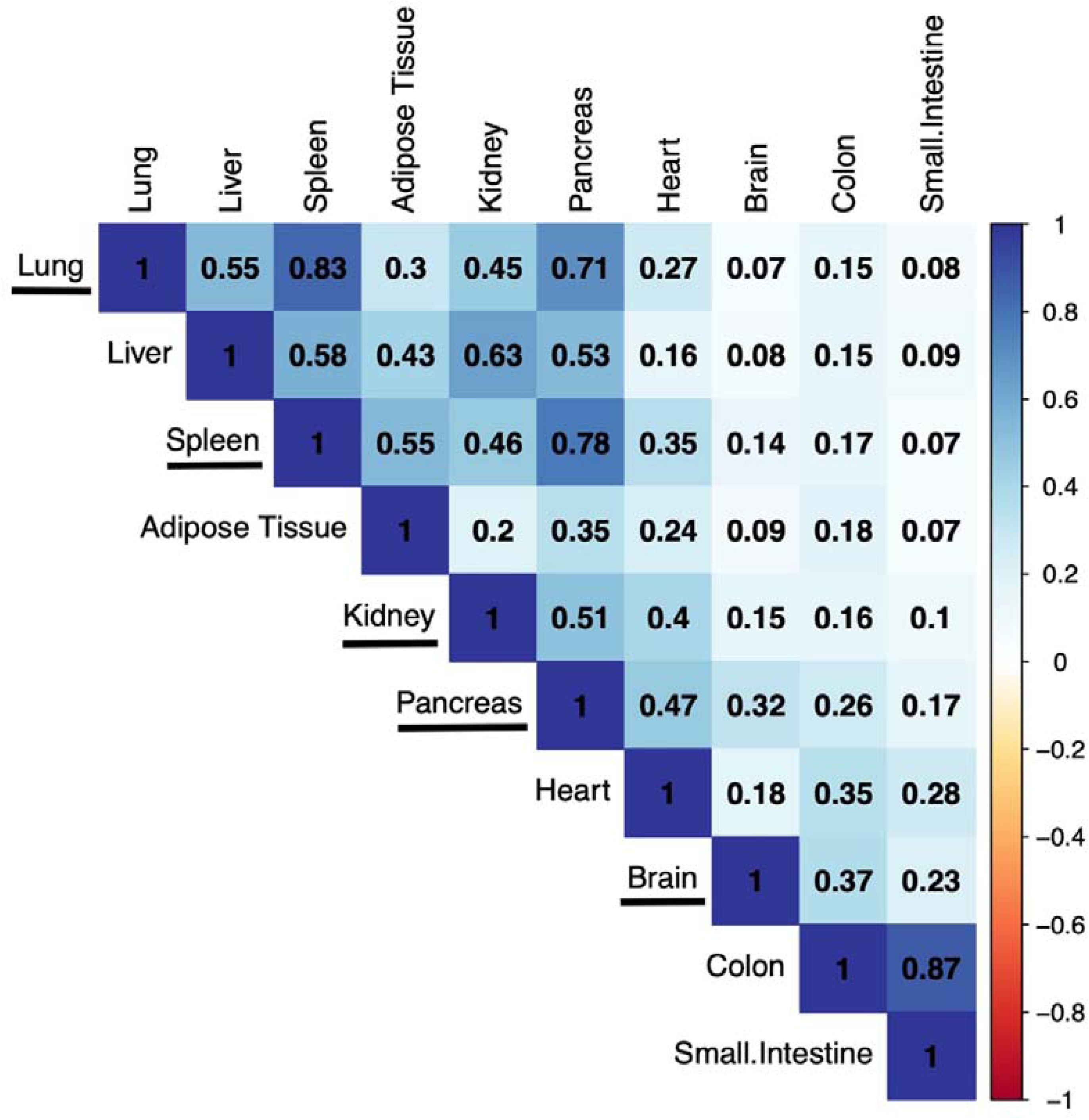
Protein expression correlation among all pig organs across the datasets included in this study. The organs underlined all come from a single dataset (PXD003204).

### 7. Comparison between gene (RNA-seq) and protein expression, and comparison with protein expression data in PaxDB

We also investigated the correlation between gene (RNA-seq based) and protein expression in baseline tissue pig datasets. For that we used the only suitable dataset available in Expression Atlas (dataset E-MTAB-5895) ^24^. We compared the normalised iBAQ protein abundances (ppb) with the baseline RNA-seq expression (FPKM). To compare expression across different organs we grouped the RNA-seq expression from various tissues into their respective organs by using their median values. We did not observe a strong correlation between protein and RNA expression across various organs (Figure S3 in Supporting File 1). The lowest correlation between protein and RNA expression was observed in lung (R^2^ = 0.06) and the highest was observed in heart (R^2^ = 0.20). In our view, these low correlations could be due to the inherent limitations in this comparison i.e. (i) samples are not paired; and (ii) the different pig breeds used (Duroc for the RNA-seq study and several others for the protein expression studies, see details in Table 1).

In addition, we compared the protein abundance of organ liver generated in this study with data from the PaxDB resource generated using a spectral counting method. We observed that protein abundance values (fraction of total (FOT) normalised ppb values) calculated using iBAQ in this study correlated till a limited extent (R^2^ = 0.47) with the semi-quantitative values available in PaxDB for (Figure S4 in Supporting File 1). Unfortunately, it was not possible to perform this correlation for other organs (see details in ‘Methods’).

## Discussion

We have previously performed three meta-analysis studies involving the reanalysis and integration in Expression Atlas of public quantitative proteomics datasets coming from cell lines and human tumour samples ^8^, from human baseline tissues ^7^, from mouse and rat baseline tissues ^6^. Here we have reanalysed 14 public proteomics datasets coming from pig tissues in baseline conditions. Our overall aim was to provide a system-wide baseline protein expression catalogue across various pig organs. We have used the same methodology as in the study involving baseline human tissues (and in the mouse/rat study), which enabled a comparison of protein expression levels across human and pig organs. To the best of our knowledge this is the first metanalysis study for pig at a protein expression level, in this case using label-free DDA data. The resource PaxDB version 5.0 ^14^ includes pig data generated using spectral counting, but their granularity is limited, providing only organ-specific expression for liver.

As done before, we reanalysed each dataset separately using MaxQuant and the same protein sequence database. The disadvantage of this approach is that the FDR statistical thresholds are applied at a dataset level and not to all datasets together as a whole. However, as also explained before ^6, 7^, using a dataset per dataset analysis approach is in our view the only sustainable manner to reanalyse and integrate quantitative proteomics datasets in resources such as Expression Atlas, where gene expression datasets are stored following the same dataset per dataset approach. It is also important to highlight that the number of commonly detected protein false positives is reduced in parallel with the increase in the number of common datasets where a given protein is detected. In this case, for proteins detected in four or more datasets, a protein FDR of less than 1% can be inferred (Figure S1 in Supporting File 1).

This overall study of protein expression in pig and its comparison with human protein expression is relevant in different contexts. First of all, systems biology research of the domestic pig is of immediate relevance for food production and animal welfare. Additionally, the domestic pig is a model organism for human biomedical research. Furthermore, the diversity of available pig models is rapidly expanding. Some possible applications of these models are research in nutrition, inflammation, and host–microbial crosstalk. In this context, pig models present great opportunities, because the pig, like humans, is an omnivore, with very similar nutritional requirements, digestive and immune systems, and gut microbial components ^12^. It is also important to highlight that mini-pig’s models are being increasingly used in drug development. Animals are still requested in the safety testing of new drug candidates, and minipigs are a potential non-rodent alternative to the use of non-human-primates (NHP), due to ethical considerations ^42^. The use of alternative *in vitro* models is still challenging due to complex biological responses in various organ systems following drug-treatment. Therefore, it is important to have access to protein expression information in pig organs (and also ideally in mini-pig, there are still very few mini-pig proteomics studies in the public domain), so that comparisons in protein abundance across different organs and species (especially between pig, mini-pig and human) can be performed.

Future directions in analogous studies will involve: (i) the inclusion of additional species, e.g. other model organisms or other species of economic importance; (ii) studies focused on particular diseases or physiological states; (iii) the inclusion of differential proteomics datasets in addition to baseline studies; and (iv) reanalysis of DIA datasets (e.g. ^43^). In conclusion we here present a meta-analysis study of public pig baseline proteomics datasets from the PRIDE database. We also performed a comparative analysis across human and pig protein abundance. The resulting protein expression data has been made available *via* Expression Atlas.

### Data availability

Expression Atlas E-PROT identifiers and the PRIDE original dataset identifiers used for the reanalysis are included in Table 1.

## Acknowledgements

We would like to thank all data submitters who made their datasets available via PRIDE and ProteomeXchange. This work has been funded by Open Targets, BBSRC [BB/T019670/1 and BB/T019557/1] and EMBL core funding. We would also like to thank Andrew Leach and the rest of the team involved in the Open Targets “Target Safety” project, for helpful discussions.

## Supporting information

- Supporting File 1:
  - Figure S1: Distribution of common reverse decoy hits across the number of datasets.
  - Figure S2: Organ specificity of canonical proteins in pig.
  - Figure S3: Correlation between gene (RNA-seq based) and protein expression in baseline tissue pig datasets.
  - Figure S4: Correlation of protein abundances across organ liver compared between PaxDB and the results found in this study.
- Supporting File 2:
  - Table S1: Median protein abundances (in ppb) for each protein group across various tissue samples included in each organ.
  - Table S2: Median binned protein abundances across various tissue samples in each pig organ.
  - Table S3: Median binned protein abundances across various pig datasets.
  - Table S4: Organ distribution of canonical proteins in pig.
  - Table S5: Gene Ontology enrichment analysis of ‘organ-enriched’ and ‘group-enriched’ proteins.
  - Table S6: Median binned protein abundances of human and pig orthologs across all organs.
  - Table S7: Elevated proteomes of three different groups in various organs after applying edit distance between human and pig homologous genes.
  - Table S8: Gene Ontology enrichment analysis of the three different groups of proteins, considering their level of expression in pig and human tissues.
  - Table S9: Protein abundances (ppb) considering only one to one mapping between human and pig orthologs across all organs.
- Supporting File 3:
  - Figure S1: Figure illustrating the binned protein abundances of all one-to-one mapped orthologs across ten common organs in human and pig.

## Abbreviations

DDA: Data Dependent Acquisition
DIA: Data Independent Acquisition
FDR: False Discovery Rate
GO: Gene Ontology
iBAQ: intensity-based absolute quantification
IDF: Investigation Description Format
MS: Mass Spectrometry
NHP: Non-Human Primate
ppb: parts per billion
PCA: Principal Component Analysis
SDRF: Sample and Data Relationship Format
UMAP: Uniform Manifold Approximation and Projection

