## Supporting File 1 for "Integrated Proteomics analysis of baseline protein expression in pig tissues"

TABLE OF CONTENT

S-2 Table of Content
S-3 Figure S1. Distribution of common reverse decoy hits across the number of datasets.
S-4 Figure S2. Organ specificity of canonical proteins in pig.
S-5 Figure S3. Correlation between gene (RNA-seq based) and protein expression in baseline tissue pig datasets.
S-6 Figure S4. Correlation of protein abundances across organ liver compared between PaxDB and this study.

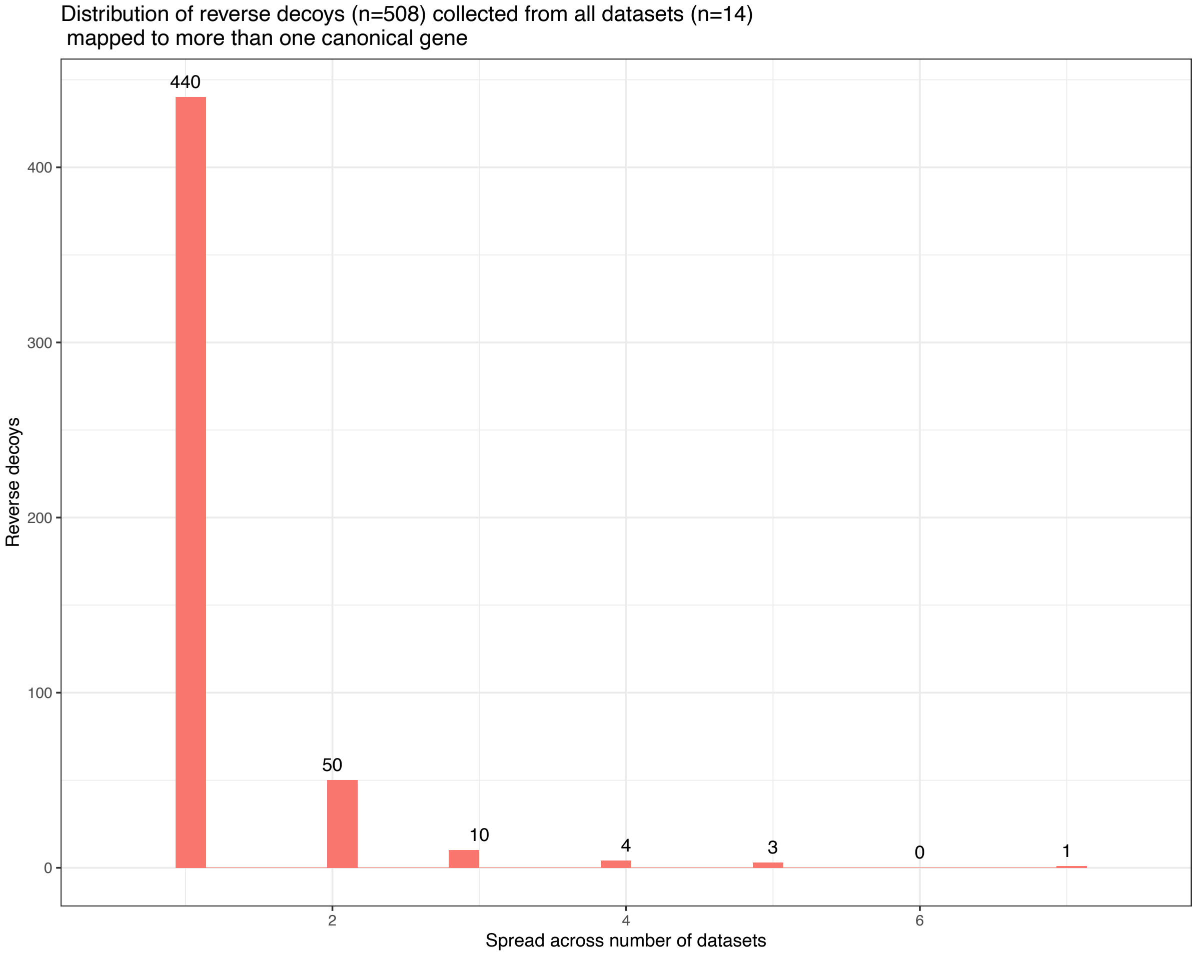


**Figure S1.** Distribution of common reverse decoy hits across the number of datasets.

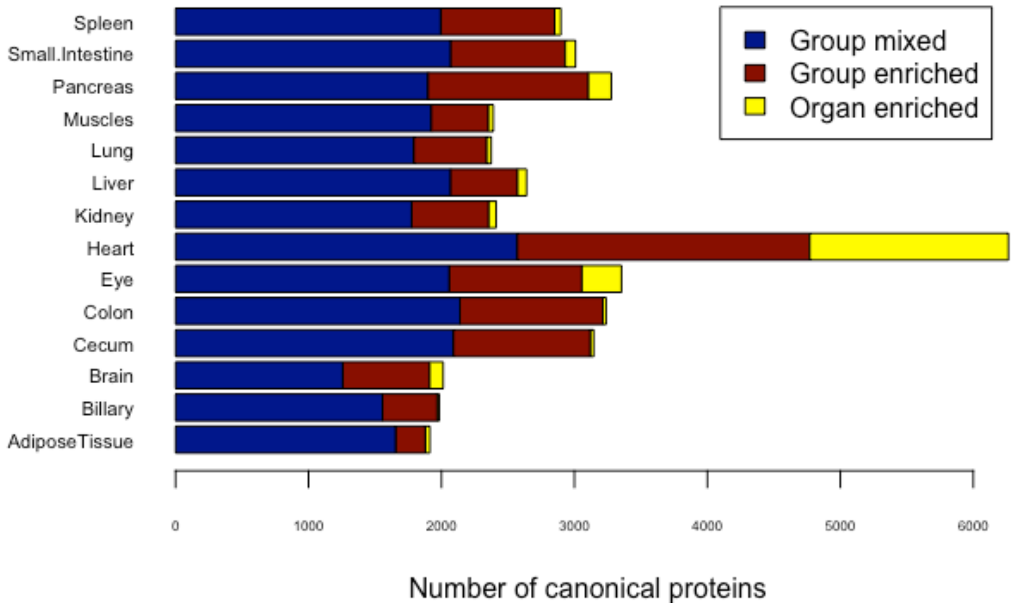


**Figure S2.** Organ specificity of canonical proteins in pig.

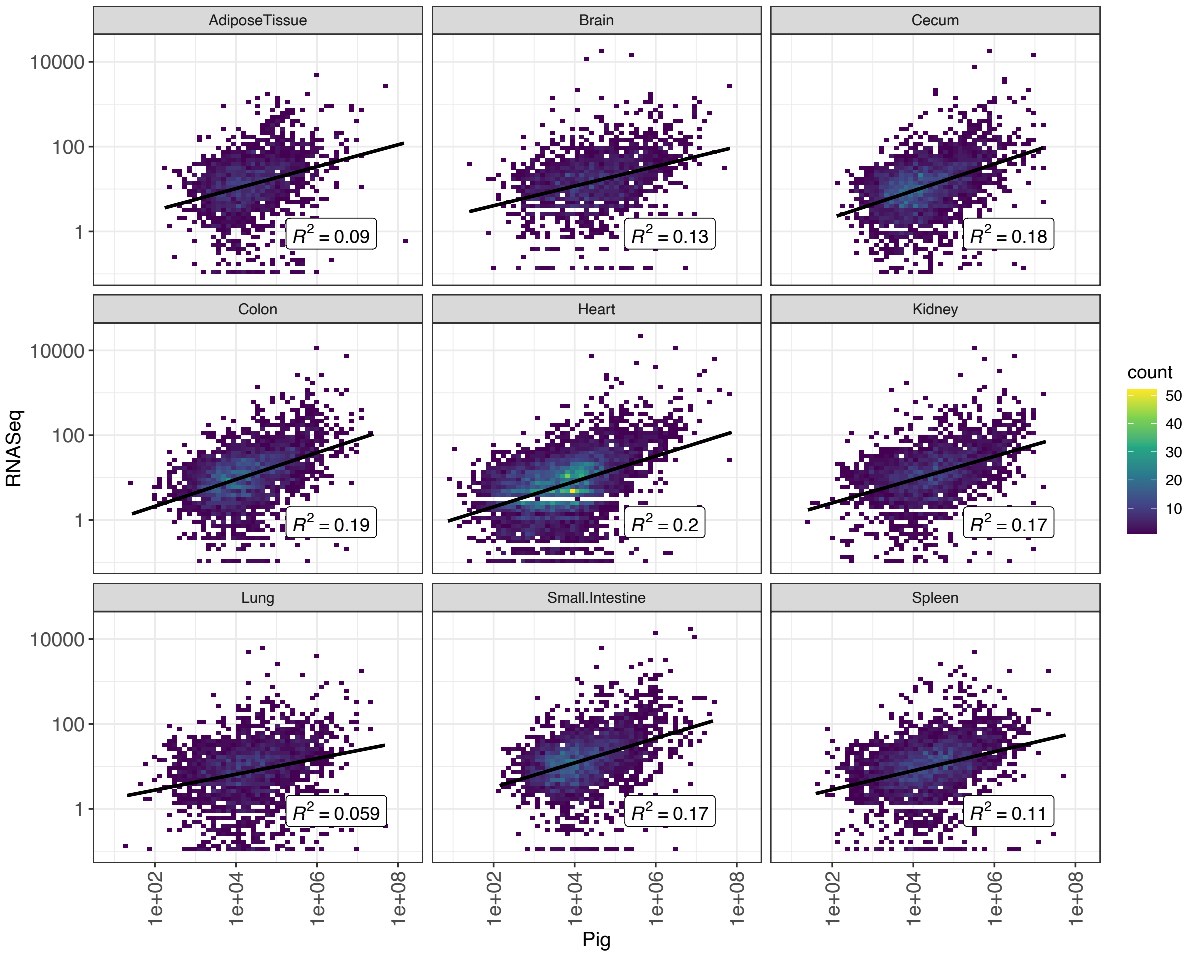


Figure S3. Correlation between gene (RNA-seq based) and protein expression in baseline tissue pig datasets.

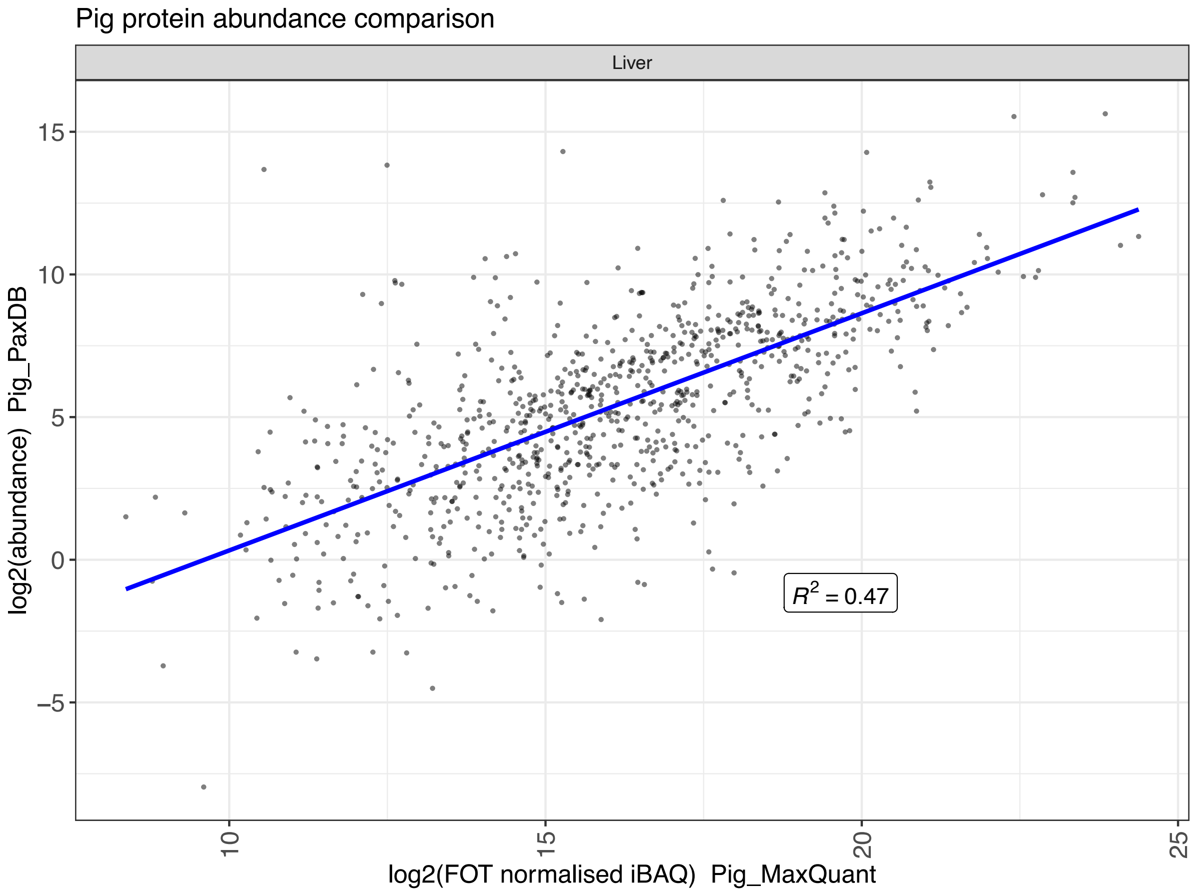


Figure S4. Correlation of protein abundances across organ liver compared between PaxDB and this study.

**Supporting File 2:**

**Table S1:** Median protein abundances (in ppb) for each protein group across various tissue samples included in each organ.
**Table S2:** Median binned protein abundances across various tissue samples in each organ of the pig.
**Table S3:** Median binned protein abundances across various pig datasets.
**Table S4:** Organ distribution of canonical proteins in pig.
**Table S5:** Gene Ontology enrichment analysis of 'organ-enriched' and 'group-enriched' proteins.
**Table S6:** Median binned protein abundances of human and pig orthologs across all organs.
**Table S7:** Elevated proteomes of three different groups in various organs after applying edit distance between human and pig homologous genes.
**Table S8:** Gene Ontology enrichment analysis of the three different groups of proteins, considering their level of expression in pig and human tissues.
**Table S9:** Protein abundances (ppb) with one-to-one mapping between human and pig orthologs across all organs.

**Supporting File 3:**

**Figure S1:** Figure illustrating the binned protein abundances of all one-to-one mapped orthologs across ten common organs in human and pig.
