## Supporting File 3 for "Integrated Proteomics analysis of baseline protein expression in pig tissues"

**Figure S1. Figure illustrating the binned protein abundances of all one-to-one mapped orthologs across ten common organs in human and pig.**

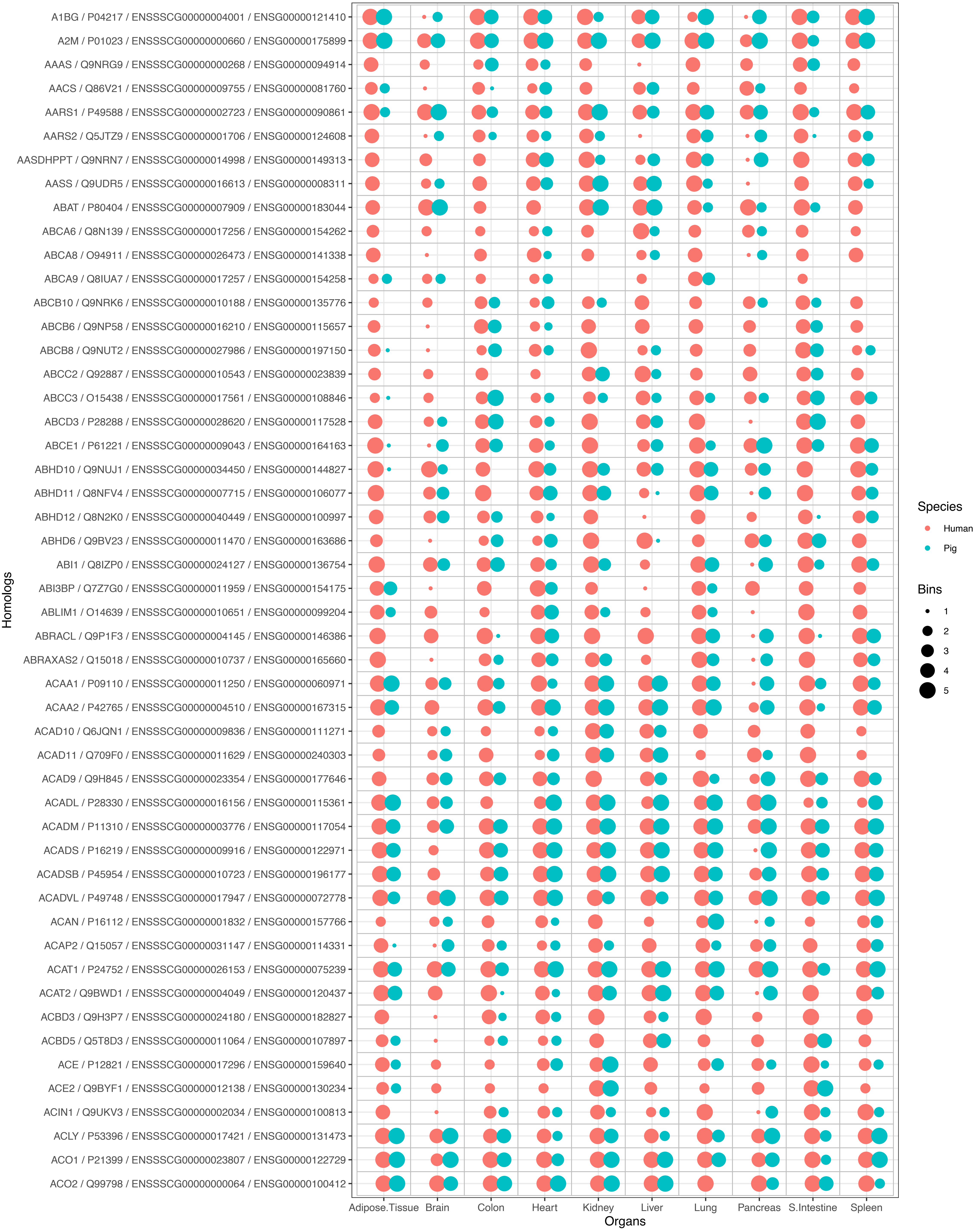
A1BG / P04217 / ENSSSCG00000004001 / ENSG00000121410
A2M / P01023 / ENSSSCG00000000660 / ENSG00000175899
AAAS / Q9NRG9 / ENSSSCG00000000268 / ENSG00000094914
AACS / Q86V21 / ENSSSCG00000009755 / ENSG00000081760
AARS1 / P49588 / ENSSSCG00000002723 / ENSG00000090861
AARS2 / Q5JTZ9 / ENSSSCG00000001706 / ENSG00000124608
AASDHPPT / Q9NRN7 / ENSSSCG00000014998 / ENSG00000149313
AASS / Q9UDR5 / ENSSSCG00000016613 / ENSG00000008311
ABAT / P80404 / ENSSSCG00000007909 / ENSG00000183044
ABCA6 / Q8N139 / ENSSSCG00000017256 / ENSG00000154262
ABCA8 / O94911 / ENSSSCG00000026473 / ENSG00000141338
ABCA9 / Q8IUA7 / ENSSSCG00000017257 / ENSG00000154258
ABCB10 / Q9NRK6 / ENSSSCG00000010188 / ENSG00000135776
ABCB6 / Q9NP58 / ENSSSCG00000016210 / ENSG00000115657
ABCB8 / Q9NUT2 / ENSSSCG00000027986 / ENSG00000197150
ABCC2 / Q92887 / ENSSSCG00000010543 / ENSG00000023839
ABCC3 / O15438 / ENSSSCG00000017561 / ENSG00000108846
ABCD3 / P28288 / ENSSSCG00000028620 / ENSG00000117528
ABCE1 / P61221 / ENSSSCG00000009043 / ENSG00000164163
ABHD10 / Q9NUJ1 / ENSSSCG00000034450 / ENSG00000144827
ABHD11 / Q8NFV4 / ENSSSCG00000007715 / ENSG00000106077
ABHD12 / Q8N2K0 / ENSSSCG00000040449 / ENSG00000100997
ABHD6 / Q9BV23 / ENSSSCG00000011470 / ENSG00000163686
ABI1 / Q8IZP0 / ENSSSCG00000024127 / ENSG00000136754
ABI3BP / Q7Z7G0 / ENSSSCG00000011959 / ENSG00000154175
ABLIM1 / O14639 / ENSSSCG00000010651 / ENSG00000099204
ABRACL / Q9P1F3 / ENSSSCG00000004145 / ENSG00000146386
ABRAXAS2 / Q15018 / ENSSSCG00000010737 / ENSG00000165660
ACAA1 / P09110 / ENSSSCG00000011250 / ENSG00000060971
ACAA2 / P42765 / ENSSSCG00000004510 / ENSG00000167315
ACAD10 / Q6JQN1 / ENSSSCG00000009836 / ENSG00000111271
ACAD11 / Q709F0 / ENSSSCG00000011629 / ENSG00000240303
ACAD9 / Q9H845 / ENSSSCG00000023354 / ENSG00000177646
ACADL / P28330 / ENSSSCG00000016156 / ENSG00000115361
ACADM / P11310 / ENSSSCG00000003776 / ENSG00000117054
ACADS / P16219 / ENSSSCG00000009916 / ENSG00000122971
ACADSB / P45954 / ENSSSCG00000010723 / ENSG00000196177
ACADVL / P49748 / ENSSSCG00000017947 / ENSG00000072778
ACAN / P16112 / ENSSSCG00000001832 / ENSG00000157766
ACAP2 / Q15057 / ENSSSCG00000031147 / ENSG00000114331
ACAT1 / P24752 / ENSSSCG00000026153 / ENSG00000075239
ACAT2 / Q9BWD1 / ENSSSCG00000004049 / ENSG00000120437
ACBD3 / Q9H3P7 / ENSSSCG00000024180 / ENSG00000182827
ACBD5 / Q5T8D3 / ENSSSCG00000011064 / ENSG00000107897
ACE / P12821 / ENSSSCG00000017296 / ENSG00000159640
ACE2 / Q9BYF1 / ENSSSCG00000012138 / ENSG00000130234
ACIN1 / Q9UKV3 / ENSSSCG00000002034 / ENSG00000100813
ACLY / P53396 / ENSSSCG00000017421 / ENSG00000131473
ACO1 / P21399 / ENSSSCG00000023807 / ENSG00000122729
ACO2 / Q99798 / ENSSSCG00000000064 / ENSG00000100412
Homologs
Adipose.Tissue
Brain
Colon
Heart
Kidney
Liver
Lung
Pancreas
S.Intestine
Spleen
Organs
Species
Human
Pig
Bins
1
2
3
4
5

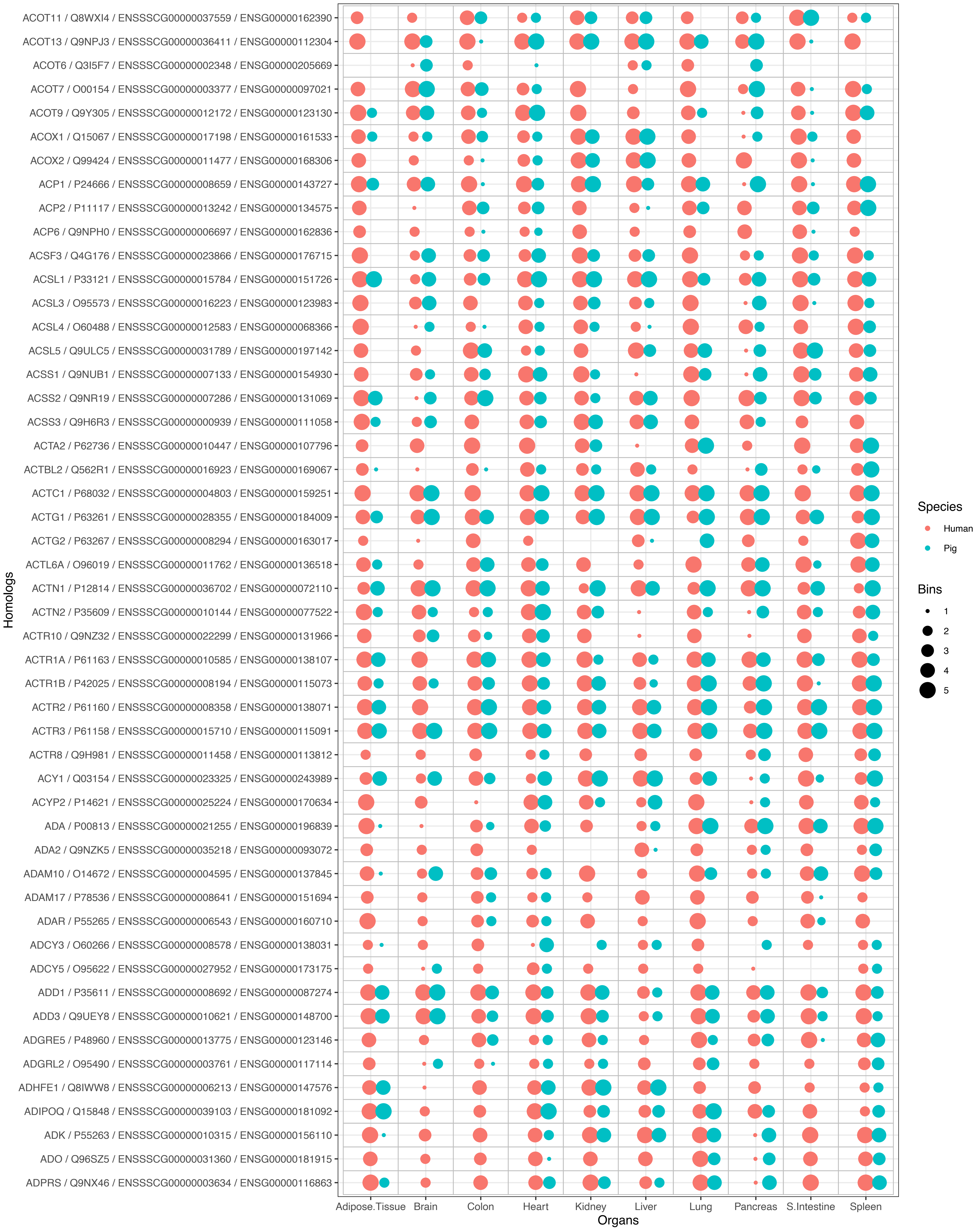
ACOT11 / Q8WXI4 / ENSSSCG00000037559 / ENSG00000162390
ACOT13 / Q9NPJ3 / ENSSSCG00000036411 / ENSG00000112304
ACOT6 / Q3I5F7 / ENSSSCG00000002348 / ENSG00000205669
ACOT7 / O00154 / ENSSSCG00000003377 / ENSG00000097021
ACOT9 / Q9Y305 / ENSSSCG00000012172 / ENSG00000123130
ACOX1 / Q15067 / ENSSSCG00000017198 / ENSG00000161533
ACOX2 / Q99424 / ENSSSCG00000011477 / ENSG00000168306
ACP1 / P24666 / ENSSSCG00000008659 / ENSG00000143727
ACP2 / P11117 / ENSSSCG00000013242 / ENSG00000134575
ACP6 / Q9NPH0 / ENSSSCG00000006697 / ENSG00000162836
ACSF3 / Q4G176 / ENSSSCG00000023866 / ENSG00000176715
ACSL1 / P33121 / ENSSSCG00000015784 / ENSG00000151726
ACSL3 / O95573 / ENSSSCG00000016223 / ENSG00000123983
ACSL4 / O60488 / ENSSSCG00000012583 / ENSG00000068366
ACSL5 / Q9ULC5 / ENSSSCG00000031789 / ENSG00000197142
ACSS1 / Q9NUB1 / ENSSSCG00000007133 / ENSG00000154930
ACSS2 / Q9NR19 / ENSSSCG00000007286 / ENSG00000131069
ACSS3 / Q9H6R3 / ENSSSCG00000000939 / ENSG00000111058
ACTA2 / P62736 / ENSSSCG00000010447 / ENSG00000107796
ACTBL2 / Q562R1 / ENSSSCG00000016923 / ENSG00000169067
ACTC1 / P68032 / ENSSSCG00000004803 / ENSG00000159251
ACTG1 / P63261 / ENSSSCG00000028355 / ENSG00000184009
ACTG2 / P63267 / ENSSSCG00000008294 / ENSG00000163017
ACTL6A / O96019 / ENSSSCG00000011762 / ENSG00000136518
ACTN1 / P12814 / ENSSSCG00000036702 / ENSG00000072110
ACTN2 / P35609 / ENSSSCG00000010144 / ENSG00000077522
ACTR10 / Q9NZ32 / ENSSSCG00000022299 / ENSG00000131966
ACTR1A / P61163 / ENSSSCG00000010585 / ENSG00000138107
ACTR1B / P42025 / ENSSSCG00000008194 / ENSG00000115073
ACTR2 / P61160 / ENSSSCG00000008358 / ENSG00000138071
ACTR3 / P61158 / ENSSSCG00000015710 / ENSG00000115091
ACTR8 / Q9H981 / ENSSSCG00000011458 / ENSG00000113812
ACY1 / Q03154 / ENSSSCG00000023325 / ENSG00000243989
ACYP2 / P14621 / ENSSSCG00000025224 / ENSG00000170634
ADA / P00813 / ENSSSCG00000021255 / ENSG00000196839
ADA2 / Q9NZK5 / ENSSSCG00000035218 / ENSG00000093072
ADAM10 / O14672 / ENSSSCG00000004595 / ENSG00000137845
ADAM17 / P78536 / ENSSSCG00000008641 / ENSG00000151694
ADAR / P55265 / ENSSSCG00000006543 / ENSG00000160710
ADCY3 / O60266 / ENSSSCG00000008578 / ENSG00000138031
ADCY5 / O95622 / ENSSSCG00000027952 / ENSG00000173175
ADD1 / P35611 / ENSSSCG00000008692 / ENSG00000087274
ADD3 / Q9UEY8 / ENSSSCG00000010621 / ENSG00000148700
ADGRE5 / P48960 / ENSSSCG00000013775 / ENSG00000123146
ADGRL2 / O95490 / ENSSSCG00000003761 / ENSG00000117114
ADHFE1 / Q8IWW8 / ENSSSCG00000006213 / ENSG00000147576
ADIPOQ / Q15848 / ENSSSCG00000039103 / ENSG00000181092
ADK / P55263 / ENSSSCG00000010315 / ENSG00000156110
ADO / Q96SZ5 / ENSSSCG00000031360 / ENSG00000181915
ADPRS / Q9NX46 / ENSSSCG00000003634 / ENSG00000116863
Homologs
Adipose.Tissue
Brain
Colon
Heart
Kidney
Liver
Lung
Pancreas
S.Intestine
Spleen
Organs
Species
Human
Pig
Bins
1
2
3
4
5

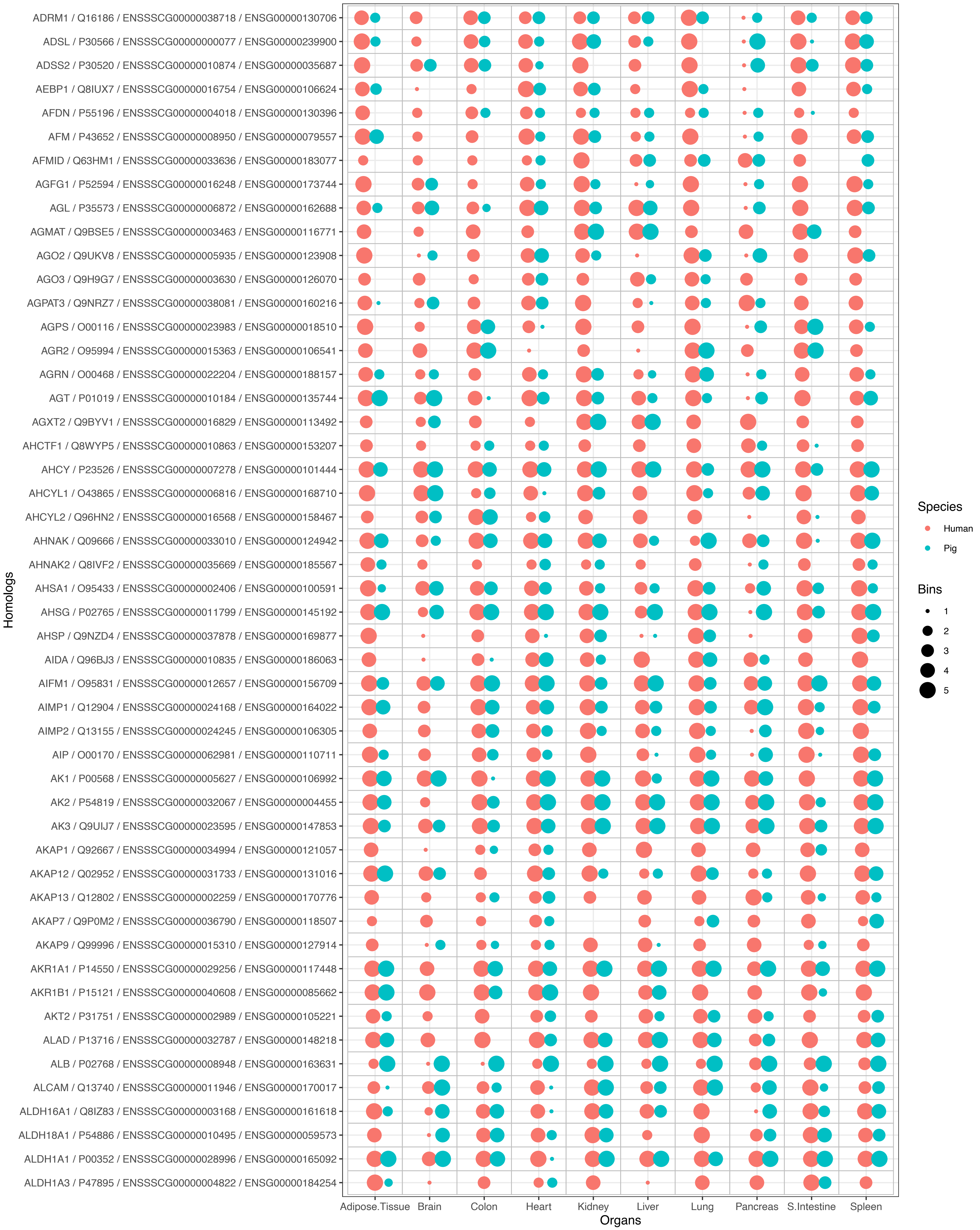

ADRM1 / Q16186 / ENSSSCG00000038718 / ENSG00000130706
ADSL / P30566 / ENSSSCG00000000077 / ENSG00000239900
ADSS2 / P30520 / ENSSSCG00000010874 / ENSG00000035687
AEBP1 / Q8IUX7 / ENSSSCG00000016754 / ENSG00000106624
AFDN / P55196 / ENSSSCG00000004018 / ENSG00000130396
AFM / P43652 / ENSSSCG00000008950 / ENSG00000079557
AFMID / Q63HM1 / ENSSSCG00000033636 / ENSG00000183077
AGFG1 / P52594 / ENSSSCG00000016248 / ENSG00000173744
AGL / P35573 / ENSSSCG00000006872 / ENSG00000162688
AGMAT / Q9BSE5 / ENSSSCG00000003463 / ENSG00000116771
AGO2 / Q9UKV8 / ENSSSCG00000005935 / ENSG00000123908
AGO3 / Q9H9G7 / ENSSSCG00000003630 / ENSG00000126070
AGPAT3 / Q9NRZ7 / ENSSSCG00000038081 / ENSG00000160216
AGPS / O00116 / ENSSSCG00000023983 / ENSG00000018510
AGR2 / O95994 / ENSSSCG00000015363 / ENSG00000106541
AGRN / O00468 / ENSSSCG00000022204 / ENSG00000188157
AGT / P01019 / ENSSSCG00000010184 / ENSG00000135744
AGXT2 / Q9BYV1 / ENSSSCG00000016829 / ENSG00000113492
AHCTF1 / Q8WYP5 / ENSSSCG00000010863 / ENSG00000153207
AHCY / P23526 / ENSSSCG00000007278 / ENSG00000101444
AHCYL1 / O43865 / ENSSSCG00000006816 / ENSG00000168710
AHCYL2 / Q96HN2 / ENSSSCG00000016568 / ENSG00000158467
AHNAK / Q09666 / ENSSSCG00000033010 / ENSG00000124942
AHNAK2 / Q8IVF2 / ENSSSCG00000035669 / ENSG00000185567
AHSA1 / O95433 / ENSSSCG00000002406 / ENSG00000100591
AHSG / P02765 / ENSSSCG00000011799 / ENSG00000145192
AHSP / Q9NZD4 / ENSSSCG00000037878 / ENSG00000169877
AIDA / Q96BJ3 / ENSSSCG00000010835 / ENSG00000186063
AIFM1 / O95831 / ENSSSCG00000012657 / ENSG00000156709
AIMP1 / Q12904 / ENSSSCG00000024168 / ENSG00000164022
AIMP2 / Q13155 / ENSSSCG00000024245 / ENSG00000106305
AIP / O00170 / ENSSSCG00000062981 / ENSG00000110711
AK1 / P00568 / ENSSSCG00000005627 / ENSG00000106992
AK2 / P54819 / ENSSSCG00000032067 / ENSG00000004455
AK3 / Q9UIJ7 / ENSSSCG00000023595 / ENSG00000147853
AKAP1 / Q92667 / ENSSSCG00000034994 / ENSG00000121057
AKAP12 / Q02952 / ENSSSCG00000031733 / ENSG00000131016
AKAP13 / Q12802 / ENSSSCG00000002259 / ENSG00000170776
AKAP7 / Q9P0M2 / ENSSSCG00000036790 / ENSG00000118507
AKAP9 / Q99996 / ENSSSCG00000015310 / ENSG00000127914
AKR1A1 / P14550 / ENSSSCG00000029256 / ENSG00000117448
AKR1B1 / P15121 / ENSSSCG00000040608 / ENSG00000085662
AKT2 / P31751 / ENSSSCG00000002989 / ENSG00000105221
ALAD / P13716 / ENSSSCG00000032787 / ENSG00000148218
ALB / P02768 / ENSSSCG00000008948 / ENSG00000163631
ALCAM / Q13740 / ENSSSCG00000011946 / ENSG00000170017
ALDH16A1 / Q8IZ83 / ENSSSCG00000003168 / ENSG00000161618
ALDH18A1 / P54886 / ENSSSCG00000010495 / ENSG00000059573
ALDH1A1 / P00352 / ENSSSCG00000028996 / ENSG00000165092
ALDH1A3 / P47895 / ENSSSCG00000004822 / ENSG00000184254
Homologs
Adipose.Tissue
Brain
Colon
Heart
Kidney
Liver
Lung
Pancreas
S.Intestine
Spleen
Organs
Species
Human
Pig
Bins
1
2
3
4
5

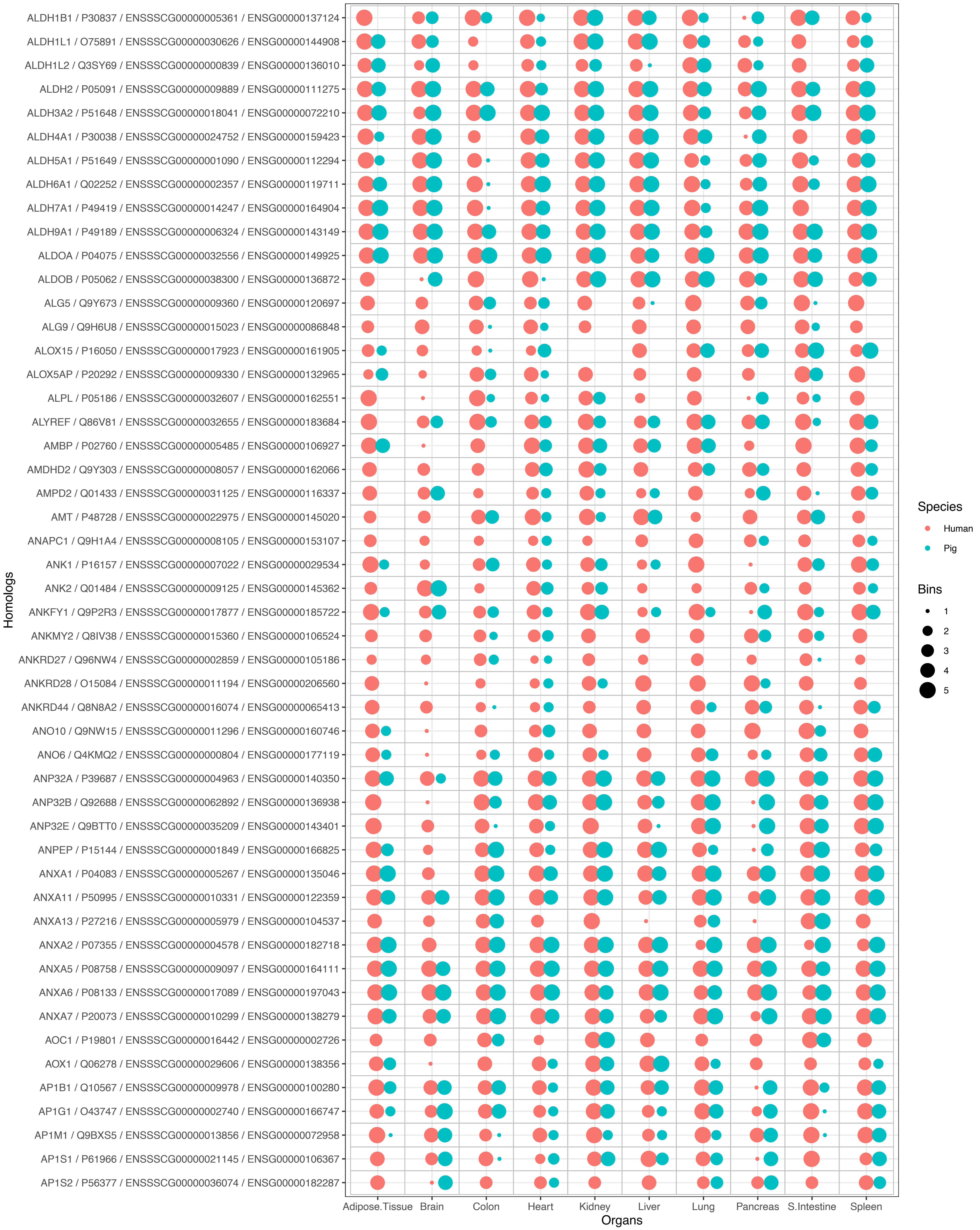

ALDH1B1 / P30837 / ENSSSCG00000005361 / ENSG00000137124
ALDH1L1 / O75891 / ENSSSCG00000030626 / ENSG00000144908
ALDH1L2 / Q3SY69 / ENSSSCG00000000839 / ENSG00000136010
ALDH2 / P05091 / ENSSSCG00000009889 / ENSG00000111275
ALDH3A2 / P51648 / ENSSSCG00000018041 / ENSG00000072210
ALDH4A1 / P30038 / ENSSSCG00000024752 / ENSG00000159423
ALDH5A1 / P51649 / ENSSSCG00000001090 / ENSG00000112294
ALDH6A1 / Q02252 / ENSSSCG00000002357 / ENSG00000119711
ALDH7A1 / P49419 / ENSSSCG00000014247 / ENSG00000164904
ALDH9A1 / P49189 / ENSSSCG00000006324 / ENSG00000143149
ALDOA / P04075 / ENSSSCG00000032556 / ENSG00000149925
ALDOB / P05062 / ENSSSCG00000038300 / ENSG00000136872
ALG5 / Q9Y673 / ENSSSCG00000009360 / ENSG00000120697
ALG9 / Q9H6U8 / ENSSSCG00000015023 / ENSG00000086848
ALOX15 / P16050 / ENSSSCG00000017923 / ENSG00000161905
ALOX5AP / P20292 / ENSSSCG00000009330 / ENSG00000132965
ALPL / P05186 / ENSSSCG00000032607 / ENSG00000162551
ALYREF / Q86V81 / ENSSSCG00000032655 / ENSG00000183684
AMBP / P02760 / ENSSSCG00000005485 / ENSG00000106927
AMDHD2 / Q9Y303 / ENSSSCG00000008057 / ENSG00000162066
AMPD2 / Q01433 / ENSSSCG00000031125 / ENSG00000116337
AMT / P48728 / ENSSSCG00000022975 / ENSG00000145020
ANAPC1 / Q9H1A4 / ENSSSCG00000008105 / ENSG00000153107
ANK1 / P16157 / ENSSSCG00000007022 / ENSG00000029534
ANK2 / Q01484 / ENSSSCG00000009125 / ENSG00000145362
ANKFY1 / Q9P2R3 / ENSSSCG00000017877 / ENSG00000185722
ANKMY2 / Q8IV38 / ENSSSCG00000015360 / ENSG00000106524
ANKRD27 / Q96NW4 / ENSSSCG00000002859 / ENSG00000105186
ANKRD28 / O15084 / ENSSSCG00000011194 / ENSG00000206560
ANKRD44 / Q8N8A2 / ENSSSCG00000016074 / ENSG00000065413
ANO10 / Q9NW15 / ENSSSCG00000011296 / ENSG00000160746
ANO6 / Q4KMQ2 / ENSSSCG00000000804 / ENSG00000177119
ANP32A / P39687 / ENSSSCG00000004963 / ENSG00000140350
ANP32B / Q92688 / ENSSSCG00000062892 / ENSG00000136938
ANP32E / Q9BTT0 / ENSSSCG00000035209 / ENSG00000143401
ANPEP / P15144 / ENSSSCG00000001849 / ENSG00000166825
ANXA1 / P04083 / ENSSSCG00000005267 / ENSG00000135046
ANXA11 / P50995 / ENSSSCG00000010331 / ENSG00000122359
ANXA13 / P27216 / ENSSSCG00000005979 / ENSG00000104537
ANXA2 / P07355 / ENSSSCG00000004578 / ENSG00000182718
ANXA5 / P08758 / ENSSSCG00000009097 / ENSG00000164111
ANXA6 / P08133 / ENSSSCG00000017089 / ENSG00000197043
ANXA7 / P20073 / ENSSSCG00000010299 / ENSG00000138279
AOC1 / P19801 / ENSSSCG00000016442 / ENSG00000002726
AOX1 / Q06278 / ENSSSCG00000029606 / ENSG00000138356
AP1B1 / Q10567 / ENSSSCG00000009978 / ENSG00000100280
AP1G1 / O43747 / ENSSSCG00000002740 / ENSG00000166747
AP1M1 / Q9BXS5 / ENSSSCG00000013856 / ENSG00000072958
AP1S1 / P61966 / ENSSSCG00000021145 / ENSG00000106367
AP1S2 / P56377 / ENSSSCG00000036074 / ENSG00000182287
Homologs
Adipose.Tissue
Brain
Colon
Heart
Kidney
Liver
Lung
Pancreas
S.Intestine
Spleen
Organs
Species
Human
Pig
Bins
1
2
3
4
5

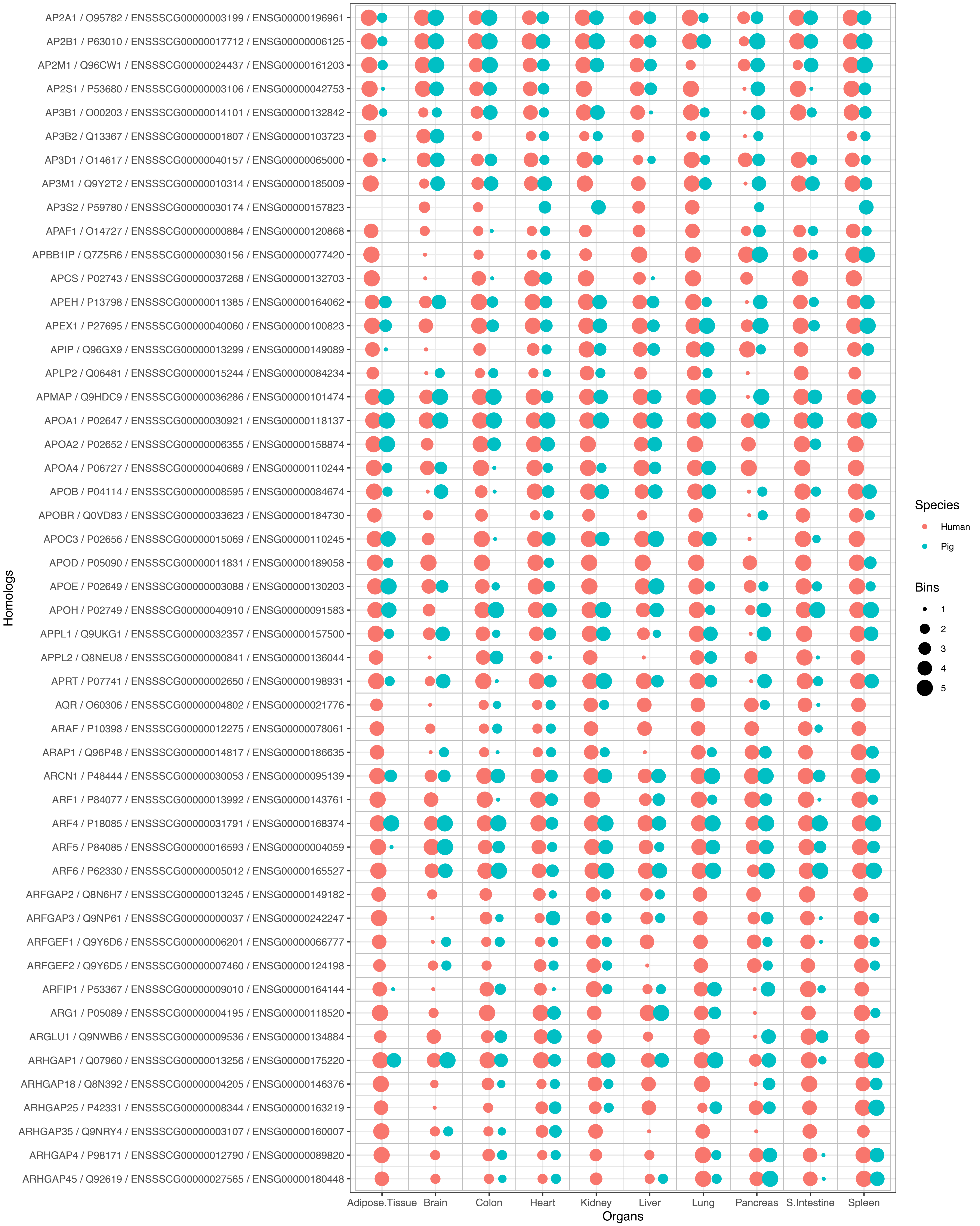

AP2A1 / O95782 / ENSSSCG00000003199 / ENSG00000196961
AP2B1 / P63010 / ENSSSCG00000017712 / ENSG00000006125
AP2M1 / Q96CW1 / ENSSSCG00000024437 / ENSG00000161203
AP2S1 / P53680 / ENSSSCG00000003106 / ENSG00000042753
AP3B1 / O00203 / ENSSSCG00000014101 / ENSG00000132842
AP3B2 / Q13367 / ENSSSCG00000001807 / ENSG00000103723
AP3D1 / O14617 / ENSSSCG00000040157 / ENSG00000065000
AP3M1 / Q9Y2T2 / ENSSSCG00000010314 / ENSG00000185009
AP3S2 / P59780 / ENSSSCG00000030174 / ENSG00000157823
APAF1 / O14727 / ENSSSCG00000000884 / ENSG00000120868
APBB1IP / Q7Z5R6 / ENSSSCG00000030156 / ENSG00000077420
APCS / P02743 / ENSSSCG00000037268 / ENSG00000132703
APEH / P13798 / ENSSSCG00000011385 / ENSG00000164062
APEX1 / P27695 / ENSSSCG00000040060 / ENSG00000100823
APIP / Q96GX9 / ENSSSCG00000013299 / ENSG00000149089
APLP2 / Q06481 / ENSSSCG00000015244 / ENSG00000084234
APMAP / Q9HDC9 / ENSSSCG00000036286 / ENSG00000101474
APOA1 / P02647 / ENSSSCG00000030921 / ENSG00000118137
APOA2 / P02652 / ENSSSCG00000006355 / ENSG00000158874
APOA4 / P06727 / ENSSSCG00000040689 / ENSG00000110244
APOB / P04114 / ENSSSCG00000008595 / ENSG00000084674
APOBR / Q0VD83 / ENSSSCG00000033623 / ENSG00000184730
APOC3 / P02656 / ENSSSCG00000015069 / ENSG00000110245
APOD / P05090 / ENSSSCG00000011831 / ENSG00000189058
APOE / P02649 / ENSSSCG00000003088 / ENSG00000130203
APOH / P02749 / ENSSSCG00000040910 / ENSG00000091583
APPL1 / Q9UKG1 / ENSSSCG00000032357 / ENSG00000157500
APPL2 / Q8NEU8 / ENSSSCG00000000841 / ENSG00000136044
APRT / P07741 / ENSSSCG00000002650 / ENSG00000198931
AQR / O60306 / ENSSSCG00000004802 / ENSG00000021776
ARAF / P10398 / ENSSSCG00000012275 / ENSG00000078061
ARAP1 / Q96P48 / ENSSSCG00000014817 / ENSG00000186635
ARCN1 / P48444 / ENSSSCG00000030053 / ENSG00000095139
ARF1 / P84077 / ENSSSCG00000013992 / ENSG00000143761
ARF4 / P18085 / ENSSSCG00000031791 / ENSG00000168374
ARF5 / P84085 / ENSSSCG00000016593 / ENSG00000004059
ARF6 / P62330 / ENSSSCG00000005012 / ENSG00000165527
ARFGAP2 / Q8N6H7 / ENSSSCG00000013245 / ENSG00000149182
ARFGAP3 / Q9NP61 / ENSSSCG00000000037 / ENSG00000242247
ARFGEF1 / Q9Y6D6 / ENSSSCG00000006201 / ENSG00000066777
ARFGEF2 / Q9Y6D5 / ENSSSCG00000007460 / ENSG00000124198
ARFIP1 / P53367 / ENSSSCG00000009010 / ENSG00000164144
ARG1 / P05089 / ENSSSCG00000004195 / ENSG00000118520
ARGLU1 / Q9NWB6 / ENSSSCG00000009536 / ENSG00000134884
ARHGAP1 / Q07960 / ENSSSCG00000013256 / ENSG00000175220
ARHGAP18 / Q8N392 / ENSSSCG00000004205 / ENSG00000146376
ARHGAP25 / P42331 / ENSSSCG00000008344 / ENSG00000163219
ARHGAP35 / Q9NRY4 / ENSSSCG00000003107 / ENSG00000160007
ARHGAP4 / P98171 / ENSSSCG00000012790 / ENSG00000089820
ARHGAP45 / Q92619 / ENSSSCG00000027565 / ENSG00000180448
Homologs
Adipose.Tissue
Brain
Colon
Heart
Kidney
Liver
Lung
Pancreas
S.Intestine
Spleen
Organs
Species
Human
Pig
Bins
1
2
3
4
5

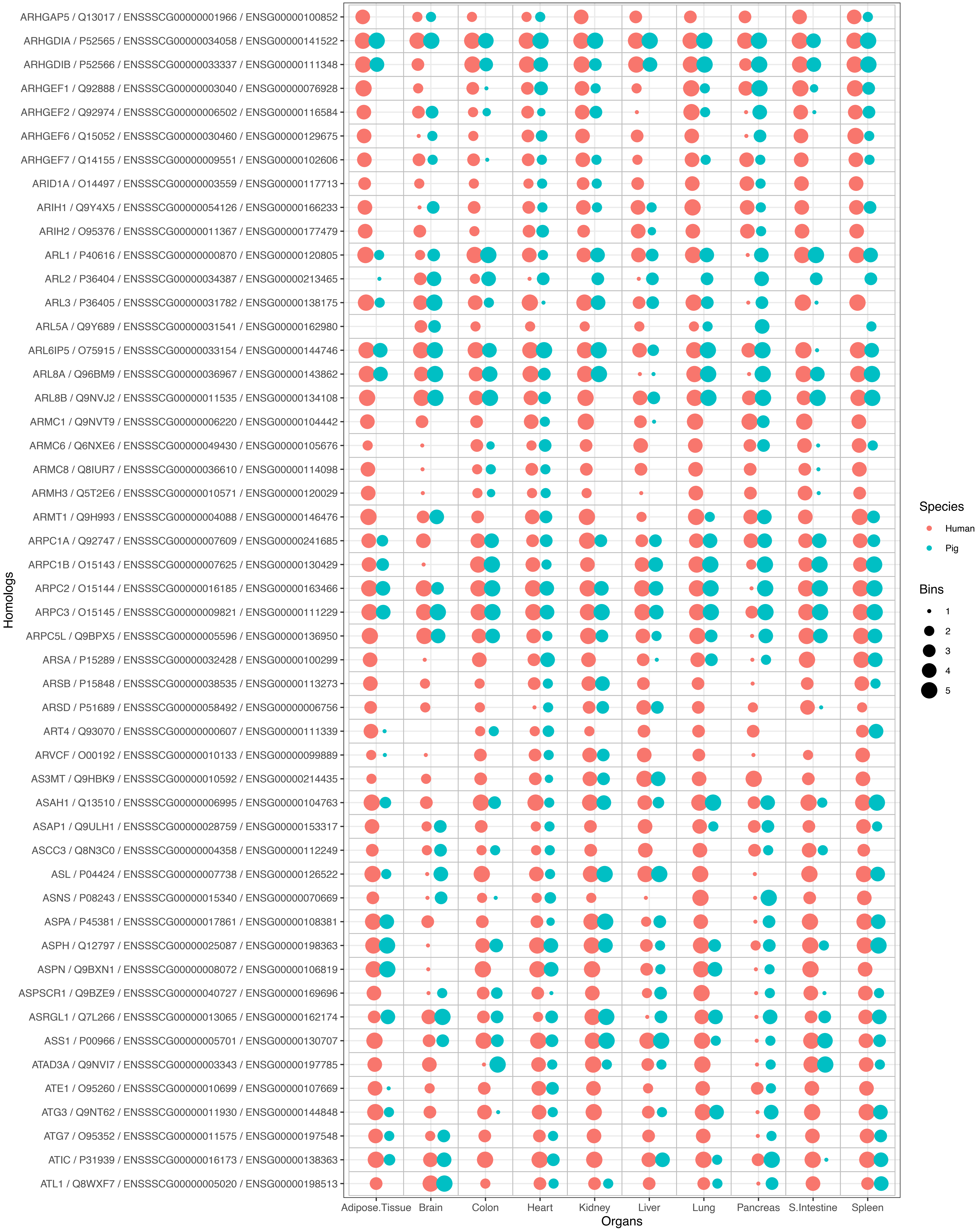
ARHGAP5 / Q13017 / ENSSSCG00000001966 / ENSG00000100852
ARHGDIA / P52565 / ENSSSCG00000034058 / ENSG00000141522
ARHGDIB / P52566 / ENSSSCG00000033337 / ENSG00000111348
ARHGEF1 / Q92888 / ENSSSCG00000003040 / ENSG00000076928
ARHGEF2 / Q92974 / ENSSSCG00000006502 / ENSG00000116584
ARHGEF6 / Q15052 / ENSSSCG00000030460 / ENSG00000129675
ARHGEF7 / Q14155 / ENSSSCG00000009551 / ENSG00000102606
ARID1A / O14497 / ENSSSCG00000003559 / ENSG00000117713
ARIH1 / Q9Y4X5 / ENSSSCG00000054126 / ENSG00000166233
ARIH2 / O95376 / ENSSSCG00000011367 / ENSG00000177479
ARL1 / P40616 / ENSSSCG00000000870 / ENSG00000120805
ARL2 / P36404 / ENSSSCG00000034387 / ENSG00000213465
ARL3 / P36405 / ENSSSCG00000031782 / ENSG00000138175
ARL5A / Q9Y689 / ENSSSCG00000031541 / ENSG00000162980
ARL6IP5 / O75915 / ENSSSCG00000033154 / ENSG00000144746
ARL8A / Q96BM9 / ENSSSCG00000036967 / ENSG00000143862
ARL8B / Q9NVJ2 / ENSSSCG00000011535 / ENSG00000134108
ARMC1 / Q9NVT9 / ENSSSCG00000006220 / ENSG00000104442
ARMC6 / Q6NXE6 / ENSSSCG00000049430 / ENSG00000105676
ARMC8 / Q8IUR7 / ENSSSCG00000036610 / ENSG00000114098
ARMH3 / Q5T2E6 / ENSSSCG00000010571 / ENSG00000120029
ARMT1 / Q9H993 / ENSSSCG00000004088 / ENSG00000146476
ARPC1A / Q92747 / ENSSSCG00000007609 / ENSG00000241685
ARPC1B / O15143 / ENSSSCG00000007625 / ENSG00000130429
ARPC2 / O15144 / ENSSSCG00000016185 / ENSG00000163466
ARPC3 / O15145 / ENSSSCG00000009821 / ENSG00000111229
ARPC5L / Q9BPX5 / ENSSSCG00000005596 / ENSG00000136950
ARSA / P15289 / ENSSSCG00000032428 / ENSG00000100299
ARSB / P15848 / ENSSSCG00000038535 / ENSG00000113273
ARSD / P51689 / ENSSSCG00000058492 / ENSG00000006756
ART4 / Q93070 / ENSSSCG00000000607 / ENSG00000111339
ARVCF / O00192 / ENSSSCG00000010133 / ENSG00000099889
AS3MT / Q9HBK9 / ENSSSCG00000010592 / ENSG00000214435
ASAH1 / Q13510 / ENSSSCG00000006995 / ENSG00000104763
ASAP1 / Q9ULH1 / ENSSSCG00000028759 / ENSG00000153317
ASCC3 / Q8N3C0 / ENSSSCG00000004358 / ENSG00000112249
ASL / P04424 / ENSSSCG00000007738 / ENSG00000126522
ASNS / P08243 / ENSSSCG00000015340 / ENSG00000070669
ASPA / P45381 / ENSSSCG00000017861 / ENSG00000108381
ASPH / Q12797 / ENSSSCG00000025087 / ENSG00000198363
ASPN / Q9BXN1 / ENSSSCG00000008072 / ENSG00000106819
ASPSCR1 / Q9BZE9 / ENSSSCG00000040727 / ENSG00000169696
ASRGL1 / Q7L266 / ENSSSCG00000013065 / ENSG00000162174
ASS1 / P00966 / ENSSSCG00000005701 / ENSG00000130707
ATAD3A / Q9NVI7 / ENSSSCG00000003343 / ENSG00000197785
ATE1 / O95260 / ENSSSCG00000010699 / ENSG00000107669
ATG3 / Q9NT62 / ENSSSCG00000011930 / ENSG00000144848
ATG7 / O95352 / ENSSSCG00000011575 / ENSG00000197548
ATIC / P31939 / ENSSSCG00000016173 / ENSG00000138363
ATL1 / Q8WXF7 / ENSSSCG00000005020 / ENSG00000198513
Homologs
Adipose.Tissue
Brain
Colon
Heart
Kidney
Liver
Lung
Pancreas
S.Intestine
Spleen
Organs
Species
Human
Pig
Bins
1
2
3
4
5

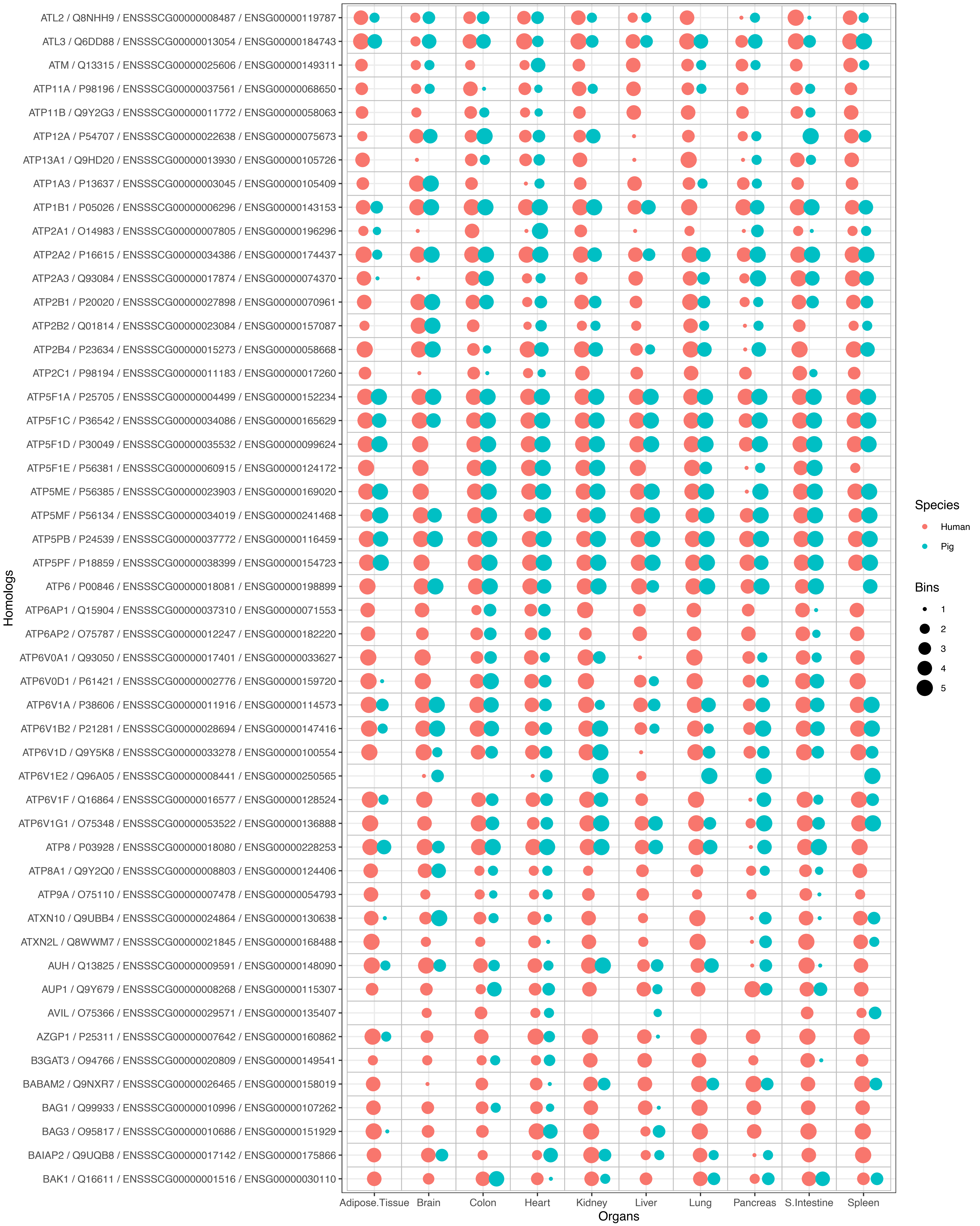
ATL2 / Q8NHH9 / ENSSSCG00000008487 / ENSG00000119787
ATL3 / Q6DD88 / ENSSSCG00000013054 / ENSG00000184743
ATM / Q13315 / ENSSSCG00000025606 / ENSG00000149311
ATP11A / P98196 / ENSSSCG00000037561 / ENSG00000068650
ATP11B / Q9Y2G3 / ENSSSCG00000011772 / ENSG00000058063
ATP12A / P54707 / ENSSSCG00000022638 / ENSG00000075673
ATP13A1 / Q9HD20 / ENSSSCG00000013930 / ENSG00000105726
ATP1A3 / P13637 / ENSSSCG00000003045 / ENSG00000105409
ATP1B1 / P05026 / ENSSSCG00000006296 / ENSG00000143153
ATP2A1 / O14983 / ENSSSCG00000007805 / ENSG00000196296
ATP2A2 / P16615 / ENSSSCG00000034386 / ENSG00000174437
ATP2A3 / Q93084 / ENSSSCG00000017874 / ENSG00000074370
ATP2B1 / P20020 / ENSSSCG00000027898 / ENSG00000070961
ATP2B2 / Q01814 / ENSSSCG00000023084 / ENSG00000157087
ATP2B4 / P23634 / ENSSSCG00000015273 / ENSG00000058668
ATP2C1 / P98194 / ENSSSCG00000011183 / ENSG00000017260
ATP5F1A / P25705 / ENSSSCG00000004499 / ENSG00000152234
ATP5F1C / P36542 / ENSSSCG00000034086 / ENSG00000165629
ATP5F1D / P30049 / ENSSSCG00000035532 / ENSG00000099624
ATP5F1E / P56381 / ENSSSCG00000060915 / ENSG00000124172
ATP5ME / P56385 / ENSSSCG00000023903 / ENSG00000169020
ATP5MF / P56134 / ENSSSCG00000034019 / ENSG00000241468
ATP5PB / P24539 / ENSSSCG00000037772 / ENSG00000116459
ATP5PF / P18859 / ENSSSCG00000038399 / ENSG00000154723
ATP6 / P00846 / ENSSSCG00000018081 / ENSG00000198899
ATP6AP1 / Q15904 / ENSSSCG00000037310 / ENSG00000071553
ATP6AP2 / O75787 / ENSSSCG00000012247 / ENSG00000182220
ATP6V0A1 / Q93050 / ENSSSCG00000017401 / ENSG00000033627
ATP6V0D1 / P61421 / ENSSSCG00000002776 / ENSG00000159720
ATP6V1A / P38606 / ENSSSCG00000011916 / ENSG00000114573
ATP6V1B2 / P21281 / ENSSSCG00000028694 / ENSG00000147416
ATP6V1D / Q9Y5K8 / ENSSSCG00000033278 / ENSG00000100554
ATP6V1E2 / Q96A05 / ENSSSCG00000008441 / ENSG00000250565
ATP6V1F / Q16864 / ENSSSCG00000016577 / ENSG00000128524
ATP6V1G1 / O75348 / ENSSSCG00000053522 / ENSG00000136888
ATP8 / P03928 / ENSSSCG00000018080 / ENSG00000228253
ATP8A1 / Q9Y2Q0 / ENSSSCG00000008803 / ENSG00000124406
ATP9A / O75110 / ENSSSCG00000007478 / ENSG00000054793
ATXN10 / Q9UBB4 / ENSSSCG00000024864 / ENSG00000130638
ATXN2L / Q8WWM7 / ENSSSCG00000021845 / ENSG00000168488
AUH / Q13825 / ENSSSCG00000009591 / ENSG00000148090
AUP1 / Q9Y679 / ENSSSCG00000008268 / ENSG00000115307
AVIL / O75366 / ENSSSCG00000029571 / ENSG00000135407
AZGP1 / P25311 / ENSSSCG00000007642 / ENSG00000160862
B3GAT3 / O94766 / ENSSSCG00000020809 / ENSG00000149541
BABAM2 / Q9NXR7 / ENSSSCG00000026465 / ENSG00000158019
BAG1 / Q99933 / ENSSSCG00000010996 / ENSG00000107262
BAG3 / O95817 / ENSSSCG00000010686 / ENSG00000151929
BAIAP2 / Q9UQB8 / ENSSSCG00000017142 / ENSG00000175866
BAK1 / Q16611 / ENSSSCG00000001516 / ENSG00000030110
Homologs
Adipose.Tissue
Brain
Colon
Heart
Kidney
Liver
Lung
Pancreas
S.Intestine
Spleen
Organs
Species
Human
Pig
Bins
1
2
3
4
5

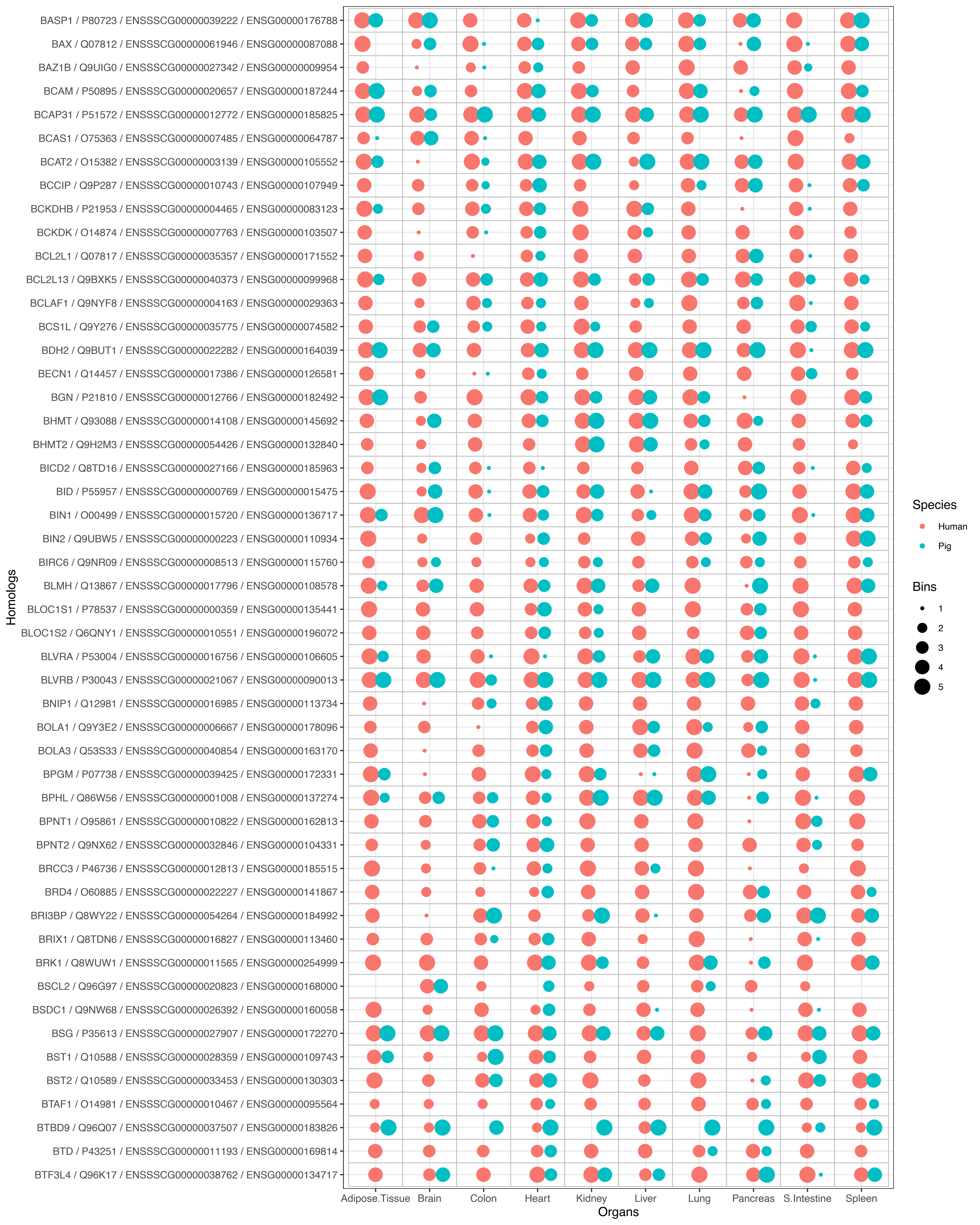
BASP1 / P80723 / ENSSSCG00000039222 / ENSG00000176788
BAX / Q07812 / ENSSSCG00000061946 / ENSG00000087088
BAZ1B / Q9UIG0 / ENSSSCG00000027342 / ENSG00000009954
BCAM / P50895 / ENSSSCG00000020657 / ENSG00000187244
BCAP31 / P51572 / ENSSSCG00000012772 / ENSG00000185825
BCAS1 / O75363 / ENSSSCG00000007485 / ENSG00000064787
BCAT2 / O15382 / ENSSSCG00000003139 / ENSG00000105552
BCCIP / Q9P287 / ENSSSCG00000010743 / ENSG00000107949
BCKDHB / P21953 / ENSSSCG00000004465 / ENSG00000083123
BCKDK / O14874 / ENSSSCG00000007763 / ENSG00000103507
BCL2L1 / Q07817 / ENSSSCG00000035357 / ENSG00000171552
BCL2L13 / Q9BXK5 / ENSSSCG00000040373 / ENSG00000099968
BCLAF1 / Q9NYF8 / ENSSSCG00000004163 / ENSG00000029363
BCS1L / Q9Y276 / ENSSSCG00000035775 / ENSG00000074582
BDH2 / Q9BUT1 / ENSSSCG00000022282 / ENSG00000164039
BECN1 / Q14457 / ENSSSCG00000017386 / ENSG00000126581
BGN / P21810 / ENSSSCG00000012766 / ENSG00000182492
BHMT / Q93088 / ENSSSCG00000014108 / ENSG00000145692
BHMT2 / Q9H2M3 / ENSSSCG00000054426 / ENSG00000132840
BICD2 / Q8TD16 / ENSSSCG00000027166 / ENSG00000185963
BID / P55957 / ENSSSCG00000000769 / ENSG00000015475
BIN1 / O00499 / ENSSSCG00000015720 / ENSG00000136717
BIN2 / Q9UBW5 / ENSSSCG00000000223 / ENSG00000110934
BIRC6 / Q9NR09 / ENSSSCG00000008513 / ENSG00000115760
BLMH / Q13867 / ENSSSCG00000017796 / ENSG00000108578
BLOC1S1 / P78537 / ENSSSCG00000000359 / ENSG00000135441
BLOC1S2 / Q6QNY1 / ENSSSCG00000010551 / ENSG00000196072
BLVRA / P53004 / ENSSSCG00000016756 / ENSG00000106605
BLVRB / P30043 / ENSSSCG00000021067 / ENSG00000090013
BNIP1 / Q12981 / ENSSSCG00000016985 / ENSG00000113734
BOLA1 / Q9Y3E2 / ENSSSCG00000006667 / ENSG00000178096
BOLA3 / Q53S33 / ENSSSCG00000040854 / ENSG00000163170
BPGM / P07738 / ENSSSCG00000039425 / ENSG00000172331
BPHL / Q86W56 / ENSSSCG00000001008 / ENSG00000137274
BPNT1 / O95861 / ENSSSCG00000010822 / ENSG00000162813
BPNT2 / Q9NX62 / ENSSSCG00000032846 / ENSG00000104331
BRCC3 / P46736 / ENSSSCG00000012813 / ENSG00000185515
BRD4 / O60885 / ENSSSCG00000022227 / ENSG00000141867
BRI3BP / Q8WY22 / ENSSSCG00000054264 / ENSG00000184992
BRIX1 / Q8TDN6 / ENSSSCG00000016827 / ENSG00000113460
BRK1 / Q8WUW1 / ENSSSCG00000011565 / ENSG00000254999
BSCL2 / Q96G97 / ENSSSCG00000020823 / ENSG00000168000
BSDC1 / Q9NW68 / ENSSSCG00000026392 / ENSG00000160058
BSG / P35613 / ENSSSCG00000027907 / ENSG00000172270
BST1 / Q10588 / ENSSSCG00000028359 / ENSG00000109743
BST2 / Q10589 / ENSSSCG00000033453 / ENSG00000130303
BTAF1 / O14981 / ENSSSCG00000010467 / ENSG00000095564
BTBD9 / Q96Q07 / ENSSSCG00000037507 / ENSG00000183826
BTD / P43251 / ENSSSCG00000011193 / ENSG00000169814
BTF3L4 / Q96K17 / ENSSSCG00000038762 / ENSG00000134717
Homologs
Adipose.Tissue
Brain
Colon
Heart
Kidney
Liver
Lung
Pancreas
S.Intestine
Spleen
Organs
Species
Human
Pig
Bins
1
2
3
4
5

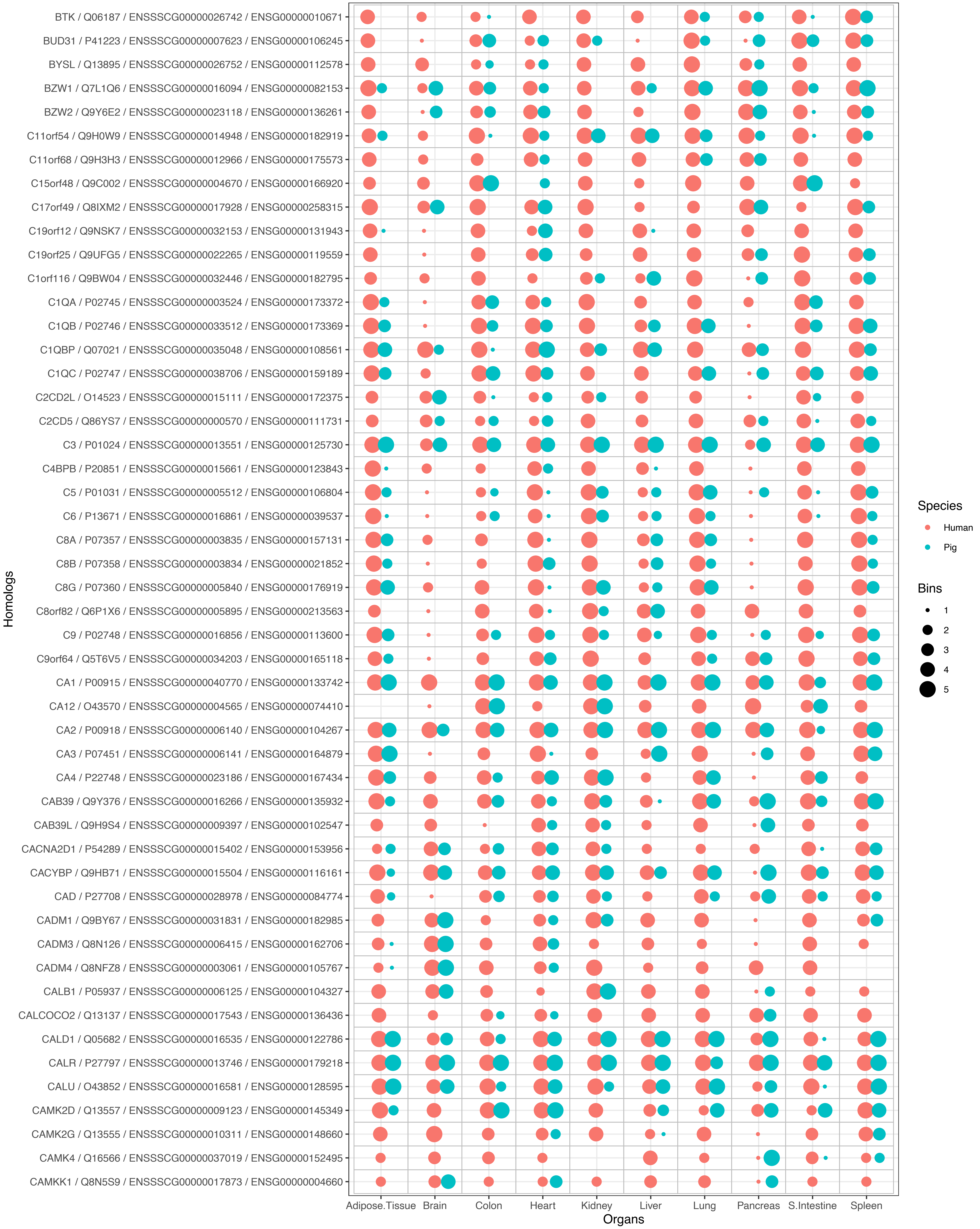

BTK / Q06187 / ENSSSCG00000026742 / ENSG00000010671
BUD31 / P41223 / ENSSSCG00000007623 / ENSG00000106245
BYSL / Q13895 / ENSSSCG00000026752 / ENSG00000112578
BZW1 / Q7L1Q6 / ENSSSCG00000016094 / ENSG00000082153
BZW2 / Q9Y6E2 / ENSSSCG00000023118 / ENSG00000136261
C11orf54 / Q9H0W9 / ENSSSCG00000014948 / ENSG00000182919
C11orf68 / Q9H3H3 / ENSSSCG00000012966 / ENSG00000175573
C15orf48 / Q9C002 / ENSSSCG00000004670 / ENSG00000166920
C17orf49 / Q8IXM2 / ENSSSCG00000017928 / ENSG00000258315
C19orf12 / Q9NSK7 / ENSSSCG00000032153 / ENSG00000131943
C19orf25 / Q9UFG5 / ENSSSCG00000022265 / ENSG00000119559
C1orf116 / Q9BW04 / ENSSSCG00000032446 / ENSG00000182795
C1QA / P02745 / ENSSSCG00000003524 / ENSG00000173372
C1QB / P02746 / ENSSSCG00000033512 / ENSG00000173369
C1QBP / Q07021 / ENSSSCG00000035048 / ENSG00000108561
C1QC / P02747 / ENSSSCG00000038706 / ENSG00000159189
C2CD2L / O14523 / ENSSSCG00000015111 / ENSG00000172375
C2CD5 / Q86YS7 / ENSSSCG00000000570 / ENSG00000111731
C3 / P01024 / ENSSSCG00000013551 / ENSG00000125730
C4BPB / P20851 / ENSSSCG00000015661 / ENSG00000123843
C5 / P01031 / ENSSSCG00000005512 / ENSG00000106804
C6 / P13671 / ENSSSCG00000016861 / ENSG00000039537
C8A / P07357 / ENSSSCG00000003835 / ENSG00000157131
C8B / P07358 / ENSSSCG00000003834 / ENSG00000021852
C8G / P07360 / ENSSSCG00000005840 / ENSG00000176919
C8orf82 / Q6P1X6 / ENSSSCG00000005895 / ENSG00000213563
C9 / P02748 / ENSSSCG00000016856 / ENSG00000113600
C9orf64 / Q5T6V5 / ENSSSCG00000034203 / ENSG00000165118
CA1 / P00915 / ENSSSCG00000040770 / ENSG00000133742
CA12 / O43570 / ENSSSCG00000004565 / ENSG00000074410
CA2 / P00918 / ENSSSCG00000006140 / ENSG00000104267
CA3 / P07451 / ENSSSCG00000006141 / ENSG00000164879
CA4 / P22748 / ENSSSCG00000023186 / ENSG00000167434
CAB39 / Q9Y376 / ENSSSCG00000016266 / ENSG00000135932
CAB39L / Q9H9S4 / ENSSSCG00000009397 / ENSG00000102547
CACNA2D1 / P54289 / ENSSSCG00000015402 / ENSG00000153956
CACYBP / Q9HB71 / ENSSSCG00000015504 / ENSG00000116161
CAD / P27708 / ENSSSCG00000028978 / ENSG00000084774
CADM1 / Q9BY67 / ENSSSCG00000031831 / ENSG00000182985
CADM3 / Q8N126 / ENSSSCG00000006415 / ENSG00000162706
CADM4 / Q8NFZ8 / ENSSSCG00000003061 / ENSG00000105767
CALB1 / P05937 / ENSSSCG00000006125 / ENSG00000104327
CALCOCO2 / Q13137 / ENSSSCG00000017543 / ENSG00000136436
CALD1 / Q05682 / ENSSSCG00000016535 / ENSG00000122786
CALR / P27797 / ENSSSCG00000013746 / ENSG00000179218
CALU / O43852 / ENSSSCG00000016581 / ENSG00000128595
CAMK2D / Q13557 / ENSSSCG00000009123 / ENSG00000145349
CAMK2G / Q13555 / ENSSSCG00000010311 / ENSG00000148660
CAMK4 / Q16566 / ENSSSCG00000037019 / ENSG00000152495
CAMKK1 / Q8N5S9 / ENSSSCG00000017873 / ENSG00000004660
Homologs
Adipose.Tissue
Brain
Colon
Heart
Kidney
Liver
Lung
Pancreas
S.Intestine
Spleen
Organs
Species
Human
Pig
Bins
1
2
3
4
5

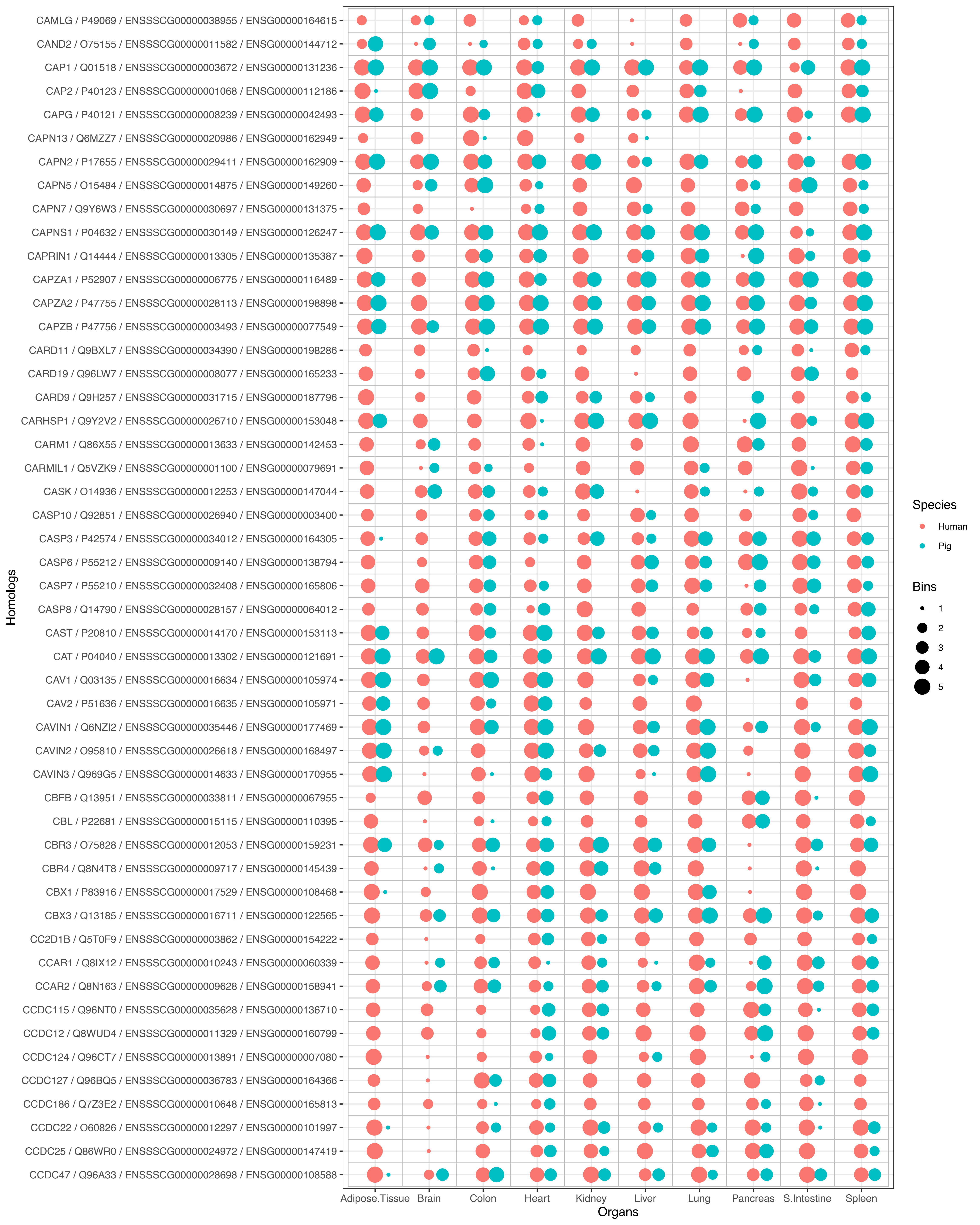
CAMLG / P49069 / ENSSSCG00000038955 / ENSG00000164615
CAND2 / O75155 / ENSSSCG00000011582 / ENSG00000144712
CAP1 / Q01518 / ENSSSCG00000003672 / ENSG00000131236
CAP2 / P40123 / ENSSSCG00000001068 / ENSG00000112186
CAPG / P40121 / ENSSSCG00000008239 / ENSG00000042493
CAPN13 / Q6MZZ7 / ENSSSCG00000020986 / ENSG00000162949
CAPN2 / P17655 / ENSSSCG00000029411 / ENSG00000162909
CAPN5 / O15484 / ENSSSCG00000014875 / ENSG00000149260
CAPN7 / Q9Y6W3 / ENSSSCG00000030697 / ENSG00000131375
CAPNS1 / P04632 / ENSSSCG00000030149 / ENSG00000126247
CAPRIN1 / Q14444 / ENSSSCG00000013305 / ENSG00000135387
CAPZA1 / P52907 / ENSSSCG00000006775 / ENSG00000116489
CAPZA2 / P47755 / ENSSSCG00000028113 / ENSG00000198898
CAPZB / P47756 / ENSSSCG00000003493 / ENSG00000077549
CARD11 / Q9BXL7 / ENSSSCG00000034390 / ENSG00000198286
CARD19 / Q96LW7 / ENSSSCG00000008077 / ENSG00000165233
CARD9 / Q9H257 / ENSSSCG00000031715 / ENSG00000187796
CARHSP1 / Q9Y2V2 / ENSSSCG00000026710 / ENSG00000153048
CARM1 / Q86X55 / ENSSSCG00000013633 / ENSG00000142453
CARMIL1 / Q5VZK9 / ENSSSCG00000001100 / ENSG00000079691
CASK / O14936 / ENSSSCG00000012253 / ENSG00000147044
CASP10 / Q92851 / ENSSSCG00000026940 / ENSG00000003400
CASP3 / P42574 / ENSSSCG00000034012 / ENSG00000164305
CASP6 / P55212 / ENSSSCG00000009140 / ENSG00000138794
CASP7 / P55210 / ENSSSCG00000032408 / ENSG00000165806
CASP8 / Q14790 / ENSSSCG00000028157 / ENSG00000064012
CAST / P20810 / ENSSSCG00000014170 / ENSG00000153113
CAT / P04040 / ENSSSCG00000013302 / ENSG00000121691
CAV1 / Q03135 / ENSSSCG00000016634 / ENSG00000105974
CAV2 / P51636 / ENSSSCG00000016635 / ENSG00000105971
CAVIN1 / Q6NZI2 / ENSSSCG00000035446 / ENSG00000177469
CAVIN2 / O95810 / ENSSSCG00000026618 / ENSG00000168497
CAVIN3 / Q969G5 / ENSSSCG00000014633 / ENSG00000170955
CBFB / Q13951 / ENSSSCG00000033811 / ENSG00000067955
CBL / P22681 / ENSSSCG00000015115 / ENSG00000110395
CBR3 / O75828 / ENSSSCG00000012053 / ENSG00000159231
CBR4 / Q8N4T8 / ENSSSCG00000009717 / ENSG00000145439
CBX1 / P83916 / ENSSSCG00000017529 / ENSG00000108468
CBX3 / Q13185 / ENSSSCG00000016711 / ENSG00000122565
CC2D1B / Q5T0F9 / ENSSSCG00000003862 / ENSG00000154222
CCAR1 / Q8IX12 / ENSSSCG00000010243 / ENSG00000060339
CCAR2 / Q8N163 / ENSSSCG00000009628 / ENSG00000158941
CCDC115 / Q96NT0 / ENSSSCG00000035628 / ENSG00000136710
CCDC12 / Q8WUD4 / ENSSSCG00000011329 / ENSG00000160799
CCDC124 / Q96CT7 / ENSSSCG00000013891 / ENSG00000007080
CCDC127 / Q96BQ5 / ENSSSCG00000036783 / ENSG00000164366
CCDC186 / Q7Z3E2 / ENSSSCG00000010648 / ENSG00000165813
CCDC22 / O60826 / ENSSSCG00000012297 / ENSG00000101997
CCDC25 / Q86WR0 / ENSSSCG00000024972 / ENSG00000147419
CCDC47 / Q96A33 / ENSSSCG00000028698 / ENSG00000108588
Homologs
Adipose.Tissue
Brain
Colon
Heart
Kidney
Liver
Lung
Pancreas
S.Intestine
Spleen
Organs
Species
Human
Pig
Bins
1
2
3
4
5

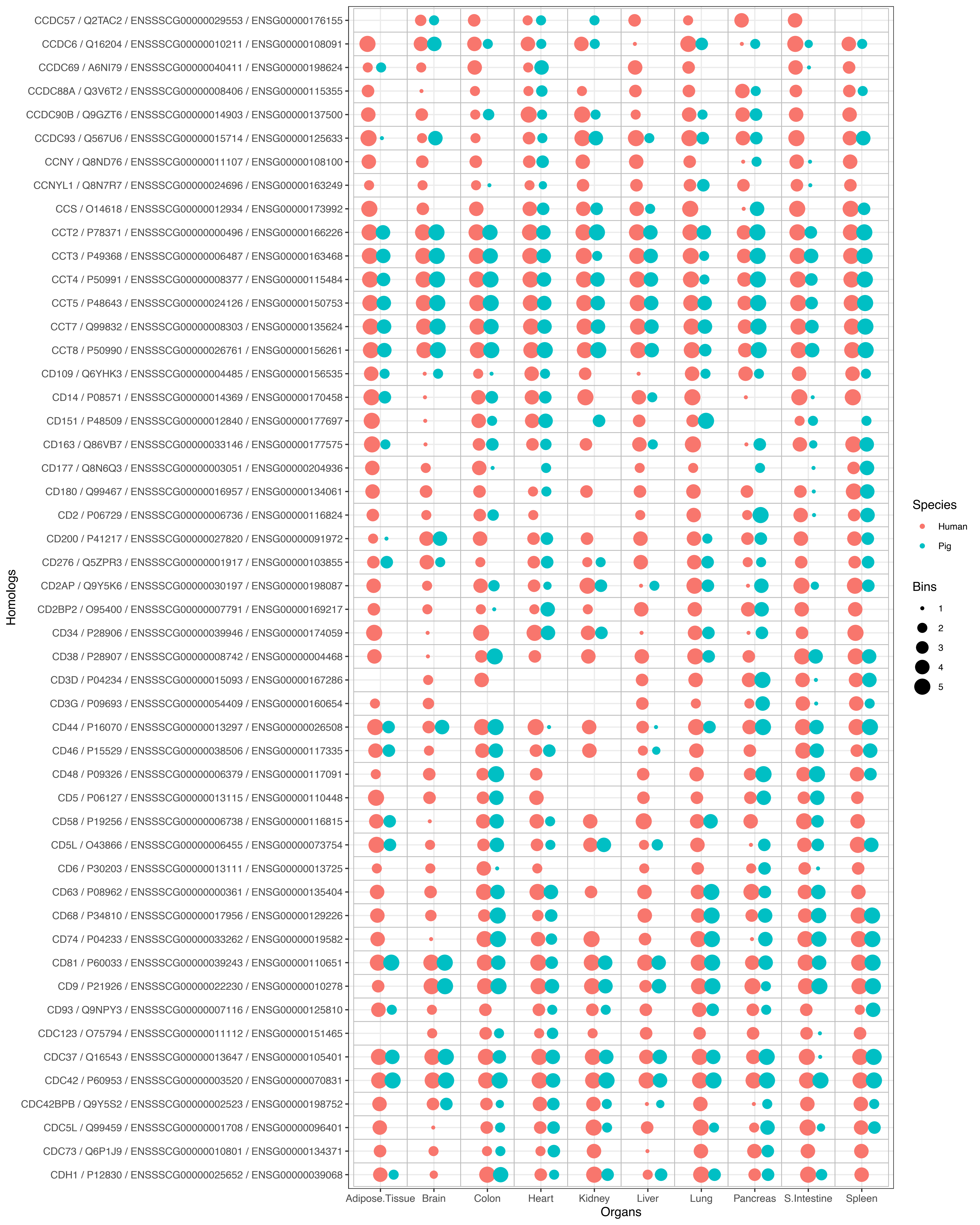

CCDC57 / Q2TAC2 / ENSSSCG00000029553 / ENSG00000176155
CCDC6 / Q16204 / ENSSSCG00000010211 / ENSG00000108091
CCDC69 / A6NI79 / ENSSSCG00000040411 / ENSG00000198624
CCDC88A / Q3V6T2 / ENSSSCG00000008406 / ENSG00000115355
CCDC90B / Q9GZT6 / ENSSSCG00000014903 / ENSG00000137500
CCDC93 / Q567U6 / ENSSSCG00000015714 / ENSG00000125633
CCNY / Q8ND76 / ENSSSCG00000011107 / ENSG00000108100
CCNYL1 / Q8N7R7 / ENSSSCG00000024696 / ENSG00000163249
CCS / O14618 / ENSSSCG00000012934 / ENSG00000173992
CCT2 / P78371 / ENSSSCG00000000496 / ENSG00000166226
CCT3 / P49368 / ENSSSCG00000006487 / ENSG00000163468
CCT4 / P50991 / ENSSSCG00000008377 / ENSG00000115484
CCT5 / P48643 / ENSSSCG00000024126 / ENSG00000150753
CCT7 / Q99832 / ENSSSCG00000008303 / ENSG00000135624
CCT8 / P50990 / ENSSSCG00000026761 / ENSG00000156261
CD109 / Q6YHK3 / ENSSSCG00000004485 / ENSG00000156535
CD14 / P08571 / ENSSSCG00000014369 / ENSG00000170458
CD151 / P48509 / ENSSSCG00000012840 / ENSG00000177697
CD163 / Q86VB7 / ENSSSCG00000033146 / ENSG00000177575
CD177 / Q8N6Q3 / ENSSSCG00000003051 / ENSG00000204936
CD180 / Q99467 / ENSSSCG00000016957 / ENSG00000134061
CD2 / P06729 / ENSSSCG00000006736 / ENSG00000116824
CD200 / P41217 / ENSSSCG00000027820 / ENSG00000091972
CD276 / Q5ZPR3 / ENSSSCG00000001917 / ENSG00000103855
CD2AP / Q9Y5K6 / ENSSSCG00000030197 / ENSG00000198087
CD2BP2 / O95400 / ENSSSCG00000007791 / ENSG00000169217
CD34 / P28906 / ENSSSCG00000039946 / ENSG00000174059
CD38 / P28907 / ENSSSCG00000008742 / ENSG00000004468
CD3D / P04234 / ENSSSCG00000015093 / ENSG00000167286
CD3G / P09693 / ENSSSCG00000054409 / ENSG00000160654
CD44 / P16070 / ENSSSCG00000013297 / ENSG00000026508
CD46 / P15529 / ENSSSCG00000038506 / ENSG00000117335
CD48 / P09326 / ENSSSCG00000006379 / ENSG00000117091
CD5 / P06127 / ENSSSCG00000013115 / ENSG00000110448
CD58 / P19256 / ENSSSCG00000006738 / ENSG00000116815
CD5L / O43866 / ENSSSCG00000006455 / ENSG00000073754
CD6 / P30203 / ENSSSCG00000013111 / ENSG00000013725
CD63 / P08962 / ENSSSCG00000000361 / ENSG00000135404
CD68 / P34810 / ENSSSCG00000017956 / ENSG00000129226
CD74 / P04233 / ENSSSCG00000033262 / ENSG00000019582
CD81 / P60033 / ENSSSCG00000039243 / ENSG00000110651
CD9 / P21926 / ENSSSCG00000022230 / ENSG00000010278
CD93 / Q9NPY3 / ENSSSCG00000007116 / ENSG00000125810
CDC123 / O75794 / ENSSSCG00000011112 / ENSG00000151465
CDC37 / Q16543 / ENSSSCG00000013647 / ENSG00000105401
CDC42 / P60953 / ENSSSCG00000003520 / ENSG00000070831
CDC42BPB / Q9Y5S2 / ENSSSCG00000002523 / ENSG00000198752
CDC5L / Q99459 / ENSSSCG00000001708 / ENSG00000096401
CDC73 / Q6P1J9 / ENSSSCG00000010801 / ENSG00000134371
CDH1 / P12830 / ENSSSCG00000025652 / ENSG00000039068
Homologs
Adipose.Tissue
Brain
Colon
Heart
Kidney
Liver
Lung
Pancreas
S.Intestine
Spleen
Organs
Species
Human
Pig
Bins
1
2
3
4
5

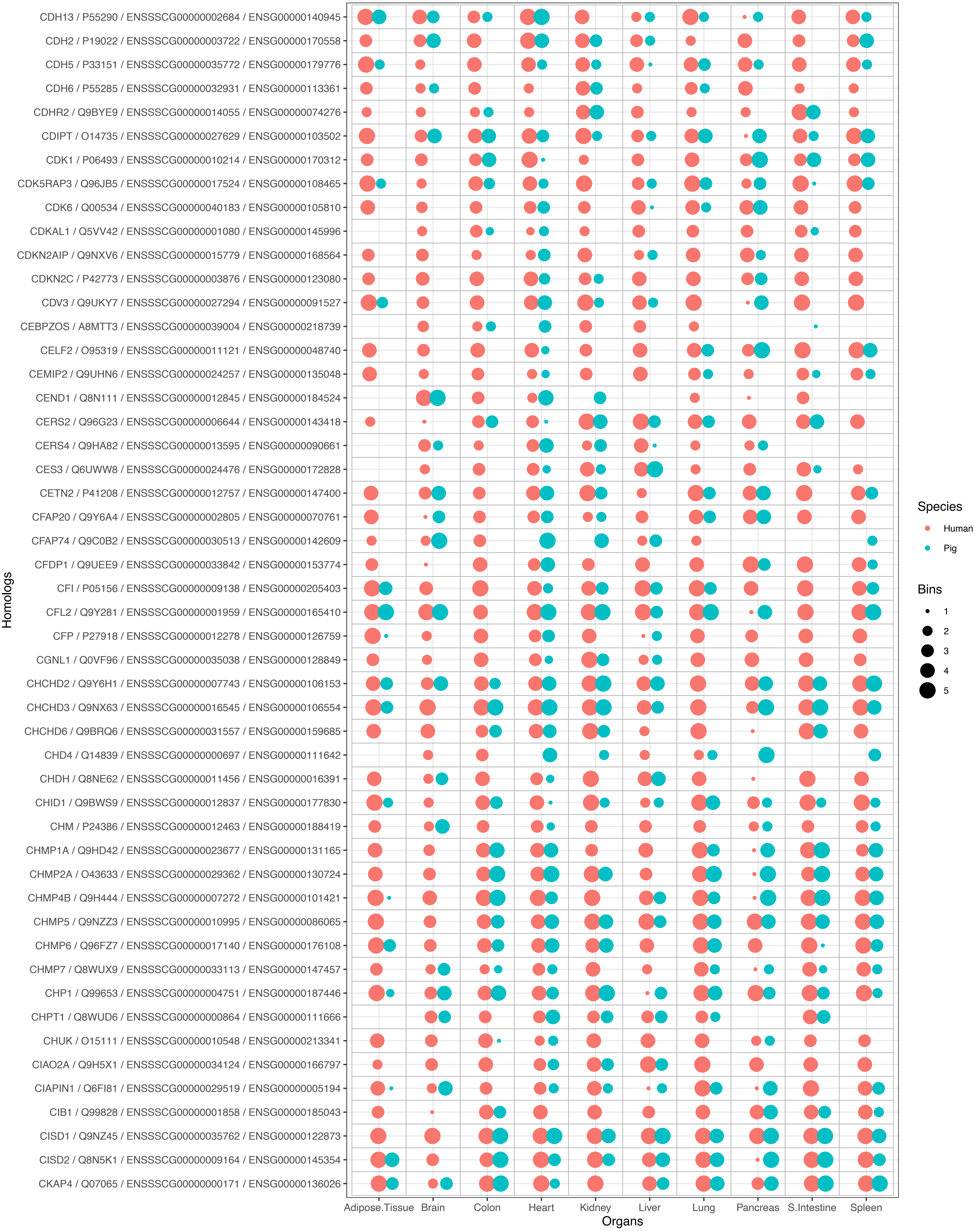
CDH13 / P55290 / ENSSSCG00000002684 / ENSG00000140945
CDH2 / P19022 / ENSSSCG00000003722 / ENSG00000170558
CDH5 / P33151 / ENSSSCG00000035772 / ENSG00000179776
CDH6 / P55285 / ENSSSCG00000032931 / ENSG00000113361
CDHR2 / Q9BYE9 / ENSSSCG00000014055 / ENSG00000074276
CDIPT / O14735 / ENSSSCG00000027629 / ENSG00000103502
CDK1 / P06493 / ENSSSCG00000010214 / ENSG00000170312
CDK5RAP3 / Q96JB5 / ENSSSCG00000017524 / ENSG00000108465
CDK6 / Q00534 / ENSSSCG00000040183 / ENSG00000105810
CDKAL1 / Q5VV42 / ENSSSCG00000001080 / ENSG00000145996
CDKN2AIP / Q9NXV6 / ENSSSCG00000015779 / ENSG00000168564
CDKN2C / P42773 / ENSSSCG00000003876 / ENSG00000123080
CDV3 / Q9UKY7 / ENSSSCG00000027294 / ENSG00000091527
CEBPZOS / A8MTT3 / ENSSSCG00000039004 / ENSG00000218739
CELF2 / O95319 / ENSSSCG00000011121 / ENSG00000048740
CEMIP2 / Q9UHN6 / ENSSSCG00000024257 / ENSG00000135048
CEND1 / Q8N111 / ENSSSCG00000012845 / ENSG00000184524
CERS2 / Q96G23 / ENSSSCG00000006644 / ENSG00000143418
CERS4 / Q9HA82 / ENSSSCG00000013595 / ENSG00000090661
CES3 / Q6UWW8 / ENSSSCG00000024476 / ENSG00000172828
CETN2 / P41208 / ENSSSCG00000012757 / ENSG00000147400
CFAP20 / Q9Y6A4 / ENSSSCG00000002805 / ENSG00000070761
CFAP74 / Q9C0B2 / ENSSSCG00000030513 / ENSG00000142609
CFDP1 / Q9UEE9 / ENSSSCG00000033842 / ENSG00000153774
CFI / P05156 / ENSSSCG00000009138 / ENSG00000205403
CFL2 / Q9Y281 / ENSSSCG00000001959 / ENSG00000165410
CFP / P27918 / ENSSSCG00000012278 / ENSG00000126759
CGNL1 / Q0VF96 / ENSSSCG00000035038 / ENSG00000128849
CHCHD2 / Q9Y6H1 / ENSSSCG00000007743 / ENSG00000106153
CHCHD3 / Q9NX63 / ENSSSCG00000016545 / ENSG00000106554
CHCHD6 / Q9BRQ6 / ENSSSCG00000031557 / ENSG00000159685
CHD4 / Q14839 / ENSSSCG00000000697 / ENSG00000111642
CHDH / Q8NE62 / ENSSSCG00000011456 / ENSG00000016391
CHID1 / Q9BWS9 / ENSSSCG00000012837 / ENSG00000177830
CHM / P24386 / ENSSSCG00000012463 / ENSG00000188419
CHMP1A / Q9HD42 / ENSSSCG00000023677 / ENSG00000131165
CHMP2A / O43633 / ENSSSCG00000029362 / ENSG00000130724
CHMP4B / Q9H444 / ENSSSCG00000007272 / ENSG00000101421
CHMP5 / Q9NZZ3 / ENSSSCG00000010995 / ENSG00000086065
CHMP6 / Q96FZ7 / ENSSSCG00000017140 / ENSG00000176108
CHMP7 / Q8WUX9 / ENSSSCG00000033113 / ENSG00000147457
CHP1 / Q99653 / ENSSSCG00000004751 / ENSG00000187446
CHPT1 / Q8WUD6 / ENSSSCG00000000864 / ENSG00000111666
CHUK / O15111 / ENSSSCG00000010548 / ENSG00000213341
CIAO2A / Q9H5X1 / ENSSSCG00000034124 / ENSG00000166797
CIAPIN1 / Q6FI81 / ENSSSCG00000029519 / ENSG00000005194
CIB1 / Q99828 / ENSSSCG00000001858 / ENSG00000185043
CISD1 / Q9NZ45 / ENSSSCG00000035762 / ENSG00000122873
CISD2 / Q8N5K1 / ENSSSCG00000009164 / ENSG00000145354
CKAP4 / Q07065 / ENSSSCG00000000171 / ENSG00000136026
Homologs
Adipose.Tissue
Brain
Colon
Heart
Kidney
Liver
Lung
Pancreas
S.Intestine
Spleen
Organs
Species
Human
Pig
Bins
1
2
3
4
5

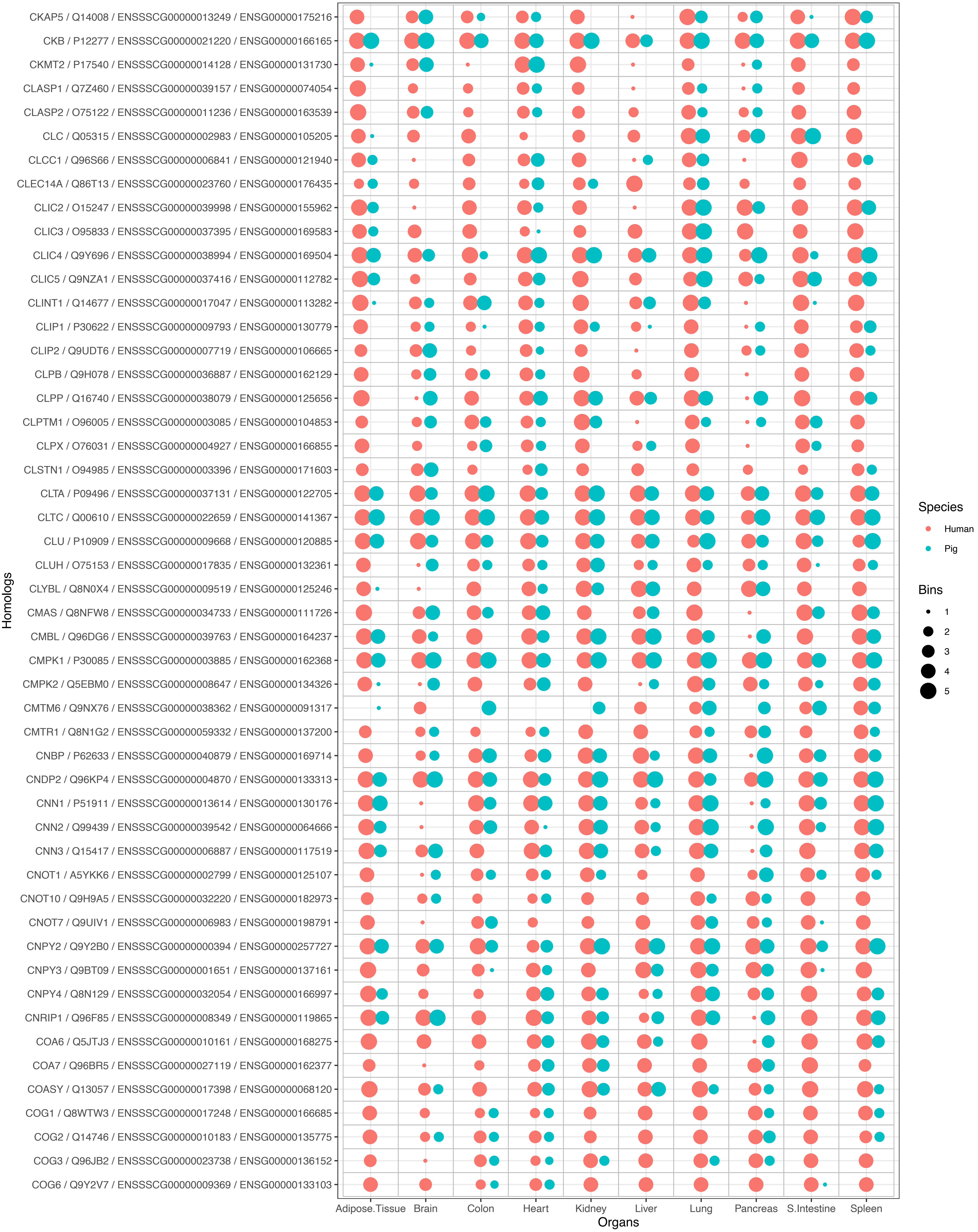
CKAP5 / Q14008 / ENSSSCG00000013249 / ENSG00000175216
CKB / P12277 / ENSSSCG00000021220 / ENSG00000166165
CKMT2 / P17540 / ENSSSCG00000014128 / ENSG00000131730
CLASP1 / Q7Z460 / ENSSSCG00000039157 / ENSG00000074054
CLASP2 / O75122 / ENSSSCG00000011236 / ENSG00000163539
CLC / Q05315 / ENSSSCG00000002983 / ENSG00000105205
CLCC1 / Q96S66 / ENSSSCG00000006841 / ENSG00000121940
CLEC14A / Q86T13 / ENSSSCG00000023760 / ENSG00000176435
CLIC2 / O15247 / ENSSSCG00000039998 / ENSG00000155962
CLIC3 / O95833 / ENSSSCG00000037395 / ENSG00000169583
CLIC4 / Q9Y696 / ENSSSCG00000038994 / ENSG00000169504
CLIC5 / Q9NZA1 / ENSSSCG00000037416 / ENSG00000112782
CLINT1 / Q14677 / ENSSSCG00000017047 / ENSG00000113282
CLIP1 / P30622 / ENSSSCG00000009793 / ENSG00000130779
CLIP2 / Q9UDT6 / ENSSSCG00000007719 / ENSG00000106665
CLPB / Q9H078 / ENSSSCG00000036887 / ENSG00000162129
CLPP / Q16740 / ENSSSCG00000038079 / ENSG00000125656
CLPTM1 / O96005 / ENSSSCG00000003085 / ENSG00000104853
CLPX / O76031 / ENSSSCG00000004927 / ENSG00000166855
CLSTN1 / O94985 / ENSSSCG00000003396 / ENSG00000171603
CLTA / P09496 / ENSSSCG00000037131 / ENSG00000122705
CLTC / Q00610 / ENSSSCG00000022659 / ENSG00000141367
CLU / P10909 / ENSSSCG00000009668 / ENSG00000120885
CLUH / O75153 / ENSSSCG00000017835 / ENSG00000132361
CLYBL / Q8N0X4 / ENSSSCG00000009519 / ENSG00000125246
CMAS / Q8NFW8 / ENSSSCG00000034733 / ENSG00000111726
CMBL / Q96DG6 / ENSSSCG00000039763 / ENSG00000164237
CMPK1 / P30085 / ENSSSCG00000003885 / ENSG00000162368
CMPK2 / Q5EBM0 / ENSSSCG00000008647 / ENSG00000134326
CMTM6 / Q9NX76 / ENSSSCG00000038362 / ENSG00000091317
CMTR1 / Q8N1G2 / ENSSSCG00000059332 / ENSG00000137200
CNBP / P62633 / ENSSSCG00000040879 / ENSG00000169714
CNDP2 / Q96KP4 / ENSSSCG00000004870 / ENSG00000133313
CNN1 / P51911 / ENSSSCG00000013614 / ENSG00000130176
CNN2 / Q99439 / ENSSSCG00000039542 / ENSG00000064666
CNN3 / Q15417 / ENSSSCG00000006887 / ENSG00000117519
CNOT1 / A5YKK6 / ENSSSCG00000002799 / ENSG00000125107
CNOT10 / Q9H9A5 / ENSSSCG00000032220 / ENSG00000182973
CNOT7 / Q9UIV1 / ENSSSCG00000006983 / ENSG00000198791
CNPY2 / Q9Y2B0 / ENSSSCG00000000394 / ENSG00000257727
CNPY3 / Q9BT09 / ENSSSCG00000001651 / ENSG00000137161
CNPY4 / Q8N129 / ENSSSCG00000032054 / ENSG00000166997
CNRIP1 / Q96F85 / ENSSSCG00000008349 / ENSG00000119865
COA6 / Q5JTJ3 / ENSSSCG00000010161 / ENSG00000168275
COA7 / Q96BR5 / ENSSSCG00000027119 / ENSG00000162377
COASY / Q13057 / ENSSSCG00000017398 / ENSG00000068120
COG1 / Q8WTW3 / ENSSSCG00000017248 / ENSG00000166685
COG2 / Q14746 / ENSSSCG00000010183 / ENSG00000135775
COG3 / Q96JB2 / ENSSSCG00000023738 / ENSG00000136152
COG6 / Q9Y2V7 / ENSSSCG00000009369 / ENSG00000133103
Homologs
Adipose.Tissue
Brain
Colon
Heart
Kidney
Liver
Lung
Pancreas
S.Intestine
Spleen
Organs
Species
Human
Pig
Bins
1
2
3
4
5

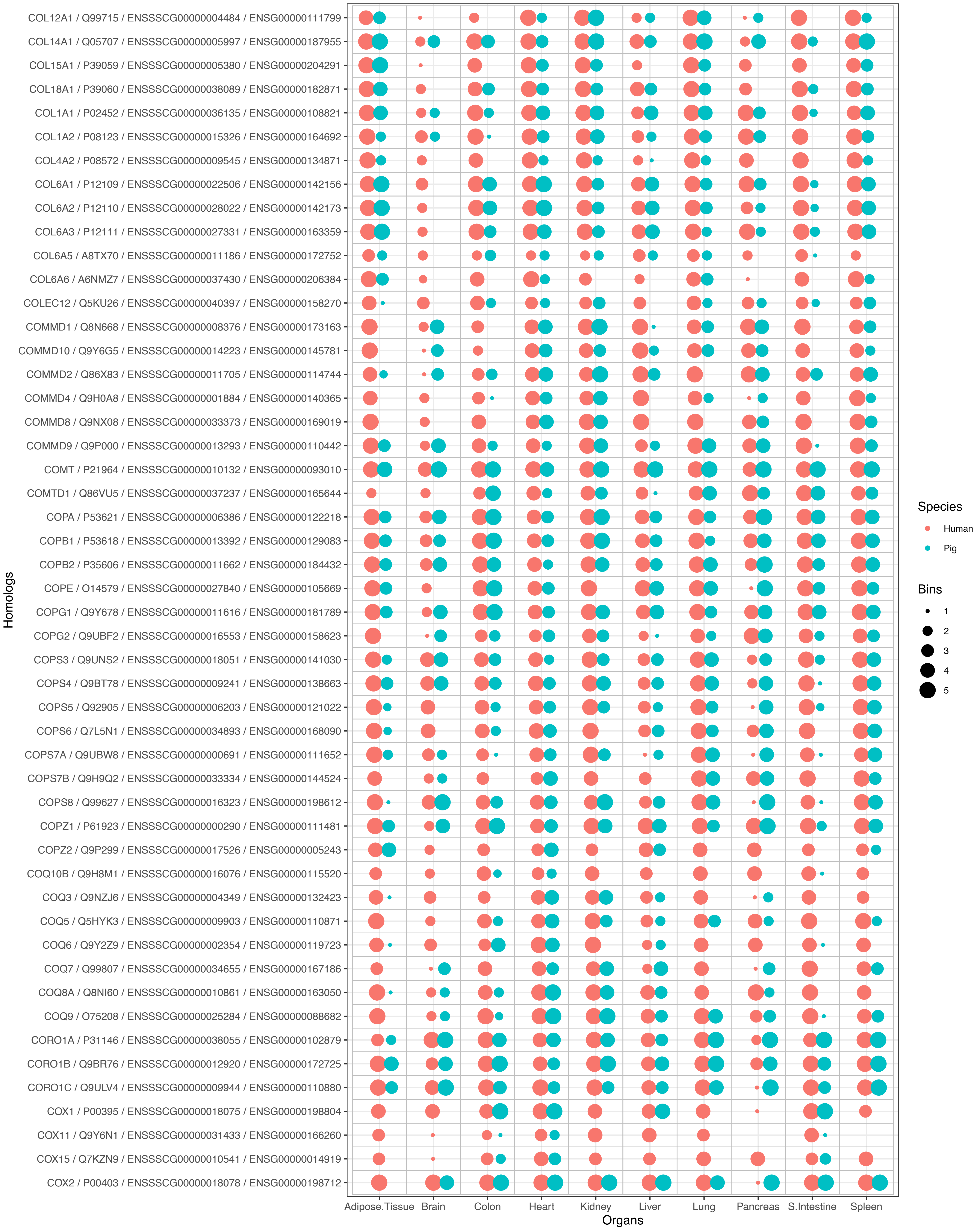
COL12A1 / Q99715 / ENSSSCG00000004484 / ENSG00000111799
COL14A1 / Q05707 / ENSSSCG00000005997 / ENSG00000187955
COL15A1 / P39059 / ENSSSCG00000005380 / ENSG00000204291
COL18A1 / P39060 / ENSSSCG00000038089 / ENSG00000182871
COL1A1 / P02452 / ENSSSCG00000036135 / ENSG00000108821
COL1A2 / P08123 / ENSSSCG00000015326 / ENSG00000164692
COL4A2 / P08572 / ENSSSCG00000009545 / ENSG00000134871
COL6A1 / P12109 / ENSSSCG00000022506 / ENSG00000142156
COL6A2 / P12110 / ENSSSCG00000028022 / ENSG00000142173
COL6A3 / P12111 / ENSSSCG00000027331 / ENSG00000163359
COL6A5 / A8TX70 / ENSSSCG00000011186 / ENSG00000172752
COL6A6 / A6NMZ7 / ENSSSCG00000037430 / ENSG00000206384
COLEC12 / Q5KU26 / ENSSSCG00000040397 / ENSG00000158270
COMMD1 / Q8N668 / ENSSSCG00000008376 / ENSG00000173163
COMMD10 / Q9Y6G5 / ENSSSCG00000014223 / ENSG00000145781
COMMD2 / Q86X83 / ENSSSCG00000011705 / ENSG00000114744
COMMD4 / Q9H0A8 / ENSSSCG00000001884 / ENSG00000140365
COMMD8 / Q9NX08 / ENSSSCG00000033373 / ENSG00000169019
COMMD9 / Q9P000 / ENSSSCG00000013293 / ENSG00000110442
COMT / P21964 / ENSSSCG00000010132 / ENSG00000093010
COMTD1 / Q86VU5 / ENSSSCG00000037237 / ENSG00000165644
COPA / P53621 / ENSSSCG00000006386 / ENSG00000122218
COPB1 / P53618 / ENSSSCG00000013392 / ENSG00000129083
COPB2 / P35606 / ENSSSCG00000011662 / ENSG00000184432
COPE / O14579 / ENSSSCG00000027840 / ENSG00000105669
COPG1 / Q9Y678 / ENSSSCG00000011616 / ENSG00000181789
COPG2 / Q9UBF2 / ENSSSCG00000016553 / ENSG00000158623
COPS3 / Q9UNS2 / ENSSSCG00000018051 / ENSG00000141030
COPS4 / Q9BT78 / ENSSSCG00000009241 / ENSG00000138663
COPS5 / Q92905 / ENSSSCG00000006203 / ENSG00000121022
COPS6 / Q7L5N1 / ENSSSCG00000034893 / ENSG00000168090
COPS7A / Q9UBW8 / ENSSSCG00000000691 / ENSG00000111652
COPS7B / Q9H9Q2 / ENSSSCG00000033334 / ENSG00000144524
COPS8 / Q99627 / ENSSSCG00000016323 / ENSG00000198612
COPZ1 / P61923 / ENSSSCG00000000290 / ENSG00000111481
COPZ2 / Q9P299 / ENSSSCG00000017526 / ENSG00000005243
COQ10B / Q9H8M1 / ENSSSCG00000016076 / ENSG00000115520
COQ3 / Q9NZJ6 / ENSSSCG00000004349 / ENSG00000132423
COQ5 / Q5HYK3 / ENSSSCG00000009903 / ENSG00000110871
COQ6 / Q9Y2Z9 / ENSSSCG00000002354 / ENSG00000119723
COQ7 / Q99807 / ENSSSCG00000034655 / ENSG00000167186
COQ8A / Q8NI60 / ENSSSCG00000010861 / ENSG00000163050
COQ9 / O75208 / ENSSSCG00000025284 / ENSG00000088682
CORO1A / P31146 / ENSSSCG00000038055 / ENSG00000102879
CORO1B / Q9BR76 / ENSSSCG00000012920 / ENSG00000172725
CORO1C / Q9ULV4 / ENSSSCG00000009944 / ENSG00000110880
COX1 / P00395 / ENSSSCG00000018075 / ENSG00000198804
COX11 / Q9Y6N1 / ENSSSCG00000031433 / ENSG00000166260
COX15 / Q7KZN9 / ENSSSCG00000010541 / ENSG00000014919
COX2 / P00403 / ENSSSCG00000018078 / ENSG00000198712
Homologs
Adipose.Tissue
Brain
Colon
Heart
Kidney
Liver
Lung
Pancreas
S.Intestine
Spleen
Organs
Species
Human
Pig
Bins
1
2
3
4
5

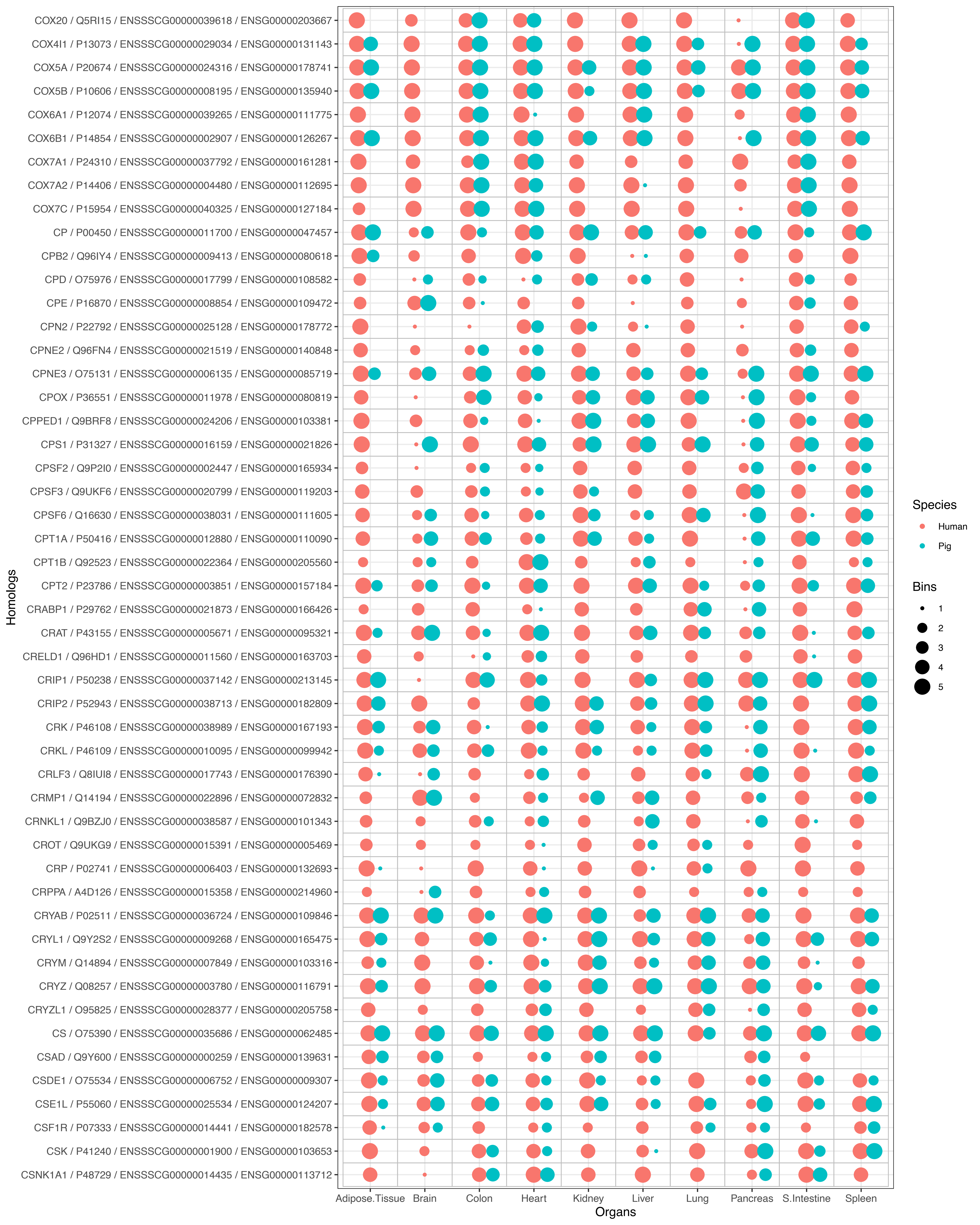
COX20 / Q5RI15 / ENSSSCG00000039618 / ENSG00000203667
COX4I1 / P13073 / ENSSSCG00000029034 / ENSG00000131143
COX5A / P20674 / ENSSSCG00000024316 / ENSG00000178741
COX5B / P10606 / ENSSSCG00000008195 / ENSG00000135940
COX6A1 / P12074 / ENSSSCG00000039265 / ENSG00000111775
COX6B1 / P14854 / ENSSSCG00000002907 / ENSG00000126267
COX7A1 / P24310 / ENSSSCG00000037792 / ENSG00000161281
COX7A2 / P14406 / ENSSSCG00000004480 / ENSG00000112695
COX7C / P15954 / ENSSSCG00000040325 / ENSG00000127184
CP / P00450 / ENSSSCG00000011700 / ENSG00000047457
CPB2 / Q96IY4 / ENSSSCG00000009413 / ENSG00000080618
CPD / O75976 / ENSSSCG00000017799 / ENSG00000108582
CPE / P16870 / ENSSSCG00000008854 / ENSG00000109472
CPN2 / P22792 / ENSSSCG00000025128 / ENSG00000178772
CPNE2 / Q96FN4 / ENSSSCG00000021519 / ENSG00000140848
CPNE3 / O75131 / ENSSSCG00000006135 / ENSG00000085719
CPOX / P36551 / ENSSSCG00000011978 / ENSG00000080819
CPPED1 / Q9BRF8 / ENSSSCG00000024206 / ENSG00000103381
CPS1 / P31327 / ENSSSCG00000016159 / ENSG00000021826
CPSF2 / Q9P2I0 / ENSSSCG00000002447 / ENSG00000165934
CPSF3 / Q9UKF6 / ENSSSCG00000020799 / ENSG00000119203
CPSF6 / Q16630 / ENSSSCG00000038031 / ENSG00000111605
CPT1A / P50416 / ENSSSCG00000012880 / ENSG00000110090
CPT1B / Q92523 / ENSSSCG00000022364 / ENSG00000205560
CPT2 / P23786 / ENSSSCG00000003851 / ENSG00000157184
CRABP1 / P29762 / ENSSSCG00000021873 / ENSG00000166426
CRAT / P43155 / ENSSSCG00000005671 / ENSG00000095321
CRELD1 / Q96HD1 / ENSSSCG00000011560 / ENSG00000163703
CRIP1 / P50238 / ENSSSCG00000037142 / ENSG00000213145
CRIP2 / P52943 / ENSSSCG00000038713 / ENSG00000182809
CRK / P46108 / ENSSSCG00000038989 / ENSG00000167193
CRKL / P46109 / ENSSSCG00000010095 / ENSG00000099942
CRLF3 / Q8IUI8 / ENSSSCG00000017743 / ENSG00000176390
CRMP1 / Q14194 / ENSSSCG00000022896 / ENSG00000072832
CRNKL1 / Q9BZJ0 / ENSSSCG00000038587 / ENSG00000101343
CROT / Q9UKG9 / ENSSSCG00000015391 / ENSG00000005469
CRP / P02741 / ENSSSCG00000006403 / ENSG00000132693
CRPPA / A4D126 / ENSSSCG00000015358 / ENSG00000214960
CRYAB / P02511 / ENSSSCG00000036724 / ENSG00000109846
CRYL1 / Q9Y2S2 / ENSSSCG00000009268 / ENSG00000165475
CRYM / Q14894 / ENSSSCG00000007849 / ENSG00000103316
CRYZ / Q08257 / ENSSSCG00000003780 / ENSG00000116791
CRYZL1 / O95825 / ENSSSCG00000028377 / ENSG00000205758
CS / O75390 / ENSSSCG00000035686 / ENSG00000062485
CSAD / Q9Y600 / ENSSSCG00000000259 / ENSG00000139631
CSDE1 / O75534 / ENSSSCG00000006752 / ENSG00000009307
CSE1L / P55060 / ENSSSCG00000025534 / ENSG00000124207
CSF1R / P07333 / ENSSSCG00000014441 / ENSG00000182578
CSK / P41240 / ENSSSCG00000001900 / ENSG00000103653
CSNK1A1 / P48729 / ENSSSCG00000014435 / ENSG00000113712
Homologs
Adipose.Tissue
Brain
Colon
Heart
Kidney
Liver
Lung
Pancreas
S.Intestine
Spleen
Organs
Species
Human
Pig
Bins
1
2
3
4
5

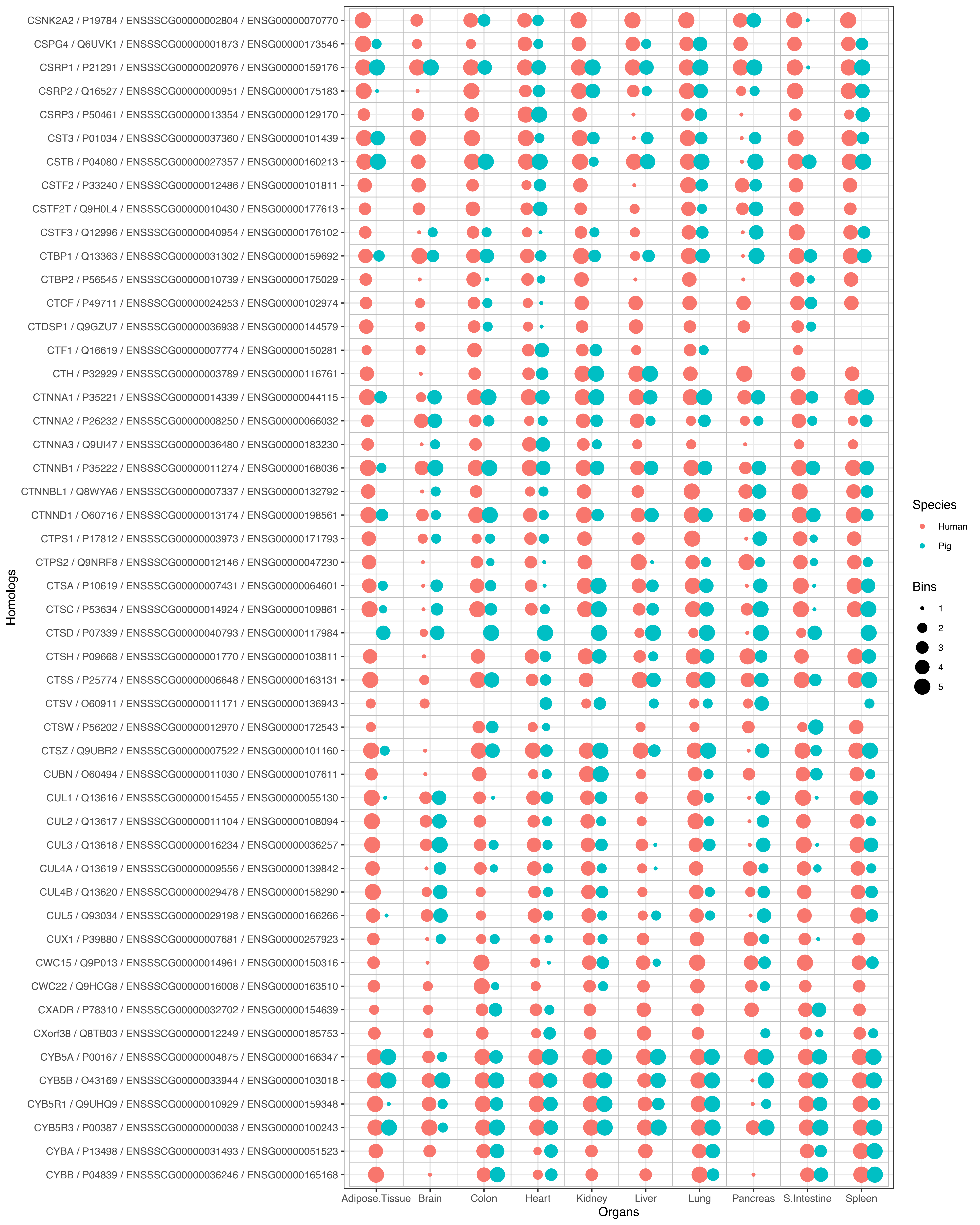
CSNK2A2 / P19784 / ENSSSCG00000002804 / ENSG00000070770
CSPG4 / Q6UVK1 / ENSSSCG00000001873 / ENSG00000173546
CSRP1 / P21291 / ENSSSCG00000020976 / ENSG00000159176
CSRP2 / Q16527 / ENSSSCG00000000951 / ENSG00000175183
CSRP3 / P50461 / ENSSSCG00000013354 / ENSG00000129170
CST3 / P01034 / ENSSSCG00000037360 / ENSG00000101439
CSTB / P04080 / ENSSSCG00000027357 / ENSG00000160213
CSTF2 / P33240 / ENSSSCG00000012486 / ENSG00000101811
CSTF2T / Q9H0L4 / ENSSSCG00000010430 / ENSG00000177613
CSTF3 / Q12996 / ENSSSCG00000040954 / ENSG00000176102
CTBP1 / Q13363 / ENSSSCG00000031302 / ENSG00000159692
CTBP2 / P56545 / ENSSSCG00000010739 / ENSG00000175029
CTCF / P49711 / ENSSSCG00000024253 / ENSG00000102974
CTDSP1 / Q9GZU7 / ENSSSCG00000036938 / ENSG00000144579
CTF1 / Q16619 / ENSSSCG00000007774 / ENSG00000150281
CTH / P32929 / ENSSSCG00000003789 / ENSG00000116761
CTNNA1 / P35221 / ENSSSCG00000014339 / ENSG00000044115
CTNNA2 / P26232 / ENSSSCG00000008250 / ENSG00000066032
CTNNA3 / Q9UI47 / ENSSSCG00000036480 / ENSG00000183230
CTNNB1 / P35222 / ENSSSCG00000011274 / ENSG00000168036
CTNNBL1 / Q8WYA6 / ENSSSCG00000007337 / ENSG00000132792
CTNND1 / O60716 / ENSSSCG00000013174 / ENSG00000198561
CTPS1 / P17812 / ENSSSCG00000003973 / ENSG00000171793
CTPS2 / Q9NRF8 / ENSSSCG00000012146 / ENSG00000047230
CTSA / P10619 / ENSSSCG00000007431 / ENSG00000064601
CTSC / P53634 / ENSSSCG00000014924 / ENSG00000109861
CTSD / P07339 / ENSSSCG00000040793 / ENSG00000117984
CTSH / P09668 / ENSSSCG00000001770 / ENSG00000103811
CTSS / P25774 / ENSSSCG00000006648 / ENSG00000163131
CTSV / O60911 / ENSSSCG00000011171 / ENSG00000136943
CTSW / P56202 / ENSSSCG00000012970 / ENSG00000172543
CTSZ / Q9UBR2 / ENSSSCG00000007522 / ENSG00000101160
CUBN / O60494 / ENSSSCG00000011030 / ENSG00000107611
CUL1 / Q13616 / ENSSSCG00000015455 / ENSG00000055130
CUL2 / Q13617 / ENSSSCG00000011104 / ENSG00000108094
CUL3 / Q13618 / ENSSSCG00000016234 / ENSG00000036257
CUL4A / Q13619 / ENSSSCG00000009556 / ENSG00000139842
CUL4B / Q13620 / ENSSSCG00000029478 / ENSG00000158290
CUL5 / Q93034 / ENSSSCG00000029198 / ENSG00000166266
CUX1 / P39880 / ENSSSCG00000007681 / ENSG00000257923
CWC15 / Q9P013 / ENSSSCG00000014961 / ENSG00000150316
CWC22 / Q9HCG8 / ENSSSCG00000016008 / ENSG00000163510
CXADR / P78310 / ENSSSCG00000032702 / ENSG00000154639
CXorf38 / Q8TB03 / ENSSSCG00000012249 / ENSG00000185753
CYB5A / P00167 / ENSSSCG00000004875 / ENSG00000166347
CYB5B / O43169 / ENSSSCG00000033944 / ENSG00000103018
CYB5R1 / Q9UHQ9 / ENSSSCG00000010929 / ENSG00000159348
CYB5R3 / P00387 / ENSSSCG00000000038 / ENSG00000100243
CYBA / P13498 / ENSSSCG00000031493 / ENSG00000051523
CYBB / P04839 / ENSSSCG00000036246 / ENSG00000165168
Homologs
Adipose.Tissue
Brain
Colon
Heart
Kidney
Liver
Lung
Pancreas
S.Intestine
Spleen
Organs
Species
Human
Pig
Bins
1
2
3
4
5

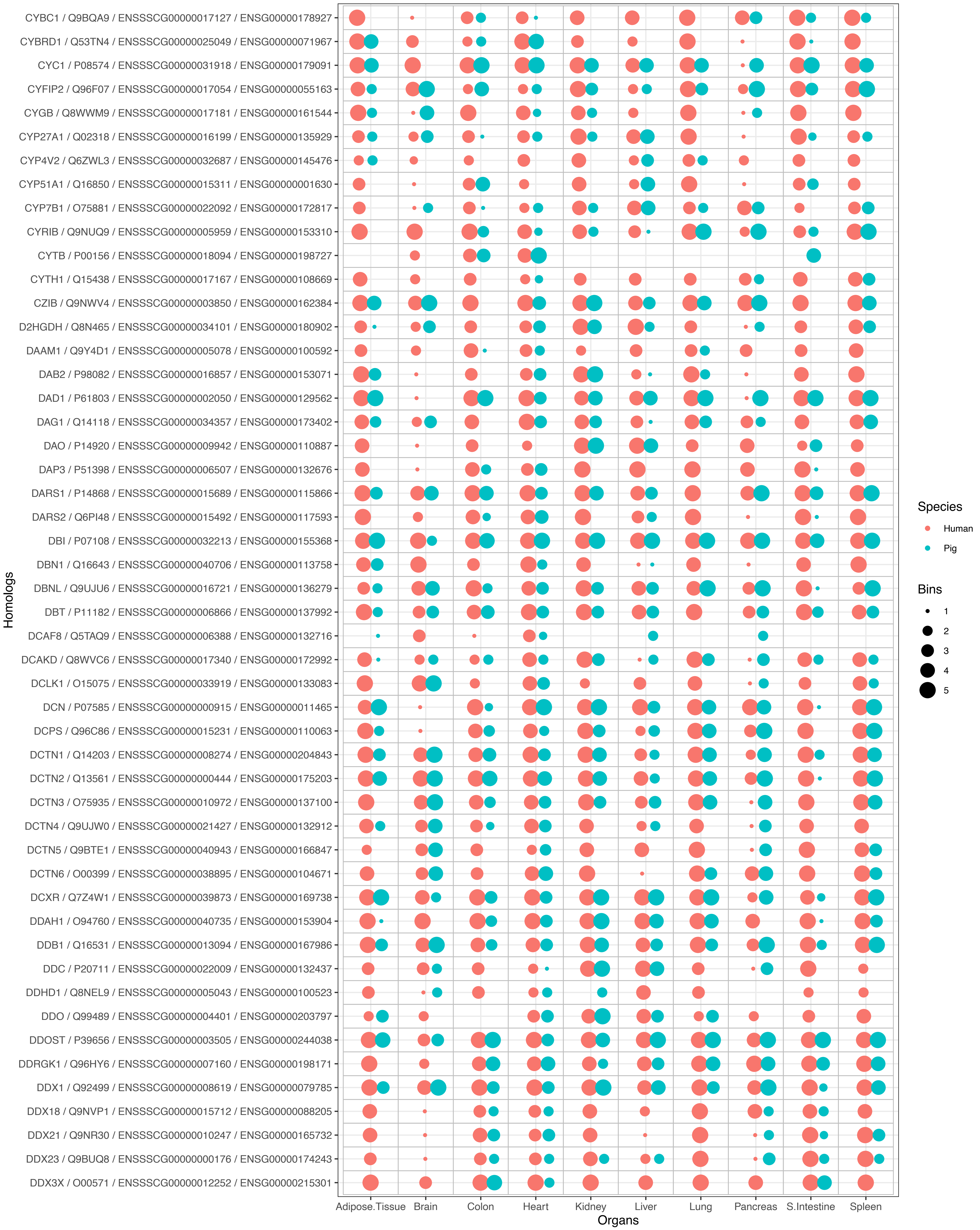

CYBC1 / Q9BQA9 / ENSSSCG00000017127 / ENSG00000178927
CYBRD1 / Q53TN4 / ENSSSCG00000025049 / ENSG00000071967
CYC1 / P08574 / ENSSSCG00000031918 / ENSG00000179091
CYFIP2 / Q96F07 / ENSSSCG00000017054 / ENSG00000055163
CYGB / Q8WWM9 / ENSSSCG00000017181 / ENSG00000161544
CYP27A1 / Q02318 / ENSSSCG00000016199 / ENSG00000135929
CYP4V2 / Q6ZWL3 / ENSSSCG00000032687 / ENSG00000145476
CYP51A1 / Q16850 / ENSSSCG00000015311 / ENSG00000001630
CYP7B1 / O75881 / ENSSSCG00000022092 / ENSG00000172817
CYRIB / Q9NUQ9 / ENSSSCG00000005959 / ENSG00000153310
CYTB / P00156 / ENSSSCG00000018094 / ENSG00000198727
CYTH1 / Q15438 / ENSSSCG00000017167 / ENSG00000108669
CZIB / Q9NWV4 / ENSSSCG00000003850 / ENSG00000162384
D2HGDH / Q8N465 / ENSSSCG00000034101 / ENSG00000180902
DAAM1 / Q9Y4D1 / ENSSSCG00000005078 / ENSG00000100592
DAB2 / P98082 / ENSSSCG00000016857 / ENSG00000153071
DAD1 / P61803 / ENSSSCG00000002050 / ENSG00000129562
DAG1 / Q14118 / ENSSSCG00000034357 / ENSG00000173402
DAO / P14920 / ENSSSCG00000009942 / ENSG00000110887
DAP3 / P51398 / ENSSSCG00000006507 / ENSG00000132676
DARS1 / P14868 / ENSSSCG00000015689 / ENSG00000115866
DARS2 / Q6PI48 / ENSSSCG00000015492 / ENSG00000117593
DBI / P07108 / ENSSSCG00000032213 / ENSG00000155368
DBN1 / Q16643 / ENSSSCG00000040706 / ENSG00000113758
DBNL / Q9UJU6 / ENSSSCG00000016721 / ENSG00000136279
DBT / P11182 / ENSSSCG00000006866 / ENSG00000137992
DCAF8 / Q5TAQ9 / ENSSSCG00000006388 / ENSG00000132716
DCAKD / Q8WVC6 / ENSSSCG00000017340 / ENSG00000172992
DCLK1 / O15075 / ENSSSCG00000033919 / ENSG00000133083
DCN / P07585 / ENSSSCG00000000915 / ENSG00000011465
DCPS / Q96C86 / ENSSSCG00000015231 / ENSG00000110063
DCTN1 / Q14203 / ENSSSCG00000008274 / ENSG00000204843
DCTN2 / Q13561 / ENSSSCG00000000444 / ENSG00000175203
DCTN3 / O75935 / ENSSSCG00000010972 / ENSG00000137100
DCTN4 / Q9UJW0 / ENSSSCG00000021427 / ENSG00000132912
DCTN5 / Q9BTE1 / ENSSSCG00000040943 / ENSG00000166847
DCTN6 / O00399 / ENSSSCG00000038895 / ENSG00000104671
DCXR / Q7Z4W1 / ENSSSCG00000039873 / ENSG00000169738
DDAH1 / O94760 / ENSSSCG00000040735 / ENSG00000153904
DDB1 / Q16531 / ENSSSCG00000013094 / ENSG00000167986
DDC / P20711 / ENSSSCG00000022009 / ENSG00000132437
DDHD1 / Q8NEL9 / ENSSSCG00000005043 / ENSG00000100523
DDO / Q99489 / ENSSSCG00000004401 / ENSG00000203797
DDOST / P39656 / ENSSSCG00000003505 / ENSG00000244038
DDRGK1 / Q96HY6 / ENSSSCG00000007160 / ENSG00000198171
DDX1 / Q92499 / ENSSSCG00000008619 / ENSG00000079785
DDX18 / Q9NVP1 / ENSSSCG00000015712 / ENSG00000088205
DDX21 / Q9NR30 / ENSSSCG00000010247 / ENSG00000165732
DDX23 / Q9BUQ8 / ENSSSCG00000000176 / ENSG00000174243
DDX3X / O00571 / ENSSSCG00000012252 / ENSG00000215301
Homologs
Adipose.Tissue
Brain
Colon
Heart
Kidney
Liver
Lung
Pancreas
S.Intestine
Spleen
Organs
Species
Human
Pig
Bins
1
2
3
4
5

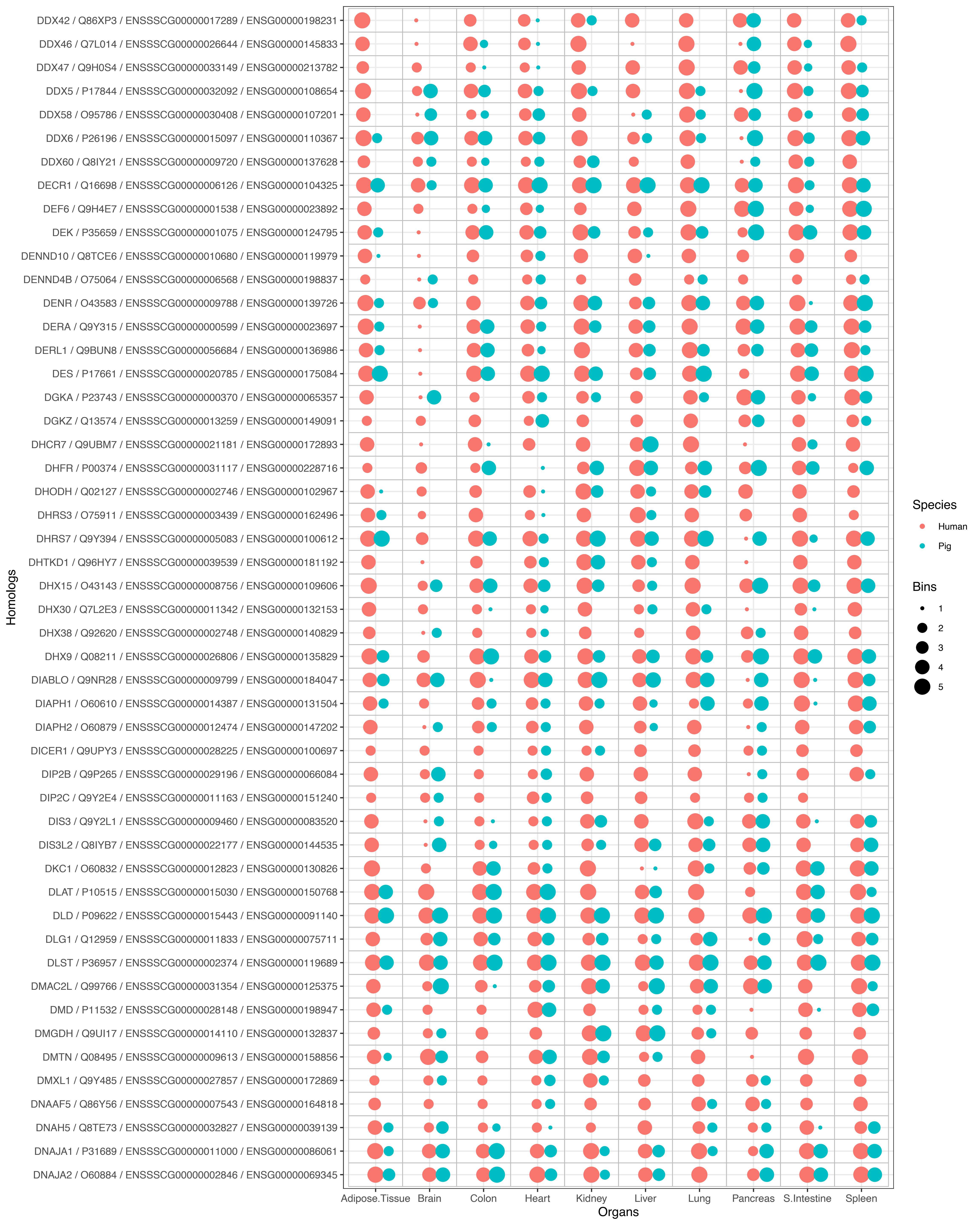
DDX42 / Q86XP3 / ENSSSCG00000017289 / ENSG00000198231
DDX46 / Q7L014 / ENSSSCG00000026644 / ENSG00000145833
DDX47 / Q9H0S4 / ENSSSCG00000033149 / ENSG00000213782
DDX5 / P17844 / ENSSSCG00000032092 / ENSG00000108654
DDX58 / O95786 / ENSSSCG00000030408 / ENSG00000107201
DDX6 / P26196 / ENSSSCG00000015097 / ENSG00000110367
DDX60 / Q8IY21 / ENSSSCG00000009720 / ENSG00000137628
DECR1 / Q16698 / ENSSSCG00000006126 / ENSG00000104325
DEF6 / Q9H4E7 / ENSSSCG00000001538 / ENSG00000023892
DEK / P35659 / ENSSSCG00000001075 / ENSG00000124795
DENND10 / Q8TCE6 / ENSSSCG00000010680 / ENSG00000119979
DENND4B / O75064 / ENSSSCG00000006568 / ENSG00000198837
DENR / O43583 / ENSSSCG00000009788 / ENSG00000139726
DERA / Q9Y315 / ENSSSCG00000000599 / ENSG00000023697
DERL1 / Q9BUN8 / ENSSSCG00000056684 / ENSG00000136986
DES / P17661 / ENSSSCG00000020785 / ENSG00000175084
DGKA / P23743 / ENSSSCG00000000370 / ENSG00000065357
DGKZ / Q13574 / ENSSSCG00000013259 / ENSG00000149091
DHCR7 / Q9UBM7 / ENSSSCG00000021181 / ENSG00000172893
DHFR / P00374 / ENSSSCG00000031117 / ENSG00000228716
DHODH / Q02127 / ENSSSCG00000002746 / ENSG00000102967
DHRS3 / O75911 / ENSSSCG00000003439 / ENSG00000162496
DHRS7 / Q9Y394 / ENSSSCG00000005083 / ENSG00000100612
DHTKD1 / Q96HY7 / ENSSSCG00000039539 / ENSG00000181192
DHX15 / O43143 / ENSSSCG00000008756 / ENSG00000109606
DHX30 / Q7L2E3 / ENSSSCG00000011342 / ENSG00000132153
DHX38 / Q92620 / ENSSSCG00000002748 / ENSG00000140829
DHX9 / Q08211 / ENSSSCG00000026806 / ENSG00000135829
DIABLO / Q9NR28 / ENSSSCG00000009799 / ENSG00000184047
DIAPH1 / O60610 / ENSSSCG00000014387 / ENSG00000131504
DIAPH2 / O60879 / ENSSSCG00000012474 / ENSG00000147202
DICER1 / Q9UPY3 / ENSSSCG00000028225 / ENSG00000100697
DIP2B / Q9P265 / ENSSSCG00000029196 / ENSG00000066084
DIP2C / Q9Y2E4 / ENSSSCG00000011163 / ENSG00000151240
DIS3 / Q9Y2L1 / ENSSSCG00000009460 / ENSG00000083520
DIS3L2 / Q8IYB7 / ENSSSCG00000022177 / ENSG00000144535
DKC1 / O60832 / ENSSSCG00000012823 / ENSG00000130826
DLAT / P10515 / ENSSSCG00000015030 / ENSG00000150768
DLD / P09622 / ENSSSCG00000015443 / ENSG00000091140
DLG1 / Q12959 / ENSSSCG00000011833 / ENSG00000075711
DLST / P36957 / ENSSSCG00000002374 / ENSG00000119689
DMAC2L / Q99766 / ENSSSCG00000031354 / ENSG00000125375
DMD / P11532 / ENSSSCG00000028148 / ENSG00000198947
DMGDH / Q9UI17 / ENSSSCG00000014110 / ENSG00000132837
DMTN / Q08495 / ENSSSCG00000009613 / ENSG00000158856
DMXL1 / Q9Y485 / ENSSSCG00000027857 / ENSG00000172869
DNAAF5 / Q86Y56 / ENSSSCG00000007543 / ENSG00000164818
DNAH5 / Q8TE73 / ENSSSCG00000032827 / ENSG00000039139
DNAJA1 / P31689 / ENSSSCG00000011000 / ENSG00000086061
DNAJA2 / O60884 / ENSSSCG00000002846 / ENSG00000069345
Homologs
Adipose.Tissue
Brain
Colon
Heart
Kidney
Liver
Lung
Pancreas
S.Intestine
Spleen
Organs
Species
Human
Pig
Bins
1
2
3
4
5

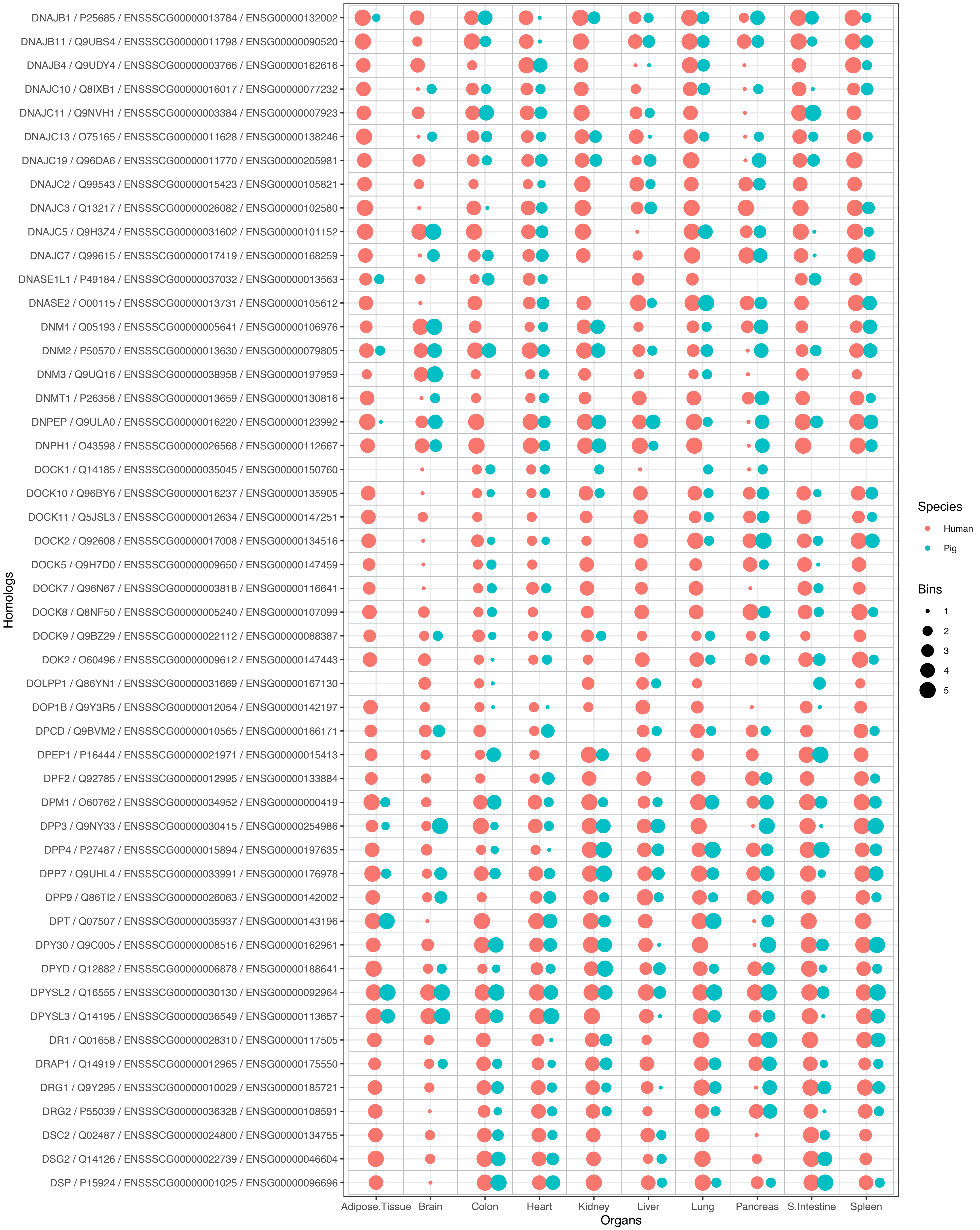
DNAJB1 / P25685 / ENSSSCG00000013784 / ENSG00000132002
DNAJB11 / Q9UBS4 / ENSSSCG00000011798 / ENSG00000090520
DNAJB4 / Q9UDY4 / ENSSSCG00000003766 / ENSG00000162616
DNAJC10 / Q8IXB1 / ENSSSCG00000016017 / ENSG00000077232
DNAJC11 / Q9NVH1 / ENSSSCG00000003384 / ENSG00000007923
DNAJC13 / O75165 / ENSSSCG00000011628 / ENSG00000138246
DNAJC19 / Q96DA6 / ENSSSCG00000011770 / ENSG00000205981
DNAJC2 / Q99543 / ENSSSCG00000015423 / ENSG00000105821
DNAJC3 / Q13217 / ENSSSCG00000026082 / ENSG00000102580
DNAJC5 / Q9H3Z4 / ENSSSCG00000031602 / ENSG00000101152
DNAJC7 / Q99615 / ENSSSCG00000017419 / ENSG00000168259
DNASE1L1 / P49184 / ENSSSCG00000037032 / ENSG00000013563
DNASE2 / O00115 / ENSSSCG00000013731 / ENSG00000105612
DNM1 / Q05193 / ENSSSCG00000005641 / ENSG00000106976
DNM2 / P50570 / ENSSSCG00000013630 / ENSG00000079805
DNM3 / Q9UQ16 / ENSSSCG00000038958 / ENSG00000197959
DNMT1 / P26358 / ENSSSCG00000013659 / ENSG00000130816
DNPEP / Q9ULA0 / ENSSSCG00000016220 / ENSG00000123992
DNPH1 / O43598 / ENSSSCG00000026568 / ENSG00000112667
DOCK1 / Q14185 / ENSSSCG00000035045 / ENSG00000150760
DOCK10 / Q96BY6 / ENSSSCG00000016237 / ENSG00000135905
DOCK11 / Q5JSL3 / ENSSSCG00000012634 / ENSG00000147251
DOCK2 / Q92608 / ENSSSCG00000017008 / ENSG00000134516
DOCK5 / Q9H7D0 / ENSSSCG00000009650 / ENSG00000147459
DOCK7 / Q96N67 / ENSSSCG00000003818 / ENSG00000116641
DOCK8 / Q8NF50 / ENSSSCG00000005240 / ENSG00000107099
DOCK9 / Q9BZ29 / ENSSSCG00000022112 / ENSG00000088387
DOK2 / O60496 / ENSSSCG00000009612 / ENSG00000147443
DOLPP1 / Q86YN1 / ENSSSCG00000031669 / ENSG00000167130
DOP1B / Q9Y3R5 / ENSSSCG00000012054 / ENSG00000142197
DPCD / Q9BVM2 / ENSSSCG00000010565 / ENSG00000166171
DPEP1 / P16444 / ENSSSCG00000021971 / ENSG00000015413
DPF2 / Q92785 / ENSSSCG00000012995 / ENSG00000133884
DPM1 / O60762 / ENSSSCG00000034952 / ENSG00000000419
DPP3 / Q9NY33 / ENSSSCG00000030415 / ENSG00000254986
DPP4 / P27487 / ENSSSCG00000015894 / ENSG00000197635
DPP7 / Q9UHL4 / ENSSSCG00000033991 / ENSG00000176978
DPP9 / Q86TI2 / ENSSSCG00000026063 / ENSG00000142002
DPT / Q07507 / ENSSSCG00000035937 / ENSG00000143196
DPY30 / Q9C005 / ENSSSCG00000008516 / ENSG00000162961
DPYD / Q12882 / ENSSSCG00000006878 / ENSG00000188641
DPYSL2 / Q16555 / ENSSSCG00000030130 / ENSG00000092964
DPYSL3 / Q14195 / ENSSSCG00000036549 / ENSG00000113657
DR1 / Q01658 / ENSSSCG00000028310 / ENSG00000117505
DRAP1 / Q14919 / ENSSSCG00000012965 / ENSG00000175550
DRG1 / Q9Y295 / ENSSSCG00000010029 / ENSG00000185721
DRG2 / P55039 / ENSSSCG00000036328 / ENSG00000108591
DSC2 / Q02487 / ENSSSCG00000024800 / ENSG00000134755
DSG2 / Q14126 / ENSSSCG00000022739 / ENSG00000046604
DSP / P15924 / ENSSSCG00000001025 / ENSG00000096696
Homologs
Adipose.Tissue
Brain
Colon
Heart
Kidney
Liver
Lung
Pancreas
S.Intestine
Spleen
Organs
Species
Human
Pig
Bins
1
2
3
4
5

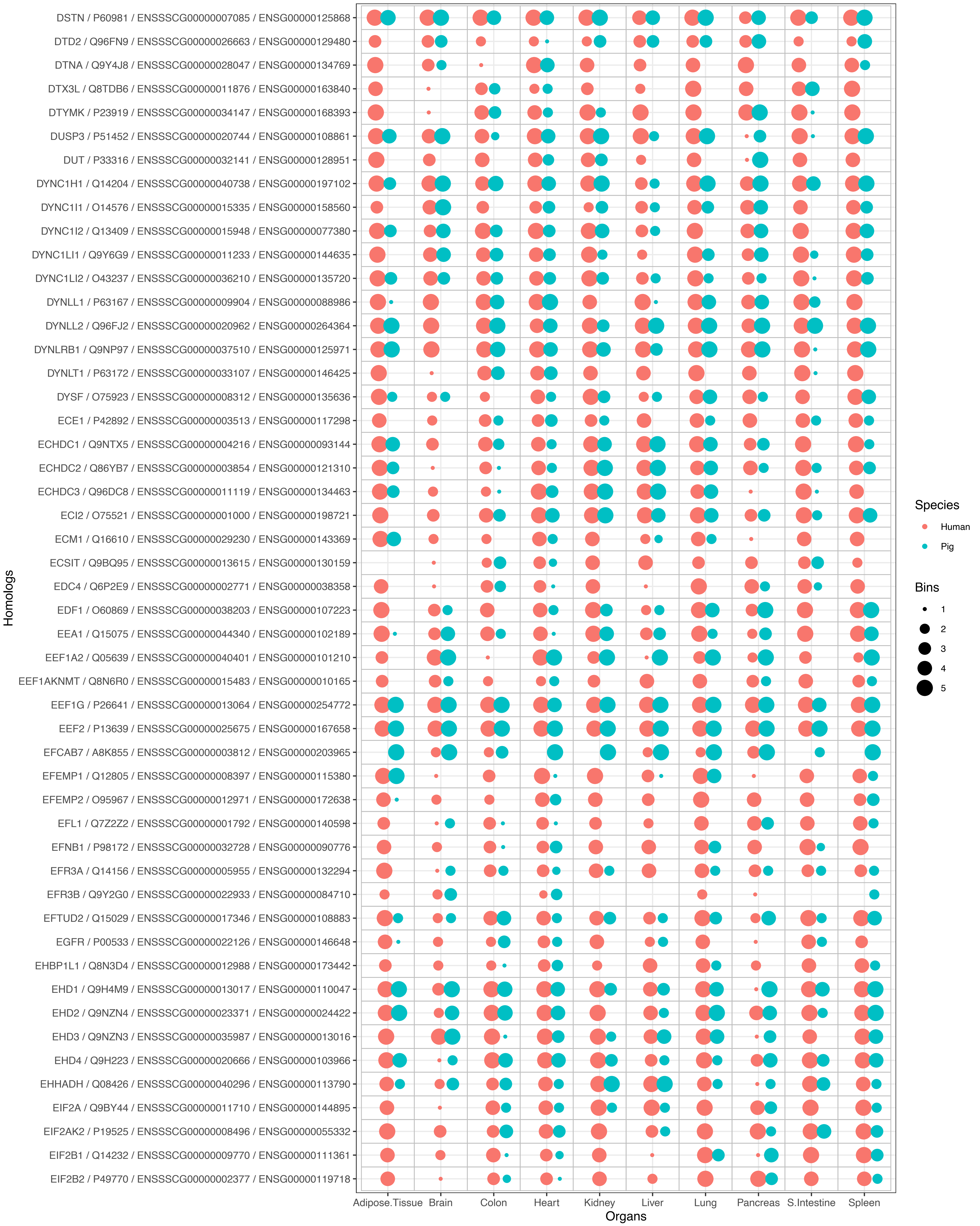
DSTN / P60981 / ENSSSCG00000007085 / ENSG00000125868
DTD2 / Q96FN9 / ENSSSCG00000026663 / ENSG00000129480
DTNA / Q9Y4J8 / ENSSSCG00000028047 / ENSG00000134769
DTX3L / Q8TDB6 / ENSSSCG00000011876 / ENSG00000163840
DTYMK / P23919 / ENSSSCG00000034147 / ENSG00000168393
DUSP3 / P51452 / ENSSSCG00000020744 / ENSG00000108861
DUT / P33316 / ENSSSCG00000032141 / ENSG00000128951
DYNC1H1 / Q14204 / ENSSSCG00000040738 / ENSG00000197102
DYNC1I1 / O14576 / ENSSSCG00000015335 / ENSG00000158560
DYNC1I2 / Q13409 / ENSSSCG00000015948 / ENSG00000077380
DYNC1LI1 / Q9Y6G9 / ENSSSCG00000011233 / ENSG00000144635
DYNC1LI2 / O43237 / ENSSSCG00000036210 / ENSG00000135720
DYNLL1 / P63167 / ENSSSCG00000009904 / ENSG00000088986
DYNLL2 / Q96FJ2 / ENSSSCG00000020962 / ENSG00000264364
DYNLRB1 / Q9NP97 / ENSSSCG00000037510 / ENSG00000125971
DYNLT1 / P63172 / ENSSSCG00000033107 / ENSG00000146425
DYSF / O75923 / ENSSSCG00000008312 / ENSG00000135636
ECE1 / P42892 / ENSSSCG00000003513 / ENSG00000117298
ECHDC1 / Q9NTX5 / ENSSSCG00000004216 / ENSG00000093144
ECHDC2 / Q86YB7 / ENSSSCG00000003854 / ENSG00000121310
ECHDC3 / Q96DC8 / ENSSSCG00000011119 / ENSG00000134463
ECI2 / O75521 / ENSSSCG00000001000 / ENSG00000198721
ECM1 / Q16610 / ENSSSCG00000029230 / ENSG00000143369
ECSIT / Q9BQ95 / ENSSSCG00000013615 / ENSG00000130159
EDC4 / Q6P2E9 / ENSSSCG00000002771 / ENSG00000038358
EDF1 / O60869 / ENSSSCG00000038203 / ENSG00000107223
EEA1 / Q15075 / ENSSSCG00000044340 / ENSG00000102189
EEF1A2 / Q05639 / ENSSSCG00000040401 / ENSG00000101210
EEF1AKNMT / Q8N6R0 / ENSSSCG00000015483 / ENSG00000010165
EEF1G / P26641 / ENSSSCG00000013064 / ENSG00000254772
EEF2 / P13639 / ENSSSCG00000025675 / ENSG00000167658
EFCAB7 / A8K855 / ENSSSCG00000003812 / ENSG00000203965
EFEMP1 / Q12805 / ENSSSCG00000008397 / ENSG00000115380
EFEMP2 / O95967 / ENSSSCG00000012971 / ENSG00000172638
EFL1 / Q7Z2Z2 / ENSSSCG00000001792 / ENSG00000140598
EFNB1 / P98172 / ENSSSCG00000032728 / ENSG00000090776
EFR3A / Q14156 / ENSSSCG00000005955 / ENSG00000132294
EFR3B / Q9Y2G0 / ENSSSCG00000022933 / ENSG00000084710
EFTUD2 / Q15029 / ENSSSCG00000017346 / ENSG00000108883
EGFR / P00533 / ENSSSCG00000022126 / ENSG00000146648
EHBP1L1 / Q8N3D4 / ENSSSCG00000012988 / ENSG00000173442
EHD1 / Q9H4M9 / ENSSSCG00000013017 / ENSG00000110047
EHD2 / Q9NZN4 / ENSSSCG00000023371 / ENSG00000024422
EHD3 / Q9NZN3 / ENSSSCG00000035987 / ENSG00000013016
EHD4 / Q9H223 / ENSSSCG00000020666 / ENSG00000103966
EHHADH / Q08426 / ENSSSCG00000040296 / ENSG00000113790
EIF2A / Q9BY44 / ENSSSCG00000011710 / ENSG00000144895
EIF2AK2 / P19525 / ENSSSCG00000008496 / ENSG00000055332
EIF2B1 / Q14232 / ENSSSCG00000009770 / ENSG00000111361
EIF2B2 / P49770 / ENSSSCG00000002377 / ENSG00000119718
Homologs
Adipose.Tissue
Brain
Colon
Heart
Kidney
Liver
Lung
Pancreas
S.Intestine
Spleen
Organs
Species
Human
Pig
Bins
1
2
3
4
5

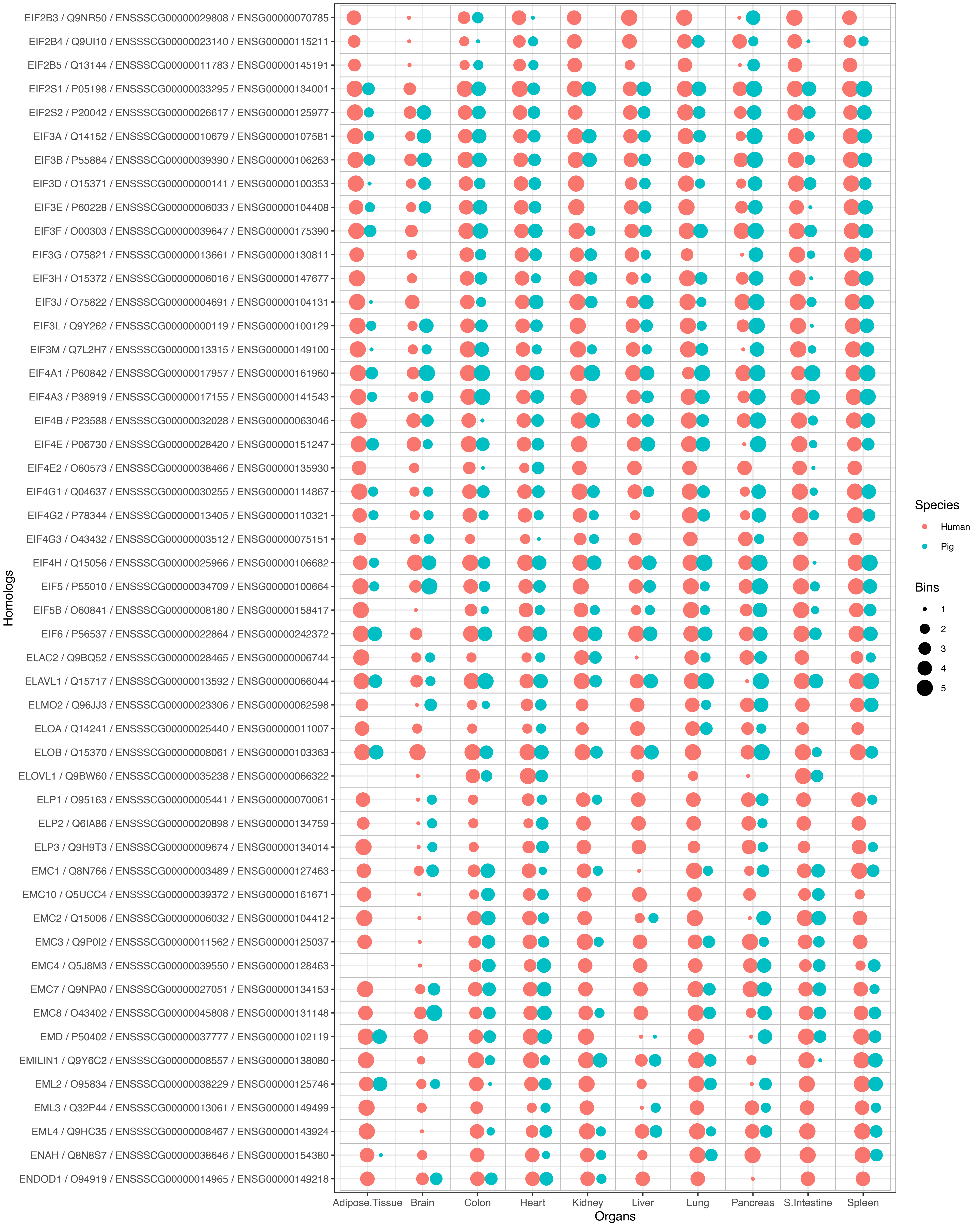

EIF2B3 / Q9NR50 / ENSSSCG00000029808 / ENSG00000070785
EIF2B4 / Q9UI10 / ENSSSCG00000023140 / ENSG00000115211
EIF2B5 / Q13144 / ENSSSCG00000011783 / ENSG00000145191
EIF2S1 / P05198 / ENSSSCG00000033295 / ENSG00000134001
EIF2S2 / P20042 / ENSSSCG00000026617 / ENSG00000125977
EIF3A / Q14152 / ENSSSCG00000010679 / ENSG00000107581
EIF3B / P55884 / ENSSSCG00000039390 / ENSG00000106263
EIF3D / O15371 / ENSSSCG00000000141 / ENSG00000100353
EIF3E / P60228 / ENSSSCG00000006033 / ENSG00000104408
EIF3F / O00303 / ENSSSCG00000039647 / ENSG00000175390
EIF3G / O75821 / ENSSSCG00000013661 / ENSG00000130811
EIF3H / O15372 / ENSSSCG00000006016 / ENSG00000147677
EIF3J / O75822 / ENSSSCG00000004691 / ENSG00000104131
EIF3L / Q9Y262 / ENSSSCG00000000119 / ENSG00000100129
EIF3M / Q7L2H7 / ENSSSCG00000013315 / ENSG00000149100
EIF4A1 / P60842 / ENSSSCG00000017957 / ENSG00000161960
EIF4A3 / P38919 / ENSSSCG00000017155 / ENSG00000141543
EIF4B / P23588 / ENSSSCG00000032028 / ENSG00000063046
EIF4E / P06730 / ENSSSCG00000028420 / ENSG00000151247
EIF4E2 / O60573 / ENSSSCG00000038466 / ENSG00000135930
EIF4G1 / Q04637 / ENSSSCG00000030255 / ENSG00000114867
EIF4G2 / P78344 / ENSSSCG00000013405 / ENSG00000110321
EIF4G3 / O43432 / ENSSSCG00000003512 / ENSG00000075151
EIF4H / Q15056 / ENSSSCG00000025966 / ENSG00000106682
EIF5 / P55010 / ENSSSCG00000034709 / ENSG00000100664
EIF5B / O60841 / ENSSSCG00000008180 / ENSG00000158417
EIF6 / P56537 / ENSSSCG00000022864 / ENSG00000242372
ELAC2 / Q9BQ52 / ENSSSCG00000028465 / ENSG00000006744
ELAVL1 / Q15717 / ENSSSCG00000013592 / ENSG00000066044
ELMO2 / Q96JJ3 / ENSSSCG00000023306 / ENSG00000062598
ELOA / Q14241 / ENSSSCG00000025440 / ENSG00000011007
ELOB / Q15370 / ENSSSCG00000008061 / ENSG00000103363
ELOVL1 / Q9BW60 / ENSSSCG00000035238 / ENSG00000066322
ELP1 / O95163 / ENSSSCG00000005441 / ENSG00000070061
ELP2 / Q6IA86 / ENSSSCG00000020898 / ENSG00000134759
ELP3 / Q9H9T3 / ENSSSCG00000009674 / ENSG00000134014
EMC1 / Q8N766 / ENSSSCG00000003489 / ENSG00000127463
EMC10 / Q5UCC4 / ENSSSCG00000039372 / ENSG00000161671
EMC2 / Q15006 / ENSSSCG00000006032 / ENSG00000104412
EMC3 / Q9P0I2 / ENSSSCG00000011562 / ENSG00000125037
EMC4 / Q5J8M3 / ENSSSCG00000039550 / ENSG00000128463
EMC7 / Q9NPA0 / ENSSSCG00000027051 / ENSG00000134153
EMC8 / O43402 / ENSSSCG00000045808 / ENSG00000131148
EMD / P50402 / ENSSSCG00000037777 / ENSG00000102119
EMILIN1 / Q9Y6C2 / ENSSSCG00000008557 / ENSG00000138080
EML2 / O95834 / ENSSSCG00000038229 / ENSG00000125746
EML3 / Q32P44 / ENSSSCG00000013061 / ENSG00000149499
EML4 / Q9HC35 / ENSSSCG00000008467 / ENSG00000143924
ENAH / Q8N8S7 / ENSSSCG00000038646 / ENSG00000154380
ENDOD1 / O94919 / ENSSSCG00000014965 / ENSG00000149218
Homologs
Adipose.Tissue
Brain
Colon
Heart
Kidney
Liver
Lung
Pancreas
S.Intestine
Spleen
Organs
Species
Human
Pig
Bins
1
2
3
4
5

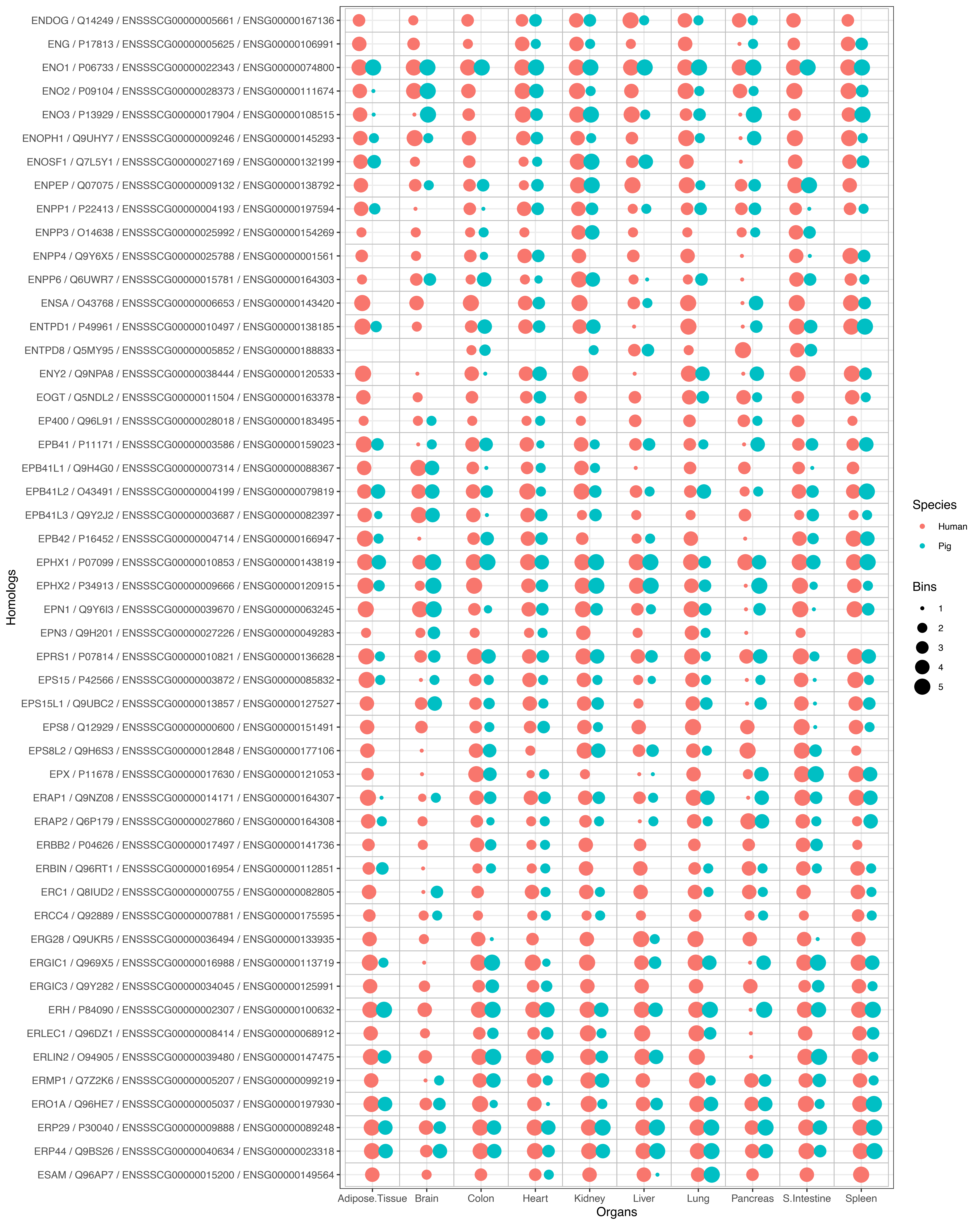
ENDOG / Q14249 / ENSSSCG00000005661 / ENSG00000167136
ENG / P17813 / ENSSSCG00000005625 / ENSG00000106991
ENO1 / P06733 / ENSSSCG00000022343 / ENSG00000074800
ENO2 / P09104 / ENSSSCG00000028373 / ENSG00000111674
ENO3 / P13929 / ENSSSCG00000017904 / ENSG00000108515
ENOPH1 / Q9UHY7 / ENSSSCG00000009246 / ENSG00000145293
ENOSF1 / Q7L5Y1 / ENSSSCG00000027169 / ENSG00000132199
ENPEP / Q07075 / ENSSSCG00000009132 / ENSG00000138792
ENPP1 / P22413 / ENSSSCG00000004193 / ENSG00000197594
ENPP3 / O14638 / ENSSSCG00000025992 / ENSG00000154269
ENPP4 / Q9Y6X5 / ENSSSCG00000025788 / ENSG00000001561
ENPP6 / Q6UWR7 / ENSSSCG00000015781 / ENSG00000164303
ENSA / O43768 / ENSSSCG00000006653 / ENSG00000143420
ENTPD1 / P49961 / ENSSSCG00000010497 / ENSG00000138185
ENTPD8 / Q5MY95 / ENSSSCG00000005852 / ENSG00000188833
ENY2 / Q9NPA8 / ENSSSCG00000038444 / ENSG00000120533
EOGT / Q5NDL2 / ENSSSCG00000011504 / ENSG00000163378
EP400 / Q96L91 / ENSSSCG00000028018 / ENSG00000183495
EPB41 / P11171 / ENSSSCG00000003586 / ENSG00000159023
EPB41L1 / Q9H4G0 / ENSSSCG00000007314 / ENSG00000088367
EPB41L2 / O43491 / ENSSSCG00000004199 / ENSG00000079819
EPB41L3 / Q9Y2J2 / ENSSSCG00000003687 / ENSG00000082397
EPB42 / P16452 / ENSSSCG00000004714 / ENSG00000166947
EPHX1 / P07099 / ENSSSCG00000010853 / ENSG00000143819
EPHX2 / P34913 / ENSSSCG00000009666 / ENSG00000120915
EPN1 / Q9Y6I3 / ENSSSCG00000039670 / ENSG00000063245
EPN3 / Q9H201 / ENSSSCG00000027226 / ENSG00000049283
EPRS1 / P07814 / ENSSSCG00000010821 / ENSG00000136628
EPS15 / P42566 / ENSSSCG00000003872 / ENSG00000085832
EPS15L1 / Q9UBC2 / ENSSSCG00000013857 / ENSG00000127527
EPS8 / Q12929 / ENSSSCG00000000600 / ENSG00000151491
EPS8L2 / Q9H6S3 / ENSSSCG00000012848 / ENSG00000177106
EPX / P11678 / ENSSSCG00000017630 / ENSG00000121053
ERAP1 / Q9NZ08 / ENSSSCG00000014171 / ENSG00000164307
ERAP2 / Q6P179 / ENSSSCG00000027860 / ENSG00000164308
ERBB2 / P04626 / ENSSSCG00000017497 / ENSG00000141736
ERBIN / Q96RT1 / ENSSSCG00000016954 / ENSG00000112851
ERC1 / Q8IUD2 / ENSSSCG00000000755 / ENSG00000082805
ERCC4 / Q92889 / ENSSSCG00000007881 / ENSG00000175595
ERG28 / Q9UKR5 / ENSSSCG00000036494 / ENSG00000133935
ERGIC1 / Q969X5 / ENSSSCG00000016988 / ENSG00000113719
ERGIC3 / Q9Y282 / ENSSSCG00000034045 / ENSG00000125991
ERH / P84090 / ENSSSCG00000002307 / ENSG00000100632
ERLEC1 / Q96DZ1 / ENSSSCG00000008414 / ENSG00000068912
ERLIN2 / O94905 / ENSSSCG00000039480 / ENSG00000147475
ERMP1 / Q7Z2K6 / ENSSSCG00000005207 / ENSG00000099219
ERO1A / Q96HE7 / ENSSSCG00000005037 / ENSG00000197930
ERP29 / P30040 / ENSSSCG00000009888 / ENSG00000089248
ERP44 / Q9BS26 / ENSSSCG00000040634 / ENSG00000023318
ESAM / Q96AP7 / ENSSSCG00000015200 / ENSG00000149564
Homologs
Adipose.Tissue
Brain
Colon
Heart
Kidney
Liver
Lung
Pancreas
S.Intestine
Spleen
Organs
Species
Human
Pig
Bins
1
2
3
4
5

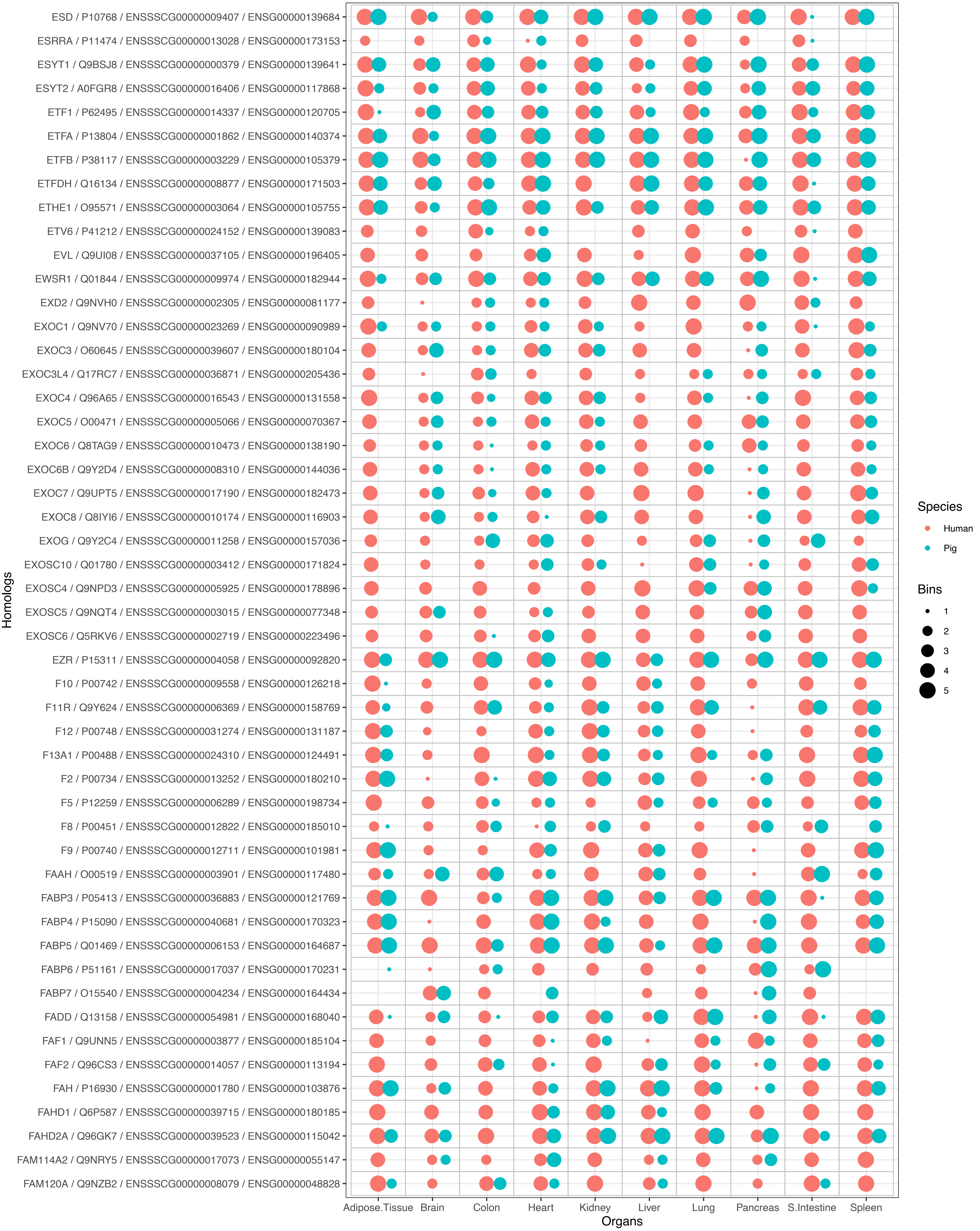
ESD / P10768 / ENSSSCG00000009407 / ENSG00000139684
ESRRA / P11474 / ENSSSCG00000013028 / ENSG00000173153
ESYT1 / Q9BSJ8 / ENSSSCG00000000379 / ENSG00000139641
ESYT2 / A0FGR8 / ENSSSCG00000016406 / ENSG00000117868
ETF1 / P62495 / ENSSSCG00000014337 / ENSG00000120705
ETFA / P13804 / ENSSSCG00000001862 / ENSG00000140374
ETFB / P38117 / ENSSSCG00000003229 / ENSG00000105379
ETFDH / Q16134 / ENSSSCG00000008877 / ENSG00000171503
ETHE1 / O95571 / ENSSSCG00000003064 / ENSG00000105755
ETV6 / P41212 / ENSSSCG00000024152 / ENSG00000139083
EVL / Q9UI08 / ENSSSCG00000037105 / ENSG00000196405
EWSR1 / Q01844 / ENSSSCG00000009974 / ENSG00000182944
EXD2 / Q9NVH0 / ENSSSCG00000002305 / ENSG00000081177
EXOC1 / Q9NV70 / ENSSSCG00000023269 / ENSG00000090989
EXOC3 / O60645 / ENSSSCG00000039607 / ENSG00000180104
EXOC3L4 / Q17RC7 / ENSSSCG00000036871 / ENSG00000205436
EXOC4 / Q96A65 / ENSSSCG00000016543 / ENSG00000131558
EXOC5 / O00471 / ENSSSCG00000005066 / ENSG00000070367
EXOC6 / Q8TAG9 / ENSSSCG00000010473 / ENSG00000138190
EXOC6B / Q9Y2D4 / ENSSSCG00000008310 / ENSG00000144036
EXOC7 / Q9UPT5 / ENSSSCG00000017190 / ENSG00000182473
EXOC8 / Q8IYI6 / ENSSSCG00000010174 / ENSG00000116903
EXOG / Q9Y2C4 / ENSSSCG00000011258 / ENSG00000157036
EXOSC10 / Q01780 / ENSSSCG00000003412 / ENSG00000171824
EXOSC4 / Q9NPD3 / ENSSSCG00000005925 / ENSG00000178896
EXOSC5 / Q9NQT4 / ENSSSCG00000003015 / ENSG00000077348
EXOSC6 / Q5RKV6 / ENSSSCG00000002719 / ENSG00000223496
EZR / P15311 / ENSSSCG00000004058 / ENSG00000092820
F10 / P00742 / ENSSSCG00000009558 / ENSG00000126218
F11R / Q9Y624 / ENSSSCG00000006369 / ENSG00000158769
F12 / P00748 / ENSSSCG00000031274 / ENSG00000131187
F13A1 / P00488 / ENSSSCG00000024310 / ENSG00000124491
F2 / P00734 / ENSSSCG00000013252 / ENSG00000180210
F5 / P12259 / ENSSSCG00000006289 / ENSG00000198734
F8 / P00451 / ENSSSCG00000012822 / ENSG00000185010
F9 / P00740 / ENSSSCG00000012711 / ENSG00000101981
FAAH / O00519 / ENSSSCG00000003901 / ENSG00000117480
FABP3 / P05413 / ENSSSCG00000036883 / ENSG00000121769
FABP4 / P15090 / ENSSSCG00000040681 / ENSG00000170323
FABP5 / Q01469 / ENSSSCG00000006153 / ENSG00000164687
FABP6 / P51161 / ENSSSCG00000017037 / ENSG00000170231
FABP7 / O15540 / ENSSSCG00000004234 / ENSG00000164434
FADD / Q13158 / ENSSSCG00000054981 / ENSG00000168040
FAF1 / Q9UNN5 / ENSSSCG00000003877 / ENSG00000185104
FAF2 / Q96CS3 / ENSSSCG00000014057 / ENSG00000113194
FAH / P16930 / ENSSSCG00000001780 / ENSG00000103876
FAHD1 / Q6P587 / ENSSSCG00000039715 / ENSG00000180185
FAHD2A / Q96GK7 / ENSSSCG00000039523 / ENSG00000115042
FAM114A2 / Q9NRY5 / ENSSSCG00000017073 / ENSG00000055147
FAM120A / Q9NZB2 / ENSSSCG00000008079 / ENSG00000048828
Homologs
Adipose.Tissue
Brain
Colon
Heart
Kidney
Liver
Lung
Pancreas
S.Intestine
Spleen
Organs
Species
Human
Pig
Bins
1
2
3
4
5

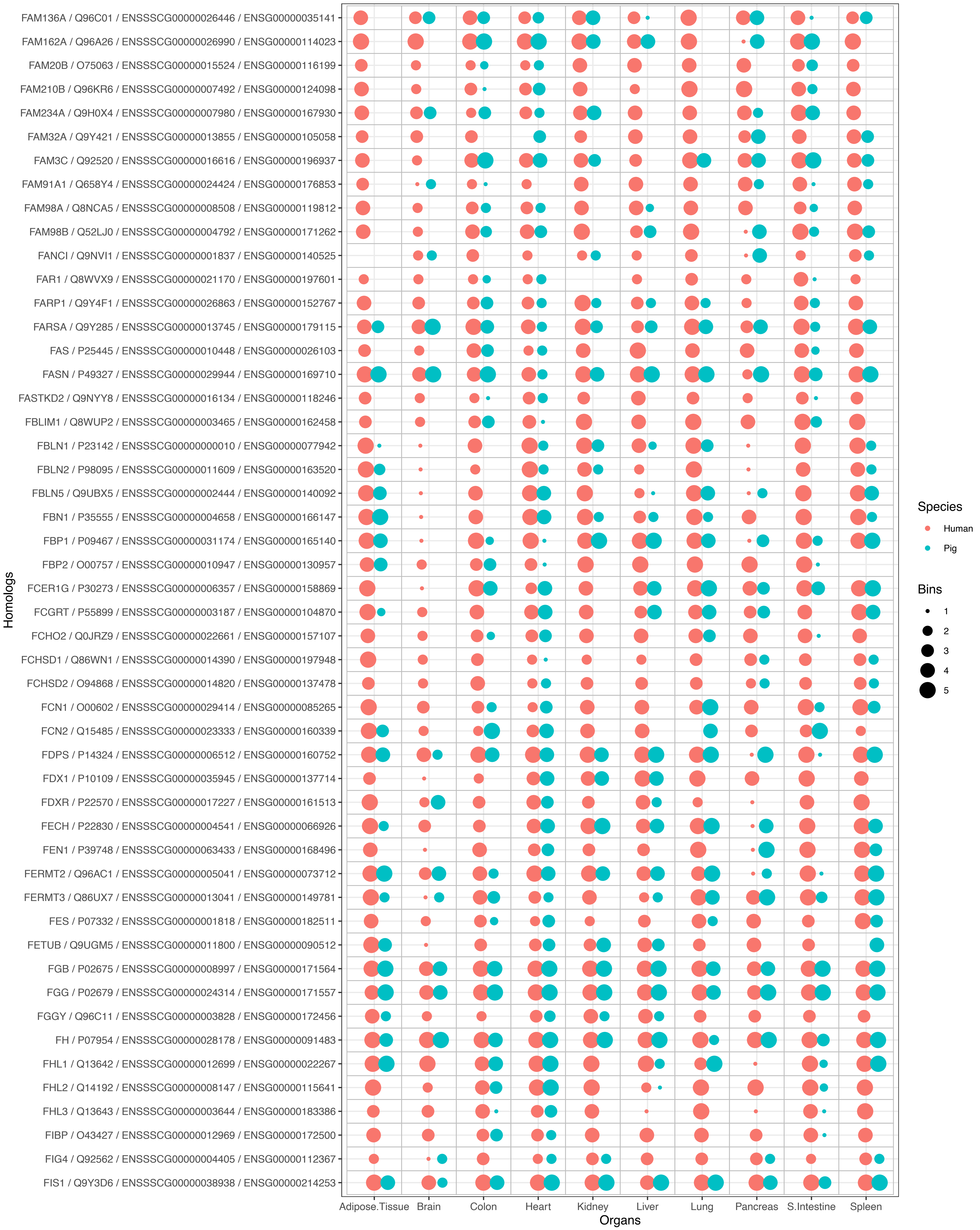
FAM136A / Q96C01 / ENSSSCG00000026446 / ENSG00000035141
FAM162A / Q96A26 / ENSSSCG00000026990 / ENSG00000114023
FAM20B / O75063 / ENSSSCG00000015524 / ENSG00000116199
FAM210B / Q96KR6 / ENSSSCG00000007492 / ENSG00000124098
FAM234A / Q9H0X4 / ENSSSCG00000007980 / ENSG00000167930
FAM32A / Q9Y421 / ENSSSCG00000013855 / ENSG00000105058
FAM3C / Q92520 / ENSSSCG00000016616 / ENSG00000196937
FAM91A1 / Q658Y4 / ENSSSCG00000024424 / ENSG00000176853
FAM98A / Q8NCA5 / ENSSSCG00000008508 / ENSG00000119812
FAM98B / Q52LJ0 / ENSSSCG00000004792 / ENSG00000171262
FANCI / Q9NVI1 / ENSSSCG00000001837 / ENSG00000140525
FAR1 / Q8WVX9 / ENSSSCG00000021170 / ENSG00000197601
FARP1 / Q9Y4F1 / ENSSSCG00000026863 / ENSG00000152767
FARSA / Q9Y285 / ENSSSCG00000013745 / ENSG00000179115
FAS / P25445 / ENSSSCG00000010448 / ENSG00000026103
FASN / P49327 / ENSSSCG00000029944 / ENSG00000169710
FASTKD2 / Q9NYY8 / ENSSSCG00000016134 / ENSG00000118246
FBLIM1 / Q8WUP2 / ENSSSCG00000003465 / ENSG00000162458
FBLN1 / P23142 / ENSSSCG00000000010 / ENSG00000077942
FBLN2 / P98095 / ENSSSCG00000011609 / ENSG00000163520
FBLN5 / Q9UBX5 / ENSSSCG00000002444 / ENSG00000140092
FBN1 / P35555 / ENSSSCG00000004658 / ENSG00000166147
FBP1 / P09467 / ENSSSCG00000031174 / ENSG00000165140
FBP2 / O00757 / ENSSSCG00000010947 / ENSG00000130957
FCER1G / P30273 / ENSSSCG00000006357 / ENSG00000158869
FCGRT / P55899 / ENSSSCG00000003187 / ENSG00000104870
FCHO2 / Q0JRZ9 / ENSSSCG00000022661 / ENSG00000157107
FCHSD1 / Q86WN1 / ENSSSCG00000014390 / ENSG00000197948
FCHSD2 / O94868 / ENSSSCG00000014820 / ENSG00000137478
FCN1 / O00602 / ENSSSCG00000029414 / ENSG00000085265
FCN2 / Q15485 / ENSSSCG00000023333 / ENSG00000160339
FDPS / P14324 / ENSSSCG00000006512 / ENSG00000160752
FDX1 / P10109 / ENSSSCG00000035945 / ENSG00000137714
FDXR / P22570 / ENSSSCG00000017227 / ENSG00000161513
FECH / P22830 / ENSSSCG00000004541 / ENSG00000066926
FEN1 / P39748 / ENSSSCG00000063433 / ENSG00000168496
FERMT2 / Q96AC1 / ENSSSCG00000005041 / ENSG00000073712
FERMT3 / Q86UX7 / ENSSSCG00000013041 / ENSG00000149781
FES / P07332 / ENSSSCG00000001818 / ENSG00000182511
FETUB / Q9UGM5 / ENSSSCG00000011800 / ENSG00000090512
FGB / P02675 / ENSSSCG00000008997 / ENSG00000171564
FGG / P02679 / ENSSSCG00000024314 / ENSG00000171557
FGGY / Q96C11 / ENSSSCG00000003828 / ENSG00000172456
FH / P07954 / ENSSSCG00000028178 / ENSG00000091483
FHL1 / Q13642 / ENSSSCG00000012699 / ENSG00000022267
FHL2 / Q14192 / ENSSSCG00000008147 / ENSG00000115641
FHL3 / Q13643 / ENSSSCG00000003644 / ENSG00000183386
FIBP / O43427 / ENSSSCG00000012969 / ENSG00000172500
FIG4 / Q92562 / ENSSSCG00000004405 / ENSG00000112367
FIS1 / Q9Y3D6 / ENSSSCG00000038938 / ENSG00000214253
Homologs
Adipose.Tissue
Brain
Colon
Heart
Kidney
Liver
Lung
Pancreas
S.Intestine
Spleen
Organs
Species
Human
Pig
Bins
1
2
3
4
5

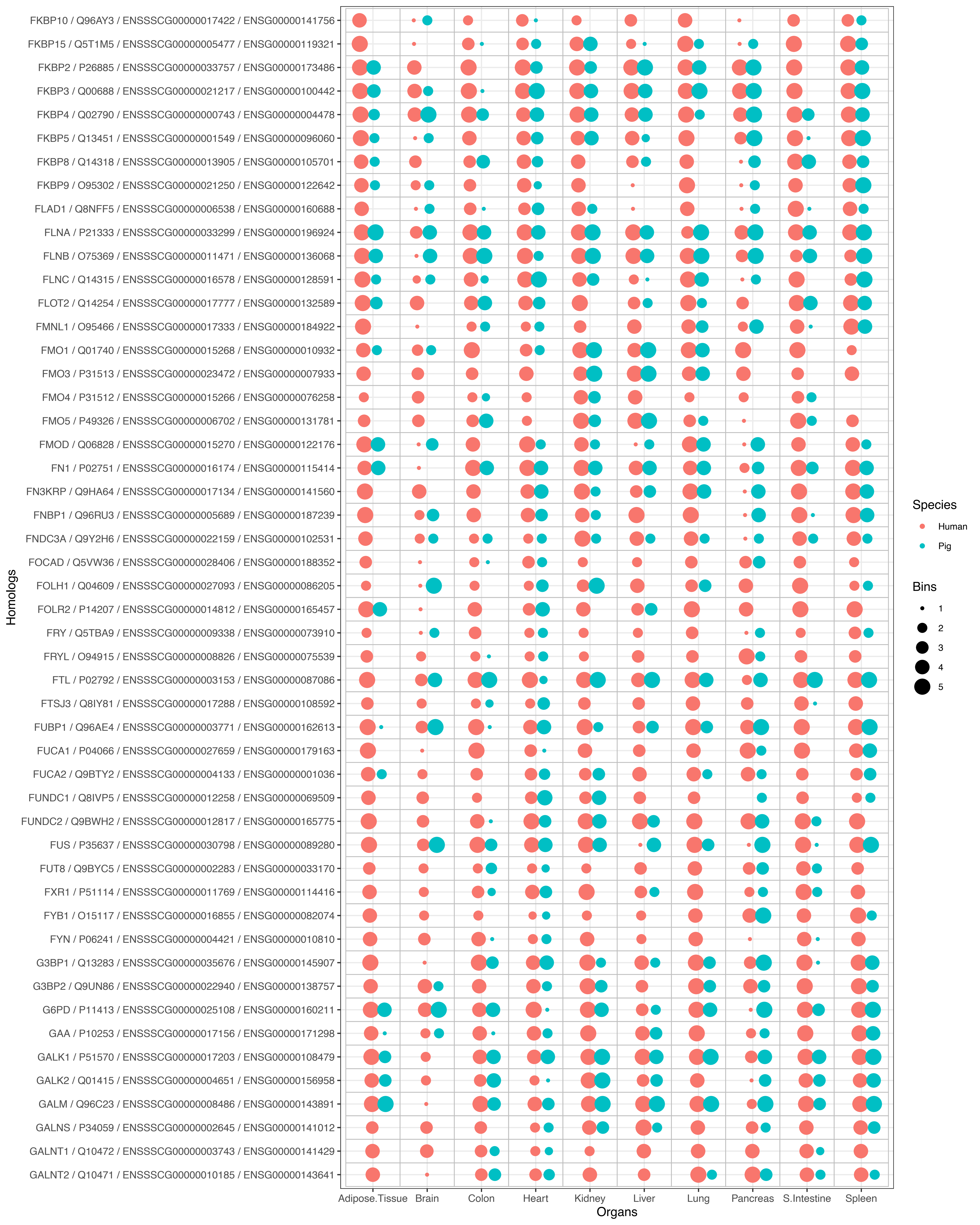

FKBP10 / Q96AY3 / ENSSSCG00000017422 / ENSG00000141756
FKBP15 / Q5T1M5 / ENSSSCG00000005477 / ENSG00000119321
FKBP2 / P26885 / ENSSSCG00000033757 / ENSG00000173486
FKBP3 / Q00688 / ENSSSCG00000021217 / ENSG00000100442
FKBP4 / Q02790 / ENSSSCG00000000743 / ENSG00000004478
FKBP5 / Q13451 / ENSSSCG00000001549 / ENSG00000096060
FKBP8 / Q14318 / ENSSSCG00000013905 / ENSG00000105701
FKBP9 / O95302 / ENSSSCG00000021250 / ENSG00000122642
FLAD1 / Q8NFF5 / ENSSSCG00000006538 / ENSG00000160688
FLNA / P21333 / ENSSSCG00000033299 / ENSG00000196924
FLNB / O75369 / ENSSSCG00000011471 / ENSG00000136068
FLNC / Q14315 / ENSSSCG00000016578 / ENSG00000128591
FLOT2 / Q14254 / ENSSSCG00000017777 / ENSG00000132589
FMNL1 / O95466 / ENSSSCG00000017333 / ENSG00000184922
FMO1 / Q01740 / ENSSSCG00000015268 / ENSG00000010932
FMO3 / P31513 / ENSSSCG00000023472 / ENSG00000007933
FMO4 / P31512 / ENSSSCG00000015266 / ENSG00000076258
FMO5 / P49326 / ENSSSCG00000006702 / ENSG00000131781
FMOD / Q06828 / ENSSSCG00000015270 / ENSG00000122176
FN1 / P02751 / ENSSSCG00000016174 / ENSG00000115414
FN3KRP / Q9HA64 / ENSSSCG00000017134 / ENSG00000141560
FNBP1 / Q96RU3 / ENSSSCG00000005689 / ENSG00000187239
FNDC3A / Q9Y2H6 / ENSSSCG00000022159 / ENSG00000102531
FOCAD / Q5VW36 / ENSSSCG00000028406 / ENSG00000188352
FOLH1 / Q04609 / ENSSSCG00000027093 / ENSG00000086205
FOLR2 / P14207 / ENSSSCG00000014812 / ENSG00000165457
FRY / Q5TBA9 / ENSSSCG00000009338 / ENSG00000073910
FRYL / O94915 / ENSSSCG00000008826 / ENSG00000075539
FTL / P02792 / ENSSSCG00000003153 / ENSG00000087086
FTSJ3 / Q8IY81 / ENSSSCG00000017288 / ENSG00000108592
FUBP1 / Q96AE4 / ENSSSCG00000003771 / ENSG00000162613
FUCA1 / P04066 / ENSSSCG00000027659 / ENSG00000179163
FUCA2 / Q9BTY2 / ENSSSCG00000004133 / ENSG00000001036
FUNDC1 / Q8IVP5 / ENSSSCG00000012258 / ENSG00000069509
FUNDC2 / Q9BWH2 / ENSSSCG00000012817 / ENSG00000165775
FUS / P35637 / ENSSSCG00000030798 / ENSG00000089280
FUT8 / Q9BYC5 / ENSSSCG00000002283 / ENSG00000033170
FXR1 / P51114 / ENSSSCG00000011769 / ENSG00000114416
FYB1 / O15117 / ENSSSCG00000016855 / ENSG00000082074
FYN / P06241 / ENSSSCG00000004421 / ENSG00000010810
G3BP1 / Q13283 / ENSSSCG00000035676 / ENSG00000145907
G3BP2 / Q9UN86 / ENSSSCG00000022940 / ENSG00000138757
G6PD / P11413 / ENSSSCG00000025108 / ENSG00000160211
GAA / P10253 / ENSSSCG00000017156 / ENSG00000171298
GALK1 / P51570 / ENSSSCG00000017203 / ENSG00000108479
GALK2 / Q01415 / ENSSSCG00000004651 / ENSG00000156958
GALM / Q96C23 / ENSSSCG00000008486 / ENSG00000143891
GALNS / P34059 / ENSSSCG00000002645 / ENSG00000141012
GALNT1 / Q10472 / ENSSSCG00000003743 / ENSG00000141429
GALNT2 / Q10471 / ENSSSCG00000010185 / ENSG00000143641
Homologs
Adipose.Tissue
Brain
Colon
Heart
Kidney
Liver
Lung
Pancreas
S.Intestine
Spleen
Organs
Species
Human
Pig
Bins
1
2
3
4
5

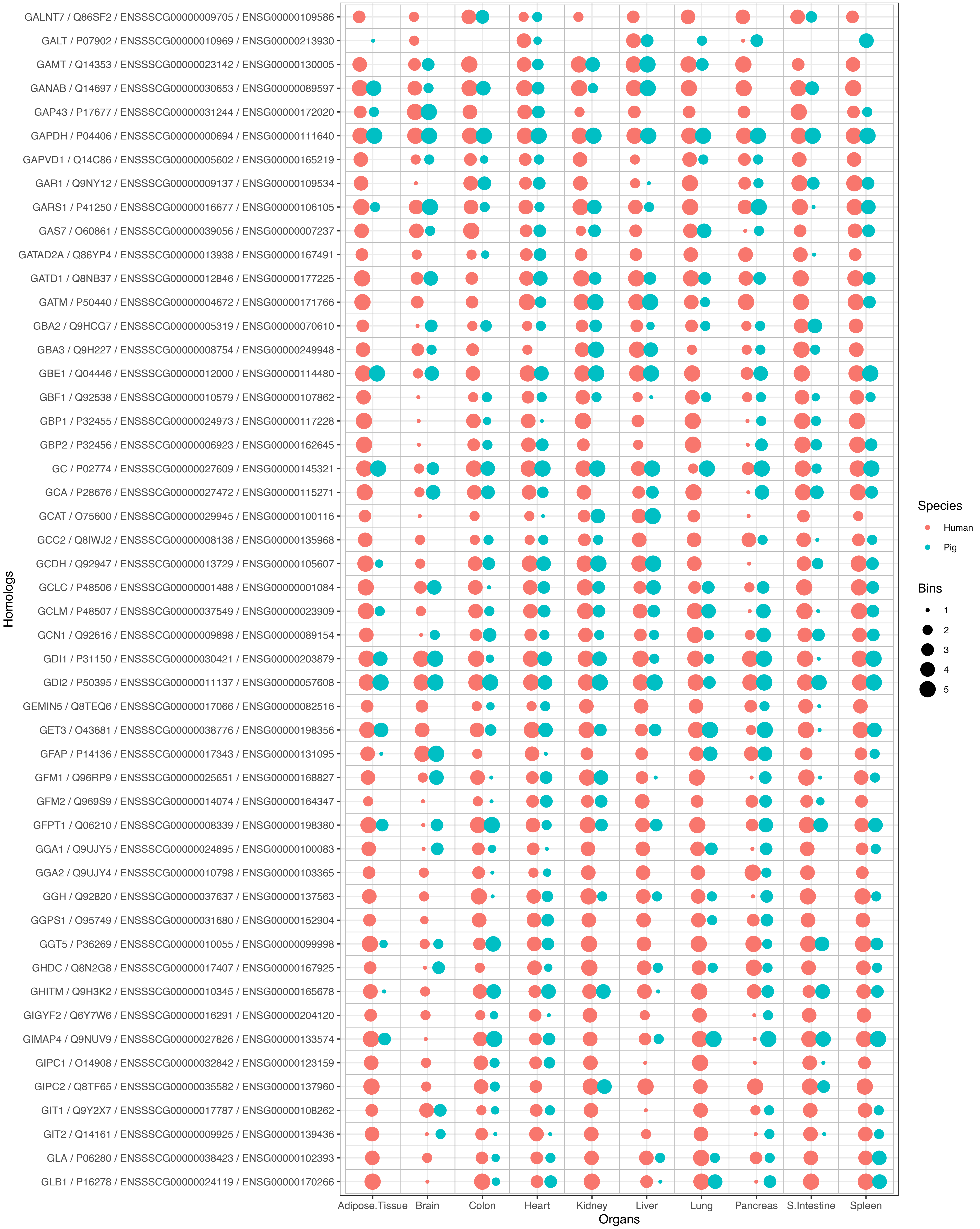
GALNT7 / Q86SF2 / ENSSSCG00000009705 / ENSG00000109586
GALT / P07902 / ENSSSCG00000010969 / ENSG00000213930
GAMT / Q14353 / ENSSSCG00000023142 / ENSG00000130005
GANAB / Q14697 / ENSSSCG00000030653 / ENSG00000089597
GAP43 / P17677 / ENSSSCG00000031244 / ENSG00000172020
GAPDH / P04406 / ENSSSCG00000000694 / ENSG00000111640
GAPVD1 / Q14C86 / ENSSSCG00000005602 / ENSG00000165219
GAR1 / Q9NY12 / ENSSSCG00000009137 / ENSG00000109534
GARS1 / P41250 / ENSSSCG00000016677 / ENSG00000106105
GAS7 / O60861 / ENSSSCG00000039056 / ENSG00000007237
GATAD2A / Q86YP4 / ENSSSCG00000013938 / ENSG00000167491
GATD1 / Q8NB37 / ENSSSCG00000012846 / ENSG00000177225
GATM / P50440 / ENSSSCG00000004672 / ENSG00000171766
GBA2 / Q9HCG7 / ENSSSCG00000005319 / ENSG00000070610
GBA3 / Q9H227 / ENSSSCG00000008754 / ENSG00000249948
GBE1 / Q04446 / ENSSSCG00000012000 / ENSG00000114480
GBF1 / Q92538 / ENSSSCG00000010579 / ENSG00000107862
GBP1 / P32455 / ENSSSCG00000024973 / ENSG00000117228
GBP2 / P32456 / ENSSSCG00000006923 / ENSG00000162645
GC / P02774 / ENSSSCG00000027609 / ENSG00000145321
GCA / P28676 / ENSSSCG00000027472 / ENSG00000115271
GCAT / O75600 / ENSSSCG00000029945 / ENSG00000100116
GCC2 / Q8IWJ2 / ENSSSCG00000008138 / ENSG00000135968
GCDH / Q92947 / ENSSSCG00000013729 / ENSG00000105607
GCLC / P48506 / ENSSSCG00000001488 / ENSG00000001084
GCLM / P48507 / ENSSSCG00000037549 / ENSG00000023909
GCN1 / Q92616 / ENSSSCG00000009898 / ENSG00000089154
GDI1 / P31150 / ENSSSCG00000030421 / ENSG00000203879
GDI2 / P50395 / ENSSSCG00000011137 / ENSG00000057608
GEMIN5 / Q8TEQ6 / ENSSSCG00000017066 / ENSG00000082516
GET3 / O43681 / ENSSSCG00000038776 / ENSG00000198356
GFAP / P14136 / ENSSSCG00000017343 / ENSG00000131095
GFM1 / Q96RP9 / ENSSSCG00000025651 / ENSG00000168827
GFM2 / Q969S9 / ENSSSCG00000014074 / ENSG00000164347
GFPT1 / Q06210 / ENSSSCG00000008339 / ENSG00000198380
GGA1 / Q9UJY5 / ENSSSCG00000024895 / ENSG00000100083
GGA2 / Q9UJY4 / ENSSSCG00000010798 / ENSG00000103365
GGH / Q92820 / ENSSSCG00000037637 / ENSG00000137563
GGPS1 / O95749 / ENSSSCG00000031680 / ENSG00000152904
GGT5 / P36269 / ENSSSCG00000010055 / ENSG00000099998
GHDC / Q8N2G8 / ENSSSCG00000017407 / ENSG00000167925
GHITM / Q9H3K2 / ENSSSCG00000010345 / ENSG00000165678
GIGYF2 / Q6Y7W6 / ENSSSCG00000016291 / ENSG00000204120
GIMAP4 / Q9NUV9 / ENSSSCG00000027826 / ENSG00000133574
GIPC1 / O14908 / ENSSSCG00000032842 / ENSG00000123159
GIPC2 / Q8TF65 / ENSSSCG00000035582 / ENSG00000137960
GIT1 / Q9Y2X7 / ENSSSCG00000017787 / ENSG00000108262
GIT2 / Q14161 / ENSSSCG00000009925 / ENSG00000139436
GLA / P06280 / ENSSSCG00000038423 / ENSG00000102393
GLB1 / P16278 / ENSSSCG00000024119 / ENSG00000170266
Homologs
Adipose.Tissue
Brain
Colon
Heart
Kidney
Liver
Lung
Pancreas
S.Intestine
Spleen
Organs
Species
Human
Pig
Bins
1
2
3
4
5

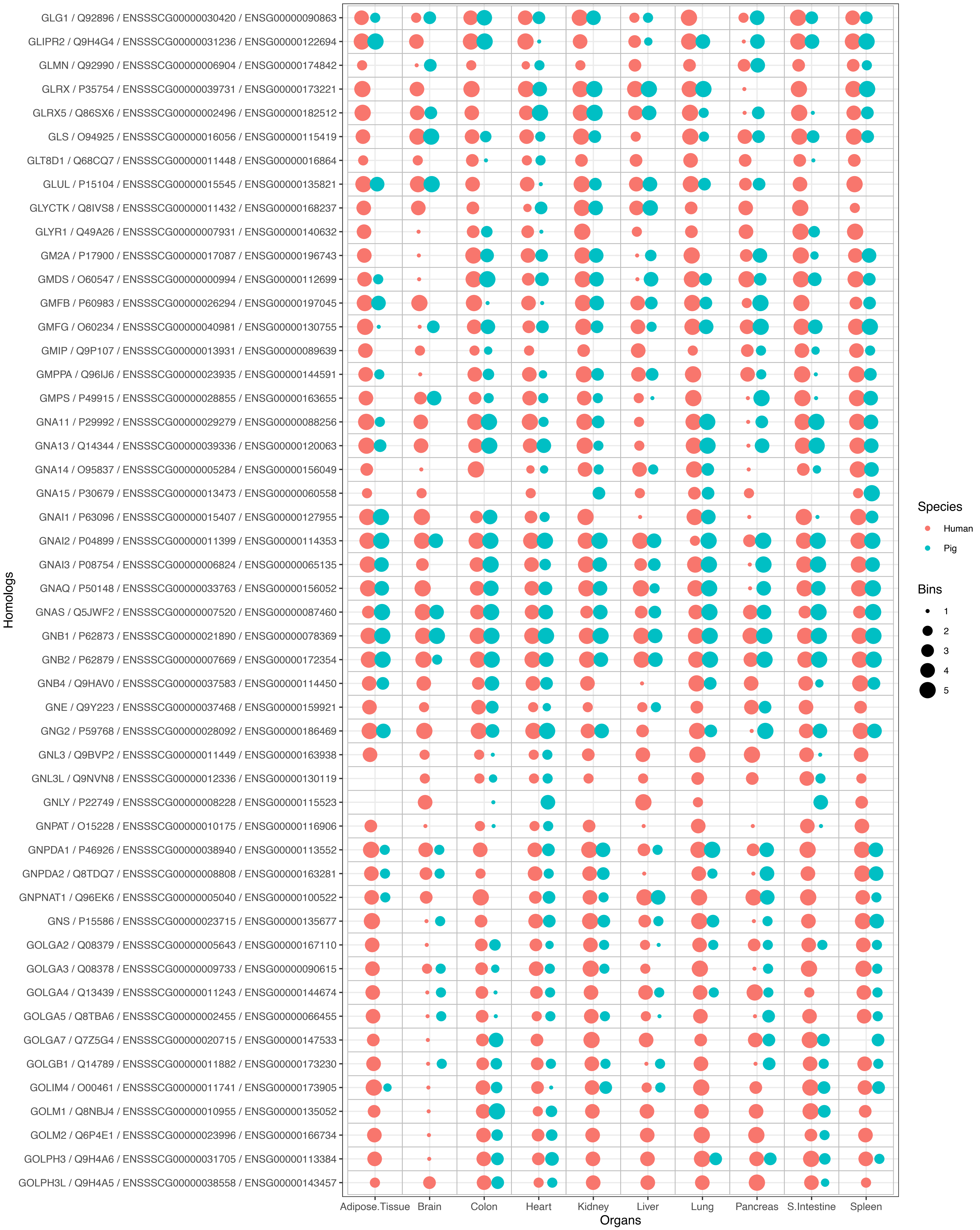
GLG1 / Q92896 / ENSSSCG00000030420 / ENSG00000090863
GLIPR2 / Q9H4G4 / ENSSSCG00000031236 / ENSG00000122694
GLMN / Q92990 / ENSSSCG00000006904 / ENSG00000174842
GLRX / P35754 / ENSSSCG00000039731 / ENSG00000173221
GLRX5 / Q86SX6 / ENSSSCG00000002496 / ENSG00000182512
GLS / O94925 / ENSSSCG00000016056 / ENSG00000115419
GLT8D1 / Q68CQ7 / ENSSSCG00000011448 / ENSG00000016864
GLUL / P15104 / ENSSSCG00000015545 / ENSG00000135821
GLYCTK / Q8IVS8 / ENSSSCG00000011432 / ENSG00000168237
GLYR1 / Q49A26 / ENSSSCG00000007931 / ENSG00000140632
GM2A / P17900 / ENSSSCG00000017087 / ENSG00000196743
GMDS / O60547 / ENSSSCG00000000994 / ENSG00000112699
GMFB / P60983 / ENSSSCG00000026294 / ENSG00000197045
GMFG / O60234 / ENSSSCG00000040981 / ENSG00000130755
GMIP / Q9P107 / ENSSSCG00000013931 / ENSG00000089639
GMPPA / Q96IJ6 / ENSSSCG00000023935 / ENSG00000144591
GMPS / P49915 / ENSSSCG00000028855 / ENSG00000163655
GNA11 / P29992 / ENSSSCG00000029279 / ENSG00000088256
GNA13 / Q14344 / ENSSSCG00000039336 / ENSG00000120063
GNA14 / O95837 / ENSSSCG00000005284 / ENSG00000156049
GNA15 / P30679 / ENSSSCG00000013473 / ENSG00000060558
GNAI1 / P63096 / ENSSSCG00000015407 / ENSG00000127955
GNAI2 / P04899 / ENSSSCG00000011399 / ENSG00000114353
GNAI3 / P08754 / ENSSSCG00000006824 / ENSG00000065135
GNAQ / P50148 / ENSSSCG00000033763 / ENSG00000156052
GNAS / Q5JWF2 / ENSSSCG00000007520 / ENSG00000087460
GNB1 / P62873 / ENSSSCG00000021890 / ENSG00000078369
GNB2 / P62879 / ENSSSCG00000007669 / ENSG00000172354
GNB4 / Q9HAV0 / ENSSSCG00000037583 / ENSG00000114450
GNE / Q9Y223 / ENSSSCG00000037468 / ENSG00000159921
GNG2 / P59768 / ENSSSCG00000028092 / ENSG00000186469
GNL3 / Q9BVP2 / ENSSSCG00000011449 / ENSG00000163938
GNL3L / Q9NVN8 / ENSSSCG00000012336 / ENSG00000130119
GNLY / P22749 / ENSSSCG00000008228 / ENSG00000115523
GNPAT / O15228 / ENSSSCG00000010175 / ENSG00000116906
GNPDA1 / P46926 / ENSSSCG00000038940 / ENSG00000113552
GNPDA2 / Q8TDQ7 / ENSSSCG00000008808 / ENSG00000163281
GNPNAT1 / Q96EK6 / ENSSSCG00000005040 / ENSG00000100522
GNS / P15586 / ENSSSCG00000023715 / ENSG00000135677
GOLGA2 / Q08379 / ENSSSCG00000005643 / ENSG00000167110
GOLGA3 / Q08378 / ENSSSCG00000009733 / ENSG00000090615
GOLGA4 / Q13439 / ENSSSCG00000011243 / ENSG00000144674
GOLGA5 / Q8TBA6 / ENSSSCG00000002455 / ENSG00000066455
GOLGA7 / Q7Z5G4 / ENSSSCG00000020715 / ENSG00000147533
GOLGB1 / Q14789 / ENSSSCG00000011882 / ENSG00000173230
GOLIM4 / O00461 / ENSSSCG00000011741 / ENSG00000173905
GOLM1 / Q8NBJ4 / ENSSSCG00000010955 / ENSG00000135052
GOLM2 / Q6P4E1 / ENSSSCG00000023996 / ENSG00000166734
GOLPH3 / Q9H4A6 / ENSSSCG00000031705 / ENSG00000113384
GOLPH3L / Q9H4A5 / ENSSSCG00000038558 / ENSG00000143457
Homologs
Adipose.Tissue
Brain
Colon
Heart
Kidney
Liver
Lung
Pancreas
S.Intestine
Spleen
Organs
Species
Human
Pig
Bins
1
2
3
4
5

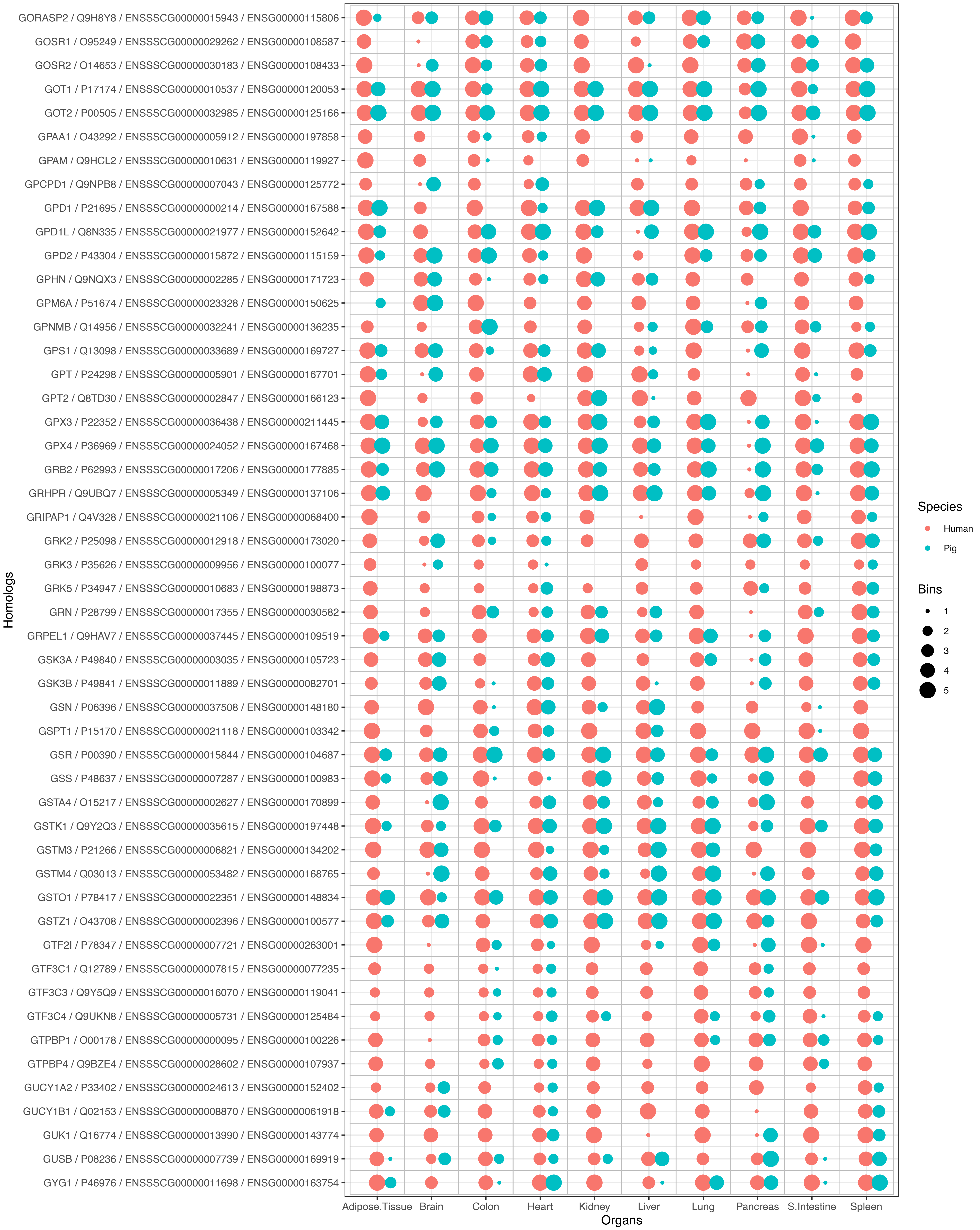
GORASP2 / Q9H8Y8 / ENSSSCG00000015943 / ENSG00000115806
GOSR1 / O95249 / ENSSSCG00000029262 / ENSG00000108587
GOSR2 / O14653 / ENSSSCG00000030183 / ENSG00000108433
GOT1 / P17174 / ENSSSCG00000010537 / ENSG00000120053
GOT2 / P00505 / ENSSSCG00000032985 / ENSG00000125166
GPAA1 / O43292 / ENSSSCG00000005912 / ENSG00000197858
GPAM / Q9HCL2 / ENSSSCG00000010631 / ENSG00000119927
GPCPD1 / Q9NPB8 / ENSSSCG00000007043 / ENSG00000125772
GPD1 / P21695 / ENSSSCG00000000214 / ENSG00000167588
GPD1L / Q8N335 / ENSSSCG00000021977 / ENSG00000152642
GPD2 / P43304 / ENSSSCG00000015872 / ENSG00000115159
GPHN / Q9NQX3 / ENSSSCG00000002285 / ENSG00000171723
GPM6A / P51674 / ENSSSCG00000023328 / ENSG00000150625
GPNMB / Q14956 / ENSSSCG00000032241 / ENSG00000136235
GPS1 / Q13098 / ENSSSCG00000033689 / ENSG00000169727
GPT / P24298 / ENSSSCG00000005901 / ENSG00000167701
GPT2 / Q8TD30 / ENSSSCG00000002847 / ENSG00000166123
GPX3 / P22352 / ENSSSCG00000036438 / ENSG00000211445
GPX4 / P36969 / ENSSSCG00000024052 / ENSG00000167468
GRB2 / P62993 / ENSSSCG00000017206 / ENSG00000177885
GRHPR / Q9UBQ7 / ENSSSCG00000005349 / ENSG00000137106
GRIPAP1 / Q4V328 / ENSSSCG00000021106 / ENSG00000068400
GRK2 / P25098 / ENSSSCG00000012918 / ENSG00000173020
GRK3 / P35626 / ENSSSCG00000009956 / ENSG00000100077
GRK5 / P34947 / ENSSSCG00000010683 / ENSG00000198873
GRN / P28799 / ENSSSCG00000017355 / ENSG00000030582
GRPEL1 / Q9HAV7 / ENSSSCG00000037445 / ENSG00000109519
GSK3A / P49840 / ENSSSCG00000003035 / ENSG00000105723
GSK3B / P49841 / ENSSSCG00000011889 / ENSG00000082701
GSN / P06396 / ENSSSCG00000037508 / ENSG00000148180
GSPT1 / P15170 / ENSSSCG00000021118 / ENSG00000103342
GSR / P00390 / ENSSSCG00000015844 / ENSG00000104687
GSS / P48637 / ENSSSCG00000007287 / ENSG00000100983
GSTA4 / O15217 / ENSSSCG00000002627 / ENSG00000170899
GSTK1 / Q9Y2Q3 / ENSSSCG00000035615 / ENSG00000197448
GSTM3 / P21266 / ENSSSCG00000006821 / ENSG00000134202
GSTM4 / Q03013 / ENSSSCG00000053482 / ENSG00000168765
GSTO1 / P78417 / ENSSSCG00000022351 / ENSG00000148834
GSTZ1 / O43708 / ENSSSCG00000002396 / ENSG00000100577
GTF2I / P78347 / ENSSSCG00000007721 / ENSG00000263001
GTF3C1 / Q12789 / ENSSSCG00000007815 / ENSG00000077235
GTF3C3 / Q9Y5Q9 / ENSSSCG00000016070 / ENSG00000119041
GTF3C4 / Q9UKN8 / ENSSSCG00000005731 / ENSG00000125484
GTPBP1 / O00178 / ENSSSCG00000000095 / ENSG00000100226
GTPBP4 / Q9BZE4 / ENSSSCG00000028602 / ENSG00000107937
GUCY1A2 / P33402 / ENSSSCG00000024613 / ENSG00000152402
GUCY1B1 / Q02153 / ENSSSCG00000008870 / ENSG00000061918
GUK1 / Q16774 / ENSSSCG00000013990 / ENSG00000143774
GUSB / P08236 / ENSSSCG00000007739 / ENSG00000169919
GYG1 / P46976 / ENSSSCG00000011698 / ENSG00000163754
Homologs
Adipose.Tissue
Brain
Colon
Heart
Kidney
Liver
Lung
Pancreas
S.Intestine
Spleen
Organs
Species
Human
Pig
Bins
1
2
3
4
5

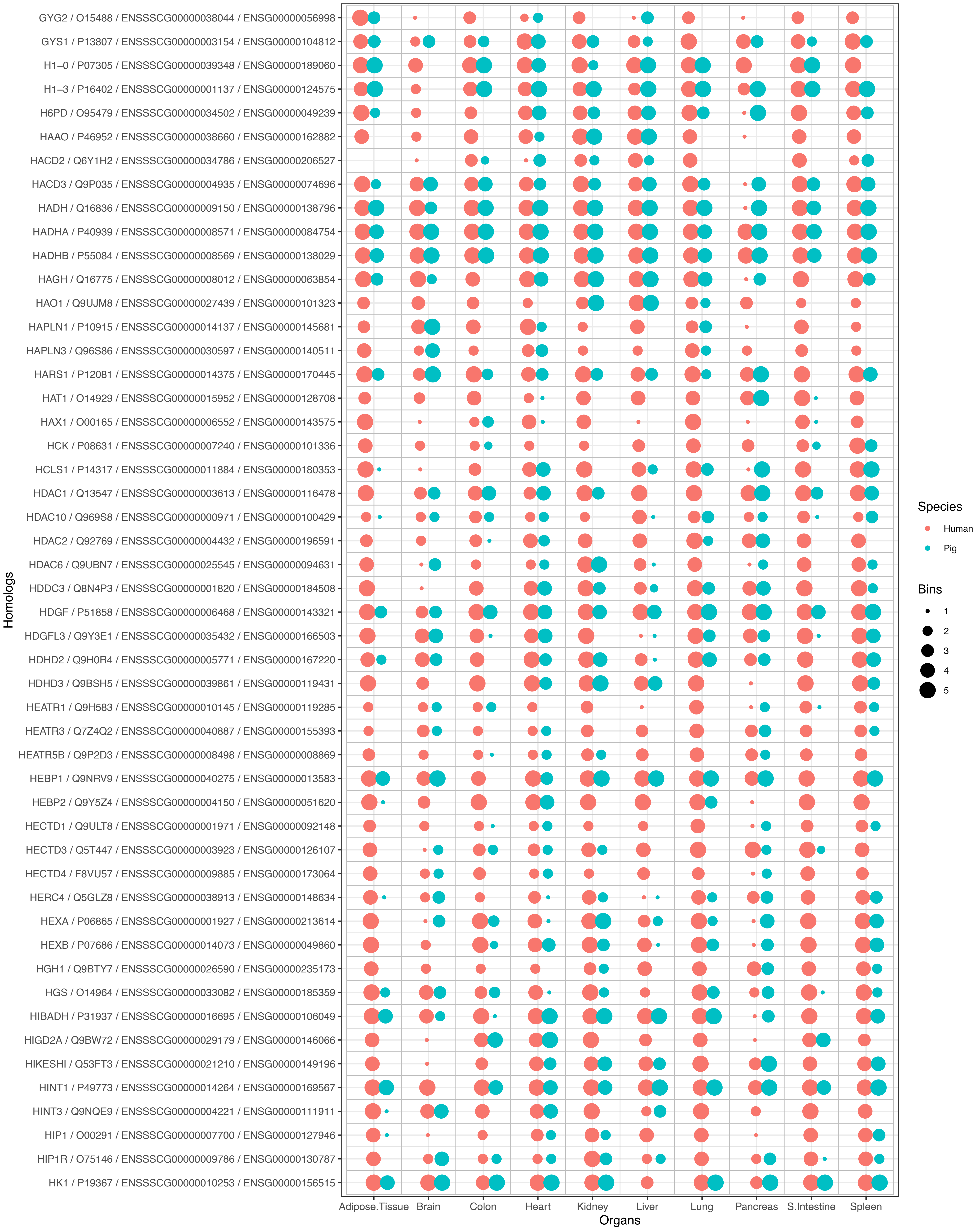
GYG2 / O15488 / ENSSSCG00000038044 / ENSG00000056998
GYS1 / P13807 / ENSSSCG00000003154 / ENSG00000104812
H1−0 / P07305 / ENSSSCG00000039348 / ENSG00000189060
H1−3 / P16402 / ENSSSCG00000001137 / ENSG00000124575
H6PD / O95479 / ENSSSCG00000034502 / ENSG00000049239
HAAO / P46952 / ENSSSCG00000038660 / ENSG00000162882
HACD2 / Q6Y1H2 / ENSSSCG00000034786 / ENSG00000206527
HACD3 / Q9P035 / ENSSSCG00000004935 / ENSG00000074696
HADH / Q16836 / ENSSSCG00000009150 / ENSG00000138796
HADHA / P40939 / ENSSSCG00000008571 / ENSG00000084754
HADHB / P55084 / ENSSSCG00000008569 / ENSG00000138029
HAGH / Q16775 / ENSSSCG00000008012 / ENSG00000063854
HAO1 / Q9UJM8 / ENSSSCG00000027439 / ENSG00000101323
HAPLN1 / P10915 / ENSSSCG00000014137 / ENSG00000145681
HAPLN3 / Q96S86 / ENSSSCG00000030597 / ENSG00000140511
HARS1 / P12081 / ENSSSCG00000014375 / ENSG00000170445
HAT1 / O14929 / ENSSSCG00000015952 / ENSG00000128708
HAX1 / O00165 / ENSSSCG00000006552 / ENSG00000143575
HCK / P08631 / ENSSSCG00000007240 / ENSG00000101336
HCLS1 / P14317 / ENSSSCG00000011884 / ENSG00000180353
HDAC1 / Q13547 / ENSSSCG00000003613 / ENSG00000116478
HDAC10 / Q969S8 / ENSSSCG00000000971 / ENSG00000100429
HDAC2 / Q92769 / ENSSSCG00000004432 / ENSG00000196591
HDAC6 / Q9UBN7 / ENSSSCG00000025545 / ENSG00000094631
HDDC3 / Q8N4P3 / ENSSSCG00000001820 / ENSG00000184508
HDGF / P51858 / ENSSSCG00000006468 / ENSG00000143321
HDGFL3 / Q9Y3E1 / ENSSSCG00000035432 / ENSG00000166503
HDHD2 / Q9H0R4 / ENSSSCG00000005771 / ENSG00000167220
HDHD3 / Q9BSH5 / ENSSSCG00000039861 / ENSG00000119431
HEATR1 / Q9H583 / ENSSSCG00000010145 / ENSG00000119285
HEATR3 / Q7Z4Q2 / ENSSSCG00000040887 / ENSG00000155393
HEATR5B / Q9P2D3 / ENSSSCG00000008498 / ENSG00000008869
HEBP1 / Q9NRV9 / ENSSSCG00000040275 / ENSG00000013583
HEBP2 / Q9Y5Z4 / ENSSSCG00000004150 / ENSG00000051620
HECTD1 / Q9ULT8 / ENSSSCG00000001971 / ENSG00000092148
HECTD3 / Q5T447 / ENSSSCG00000003923 / ENSG00000126107
HECTD4 / F8VU57 / ENSSSCG00000009885 / ENSG00000173064
HERC4 / Q5GLZ8 / ENSSSCG00000038913 / ENSG00000148634
HEXA / P06865 / ENSSSCG00000001927 / ENSG00000213614
HEXB / P07686 / ENSSSCG00000014073 / ENSG00000049860
HGH1 / Q9BTY7 / ENSSSCG00000026590 / ENSG00000235173
HGS / O14964 / ENSSSCG00000033082 / ENSG00000185359
HIBADH / P31937 / ENSSSCG00000016695 / ENSG00000106049
HIGD2A / Q9BW72 / ENSSSCG00000029179 / ENSG00000146066
HIKESHI / Q53FT3 / ENSSSCG00000021210 / ENSG00000149196
HINT1 / P49773 / ENSSSCG00000014264 / ENSG00000169567
HINT3 / Q9NQE9 / ENSSSCG00000004221 / ENSG00000111911
HIP1 / O00291 / ENSSSCG00000007700 / ENSG00000127946
HIP1R / O75146 / ENSSSCG00000009786 / ENSG00000130787
HK1 / P19367 / ENSSSCG00000010253 / ENSG00000156515
Homologs
Adipose.Tissue
Brain
Colon
Heart
Kidney
Liver
Lung
Pancreas
S.Intestine
Spleen
Organs
Species
Human
Pig
Bins
1
2
3
4
5

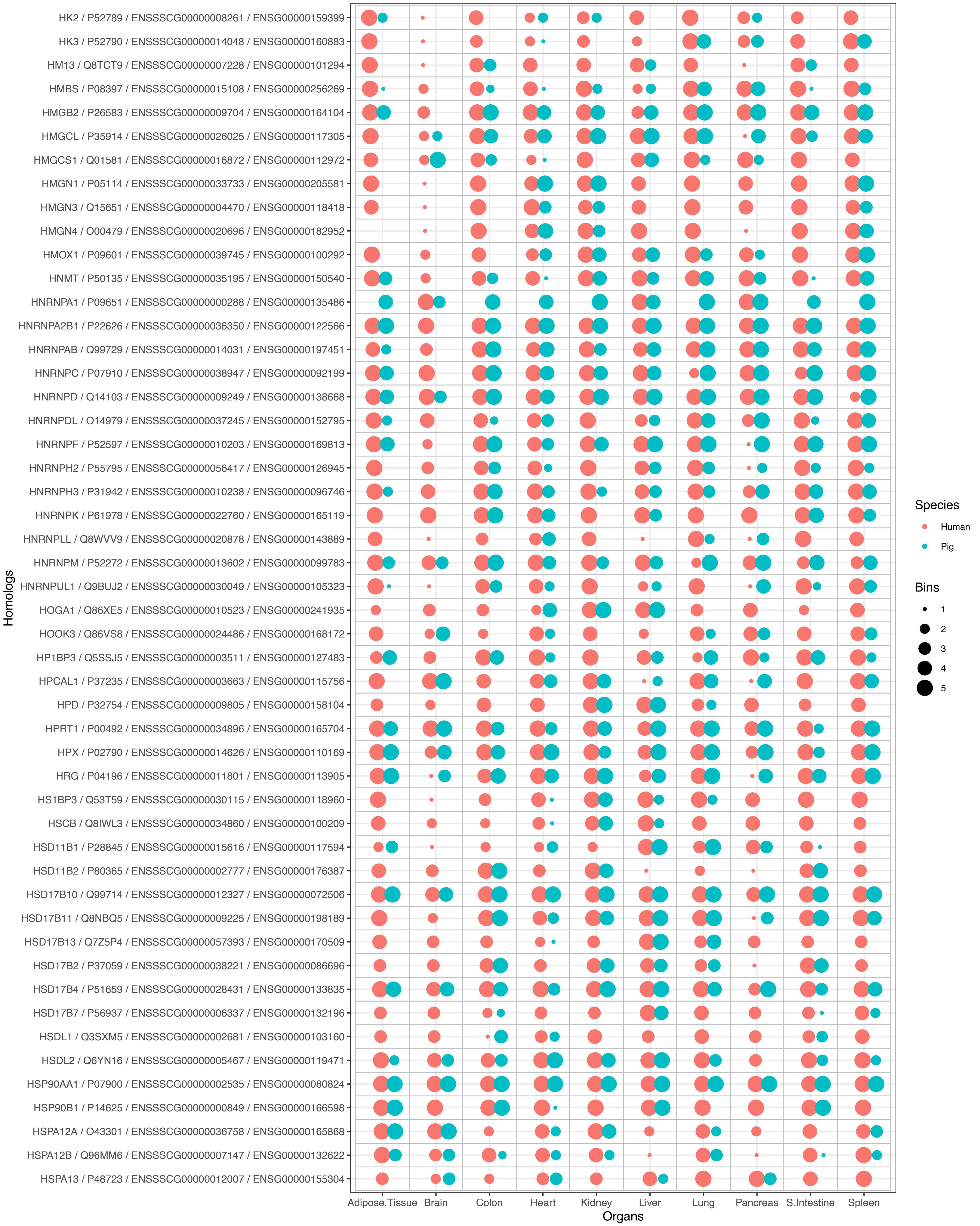

HK2 / P52789 / ENSSSCG00000008261 / ENSG00000159399
HK3 / P52790 / ENSSSCG00000014048 / ENSG00000160883
HM13 / Q8TCT9 / ENSSSCG00000007228 / ENSG00000101294
HMBS / P08397 / ENSSSCG00000015108 / ENSG00000256269
HMGB2 / P26583 / ENSSSCG00000009704 / ENSG00000164104
HMGCL / P35914 / ENSSSCG00000026025 / ENSG00000117305
HMGCS1 / Q01581 / ENSSSCG00000016872 / ENSG00000112972
HMGN1 / P05114 / ENSSSCG00000033733 / ENSG00000205581
HMGN3 / Q15651 / ENSSSCG00000004470 / ENSG00000118418
HMGN4 / O00479 / ENSSSCG00000020696 / ENSG00000182952
HMOX1 / P09601 / ENSSSCG00000039745 / ENSG00000100292
HNMT / P50135 / ENSSSCG00000035195 / ENSG00000150540
HNRNPA1 / P09651 / ENSSSCG00000000288 / ENSG00000135486
HNRNPA2B1 / P22626 / ENSSSCG00000036350 / ENSG00000122566
HNRNPAB / Q99729 / ENSSSCG00000014031 / ENSG00000197451
HNRNPC / P07910 / ENSSSCG00000038947 / ENSG00000092199
HNRNPD / Q14103 / ENSSSCG00000009249 / ENSG00000138668
HNRNPDL / O14979 / ENSSSCG00000037245 / ENSG00000152795
HNRNPF / P52597 / ENSSSCG00000010203 / ENSG00000169813
HNRNPH2 / P55795 / ENSSSCG00000056417 / ENSG00000126945
HNRNPH3 / P31942 / ENSSSCG00000010238 / ENSG00000096746
HNRNPK / P61978 / ENSSSCG00000022760 / ENSG00000165119
HNRNPLL / Q8WVV9 / ENSSSCG00000020878 / ENSG00000143889
HNRNPM / P52272 / ENSSSCG00000013602 / ENSG00000099783
HNRNPUL1 / Q9BUJ2 / ENSSSCG00000030049 / ENSG00000105323
HOGA1 / Q86XE5 / ENSSSCG00000010523 / ENSG00000241935
HOOK3 / Q86VS8 / ENSSSCG00000024486 / ENSG00000168172
HP1BP3 / Q5SSJ5 / ENSSSCG00000003511 / ENSG00000127483
HPCAL1 / P37235 / ENSSSCG00000003663 / ENSG00000115756
HPD / P32754 / ENSSSCG00000009805 / ENSG00000158104
HPRT1 / P00492 / ENSSSCG00000034896 / ENSG00000165704
HPX / P02790 / ENSSSCG00000014626 / ENSG00000110169
HRG / P04196 / ENSSSCG00000011801 / ENSG00000113905
HS1BP3 / Q53T59 / ENSSSCG00000030115 / ENSG00000118960
HSCB / Q8IWL3 / ENSSSCG00000034860 / ENSG00000100209
HSD11B1 / P28845 / ENSSSCG00000015616 / ENSG00000117594
HSD11B2 / P80365 / ENSSSCG00000002777 / ENSG00000176387
HSD17B10 / Q99714 / ENSSSCG00000012327 / ENSG00000072506
HSD17B11 / Q8NBQ5 / ENSSSCG00000009225 / ENSG00000198189
HSD17B13 / Q7Z5P4 / ENSSSCG00000057393 / ENSG00000170509
HSD17B2 / P37059 / ENSSSCG00000038221 / ENSG00000086696
HSD17B4 / P51659 / ENSSSCG00000028431 / ENSG00000133835
HSD17B7 / P56937 / ENSSSCG00000006337 / ENSG00000132196
HSDL1 / Q3SXM5 / ENSSSCG00000002681 / ENSG00000103160
HSDL2 / Q6YN16 / ENSSSCG00000005467 / ENSG00000119471
HSP90AA1 / P07900 / ENSSSCG00000002535 / ENSG00000080824
HSP90B1 / P14625 / ENSSSCG00000000849 / ENSG00000166598
HSPA12A / O43301 / ENSSSCG00000036758 / ENSG00000165868
HSPA12B / Q96MM6 / ENSSSCG00000007147 / ENSG00000132622
HSPA13 / P48723 / ENSSSCG00000012007 / ENSG00000155304
Homologs
Adipose.Tissue
Brain
Colon
Heart
Kidney
Liver
Lung
Pancreas
S.Intestine
Spleen
Organs
Species
Human
Pig
Bins
1
2
3
4
5

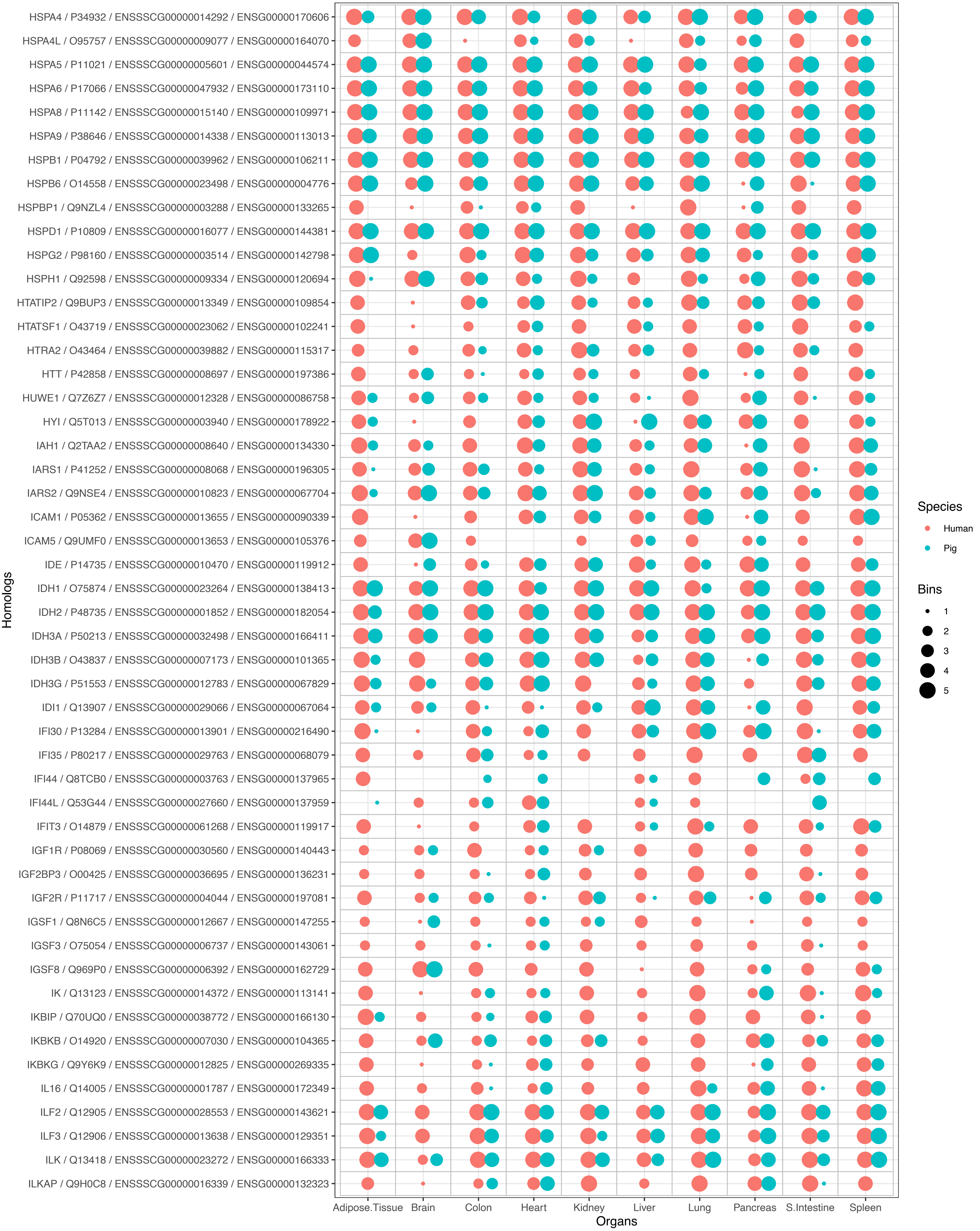
HSPA4 / P34932 / ENSSSCG00000014292 / ENSG00000170606
HSPA4L / O95757 / ENSSSCG00000009077 / ENSG00000164070
HSPA5 / P11021 / ENSSSCG00000005601 / ENSG00000044574
HSPA6 / P17066 / ENSSSCG00000047932 / ENSG00000173110
HSPA8 / P11142 / ENSSSCG00000015140 / ENSG00000109971
HSPA9 / P38646 / ENSSSCG00000014338 / ENSG00000113013
HSPB1 / P04792 / ENSSSCG00000039962 / ENSG00000106211
HSPB6 / O14558 / ENSSSCG00000023498 / ENSG00000004776
HSPBP1 / Q9NZL4 / ENSSSCG00000003288 / ENSG00000133265
HSPD1 / P10809 / ENSSSCG00000016077 / ENSG00000144381
HSPG2 / P98160 / ENSSSCG00000003514 / ENSG00000142798
HSPH1 / Q92598 / ENSSSCG00000009334 / ENSG00000120694
HTATIP2 / Q9BUP3 / ENSSSCG00000013349 / ENSG00000109854
HTATSF1 / O43719 / ENSSSCG00000023062 / ENSG00000102241
HTRA2 / O43464 / ENSSSCG00000039882 / ENSG00000115317
HTT / P42858 / ENSSSCG00000008697 / ENSG00000197386
HUWE1 / Q7Z6Z7 / ENSSSCG00000012328 / ENSG00000086758
HYI / Q5T013 / ENSSSCG00000003940 / ENSG00000178922
IAH1 / Q2TAA2 / ENSSSCG00000008640 / ENSG00000134330
IARS1 / P41252 / ENSSSCG00000008068 / ENSG00000196305
IARS2 / Q9NSE4 / ENSSSCG00000010823 / ENSG00000067704
ICAM1 / P05362 / ENSSSCG00000013655 / ENSG00000090339
ICAM5 / Q9UMF0 / ENSSSCG00000013653 / ENSG00000105376
IDE / P14735 / ENSSSCG00000010470 / ENSG00000119912
IDH1 / O75874 / ENSSSCG00000023264 / ENSG00000138413
IDH2 / P48735 / ENSSSCG00000001852 / ENSG00000182054
IDH3A / P50213 / ENSSSCG00000032498 / ENSG00000166411
IDH3B / O43837 / ENSSSCG00000007173 / ENSG00000101365
IDH3G / P51553 / ENSSSCG00000012783 / ENSG00000067829
IDI1 / Q13907 / ENSSSCG00000029066 / ENSG00000067064
IFI30 / P13284 / ENSSSCG00000013901 / ENSG00000216490
IFI35 / P80217 / ENSSSCG00000029763 / ENSG00000068079
IFI44 / Q8TCB0 / ENSSSCG00000003763 / ENSG00000137965
IFI44L / Q53G44 / ENSSSCG00000027660 / ENSG00000137959
IFIT3 / O14879 / ENSSSCG00000061268 / ENSG00000119917
IGF1R / P08069 / ENSSSCG00000030560 / ENSG00000140443
IGF2BP3 / O00425 / ENSSSCG00000036695 / ENSG00000136231
IGF2R / P11717 / ENSSSCG00000004044 / ENSG00000197081
IGSF1 / Q8N6C5 / ENSSSCG00000012667 / ENSG00000147255
IGSF3 / O75054 / ENSSSCG00000006737 / ENSG00000143061
IGSF8 / Q969P0 / ENSSSCG00000006392 / ENSG00000162729
IK / Q13123 / ENSSSCG00000014372 / ENSG00000113141
IKBIP / Q70UQ0 / ENSSSCG00000038772 / ENSG00000166130
IKBKB / O14920 / ENSSSCG00000007030 / ENSG00000104365
IKBKG / Q9Y6K9 / ENSSSCG00000012825 / ENSG00000269335
IL16 / Q14005 / ENSSSCG00000001787 / ENSG00000172349
ILF2 / Q12905 / ENSSSCG00000028553 / ENSG00000143621
ILF3 / Q12906 / ENSSSCG00000013638 / ENSG00000129351
ILK / Q13418 / ENSSSCG00000023272 / ENSG00000166333
ILKAP / Q9H0C8 / ENSSSCG00000016339 / ENSG00000132323
Homologs
Adipose.Tissue
Brain
Colon
Heart
Kidney
Liver
Lung
Pancreas
S.Intestine
Spleen
Organs
Species
Human
Pig
Bins
1
2
3
4
5

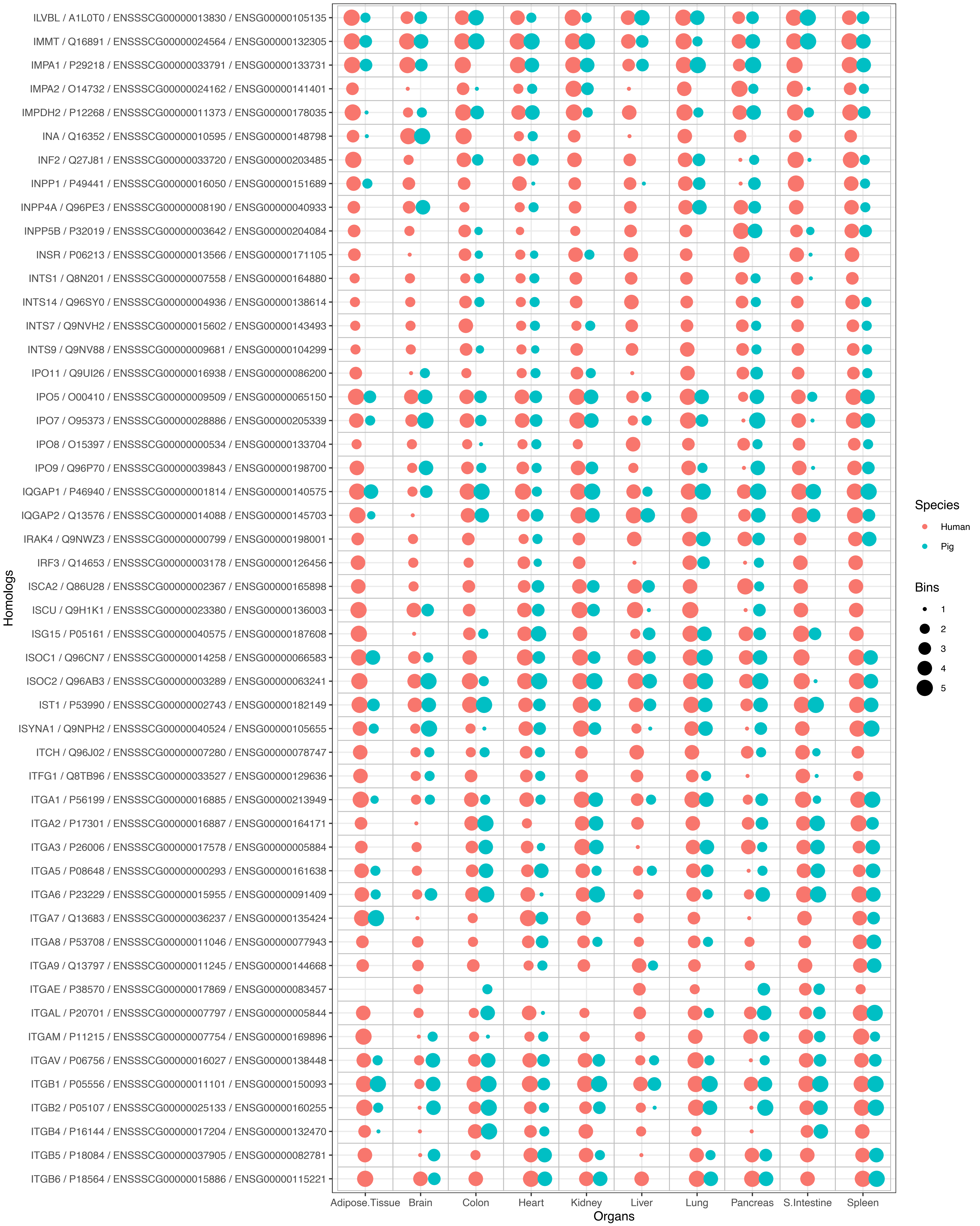
ILVBL / A1L0T0 / ENSSSCG00000013830 / ENSG00000105135
IMMT / Q16891 / ENSSSCG00000024564 / ENSG00000132305
IMPA1 / P29218 / ENSSSCG00000033791 / ENSG00000133731
IMPA2 / O14732 / ENSSSCG00000024162 / ENSG00000141401
IMPDH2 / P12268 / ENSSSCG00000011373 / ENSG00000178035
INA / Q16352 / ENSSSCG00000010595 / ENSG00000148798
INF2 / Q27J81 / ENSSSCG00000033720 / ENSG00000203485
INPP1 / P49441 / ENSSSCG00000016050 / ENSG00000151689
INPP4A / Q96PE3 / ENSSSCG00000008190 / ENSG00000040933
INPP5B / P32019 / ENSSSCG00000003642 / ENSG00000204084
INSR / P06213 / ENSSSCG00000013566 / ENSG00000171105
INTS1 / Q8N201 / ENSSSCG00000007558 / ENSG00000164880
INTS14 / Q96SY0 / ENSSSCG00000004936 / ENSG00000138614
INTS7 / Q9NVH2 / ENSSSCG00000015602 / ENSG00000143493
INTS9 / Q9NV88 / ENSSSCG00000009681 / ENSG00000104299
IPO11 / Q9UI26 / ENSSSCG00000016938 / ENSG00000086200
IPO5 / O00410 / ENSSSCG00000009509 / ENSG00000065150
IPO7 / O95373 / ENSSSCG00000028886 / ENSG00000205339
IPO8 / O15397 / ENSSSCG00000000534 / ENSG00000133704
IPO9 / Q96P70 / ENSSSCG00000039843 / ENSG00000198700
IQGAP1 / P46940 / ENSSSCG00000001814 / ENSG00000140575
IQGAP2 / Q13576 / ENSSSCG00000014088 / ENSG00000145703
IRAK4 / Q9NWZ3 / ENSSSCG00000000799 / ENSG00000198001
IRF3 / Q14653 / ENSSSCG00000003178 / ENSG00000126456
ISCA2 / Q86U28 / ENSSSCG00000002367 / ENSG00000165898
ISCU / Q9H1K1 / ENSSSCG00000023380 / ENSG00000136003
ISG15 / P05161 / ENSSSCG00000040575 / ENSG00000187608
ISOC1 / Q96CN7 / ENSSSCG00000014258 / ENSG00000066583
ISOC2 / Q96AB3 / ENSSSCG00000003289 / ENSG00000063241
IST1 / P53990 / ENSSSCG00000002743 / ENSG00000182149
ISYNA1 / Q9NPH2 / ENSSSCG00000040524 / ENSG00000105655
ITCH / Q96J02 / ENSSSCG00000007280 / ENSG00000078747
ITFG1 / Q8TB96 / ENSSSCG00000033527 / ENSG00000129636
ITGA1 / P56199 / ENSSSCG00000016885 / ENSG00000213949
ITGA2 / P17301 / ENSSSCG00000016887 / ENSG00000164171
ITGA3 / P26006 / ENSSSCG00000017578 / ENSG00000005884
ITGA5 / P08648 / ENSSSCG00000000293 / ENSG00000161638
ITGA6 / P23229 / ENSSSCG00000015955 / ENSG00000091409
ITGA7 / Q13683 / ENSSSCG00000036237 / ENSG00000135424
ITGA8 / P53708 / ENSSSCG00000011046 / ENSG00000077943
ITGA9 / Q13797 / ENSSSCG00000011245 / ENSG00000144668
ITGAE / P38570 / ENSSSCG00000017869 / ENSG00000083457
ITGAL / P20701 / ENSSSCG00000007797 / ENSG00000005844
ITGAM / P11215 / ENSSSCG00000007754 / ENSG00000169896
ITGAV / P06756 / ENSSSCG00000016027 / ENSG00000138448
ITGB1 / P05556 / ENSSSCG00000011101 / ENSG00000150093
ITGB2 / P05107 / ENSSSCG00000025133 / ENSG00000160255
ITGB4 / P16144 / ENSSSCG00000017204 / ENSG00000132470
ITGB5 / P18084 / ENSSSCG00000037905 / ENSG00000082781
ITGB6 / P18564 / ENSSSCG00000015886 / ENSG00000115221
Homologs
Adipose.Tissue
Brain
Colon
Heart
Kidney
Liver
Lung
Pancreas
S.Intestine
Spleen
Organs
Species
Human
Pig
Bins
1
2
3
4
5

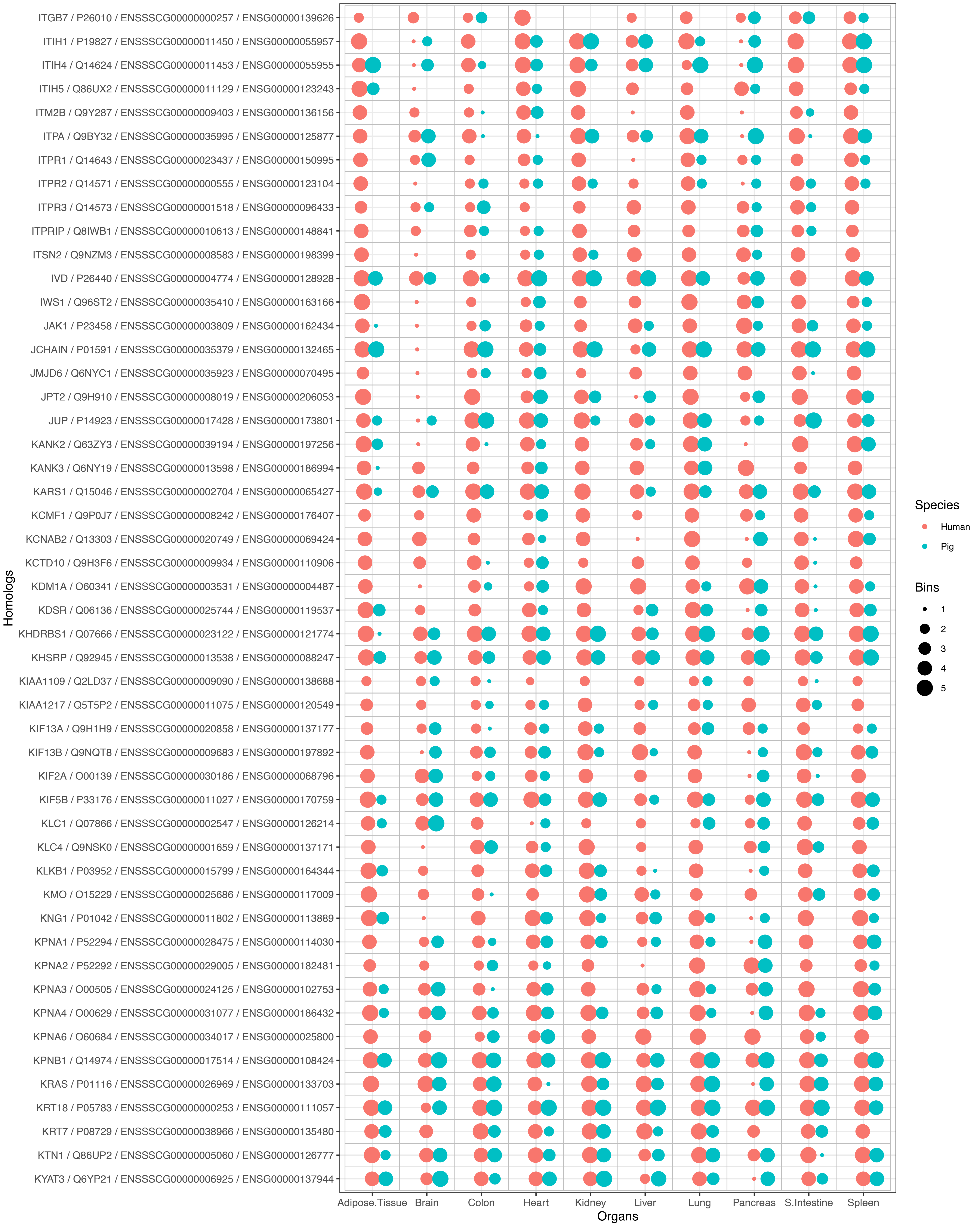
ITGB7 / P26010 / ENSSSCG00000000257 / ENSG00000139626
ITIH1 / P19827 / ENSSSCG00000011450 / ENSG00000055957
ITIH4 / Q14624 / ENSSSCG00000011453 / ENSG00000055955
ITIH5 / Q86UX2 / ENSSSCG00000011129 / ENSG00000123243
ITM2B / Q9Y287 / ENSSSCG00000009403 / ENSG00000136156
ITPA / Q9BY32 / ENSSSCG00000035995 / ENSG00000125877
ITPR1 / Q14643 / ENSSSCG00000023437 / ENSG00000150995
ITPR2 / Q14571 / ENSSSCG00000000555 / ENSG00000123104
ITPR3 / Q14573 / ENSSSCG00000001518 / ENSG00000096433
ITPRIP / Q8IWB1 / ENSSSCG00000010613 / ENSG00000148841
ITSN2 / Q9NZM3 / ENSSSCG00000008583 / ENSG00000198399
IVD / P26440 / ENSSSCG00000004774 / ENSG00000128928
IWS1 / Q96ST2 / ENSSSCG00000035410 / ENSG00000163166
JAK1 / P23458 / ENSSSCG00000003809 / ENSG00000162434
JCHAIN / P01591 / ENSSSCG00000035379 / ENSG00000132465
JMJD6 / Q6NYC1 / ENSSSCG00000035923 / ENSG00000070495
JPT2 / Q9H910 / ENSSSCG00000008019 / ENSG00000206053
JUP / P14923 / ENSSSCG00000017428 / ENSG00000173801
KANK2 / Q63ZY3 / ENSSSCG00000039194 / ENSG00000197256
KANK3 / Q6NY19 / ENSSSCG00000013598 / ENSG00000186994
KARS1 / Q15046 / ENSSSCG00000002704 / ENSG00000065427
KCMF1 / Q9P0J7 / ENSSSCG00000008242 / ENSG00000176407
KCNAB2 / Q13303 / ENSSSCG00000020749 / ENSG00000069424
KCTD10 / Q9H3F6 / ENSSSCG00000009934 / ENSG00000110906
KDM1A / O60341 / ENSSSCG00000003531 / ENSG00000004487
KDSR / Q06136 / ENSSSCG00000025744 / ENSG00000119537
KHDRBS1 / Q07666 / ENSSSCG00000023122 / ENSG00000121774
KHSRP / Q92945 / ENSSSCG00000013538 / ENSG00000088247
KIAA1109 / Q2LD37 / ENSSSCG00000009090 / ENSG00000138688
KIAA1217 / Q5T5P2 / ENSSSCG00000011075 / ENSG00000120549
KIF13A / Q9H1H9 / ENSSSCG00000020858 / ENSG00000137177
KIF13B / Q9NQT8 / ENSSSCG00000009683 / ENSG00000197892
KIF2A / O00139 / ENSSSCG00000030186 / ENSG00000068796
KIF5B / P33176 / ENSSSCG00000011027 / ENSG00000170759
KLC1 / Q07866 / ENSSSCG00000002547 / ENSG00000126214
KLC4 / Q9NSK0 / ENSSSCG00000001659 / ENSG00000137171
KLKB1 / P03952 / ENSSSCG00000015799 / ENSG00000164344
KMO / O15229 / ENSSSCG00000025686 / ENSG00000117009
KNG1 / P01042 / ENSSSCG00000011802 / ENSG00000113889
KPNA1 / P52294 / ENSSSCG00000028475 / ENSG00000114030
KPNA2 / P52292 / ENSSSCG00000029005 / ENSG00000182481
KPNA3 / O00505 / ENSSSCG00000024125 / ENSG00000102753
KPNA4 / O00629 / ENSSSCG00000031077 / ENSG00000186432
KPNA6 / O60684 / ENSSSCG00000034017 / ENSG00000025800
KPNB1 / Q14974 / ENSSSCG00000017514 / ENSG00000108424
KRAS / P01116 / ENSSSCG00000026969 / ENSG00000133703
KRT18 / P05783 / ENSSSCG00000000253 / ENSG00000111057
KRT7 / P08729 / ENSSSCG00000038966 / ENSG00000135480
KTN1 / Q86UP2 / ENSSSCG00000005060 / ENSG00000126777
KYAT3 / Q6YP21 / ENSSSCG00000006925 / ENSG00000137944
Homologs
Adipose.Tissue
Brain
Colon
Heart
Kidney
Liver
Lung
Pancreas
S.Intestine
Spleen
Organs
Species
Human
Pig
Bins
1
2
3
4
5

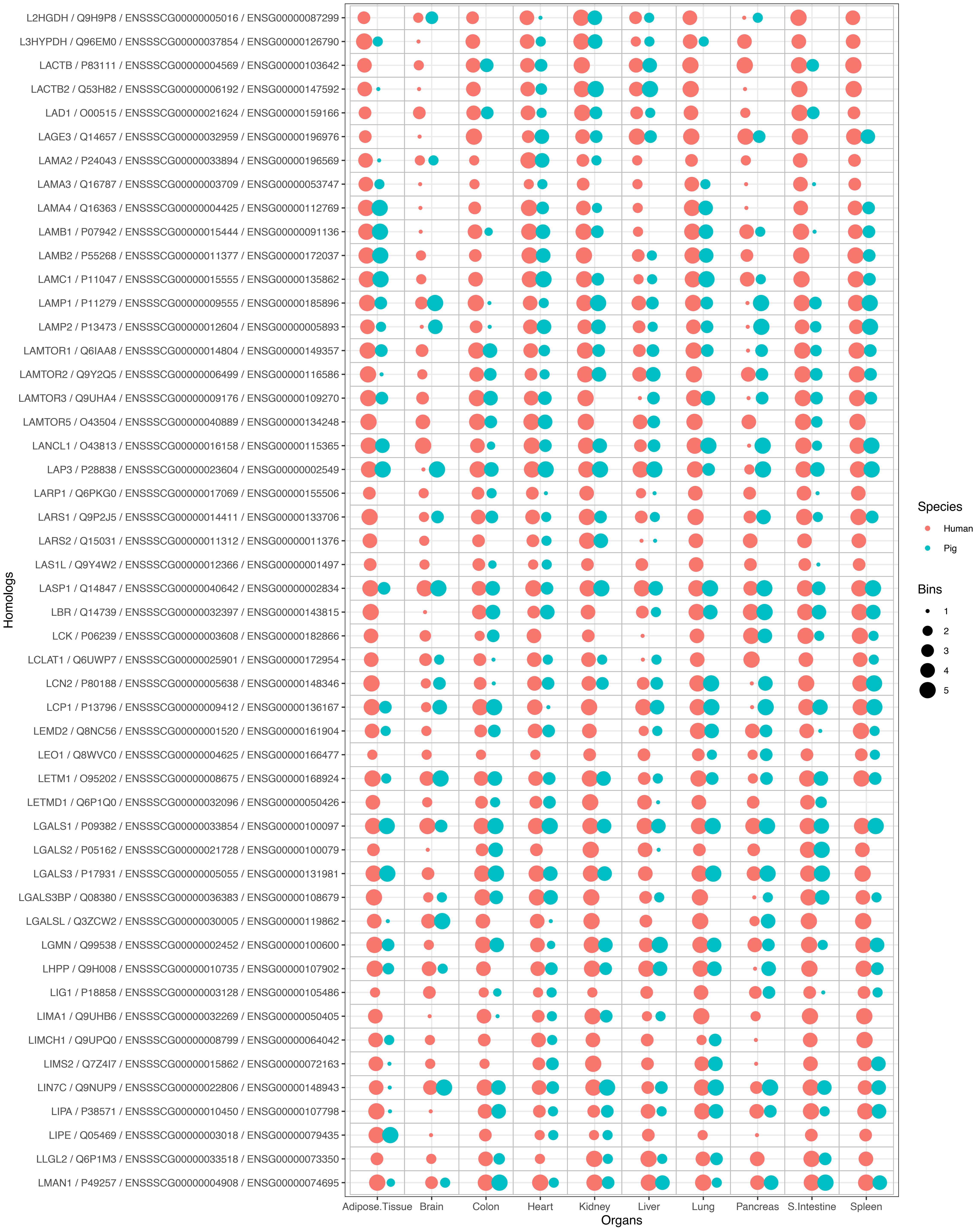

L2HGDH / Q9H9P8 / ENSSSCG00000005016 / ENSG00000087299
L3HYPDH / Q96EM0 / ENSSSCG00000037854 / ENSG00000126790
LACTB / P83111 / ENSSSCG00000004569 / ENSG00000103642
LACTB2 / Q53H82 / ENSSSCG00000006192 / ENSG00000147592
LAD1 / O00515 / ENSSSCG00000021624 / ENSG00000159166
LAGE3 / Q14657 / ENSSSCG00000032959 / ENSG00000196976
LAMA2 / P24043 / ENSSSCG00000033894 / ENSG00000196569
LAMA3 / Q16787 / ENSSSCG00000003709 / ENSG00000053747
LAMA4 / Q16363 / ENSSSCG00000004425 / ENSG00000112769
LAMB1 / P07942 / ENSSSCG00000015444 / ENSG00000091136
LAMB2 / P55268 / ENSSSCG00000011377 / ENSG00000172037
LAMC1 / P11047 / ENSSSCG00000015555 / ENSG00000135862
LAMP1 / P11279 / ENSSSCG00000009555 / ENSG00000185896
LAMP2 / P13473 / ENSSSCG00000012604 / ENSG00000005893
LAMTOR1 / Q6IAA8 / ENSSSCG00000014804 / ENSG00000149357
LAMTOR2 / Q9Y2Q5 / ENSSSCG00000006499 / ENSG00000116586
LAMTOR3 / Q9UHA4 / ENSSSCG00000009176 / ENSG00000109270
LAMTOR5 / O43504 / ENSSSCG00000040889 / ENSG00000134248
LANCL1 / O43813 / ENSSSCG00000016158 / ENSG00000115365
LAP3 / P28838 / ENSSSCG00000023604 / ENSG00000002549
LARP1 / Q6PKG0 / ENSSSCG00000017069 / ENSG00000155506
LARS1 / Q9P2J5 / ENSSSCG00000014411 / ENSG00000133706
LARS2 / Q15031 / ENSSSCG00000011312 / ENSG00000011376
LAS1L / Q9Y4W2 / ENSSSCG00000012366 / ENSG00000001497
LASP1 / Q14847 / ENSSSCG00000040642 / ENSG00000002834
LBR / Q14739 / ENSSSCG00000032397 / ENSG00000143815
LCK / P06239 / ENSSSCG00000003608 / ENSG00000182866
LCLAT1 / Q6UWP7 / ENSSSCG00000025901 / ENSG00000172954
LCN2 / P80188 / ENSSSCG00000005638 / ENSG00000148346
LCP1 / P13796 / ENSSSCG00000009412 / ENSG00000136167
LEMD2 / Q8NC56 / ENSSSCG00000001520 / ENSG00000161904
LEO1 / Q8WVC0 / ENSSSCG00000004625 / ENSG00000166477
LETM1 / O95202 / ENSSSCG00000008675 / ENSG00000168924
LETMD1 / Q6P1Q0 / ENSSSCG00000032096 / ENSG00000050426
LGALS1 / P09382 / ENSSSCG00000033854 / ENSG00000100097
LGALS2 / P05162 / ENSSSCG00000021728 / ENSG00000100079
LGALS3 / P17931 / ENSSSCG00000005055 / ENSG00000131981
LGALS3BP / Q08380 / ENSSSCG00000036383 / ENSG00000108679
LGALSL / Q3ZCW2 / ENSSSCG00000030005 / ENSG00000119862
LGMN / Q99538 / ENSSSCG00000002452 / ENSG00000100600
LHPP / Q9H008 / ENSSSCG00000010735 / ENSG00000107902
LIG1 / P18858 / ENSSSCG00000003128 / ENSG00000105486
LIMA1 / Q9UHB6 / ENSSSCG00000032269 / ENSG00000050405
LIMCH1 / Q9UPQ0 / ENSSSCG00000008799 / ENSG00000064042
LIMS2 / Q7Z4I7 / ENSSSCG00000015862 / ENSG00000072163
LIN7C / Q9NUP9 / ENSSSCG00000022806 / ENSG00000148943
LIPA / P38571 / ENSSSCG00000010450 / ENSG00000107798
LIPE / Q05469 / ENSSSCG00000003018 / ENSG00000079435
LLGL2 / Q6P1M3 / ENSSSCG00000033518 / ENSG00000073350
LMAN1 / P49257 / ENSSSCG00000004908 / ENSG00000074695
Homologs
Adipose.Tissue
Brain
Colon
Heart
Kidney
Liver
Lung
Pancreas
S.Intestine
Spleen
Organs
Species
Human
Pig
Bins
1
2
3
4
5

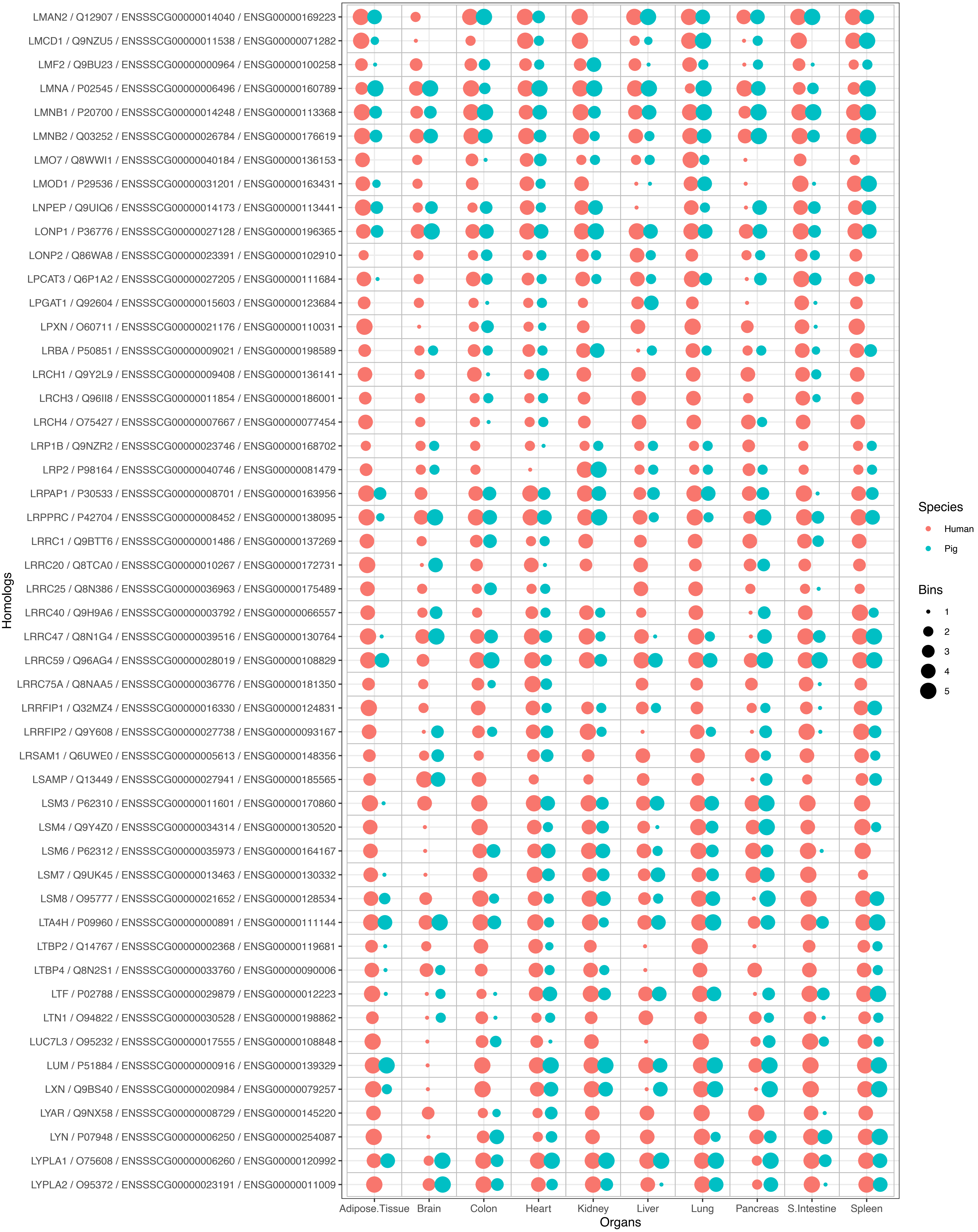
LMAN2 / Q12907 / ENSSSCG00000014040 / ENSG00000169223
LMCD1 / Q9NZU5 / ENSSSCG00000011538 / ENSG00000071282
LMF2 / Q9BU23 / ENSSSCG00000000964 / ENSG00000100258
LMNA / P02545 / ENSSSCG00000006496 / ENSG00000160789
LMNB1 / P20700 / ENSSSCG00000014248 / ENSG00000113368
LMNB2 / Q03252 / ENSSSCG00000026784 / ENSG00000176619
LMO7 / Q8WWI1 / ENSSSCG00000040184 / ENSG00000136153
LMOD1 / P29536 / ENSSSCG00000031201 / ENSG00000163431
LNPEP / Q9UIQ6 / ENSSSCG00000014173 / ENSG00000113441
LONP1 / P36776 / ENSSSCG00000027128 / ENSG00000196365
LONP2 / Q86WA8 / ENSSSCG00000023391 / ENSG00000102910
LPCAT3 / Q6P1A2 / ENSSSCG00000027205 / ENSG00000111684
LPGAT1 / Q92604 / ENSSSCG00000015603 / ENSG00000123684
LPXN / O60711 / ENSSSCG00000021176 / ENSG00000110031
LRBA / P50851 / ENSSSCG00000009021 / ENSG00000198589
LRCH1 / Q9Y2L9 / ENSSSCG00000009408 / ENSG00000136141
LRCH3 / Q96II8 / ENSSSCG00000011854 / ENSG00000186001
LRCH4 / O75427 / ENSSSCG00000007667 / ENSG00000077454
LRP1B / Q9NZR2 / ENSSSCG00000023746 / ENSG00000168702
LRP2 / P98164 / ENSSSCG00000040746 / ENSG00000081479
LRPAP1 / P30533 / ENSSSCG00000008701 / ENSG00000163956
LRPPRC / P42704 / ENSSSCG00000008452 / ENSG00000138095
LRRC1 / Q9BTT6 / ENSSSCG00000001486 / ENSG00000137269
LRRC20 / Q8TCA0 / ENSSSCG00000010267 / ENSG00000172731
LRRC25 / Q8N386 / ENSSSCG00000036963 / ENSG00000175489
LRRC40 / Q9H9A6 / ENSSSCG00000003792 / ENSG00000066557
LRRC47 / Q8N1G4 / ENSSSCG00000039516 / ENSG00000130764
LRRC59 / Q96AG4 / ENSSSCG00000028019 / ENSG00000108829
LRRC75A / Q8NAA5 / ENSSSCG00000036776 / ENSG00000181350
LRRFIP1 / Q32MZ4 / ENSSSCG00000016330 / ENSG00000124831
LRRFIP2 / Q9Y608 / ENSSSCG00000027738 / ENSG00000093167
LRSAM1 / Q6UWE0 / ENSSSCG00000005613 / ENSG00000148356
LSAMP / Q13449 / ENSSSCG00000027941 / ENSG00000185565
LSM3 / P62310 / ENSSSCG00000011601 / ENSG00000170860
LSM4 / Q9Y4Z0 / ENSSSCG00000034314 / ENSG00000130520
LSM6 / P62312 / ENSSSCG00000035973 / ENSG00000164167
LSM7 / Q9UK45 / ENSSSCG00000013463 / ENSG00000130332
LSM8 / O95777 / ENSSSCG00000021652 / ENSG00000128534
LTA4H / P09960 / ENSSSCG00000000891 / ENSG00000111144
LTBP2 / Q14767 / ENSSSCG00000002368 / ENSG00000119681
LTBP4 / Q8N2S1 / ENSSSCG00000033760 / ENSG00000090006
LTF / P02788 / ENSSSCG00000029879 / ENSG00000012223
LTN1 / O94822 / ENSSSCG00000030528 / ENSG00000198862
LUC7L3 / O95232 / ENSSSCG00000017555 / ENSG00000108848
LUM / P51884 / ENSSSCG00000000916 / ENSG00000139329
LXN / Q9BS40 / ENSSSCG00000020984 / ENSG00000079257
LYAR / Q9NX58 / ENSSSCG00000008729 / ENSG00000145220
LYN / P07948 / ENSSSCG00000006250 / ENSG00000254087
LYPLA1 / O75608 / ENSSSCG00000006260 / ENSG00000120992
LYPLA2 / O95372 / ENSSSCG00000023191 / ENSG00000011009
Homologs
Adipose.Tissue
Brain
Colon
Heart
Kidney
Liver
Lung
Pancreas
S.Intestine
Spleen
Organs
Species
Human
Pig
Bins
1
2
3
4
5

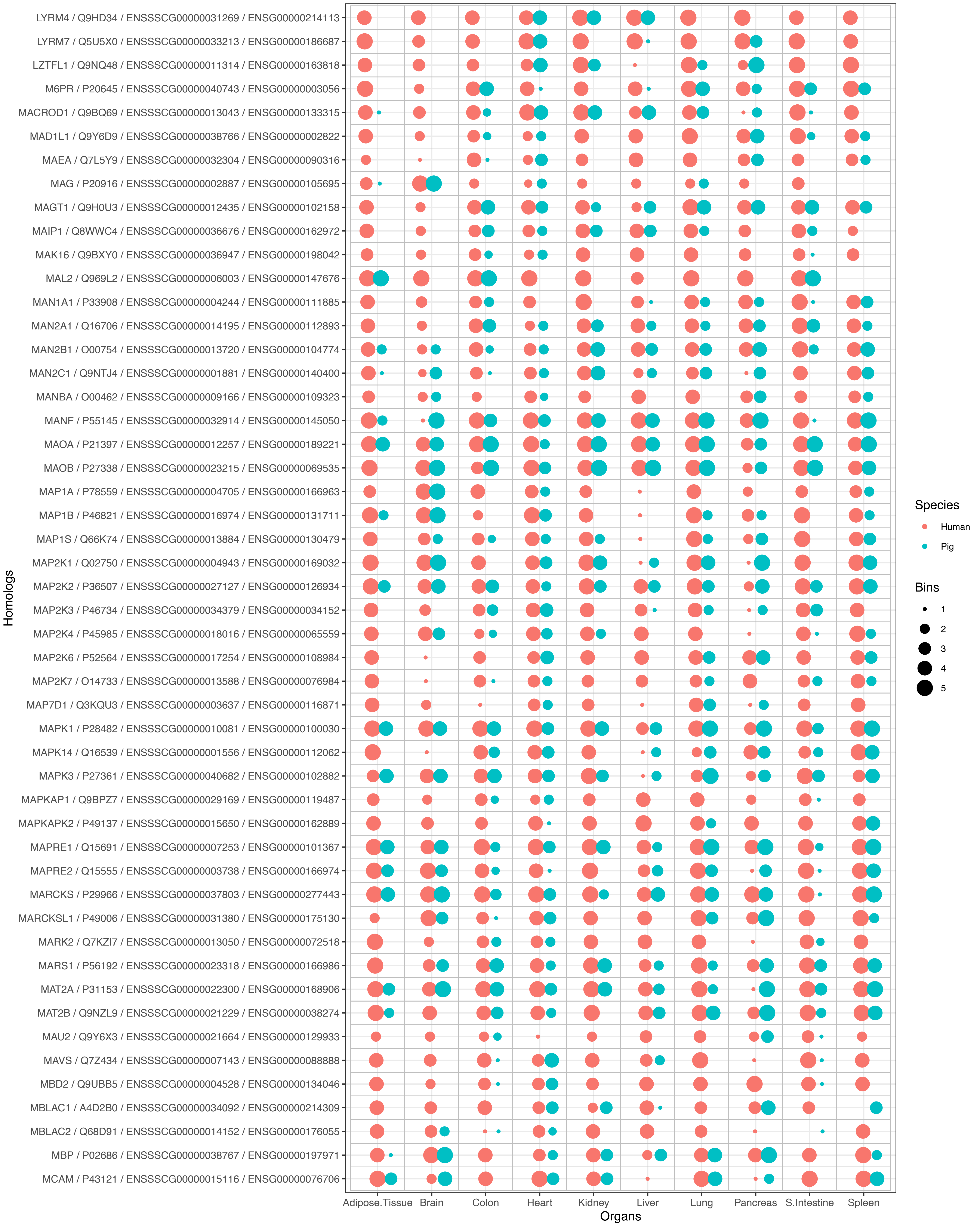
LYRM4 / Q9HD34 / ENSSSCG00000031269 / ENSG00000214113
LYRM7 / Q5U5X0 / ENSSSCG00000033213 / ENSG00000186687
LZTFL1 / Q9NQ48 / ENSSSCG00000011314 / ENSG00000163818
M6PR / P20645 / ENSSSCG00000040743 / ENSG00000003056
MACROD1 / Q9BQ69 / ENSSSCG00000013043 / ENSG00000133315
MAD1L1 / Q9Y6D9 / ENSSSCG00000038766 / ENSG00000002822
MAEA / Q7L5Y9 / ENSSSCG00000032304 / ENSG00000090316
MAG / P20916 / ENSSSCG00000002887 / ENSG00000105695
MAGT1 / Q9H0U3 / ENSSSCG00000012435 / ENSG00000102158
MAIP1 / Q8WWC4 / ENSSSCG00000036676 / ENSG00000162972
MAK16 / Q9BXY0 / ENSSSCG00000036947 / ENSG00000198042
MAL2 / Q969L2 / ENSSSCG00000006003 / ENSG00000147676
MAN1A1 / P33908 / ENSSSCG00000004244 / ENSG00000111885
MAN2A1 / Q16706 / ENSSSCG00000014195 / ENSG00000112893
MAN2B1 / O00754 / ENSSSCG00000013720 / ENSG00000104774
MAN2C1 / Q9NTJ4 / ENSSSCG00000001881 / ENSG00000140400
MANBA / O00462 / ENSSSCG00000009166 / ENSG00000109323
MANF / P55145 / ENSSSCG00000032914 / ENSG00000145050
MAOA / P21397 / ENSSSCG00000012257 / ENSG00000189221
MAOB / P27338 / ENSSSCG00000023215 / ENSG00000069535
MAP1A / P78559 / ENSSSCG00000004705 / ENSG00000166963
MAP1B / P46821 / ENSSSCG00000016974 / ENSG00000131711
MAP1S / Q66K74 / ENSSSCG00000013884 / ENSG00000130479
MAP2K1 / Q02750 / ENSSSCG00000004943 / ENSG00000169032
MAP2K2 / P36507 / ENSSSCG00000027127 / ENSG00000126934
MAP2K3 / P46734 / ENSSSCG00000034379 / ENSG00000034152
MAP2K4 / P45985 / ENSSSCG00000018016 / ENSG00000065559
MAP2K6 / P52564 / ENSSSCG00000017254 / ENSG00000108984
MAP2K7 / O14733 / ENSSSCG00000013588 / ENSG00000076984
MAP7D1 / Q3KQU3 / ENSSSCG00000003637 / ENSG00000116871
MAPK1 / P28482 / ENSSSCG00000010081 / ENSG00000100030
MAPK14 / Q16539 / ENSSSCG00000001556 / ENSG00000112062
MAPK3 / P27361 / ENSSSCG00000040682 / ENSG00000102882
MAPKAP1 / Q9BPZ7 / ENSSSCG00000029169 / ENSG00000119487
MAPKAPK2 / P49137 / ENSSSCG00000015650 / ENSG00000162889
MAPRE1 / Q15691 / ENSSSCG00000007253 / ENSG00000101367
MAPRE2 / Q15555 / ENSSSCG00000003738 / ENSG00000166974
MARCKS / P29966 / ENSSSCG00000037803 / ENSG00000277443
MARCKSL1 / P49006 / ENSSSCG00000031380 / ENSG00000175130
MARK2 / Q7KZI7 / ENSSSCG00000013050 / ENSG00000072518
MARS1 / P56192 / ENSSSCG00000023318 / ENSG00000166986
MAT2A / P31153 / ENSSSCG00000022300 / ENSG00000168906
MAT2B / Q9NZL9 / ENSSSCG00000021229 / ENSG00000038274
MAU2 / Q9Y6X3 / ENSSSCG00000021664 / ENSG00000129933
MAVS / Q7Z434 / ENSSSCG00000007143 / ENSG00000088888
MBD2 / Q9UBB5 / ENSSSCG00000004528 / ENSG00000134046
MBLAC1 / A4D2B0 / ENSSSCG00000034092 / ENSG00000214309
MBLAC2 / Q68D91 / ENSSSCG00000014152 / ENSG00000176055
MBP / P02686 / ENSSSCG00000038767 / ENSG00000197971
MCAM / P43121 / ENSSSCG00000015116 / ENSG00000076706
Homologs
Adipose.Tissue
Brain
Colon
Heart
Kidney
Liver
Lung
Pancreas
S.Intestine
Spleen
Organs
Species
Human
Pig
Bins
1
2
3
4
5

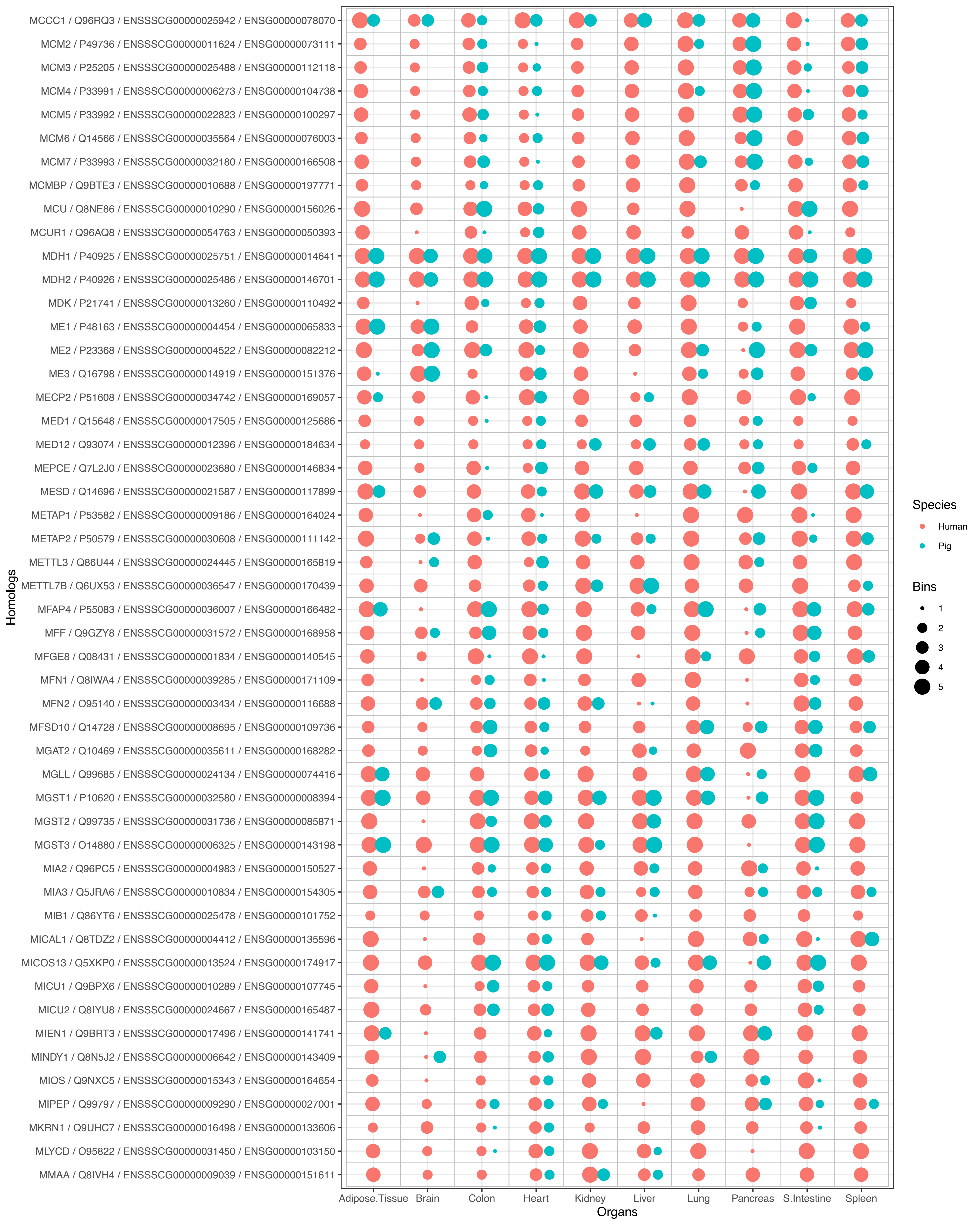
MCCC1 / Q96RQ3 / ENSSSCG00000025942 / ENSG00000078070
MCM2 / P49736 / ENSSSCG00000011624 / ENSG00000073111
MCM3 / P25205 / ENSSSCG00000025488 / ENSG00000112118
MCM4 / P33991 / ENSSSCG00000006273 / ENSG00000104738
MCM5 / P33992 / ENSSSCG00000022823 / ENSG00000100297
MCM6 / Q14566 / ENSSSCG00000035564 / ENSG00000076003
MCM7 / P33993 / ENSSSCG00000032180 / ENSG00000166508
MCMBP / Q9BTE3 / ENSSSCG00000010688 / ENSG00000197771
MCU / Q8NE86 / ENSSSCG00000010290 / ENSG00000156026
MCUR1 / Q96AQ8 / ENSSSCG00000054763 / ENSG00000050393
MDH1 / P40925 / ENSSSCG00000025751 / ENSG00000014641
MDH2 / P40926 / ENSSSCG00000025486 / ENSG00000146701
MDK / P21741 / ENSSSCG00000013260 / ENSG00000110492
ME1 / P48163 / ENSSSCG00000004454 / ENSG00000065833
ME2 / P23368 / ENSSSCG00000004522 / ENSG00000082212
ME3 / Q16798 / ENSSSCG00000014919 / ENSG00000151376
MECP2 / P51608 / ENSSSCG00000034742 / ENSG00000169057
MED1 / Q15648 / ENSSSCG00000017505 / ENSG00000125686
MED12 / Q93074 / ENSSSCG00000012396 / ENSG00000184634
MEPCE / Q7L2J0 / ENSSSCG00000023680 / ENSG00000146834
MESD / Q14696 / ENSSSCG00000021587 / ENSG00000117899
METAP1 / P53582 / ENSSSCG00000009186 / ENSG00000164024
METAP2 / P50579 / ENSSSCG00000030608 / ENSG00000111142
METTL3 / Q86U44 / ENSSSCG00000024445 / ENSG00000165819
METTL7B / Q6UX53 / ENSSSCG00000036547 / ENSG00000170439
MFAP4 / P55083 / ENSSSCG00000036007 / ENSG00000166482
MFF / Q9GZY8 / ENSSSCG00000031572 / ENSG00000168958
MFGE8 / Q08431 / ENSSSCG00000001834 / ENSG00000140545
MFN1 / Q8IWA4 / ENSSSCG00000039285 / ENSG00000171109
MFN2 / O95140 / ENSSSCG00000003434 / ENSG00000116688
MFSD10 / Q14728 / ENSSSCG00000008695 / ENSG00000109736
MGAT2 / Q10469 / ENSSSCG00000035611 / ENSG00000168282
MGLL / Q99685 / ENSSSCG00000024134 / ENSG00000074416
MGST1 / P10620 / ENSSSCG00000032580 / ENSG00000008394
MGST2 / Q99735 / ENSSSCG00000031736 / ENSG00000085871
MGST3 / O14880 / ENSSSCG00000006325 / ENSG00000143198
MIA2 / Q96PC5 / ENSSSCG00000004983 / ENSG00000150527
MIA3 / Q5JRA6 / ENSSSCG00000010834 / ENSG00000154305
MIB1 / Q86YT6 / ENSSSCG00000025478 / ENSG00000101752
MICAL1 / Q8TDZ2 / ENSSSCG00000004412 / ENSG00000135596
MICOS13 / Q5XKP0 / ENSSSCG00000013524 / ENSG00000174917
MICU1 / Q9BPX6 / ENSSSCG00000010289 / ENSG00000107745
MICU2 / Q8IYU8 / ENSSSCG00000024667 / ENSG00000165487
MIEN1 / Q9BRT3 / ENSSSCG00000017496 / ENSG00000141741
MINDY1 / Q8N5J2 / ENSSSCG00000006642 / ENSG00000143409
MIOS / Q9NXC5 / ENSSSCG00000015343 / ENSG00000164654
MIPEP / Q99797 / ENSSSCG00000009290 / ENSG00000027001
MKRN1 / Q9UHC7 / ENSSSCG00000016498 / ENSG00000133606
MLYCD / O95822 / ENSSSCG00000031450 / ENSG00000103150
MMAA / Q8IVH4 / ENSSSCG00000009039 / ENSG00000151611
Homologs
Adipose.Tissue
Brain
Colon
Heart
Kidney
Liver
Lung
Pancreas
S.Intestine
Spleen
Organs
Species
Human
Pig
Bins
1
2
3
4
5

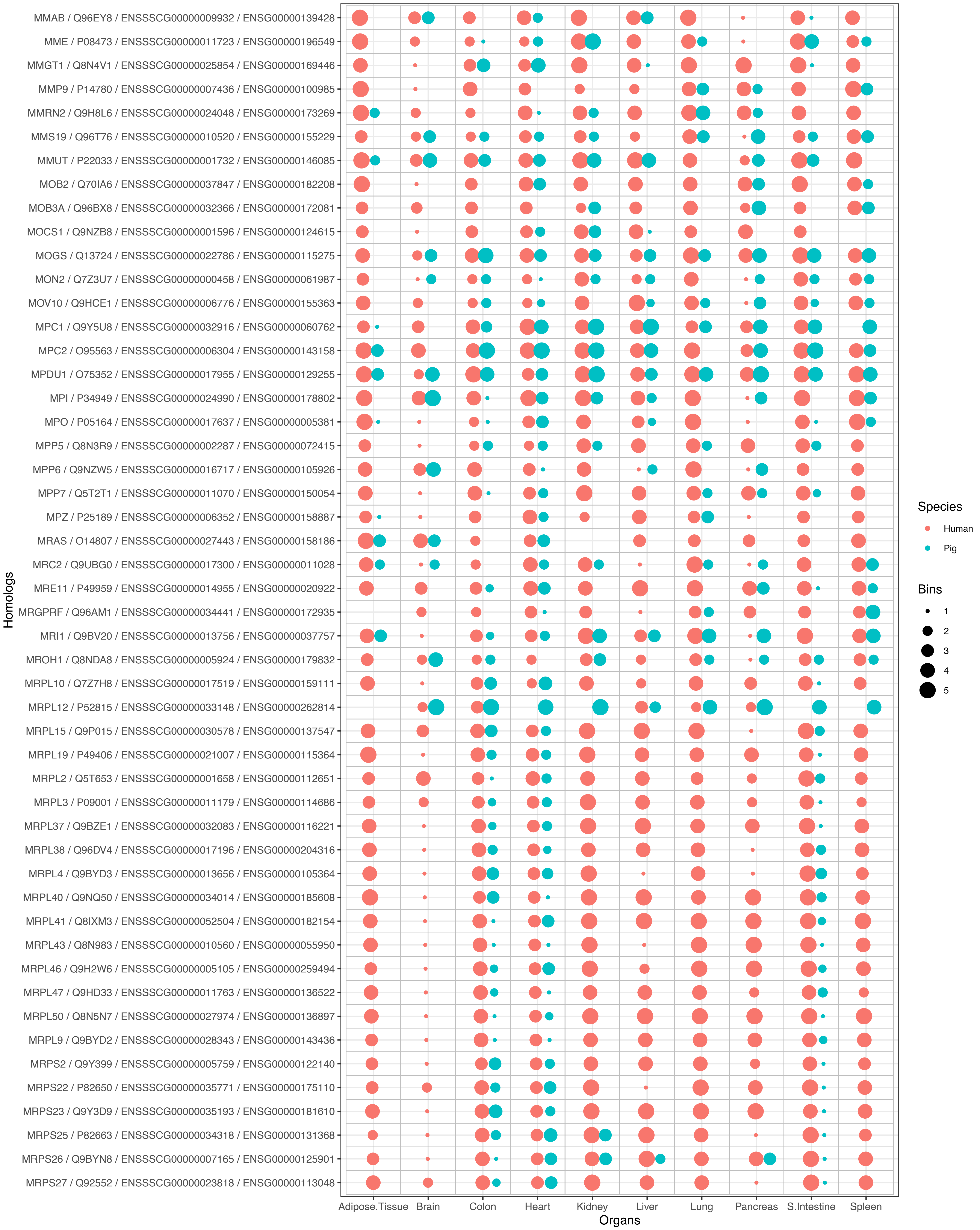
MMAB / Q96EY8 / ENSSSCG00000009932 / ENSG00000139428
MME / P08473 / ENSSSCG00000011723 / ENSG00000196549
MMGT1 / Q8N4V1 / ENSSSCG00000025854 / ENSG00000169446
MMP9 / P14780 / ENSSSCG00000007436 / ENSG00000100985
MMRN2 / Q9H8L6 / ENSSSCG00000024048 / ENSG00000173269
MMS19 / Q96T76 / ENSSSCG00000010520 / ENSG00000155229
MMUT / P22033 / ENSSSCG00000001732 / ENSG00000146085
MOB2 / Q70IA6 / ENSSSCG00000037847 / ENSG00000182208
MOB3A / Q96BX8 / ENSSSCG00000032366 / ENSG00000172081
MOCS1 / Q9NZB8 / ENSSSCG00000001596 / ENSG00000124615
MOGS / Q13724 / ENSSSCG00000022786 / ENSG00000115275
MON2 / Q7Z3U7 / ENSSSCG00000000458 / ENSG00000061987
MOV10 / Q9HCE1 / ENSSSCG00000006776 / ENSG00000155363
MPC1 / Q9Y5U8 / ENSSSCG00000032916 / ENSG00000060762
MPC2 / O95563 / ENSSSCG00000006304 / ENSG00000143158
MPDU1 / O75352 / ENSSSCG00000017955 / ENSG00000129255
MPI / P34949 / ENSSSCG00000024990 / ENSG00000178802
MPO / P05164 / ENSSSCG00000017637 / ENSG00000005381
MPP5 / Q8N3R9 / ENSSSCG00000002287 / ENSG00000072415
MPP6 / Q9NZW5 / ENSSSCG00000016717 / ENSG00000105926
MPP7 / Q5T2T1 / ENSSSCG00000011070 / ENSG00000150054
MPZ / P25189 / ENSSSCG00000006352 / ENSG00000158887
MRAS / O14807 / ENSSSCG00000027443 / ENSG00000158186
MRC2 / Q9UBG0 / ENSSSCG00000017300 / ENSG00000011028
MRE11 / P49959 / ENSSSCG00000014955 / ENSG00000020922
MRGPRF / Q96AM1 / ENSSSCG00000034441 / ENSG00000172935
MRI1 / Q9BV20 / ENSSSCG00000013756 / ENSG00000037757
MROH1 / Q8NDA8 / ENSSSCG00000005924 / ENSG00000179832
MRPL10 / Q7Z7H8 / ENSSSCG00000017519 / ENSG00000159111
MRPL12 / P52815 / ENSSSCG00000033148 / ENSG00000262814
MRPL15 / Q9P015 / ENSSSCG00000030578 / ENSG00000137547
MRPL19 / P49406 / ENSSSCG00000021007 / ENSG00000115364
MRPL2 / Q5T653 / ENSSSCG00000001658 / ENSG00000112651
MRPL3 / P09001 / ENSSSCG00000011179 / ENSG00000114686
MRPL37 / Q9BZE1 / ENSSSCG00000032083 / ENSG00000116221
MRPL38 / Q96DV4 / ENSSSCG00000017196 / ENSG00000204316
MRPL4 / Q9BYD3 / ENSSSCG00000013656 / ENSG00000105364
MRPL40 / Q9NQ50 / ENSSSCG00000034014 / ENSG00000185608
MRPL41 / Q8IXM3 / ENSSSCG00000052504 / ENSG00000182154
MRPL43 / Q8N983 / ENSSSCG00000010560 / ENSG00000055950
MRPL46 / Q9H2W6 / ENSSSCG00000005105 / ENSG00000259494
MRPL47 / Q9HD33 / ENSSSCG00000011763 / ENSG00000136522
MRPL50 / Q8N5N7 / ENSSSCG00000027974 / ENSG00000136897
MRPL9 / Q9BYD2 / ENSSSCG00000028343 / ENSG00000143436
MRPS2 / Q9Y399 / ENSSSCG00000005759 / ENSG00000122140
MRPS22 / P82650 / ENSSSCG00000035771 / ENSG00000175110
MRPS23 / Q9Y3D9 / ENSSSCG00000035193 / ENSG00000181610
MRPS25 / P82663 / ENSSSCG00000034318 / ENSG00000131368
MRPS26 / Q9BYN8 / ENSSSCG00000007165 / ENSG00000125901
MRPS27 / Q92552 / ENSSSCG00000023818 / ENSG00000113048
Homologs
Adipose.Tissue
Brain
Colon
Heart
Kidney
Liver
Lung
Pancreas
S.Intestine
Spleen
Organs
Species
Human
Pig
Bins
1
2
3
4
5

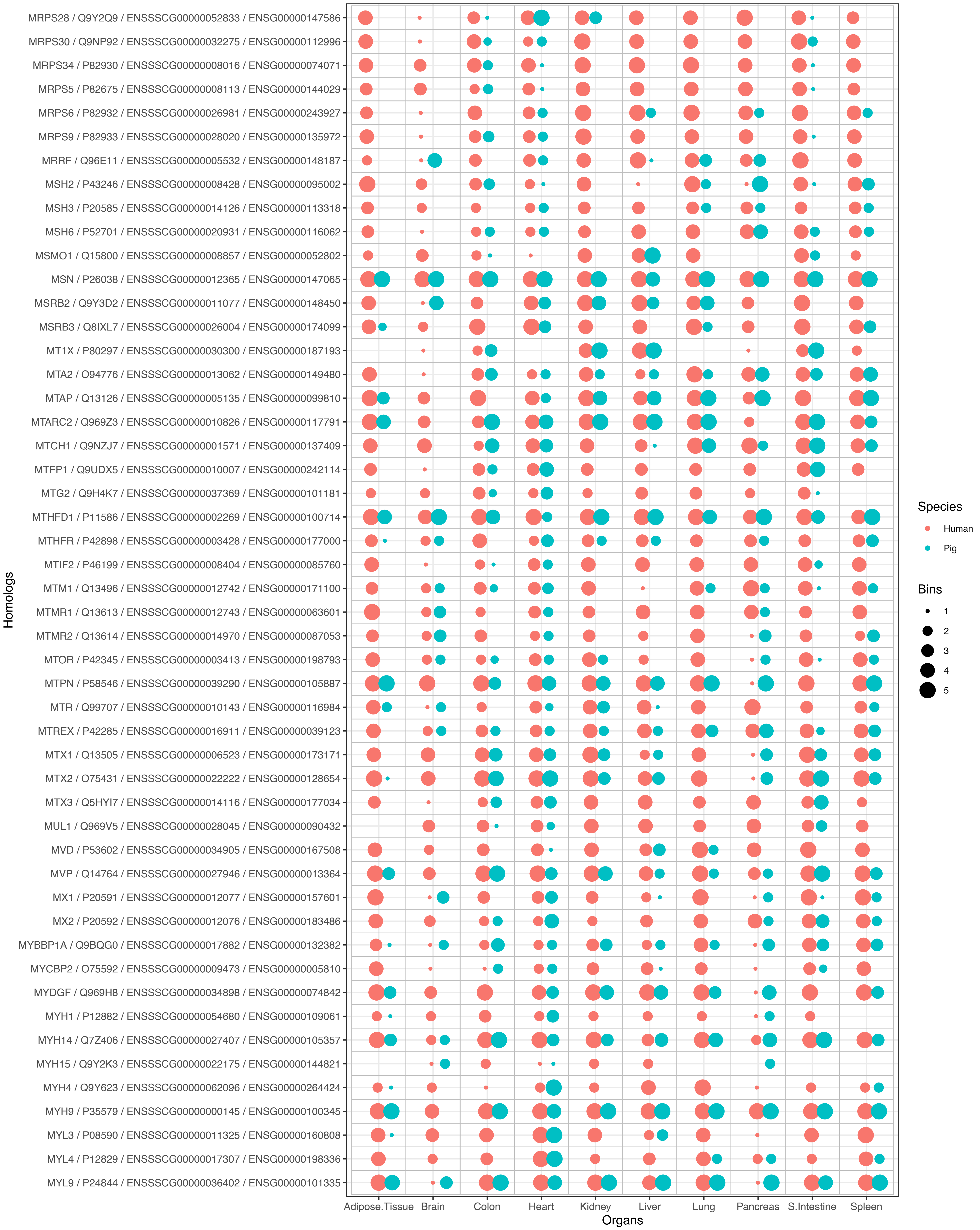
MRPS28 / Q9Y2Q9 / ENSSSCG00000052833 / ENSG00000147586
MRPS30 / Q9NP92 / ENSSSCG00000032275 / ENSG00000112996
MRPS34 / P82930 / ENSSSCG00000008016 / ENSG00000074071
MRPS5 / P82675 / ENSSSCG00000008113 / ENSG00000144029
MRPS6 / P82932 / ENSSSCG00000026981 / ENSG00000243927
MRPS9 / P82933 / ENSSSCG00000028020 / ENSG00000135972
MRRF / Q96E11 / ENSSSCG00000005532 / ENSG00000148187
MSH2 / P43246 / ENSSSCG00000008428 / ENSG00000095002
MSH3 / P20585 / ENSSSCG00000014126 / ENSG00000113318
MSH6 / P52701 / ENSSSCG00000020931 / ENSG00000116062
MSMO1 / Q15800 / ENSSSCG00000008857 / ENSG00000052802
MSN / P26038 / ENSSSCG00000012365 / ENSG00000147065
MSRB2 / Q9Y3D2 / ENSSSCG00000011077 / ENSG00000148450
MSRB3 / Q8IXL7 / ENSSSCG00000026004 / ENSG00000174099
MT1X / P80297 / ENSSSCG00000030300 / ENSG00000187193
MTA2 / O94776 / ENSSSCG00000013062 / ENSG00000149480
MTAP / Q13126 / ENSSSCG00000005135 / ENSG00000099810
MTARC2 / Q969Z3 / ENSSSCG00000010826 / ENSG00000117791
MTCH1 / Q9NZJ7 / ENSSSCG00000001571 / ENSG00000137409
MTFP1 / Q9UDX5 / ENSSSCG00000010007 / ENSG00000242114
MTG2 / Q9H4K7 / ENSSSCG00000037369 / ENSG00000101181
MTHFD1 / P11586 / ENSSSCG00000002269 / ENSG00000100714
MTHFR / P42898 / ENSSSCG00000003428 / ENSG00000177000
MTIF2 / P46199 / ENSSSCG00000008404 / ENSG00000085760
MTM1 / Q13496 / ENSSSCG00000012742 / ENSG00000171100
MTMR1 / Q13613 / ENSSSCG00000012743 / ENSG00000063601
MTMR2 / Q13614 / ENSSSCG00000014970 / ENSG00000087053
MTOR / P42345 / ENSSSCG00000003413 / ENSG00000198793
MTPN / P58546 / ENSSSCG00000039290 / ENSG00000105887
MTR / Q99707 / ENSSSCG00000010143 / ENSG00000116984
MTREX / P42285 / ENSSSCG00000016911 / ENSG00000039123
MTX1 / Q13505 / ENSSSCG00000006523 / ENSG00000173171
MTX2 / O75431 / ENSSSCG00000022222 / ENSG00000128654
MTX3 / Q5HYI7 / ENSSSCG00000014116 / ENSG00000177034
MUL1 / Q969V5 / ENSSSCG00000028045 / ENSG00000090432
MVD / P53602 / ENSSSCG00000034905 / ENSG00000167508
MVP / Q14764 / ENSSSCG00000027946 / ENSG00000013364
MX1 / P20591 / ENSSSCG00000012077 / ENSG00000157601
MX2 / P20592 / ENSSSCG00000012076 / ENSG00000183486
MYBBP1A / Q9BQG0 / ENSSSCG00000017882 / ENSG00000132382
MYCBP2 / O75592 / ENSSSCG00000009473 / ENSG00000005810
MYDGF / Q969H8 / ENSSSCG00000034898 / ENSG00000074842
MYH1 / P12882 / ENSSSCG00000054680 / ENSG00000109061
MYH14 / Q7Z406 / ENSSSCG00000027407 / ENSG00000105357
MYH15 / Q9Y2K3 / ENSSSCG00000022175 / ENSG00000144821
MYH4 / Q9Y623 / ENSSSCG00000062096 / ENSG00000264424
MYH9 / P35579 / ENSSSCG00000000145 / ENSG00000100345
MYL3 / P08590 / ENSSSCG00000011325 / ENSG00000160808
MYL4 / P12829 / ENSSSCG00000017307 / ENSG00000198336
MYL9 / P24844 / ENSSSCG00000036402 / ENSG00000101335
Homologs
Adipose.Tissue
Brain
Colon
Heart
Kidney
Liver
Lung
Pancreas
S.Intestine
Spleen
Organs
Species
Human
Pig
Bins
1
2
3
4
5

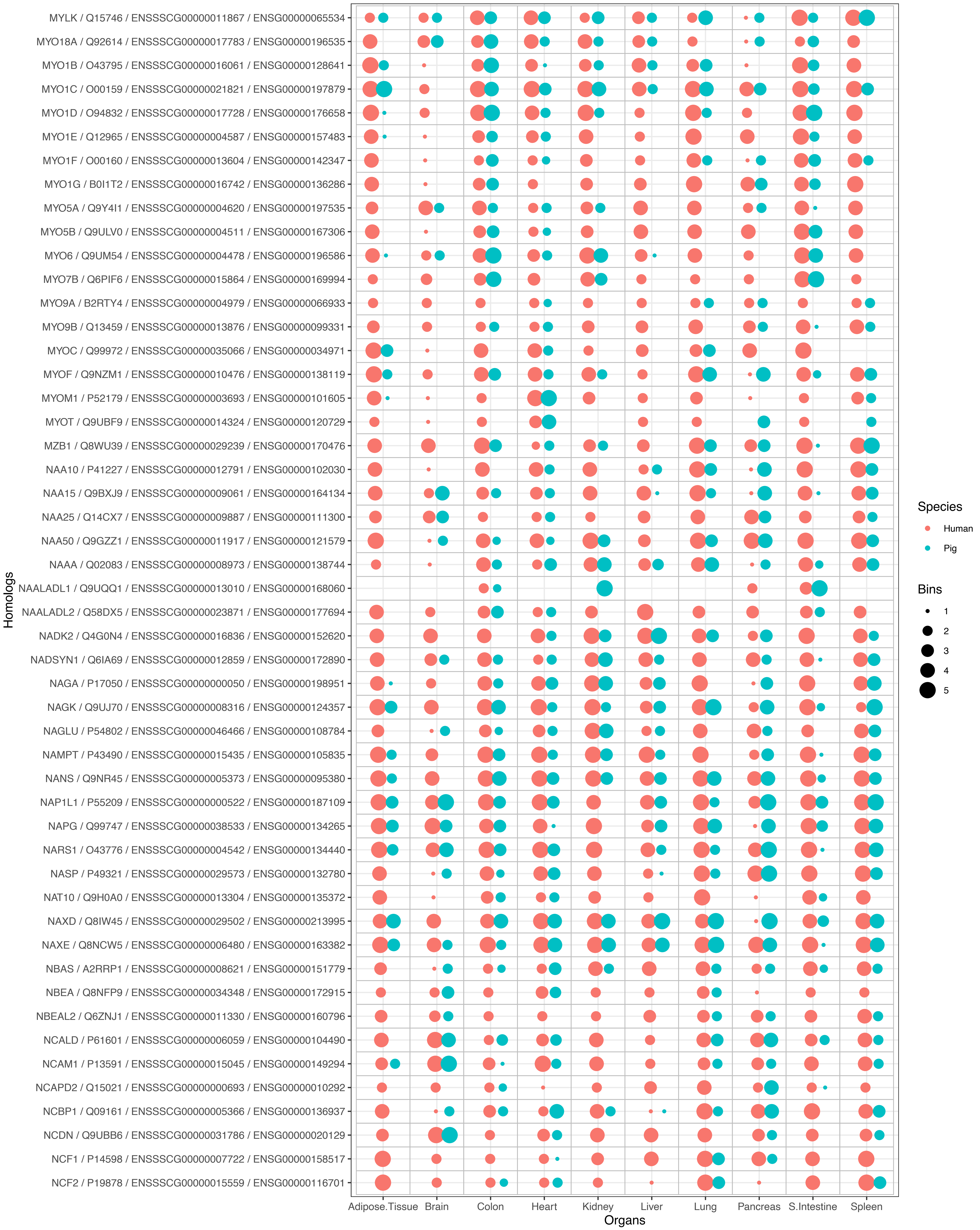

MYLK / Q15746 / ENSSSCG00000011867 / ENSG00000065534
MYO18A / Q92614 / ENSSSCG00000017783 / ENSG00000196535
MYO1B / O43795 / ENSSSCG00000016061 / ENSG00000128641
MYO1C / O00159 / ENSSSCG00000021821 / ENSG00000197879
MYO1D / O94832 / ENSSSCG00000017728 / ENSG00000176658
MYO1E / Q12965 / ENSSSCG00000004587 / ENSG00000157483
MYO1F / O00160 / ENSSSCG00000013604 / ENSG00000142347
MYO1G / B0I1T2 / ENSSSCG00000016742 / ENSG00000136286
MYO5A / Q9Y4I1 / ENSSSCG00000004620 / ENSG00000197535
MYO5B / Q9ULV0 / ENSSSCG00000004511 / ENSG00000167306
MYO6 / Q9UM54 / ENSSSCG00000004478 / ENSG00000196586
MYO7B / Q6PIF6 / ENSSSCG00000015864 / ENSG00000169994
MYO9A / B2RTY4 / ENSSSCG00000004979 / ENSG00000066933
MYO9B / Q13459 / ENSSSCG00000013876 / ENSG00000099331
MYOC / Q99972 / ENSSSCG00000035066 / ENSG00000034971
MYOF / Q9NZM1 / ENSSSCG00000010476 / ENSG00000138119
MYOM1 / P52179 / ENSSSCG00000003693 / ENSG00000101605
MYOT / Q9UBF9 / ENSSSCG00000014324 / ENSG00000120729
MZB1 / Q8WU39 / ENSSSCG00000029239 / ENSG00000170476
NAA10 / P41227 / ENSSSCG00000012791 / ENSG00000102030
NAA15 / Q9BXJ9 / ENSSSCG00000009061 / ENSG00000164134
NAA25 / Q14CX7 / ENSSSCG00000009887 / ENSG00000111300
NAA50 / Q9GZZ1 / ENSSSCG00000011917 / ENSG00000121579
NAAA / Q02083 / ENSSSCG00000008973 / ENSG00000138744
NAALADL1 / Q9UQQ1 / ENSSSCG00000013010 / ENSG00000168060
NAALADL2 / Q58DX5 / ENSSSCG00000023871 / ENSG00000177694
NADK2 / Q4G0N4 / ENSSSCG00000016836 / ENSG00000152620
NADSYN1 / Q6IA69 / ENSSSCG00000012859 / ENSG00000172890
NAGA / P17050 / ENSSSCG00000000050 / ENSG00000198951
NAGK / Q9UJ70 / ENSSSCG00000008316 / ENSG00000124357
NAGLU / P54802 / ENSSSCG00000046466 / ENSG00000108784
NAMPT / P43490 / ENSSSCG00000015435 / ENSG00000105835
NANS / Q9NR45 / ENSSSCG00000005373 / ENSG00000095380
NAP1L1 / P55209 / ENSSSCG00000000522 / ENSG00000187109
NAPG / Q99747 / ENSSSCG00000038533 / ENSG00000134265
NARS1 / O43776 / ENSSSCG00000004542 / ENSG00000134440
NASP / P49321 / ENSSSCG00000029573 / ENSG00000132780
NAT10 / Q9H0A0 / ENSSSCG00000013304 / ENSG00000135372
NAXD / Q8IW45 / ENSSSCG00000029502 / ENSG00000213995
NAXE / Q8NCW5 / ENSSSCG00000006480 / ENSG00000163382
NBAS / A2RRP1 / ENSSSCG00000008621 / ENSG00000151779
NBEA / Q8NFP9 / ENSSSCG00000034348 / ENSG00000172915
NBEAL2 / Q6ZNJ1 / ENSSSCG00000011330 / ENSG00000160796
NCALD / P61601 / ENSSSCG00000006059 / ENSG00000104490
NCAM1 / P13591 / ENSSSCG00000015045 / ENSG00000149294
NCAPD2 / Q15021 / ENSSSCG00000000693 / ENSG00000010292
NCBP1 / Q09161 / ENSSSCG00000005366 / ENSG00000136937
NCDN / Q9UBB6 / ENSSSCG00000031786 / ENSG00000020129
NCF1 / P14598 / ENSSSCG00000007722 / ENSG00000158517
NCF2 / P19878 / ENSSSCG00000015559 / ENSG00000116701
Homologs
Adipose.Tissue
Brain
Colon
Heart
Kidney
Liver
Lung
Pancreas
S.Intestine
Spleen
Organs
Species
Human
Pig
Bins
1
2
3
4
5

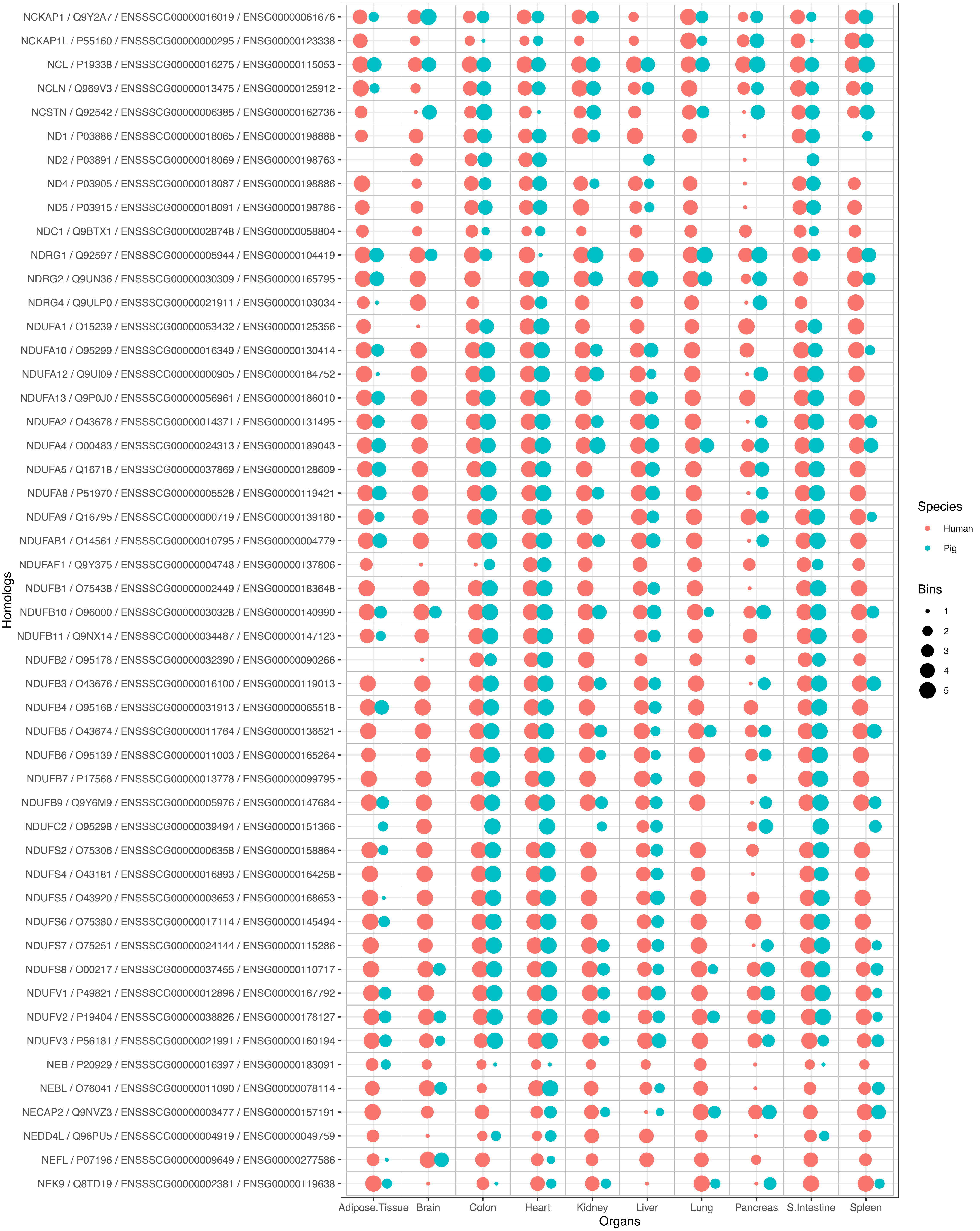
NCKAP1 / Q9Y2A7 / ENSSSCG00000016019 / ENSG00000061676
NCKAP1L / P55160 / ENSSSCG00000000295 / ENSG00000123338
NCL / P19338 / ENSSSCG00000016275 / ENSG00000115053
NCLN / Q969V3 / ENSSSCG00000013475 / ENSG00000125912
NCSTN / Q92542 / ENSSSCG00000006385 / ENSG00000162736
ND1 / P03886 / ENSSSCG00000018065 / ENSG00000198888
ND2 / P03891 / ENSSSCG00000018069 / ENSG00000198763
ND4 / P03905 / ENSSSCG00000018087 / ENSG00000198886
ND5 / P03915 / ENSSSCG00000018091 / ENSG00000198786
NDC1 / Q9BTX1 / ENSSSCG00000028748 / ENSG00000058804
NDRG1 / Q92597 / ENSSSCG00000005944 / ENSG00000104419
NDRG2 / Q9UN36 / ENSSSCG00000030309 / ENSG00000165795
NDRG4 / Q9ULP0 / ENSSSCG00000021911 / ENSG00000103034
NDUFA1 / O15239 / ENSSSCG00000053432 / ENSG00000125356
NDUFA10 / O95299 / ENSSSCG00000016349 / ENSG00000130414
NDUFA12 / Q9UI09 / ENSSSCG00000000905 / ENSG00000184752
NDUFA13 / Q9P0J0 / ENSSSCG00000056961 / ENSG00000186010
NDUFA2 / O43678 / ENSSSCG00000014371 / ENSG00000131495
NDUFA4 / O00483 / ENSSSCG00000024313 / ENSG00000189043
NDUFA5 / Q16718 / ENSSSCG00000037869 / ENSG00000128609
NDUFA8 / P51970 / ENSSSCG00000005528 / ENSG00000119421
NDUFA9 / Q16795 / ENSSSCG00000000719 / ENSG00000139180
NDUFAB1 / O14561 / ENSSSCG00000010795 / ENSG00000004779
NDUFAF1 / Q9Y375 / ENSSSCG00000004748 / ENSG00000137806
NDUFB1 / O75438 / ENSSSCG00000002449 / ENSG00000183648
NDUFB10 / O96000 / ENSSSCG00000030328 / ENSG00000140990
NDUFB11 / Q9NX14 / ENSSSCG00000034487 / ENSG00000147123
NDUFB2 / O95178 / ENSSSCG00000032390 / ENSG00000090266
NDUFB3 / O43676 / ENSSSCG00000016100 / ENSG00000119013
NDUFB4 / O95168 / ENSSSCG00000031913 / ENSG00000065518
NDUFB5 / O43674 / ENSSSCG00000011764 / ENSG00000136521
NDUFB6 / O95139 / ENSSSCG00000011003 / ENSG00000165264
NDUFB7 / P17568 / ENSSSCG00000013778 / ENSG00000099795
NDUFB9 / Q9Y6M9 / ENSSSCG00000005976 / ENSG00000147684
NDUFC2 / O95298 / ENSSSCG00000039494 / ENSG00000151366
NDUFS2 / O75306 / ENSSSCG00000006358 / ENSG00000158864
NDUFS4 / O43181 / ENSSSCG00000016893 / ENSG00000164258
NDUFS5 / O43920 / ENSSSCG00000003653 / ENSG00000168653
NDUFS6 / O75380 / ENSSSCG00000017114 / ENSG00000145494
NDUFS7 / O75251 / ENSSSCG00000024144 / ENSG00000115286
NDUFS8 / O00217 / ENSSSCG00000037455 / ENSG00000110717
NDUFV1 / P49821 / ENSSSCG00000012896 / ENSG00000167792
NDUFV2 / P19404 / ENSSSCG00000038826 / ENSG00000178127
NDUFV3 / P56181 / ENSSSCG00000021991 / ENSG00000160194
NEB / P20929 / ENSSSCG00000016397 / ENSG00000183091
NEBL / O76041 / ENSSSCG00000011090 / ENSG00000078114
NECAP2 / Q9NVZ3 / ENSSSCG00000003477 / ENSG00000157191
NEDD4L / Q96PU5 / ENSSSCG00000004919 / ENSG00000049759
NEFL / P07196 / ENSSSCG00000009649 / ENSG00000277586
NEK9 / Q8TD19 / ENSSSCG00000002381 / ENSG00000119638
Homologs
Adipose.Tissue
Brain
Colon
Heart
Kidney
Liver
Lung
Pancreas
S.Intestine
Spleen
Organs
Species
Human
Pig
Bins
1
2
3
4
5

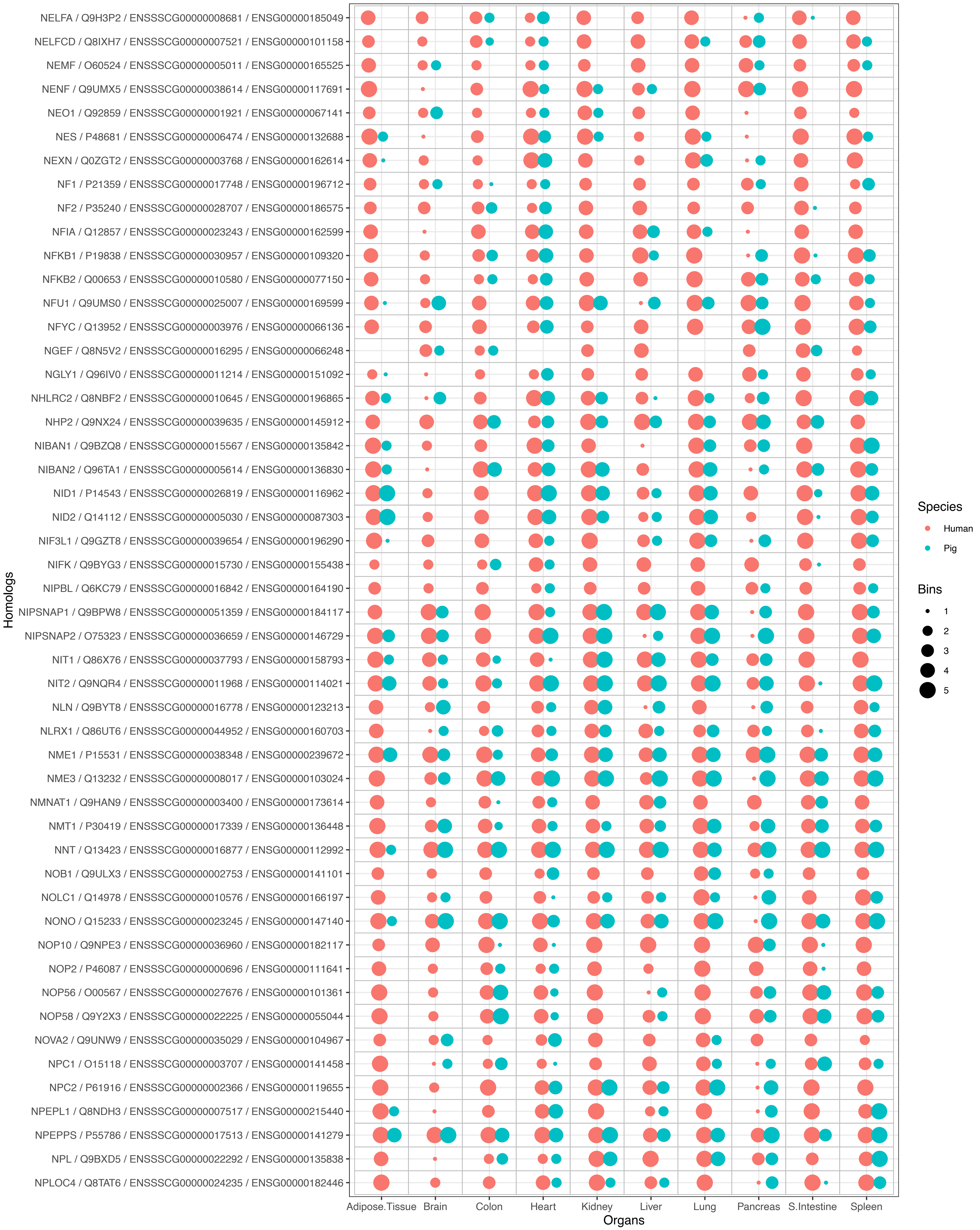

NELFA / Q9H3P2 / ENSSSCG00000008681 / ENSG00000185049
NELFCD / Q8IXH7 / ENSSSCG00000007521 / ENSG00000101158
NEMF / O60524 / ENSSSCG00000005011 / ENSG00000165525
NENF / Q9UMX5 / ENSSSCG00000038614 / ENSG00000117691
NEO1 / Q92859 / ENSSSCG00000001921 / ENSG00000067141
NES / P48681 / ENSSSCG00000006474 / ENSG00000132688
NEXN / Q0ZGT2 / ENSSSCG00000003768 / ENSG00000162614
NF1 / P21359 / ENSSSCG00000017748 / ENSG00000196712
NF2 / P35240 / ENSSSCG00000028707 / ENSG00000186575
NFIA / Q12857 / ENSSSCG00000023243 / ENSG00000162599
NFKB1 / P19838 / ENSSSCG00000030957 / ENSG00000109320
NFKB2 / Q00653 / ENSSSCG00000010580 / ENSG00000077150
NFU1 / Q9UMS0 / ENSSSCG00000025007 / ENSG00000169599
NFYC / Q13952 / ENSSSCG00000003976 / ENSG00000066136
NGEF / Q8N5V2 / ENSSSCG00000016295 / ENSG00000066248
NGLY1 / Q96IV0 / ENSSSCG00000011214 / ENSG00000151092
NHLRC2 / Q8NBF2 / ENSSSCG00000010645 / ENSG00000196865
NHP2 / Q9NX24 / ENSSSCG00000039635 / ENSG00000145912
NIBAN1 / Q9BZQ8 / ENSSSCG00000015567 / ENSG00000135842
NIBAN2 / Q96TA1 / ENSSSCG00000005614 / ENSG00000136830
NID1 / P14543 / ENSSSCG00000026819 / ENSG00000116962
NID2 / Q14112 / ENSSSCG00000005030 / ENSG00000087303
NIF3L1 / Q9GZT8 / ENSSSCG00000039654 / ENSG00000196290
NIFK / Q9BYG3 / ENSSSCG00000015730 / ENSG00000155438
NIPBL / Q6KC79 / ENSSSCG00000016842 / ENSG00000164190
NIPSNAP1 / Q9BPW8 / ENSSSCG00000051359 / ENSG00000184117
NIPSNAP2 / O75323 / ENSSSCG00000036659 / ENSG00000146729
NIT1 / Q86X76 / ENSSSCG00000037793 / ENSG00000158793
NIT2 / Q9NQR4 / ENSSSCG00000011968 / ENSG00000114021
NLN / Q9BYT8 / ENSSSCG00000016778 / ENSG00000123213
NLRX1 / Q86UT6 / ENSSSCG00000044952 / ENSG00000160703
NME1 / P15531 / ENSSSCG00000038348 / ENSG00000239672
NME3 / Q13232 / ENSSSCG00000008017 / ENSG00000103024
NMNAT1 / Q9HAN9 / ENSSSCG00000003400 / ENSG00000173614
NMT1 / P30419 / ENSSSCG00000017339 / ENSG00000136448
NNT / Q13423 / ENSSSCG00000016877 / ENSG00000112992
NOB1 / Q9ULX3 / ENSSSCG00000002753 / ENSG00000141101
NOLC1 / Q14978 / ENSSSCG00000010576 / ENSG00000166197
NONO / Q15233 / ENSSSCG00000023245 / ENSG00000147140
NOP10 / Q9NPE3 / ENSSSCG00000036960 / ENSG00000182117
NOP2 / P46087 / ENSSSCG00000000696 / ENSG00000111641
NOP56 / O00567 / ENSSSCG00000027676 / ENSG00000101361
NOP58 / Q9Y2X3 / ENSSSCG00000022225 / ENSG00000055044
NOVA2 / Q9UNW9 / ENSSSCG00000035029 / ENSG00000104967
NPC1 / O15118 / ENSSSCG00000003707 / ENSG00000141458
NPC2 / P61916 / ENSSSCG00000002366 / ENSG00000119655
NPEPL1 / Q8NDH3 / ENSSSCG00000007517 / ENSG00000215440
NPEPPS / P55786 / ENSSSCG00000017513 / ENSG00000141279
NPL / Q9BXD5 / ENSSSCG00000022292 / ENSG00000135838
NPLOC4 / Q8TAT6 / ENSSSCG00000024235 / ENSG00000182446
Homologs
Adipose.Tissue
Brain
Colon
Heart
Kidney
Liver
Lung
Pancreas
S.Intestine
Spleen
Organs
Species
Human
Pig
Bins
1
2
3
4
5

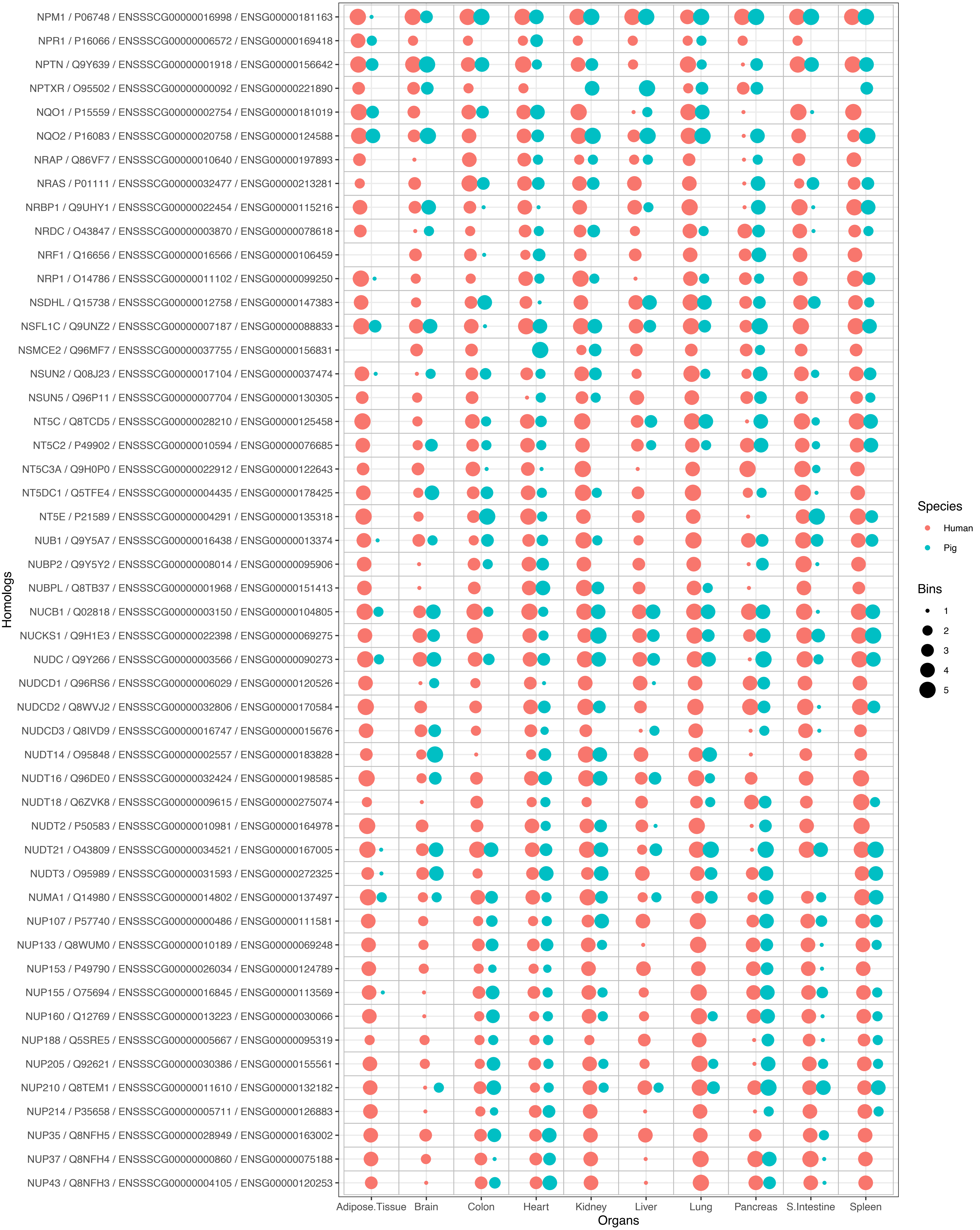

NPM1 / P06748 / ENSSSCG00000016998 / ENSG00000181163
NPR1 / P16066 / ENSSSCG00000006572 / ENSG00000169418
NPTN / Q9Y639 / ENSSSCG00000001918 / ENSG00000156642
NPTXR / O95502 / ENSSSCG00000000092 / ENSG00000221890
NQO1 / P15559 / ENSSSCG00000002754 / ENSG00000181019
NQO2 / P16083 / ENSSSCG00000020758 / ENSG00000124588
NRAP / Q86VF7 / ENSSSCG00000010640 / ENSG00000197893
NRAS / P01111 / ENSSSCG00000032477 / ENSG00000213281
NRBP1 / Q9UHY1 / ENSSSCG00000022454 / ENSG00000115216
NRDC / O43847 / ENSSSCG00000003870 / ENSG00000078618
NRF1 / Q16656 / ENSSSCG00000016566 / ENSG00000106459
NRP1 / O14786 / ENSSSCG00000011102 / ENSG00000099250
NSDHL / Q15738 / ENSSSCG00000012758 / ENSG00000147383
NSFL1C / Q9UNZ2 / ENSSSCG00000007187 / ENSG00000088833
NSMCE2 / Q96MF7 / ENSSSCG00000037755 / ENSG00000156831
NSUN2 / Q08J23 / ENSSSCG00000017104 / ENSG00000037474
NSUN5 / Q96P11 / ENSSSCG00000007704 / ENSG00000130305
NT5C / Q8TCD5 / ENSSSCG00000028210 / ENSG00000125458
NT5C2 / P49902 / ENSSSCG00000010594 / ENSG00000076685
NT5C3A / Q9H0P0 / ENSSSCG00000022912 / ENSG00000122643
NT5DC1 / Q5TFE4 / ENSSSCG00000004435 / ENSG00000178425
NT5E / P21589 / ENSSSCG00000004291 / ENSG00000135318
NUB1 / Q9Y5A7 / ENSSSCG00000016438 / ENSG00000013374
NUBP2 / Q9Y5Y2 / ENSSSCG00000008014 / ENSG00000095906
NUBPL / Q8TB37 / ENSSSCG00000001968 / ENSG00000151413
NUCB1 / Q02818 / ENSSSCG00000003150 / ENSG00000104805
NUCKS1 / Q9H1E3 / ENSSSCG00000022398 / ENSG00000069275
NUDC / Q9Y266 / ENSSSCG00000003566 / ENSG00000090273
NUDCD1 / Q96RS6 / ENSSSCG00000006029 / ENSG00000120526
NUDCD2 / Q8WVJ2 / ENSSSCG00000032806 / ENSG00000170584
NUDCD3 / Q8IVD9 / ENSSSCG00000016747 / ENSG00000015676
NUDT14 / O95848 / ENSSSCG00000002557 / ENSG00000183828
NUDT16 / Q96DE0 / ENSSSCG00000032424 / ENSG00000198585
NUDT18 / Q6ZVK8 / ENSSSCG00000009615 / ENSG00000275074
NUDT2 / P50583 / ENSSSCG00000010981 / ENSG00000164978
NUDT21 / O43809 / ENSSSCG00000034521 / ENSG00000167005
NUDT3 / O95989 / ENSSSCG00000031593 / ENSG00000272325
NUMA1 / Q14980 / ENSSSCG00000014802 / ENSG00000137497
NUP107 / P57740 / ENSSSCG00000000486 / ENSG00000111581
NUP133 / Q8WUM0 / ENSSSCG00000010189 / ENSG00000069248
NUP153 / P49790 / ENSSSCG00000026034 / ENSG00000124789
NUP155 / O75694 / ENSSSCG00000016845 / ENSG00000113569
NUP160 / Q12769 / ENSSSCG00000013223 / ENSG00000030066
NUP188 / Q5SRE5 / ENSSSCG00000005667 / ENSG00000095319
NUP205 / Q92621 / ENSSSCG00000030386 / ENSG00000155561
NUP210 / Q8TEM1 / ENSSSCG00000011610 / ENSG00000132182
NUP214 / P35658 / ENSSSCG00000005711 / ENSG00000126883
NUP35 / Q8NFH5 / ENSSSCG00000028949 / ENSG00000163002
NUP37 / Q8NFH4 / ENSSSCG00000000860 / ENSG00000075188
NUP43 / Q8NFH3 / ENSSSCG00000004105 / ENSG00000120253
Homologs
Adipose.Tissue
Brain
Colon
Heart
Kidney
Liver
Lung
Pancreas
S.Intestine
Spleen
Organs
Species
Human
Pig
Bins
1
2
3
4
5

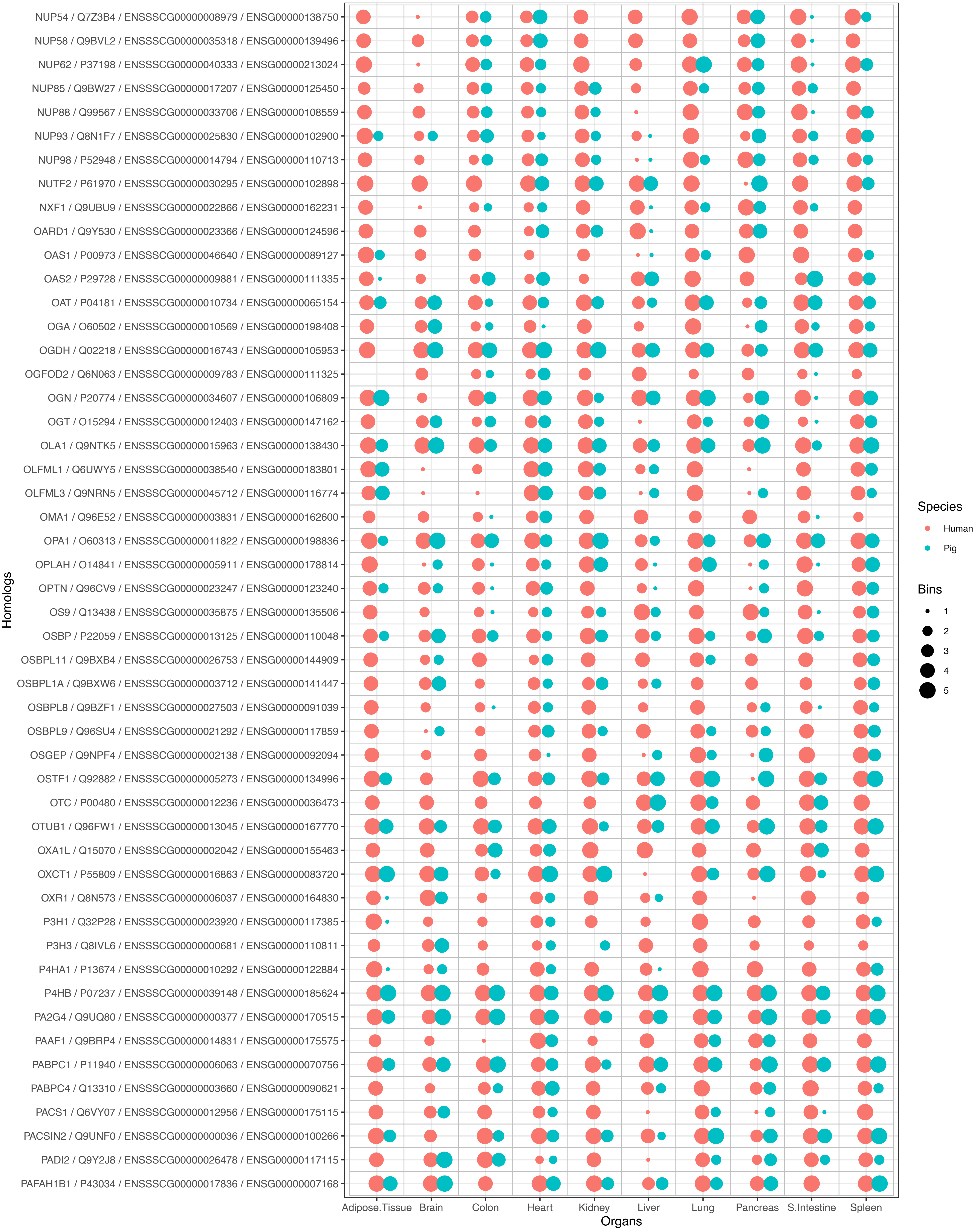
NUP54 / Q7Z3B4 / ENSSSCG00000008979 / ENSG00000138750
NUP58 / Q9BVL2 / ENSSSCG00000035318 / ENSG00000139496
NUP62 / P37198 / ENSSSCG00000040333 / ENSG00000213024
NUP85 / Q9BW27 / ENSSSCG00000017207 / ENSG00000125450
NUP88 / Q99567 / ENSSSCG00000033706 / ENSG00000108559
NUP93 / Q8N1F7 / ENSSSCG00000025830 / ENSG00000102900
NUP98 / P52948 / ENSSSCG00000014794 / ENSG00000110713
NUTF2 / P61970 / ENSSSCG00000030295 / ENSG00000102898
NXF1 / Q9UBU9 / ENSSSCG00000022866 / ENSG00000162231
OARD1 / Q9Y530 / ENSSSCG00000023366 / ENSG00000124596
OAS1 / P00973 / ENSSSCG00000046640 / ENSG00000089127
OAS2 / P29728 / ENSSSCG00000009881 / ENSG00000111335
OAT / P04181 / ENSSSCG00000010734 / ENSG00000065154
OGA / O60502 / ENSSSCG00000010569 / ENSG00000198408
OGDH / Q02218 / ENSSSCG00000016743 / ENSG00000105953
OGFOD2 / Q6N063 / ENSSSCG00000009783 / ENSG00000111325
OGN / P20774 / ENSSSCG00000034607 / ENSG00000106809
OGT / O15294 / ENSSSCG00000012403 / ENSG00000147162
OLA1 / Q9NTK5 / ENSSSCG00000015963 / ENSG00000138430
OLFML1 / Q6UWY5 / ENSSSCG00000038540 / ENSG00000183801
OLFML3 / Q9NRN5 / ENSSSCG00000045712 / ENSG00000116774
OMA1 / Q96E52 / ENSSSCG00000003831 / ENSG00000162600
OPA1 / O60313 / ENSSSCG00000011822 / ENSG00000198836
OPLAH / O14841 / ENSSSCG00000005911 / ENSG00000178814
OPTN / Q96CV9 / ENSSSCG00000023247 / ENSG00000123240
OS9 / Q13438 / ENSSSCG00000035875 / ENSG00000135506
OSBP / P22059 / ENSSSCG00000013125 / ENSG00000110048
OSBPL11 / Q9BXB4 / ENSSSCG00000026753 / ENSG00000144909
OSBPL1A / Q9BXW6 / ENSSSCG00000003712 / ENSG00000141447
OSBPL8 / Q9BZF1 / ENSSSCG00000027503 / ENSG00000091039
OSBPL9 / Q96SU4 / ENSSSCG00000021292 / ENSG00000117859
OSGEP / Q9NPF4 / ENSSSCG00000002138 / ENSG00000092094
OSTF1 / Q92882 / ENSSSCG00000005273 / ENSG00000134996
OTC / P00480 / ENSSSCG00000012236 / ENSG00000036473
OTUB1 / Q96FW1 / ENSSSCG00000013045 / ENSG00000167770
OXA1L / Q15070 / ENSSSCG00000002042 / ENSG00000155463
OXCT1 / P55809 / ENSSSCG00000016863 / ENSG00000083720
OXR1 / Q8N573 / ENSSSCG00000006037 / ENSG00000164830
P3H1 / Q32P28 / ENSSSCG00000023920 / ENSG00000117385
P3H3 / Q8IVL6 / ENSSSCG00000000681 / ENSG00000110811
P4HA1 / P13674 / ENSSSCG00000010292 / ENSG00000122884
P4HB / P07237 / ENSSSCG00000039148 / ENSG00000185624
PA2G4 / Q9UQ80 / ENSSSCG00000000377 / ENSG00000170515
PAAF1 / Q9BRP4 / ENSSSCG00000014831 / ENSG00000175575
PABPC1 / P11940 / ENSSSCG00000006063 / ENSG00000070756
PABPC4 / Q13310 / ENSSSCG00000003660 / ENSG00000090621
PACS1 / Q6VY07 / ENSSSCG00000012956 / ENSG00000175115
PACSIN2 / Q9UNF0 / ENSSSCG00000000036 / ENSG00000100266
PADI2 / Q9Y2J8 / ENSSSCG00000026478 / ENSG00000117115
PAFAH1B1 / P43034 / ENSSSCG00000017836 / ENSG00000007168
Homologs
Adipose.Tissue
Brain
Colon
Heart
Kidney
Liver
Lung
Pancreas
S.Intestine
Spleen
Organs
Species
Human
Pig
Bins
1
2
3
4
5

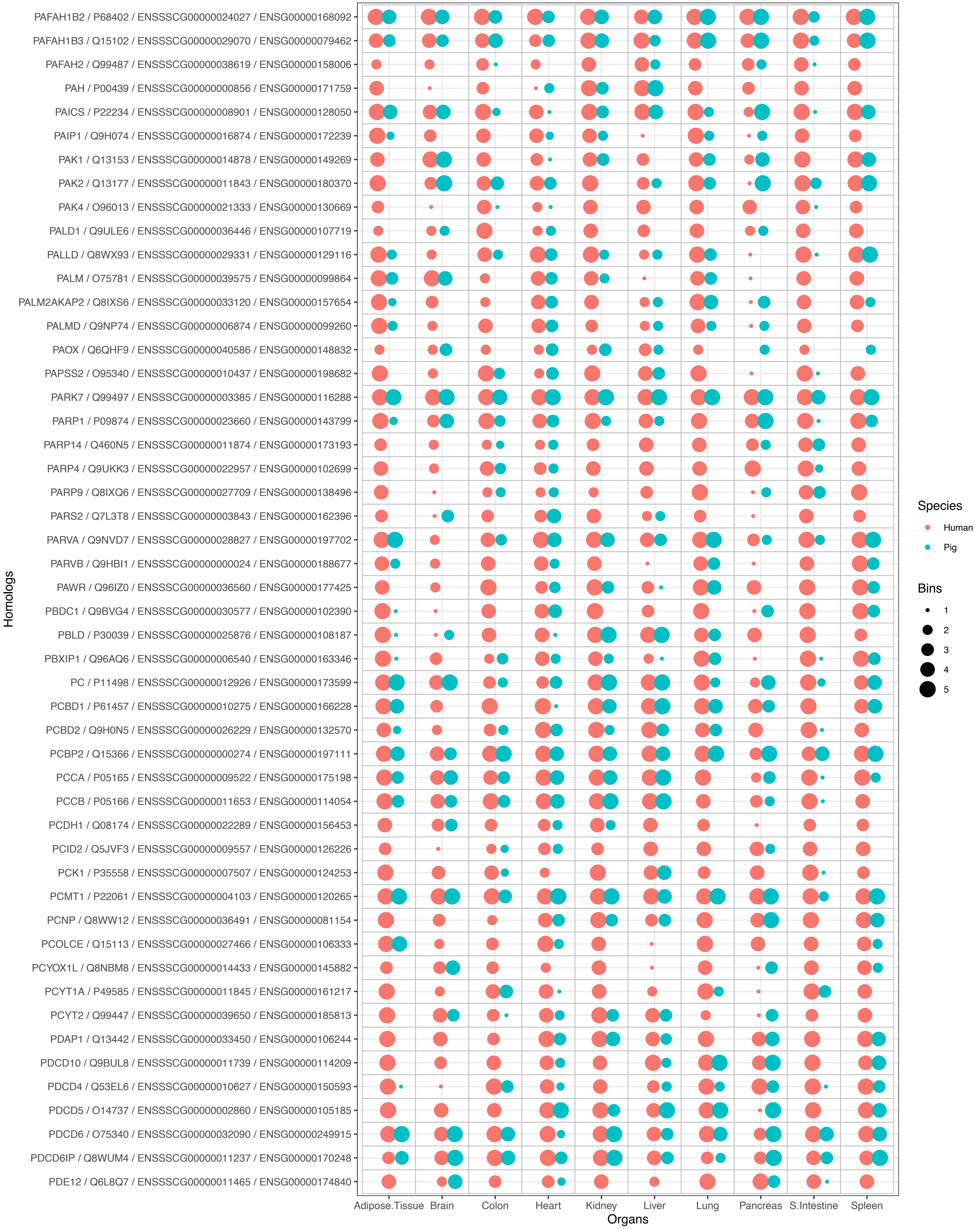

PAFAH1B2 / P68402 / ENSSSCG00000024027 / ENSG00000168092
PAFAH1B3 / Q15102 / ENSSSCG00000029070 / ENSG00000079462
PAFAH2 / Q99487 / ENSSSCG00000038619 / ENSG00000158006
PAH / P00439 / ENSSSCG00000000856 / ENSG00000171759
PAICS / P22234 / ENSSSCG00000008901 / ENSG00000128050
PAIP1 / Q9H074 / ENSSSCG00000016874 / ENSG00000172239
PAK1 / Q13153 / ENSSSCG00000014878 / ENSG00000149269
PAK2 / Q13177 / ENSSSCG00000011843 / ENSG00000180370
PAK4 / O96013 / ENSSSCG00000021333 / ENSG00000130669
PALD1 / Q9ULE6 / ENSSSCG00000036446 / ENSG00000107719
PALLD / Q8WX93 / ENSSSCG00000029331 / ENSG00000129116
PALM / O75781 / ENSSSCG00000039575 / ENSG00000099864
PALM2AKAP2 / Q8IXS6 / ENSSSCG00000033120 / ENSG00000157654
PALMD / Q9NP74 / ENSSSCG00000006874 / ENSG00000099260
PAOX / Q6QHF9 / ENSSSCG00000040586 / ENSG00000148832
PAPSS2 / O95340 / ENSSSCG00000010437 / ENSG00000198682
PARK7 / Q99497 / ENSSSCG00000003385 / ENSG00000116288
PARP1 / P09874 / ENSSSCG00000023660 / ENSG00000143799
PARP14 / Q460N5 / ENSSSCG00000011874 / ENSG00000173193
PARP4 / Q9UKK3 / ENSSSCG00000022957 / ENSG00000102699
PARP9 / Q8IXQ6 / ENSSSCG00000027709 / ENSG00000138496
PARS2 / Q7L3T8 / ENSSSCG00000003843 / ENSG00000162396
PARVA / Q9NVD7 / ENSSSCG00000028827 / ENSG00000197702
PARVB / Q9HBI1 / ENSSSCG00000000024 / ENSG00000188677
PAWR / Q96IZ0 / ENSSSCG00000036560 / ENSG00000177425
PBDC1 / Q9BVG4 / ENSSSCG00000030577 / ENSG00000102390
PBLD / P30039 / ENSSSCG00000025876 / ENSG00000108187
PBXIP1 / Q96AQ6 / ENSSSCG00000006540 / ENSG00000163346
PC / P11498 / ENSSSCG00000012926 / ENSG00000173599
PCBD1 / P61457 / ENSSSCG00000010275 / ENSG00000166228
PCBD2 / Q9H0N5 / ENSSSCG00000026229 / ENSG00000132570
PCBP2 / Q15366 / ENSSSCG00000000274 / ENSG00000197111
PCCA / P05165 / ENSSSCG00000009522 / ENSG00000175198
PCCB / P05166 / ENSSSCG00000011653 / ENSG00000114054
PCDH1 / Q08174 / ENSSSCG00000022289 / ENSG00000156453
PCID2 / Q5JVF3 / ENSSSCG00000009557 / ENSG00000126226
PCK1 / P35558 / ENSSSCG00000007507 / ENSG00000124253
PCMT1 / P22061 / ENSSSCG00000004103 / ENSG00000120265
PCNP / Q8WW12 / ENSSSCG00000036491 / ENSG00000081154
PCOLCE / Q15113 / ENSSSCG00000027466 / ENSG00000106333
PCYOX1L / Q8NBM8 / ENSSSCG00000014433 / ENSG00000145882
PCYT1A / P49585 / ENSSSCG00000011845 / ENSG00000161217
PCYT2 / Q99447 / ENSSSCG00000039650 / ENSG00000185813
PDAP1 / Q13442 / ENSSSCG00000033450 / ENSG00000106244
PDCD10 / Q9BUL8 / ENSSSCG00000011739 / ENSG00000114209
PDCD4 / Q53EL6 / ENSSSCG00000010627 / ENSG00000150593
PDCD5 / O14737 / ENSSSCG00000002860 / ENSG00000105185
PDCD6 / O75340 / ENSSSCG00000032090 / ENSG00000249915
PDCD6IP / Q8WUM4 / ENSSSCG00000011237 / ENSG00000170248
PDE12 / Q6L8Q7 / ENSSSCG00000011465 / ENSG00000174840
Homologs
Adipose.Tissue
Brain
Colon
Heart
Kidney
Liver
Lung
Pancreas
S.Intestine
Spleen
Organs
Species
Human
Pig
Bins
1
2
3
4
5

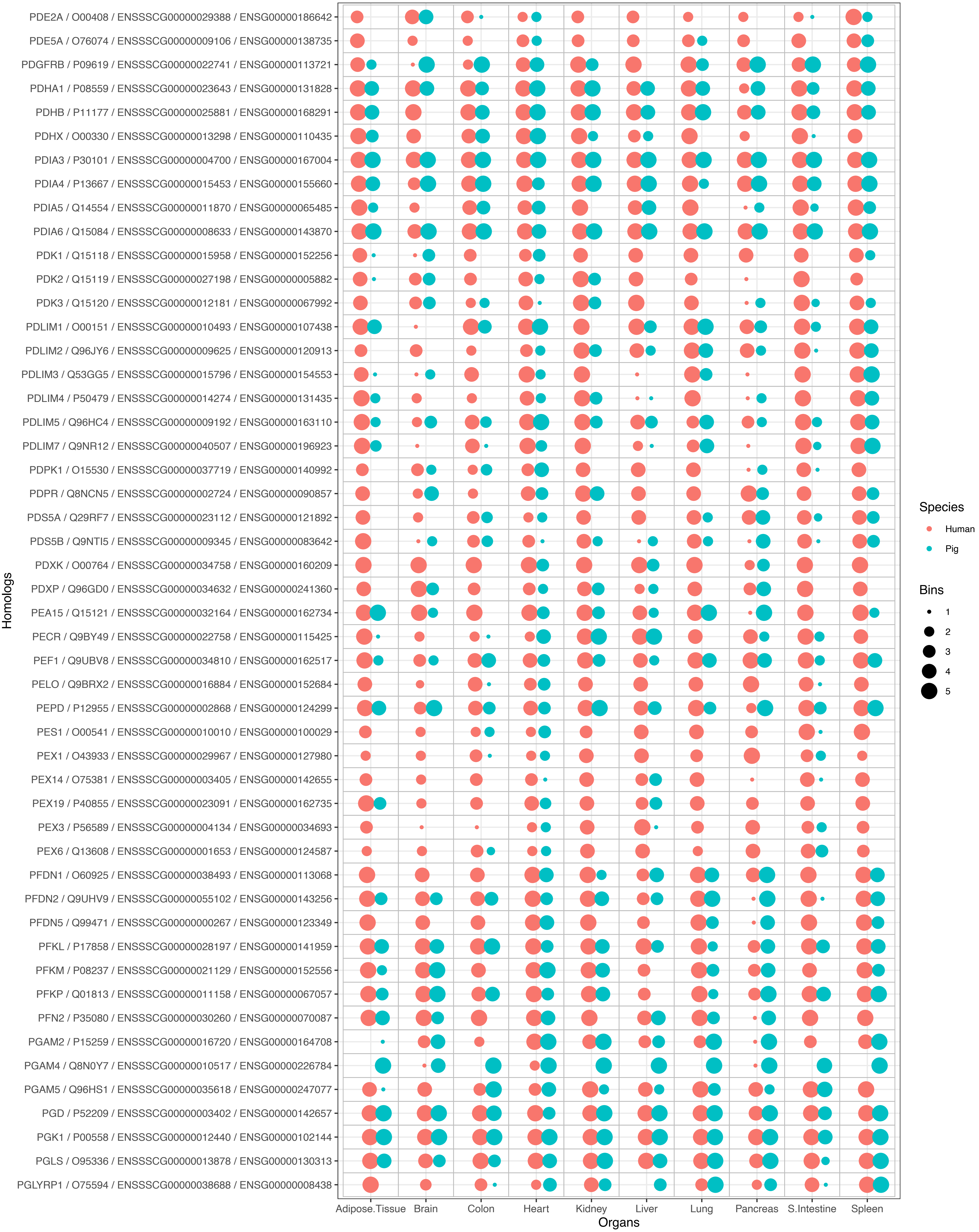

PDE2A / O00408 / ENSSSCG00000029388 / ENSG00000186642
PDE5A / O76074 / ENSSSCG00000009106 / ENSG00000138735
PDGFRB / P09619 / ENSSSCG00000022741 / ENSG00000113721
PDHA1 / P08559 / ENSSSCG00000023643 / ENSG00000131828
PDHB / P11177 / ENSSSCG00000025881 / ENSG00000168291
PDHX / O00330 / ENSSSCG00000013298 / ENSG00000110435
PDIA3 / P30101 / ENSSSCG00000004700 / ENSG00000167004
PDIA4 / P13667 / ENSSSCG00000015453 / ENSG00000155660
PDIA5 / Q14554 / ENSSSCG00000011870 / ENSG00000065485
PDIA6 / Q15084 / ENSSSCG00000008633 / ENSG00000143870
PDK1 / Q15118 / ENSSSCG00000015958 / ENSG00000152256
PDK2 / Q15119 / ENSSSCG00000027198 / ENSG00000005882
PDK3 / Q15120 / ENSSSCG00000012181 / ENSG00000067992
PDLIM1 / O00151 / ENSSSCG00000010493 / ENSG00000107438
PDLIM2 / Q96JY6 / ENSSSCG00000009625 / ENSG00000120913
PDLIM3 / Q53GG5 / ENSSSCG00000015796 / ENSG00000154553
PDLIM4 / P50479 / ENSSSCG00000014274 / ENSG00000131435
PDLIM5 / Q96HC4 / ENSSSCG00000009192 / ENSG00000163110
PDLIM7 / Q9NR12 / ENSSSCG00000040507 / ENSG00000196923
PDPK1 / O15530 / ENSSSCG00000037719 / ENSG00000140992
PDPR / Q8NCN5 / ENSSSCG00000002724 / ENSG00000090857
PDS5A / Q29RF7 / ENSSSCG00000023112 / ENSG00000121892
PDS5B / Q9NTI5 / ENSSSCG00000009345 / ENSG00000083642
PDXK / O00764 / ENSSSCG00000034758 / ENSG00000160209
PDXP / Q96GD0 / ENSSSCG00000034632 / ENSG00000241360
PEA15 / Q15121 / ENSSSCG00000032164 / ENSG00000162734
PECR / Q9BY49 / ENSSSCG00000022758 / ENSG00000115425
PEF1 / Q9UBV8 / ENSSSCG00000034810 / ENSG00000162517
PELO / Q9BRX2 / ENSSSCG00000016884 / ENSG00000152684
PEPD / P12955 / ENSSSCG00000002868 / ENSG00000124299
PES1 / O00541 / ENSSSCG00000010010 / ENSG00000100029
PEX1 / O43933 / ENSSSCG00000029967 / ENSG00000127980
PEX14 / O75381 / ENSSSCG00000003405 / ENSG00000142655
PEX19 / P40855 / ENSSSCG00000023091 / ENSG00000162735
PEX3 / P56589 / ENSSSCG00000004134 / ENSG00000034693
PEX6 / Q13608 / ENSSSCG00000001653 / ENSG00000124587
PFDN1 / O60925 / ENSSSCG00000038493 / ENSG00000113068
PFDN2 / Q9UHV9 / ENSSSCG00000055102 / ENSG00000143256
PFDN5 / Q99471 / ENSSSCG00000000267 / ENSG00000123349
PFKL / P17858 / ENSSSCG00000028197 / ENSG00000141959
PFKM / P08237 / ENSSSCG00000021129 / ENSG00000152556
PFKP / Q01813 / ENSSSCG00000011158 / ENSG00000067057
PFN2 / P35080 / ENSSSCG00000030260 / ENSG00000070087
PGAM2 / P15259 / ENSSSCG00000016720 / ENSG00000164708
PGAM4 / Q8N0Y7 / ENSSSCG00000010517 / ENSG00000226784
PGAM5 / Q96HS1 / ENSSSCG00000035618 / ENSG00000247077
PGD / P52209 / ENSSSCG00000003402 / ENSG00000142657
PGK1 / P00558 / ENSSSCG00000012440 / ENSG00000102144
PGLS / O95336 / ENSSSCG00000013878 / ENSG00000130313
PGLYRP1 / O75594 / ENSSSCG00000038688 / ENSG00000008438
Homologs
Adipose.Tissue
Brain
Colon
Heart
Kidney
Liver
Lung
Pancreas
S.Intestine
Spleen
Organs
Species
Human
Pig
Bins
1
2
3
4
5

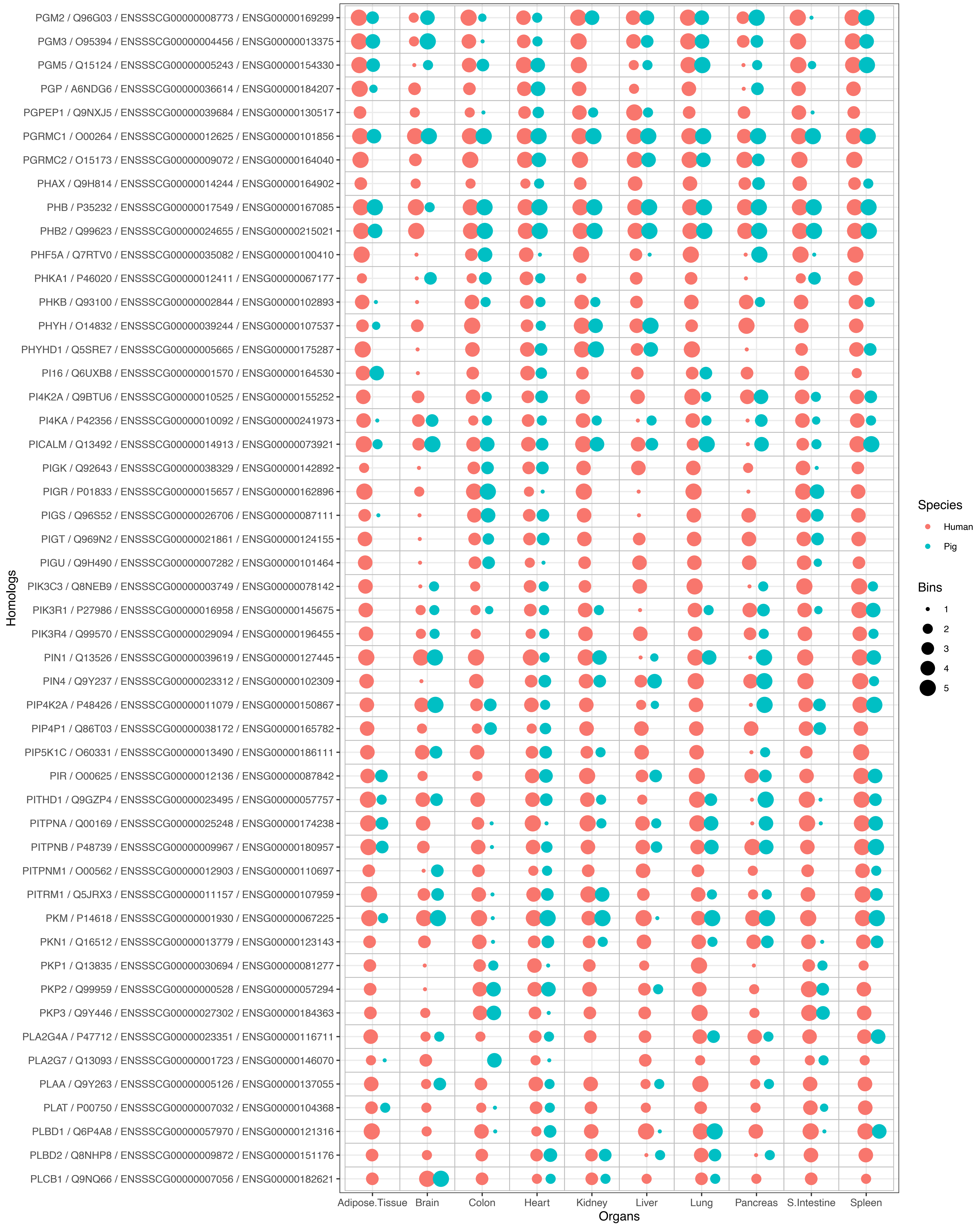

PGM2 / Q96G03 / ENSSSCG00000008773 / ENSG00000169299
PGM3 / O95394 / ENSSSCG00000004456 / ENSG00000013375
PGM5 / Q15124 / ENSSSCG00000005243 / ENSG00000154330
PGP / A6NDG6 / ENSSSCG00000036614 / ENSG00000184207
PGPEP1 / Q9NXJ5 / ENSSSCG00000039684 / ENSG00000130517
PGRMC1 / O00264 / ENSSSCG00000012625 / ENSG00000101856
PGRMC2 / O15173 / ENSSSCG00000009072 / ENSG00000164040
PHAX / Q9H814 / ENSSSCG00000014244 / ENSG00000164902
PHB / P35232 / ENSSSCG00000017549 / ENSG00000167085
PHB2 / Q99623 / ENSSSCG00000024655 / ENSG00000215021
PHF5A / Q7RTV0 / ENSSSCG00000035082 / ENSG00000100410
PHKA1 / P46020 / ENSSSCG00000012411 / ENSG00000067177
PHKB / Q93100 / ENSSSCG00000002844 / ENSG00000102893
PHYH / O14832 / ENSSSCG00000039244 / ENSG00000107537
PHYHD1 / Q5SRE7 / ENSSSCG00000005665 / ENSG00000175287
PI16 / Q6UXB8 / ENSSSCG00000001570 / ENSG00000164530
PI4K2A / Q9BTU6 / ENSSSCG00000010525 / ENSG00000155252
PI4KA / P42356 / ENSSSCG00000010092 / ENSG00000241973
PICALM / Q13492 / ENSSSCG00000014913 / ENSG00000073921
PIGK / Q92643 / ENSSSCG00000038329 / ENSG00000142892
PIGR / P01833 / ENSSSCG00000015657 / ENSG00000162896
PIGS / Q96S52 / ENSSSCG00000026706 / ENSG00000087111
PIGT / Q969N2 / ENSSSCG00000021861 / ENSG00000124155
PIGU / Q9H490 / ENSSSCG00000007282 / ENSG00000101464
PIK3C3 / Q8NEB9 / ENSSSCG00000003749 / ENSG00000078142
PIK3R1 / P27986 / ENSSSCG00000016958 / ENSG00000145675
PIK3R4 / Q99570 / ENSSSCG00000029094 / ENSG00000196455
PIN1 / Q13526 / ENSSSCG00000039619 / ENSG00000127445
PIN4 / Q9Y237 / ENSSSCG00000023312 / ENSG00000102309
PIP4K2A / P48426 / ENSSSCG00000011079 / ENSG00000150867
PIP4P1 / Q86T03 / ENSSSCG00000038172 / ENSG00000165782
PIP5K1C / O60331 / ENSSSCG00000013490 / ENSG00000186111
PIR / O00625 / ENSSSCG00000012136 / ENSG00000087842
PITHD1 / Q9GZP4 / ENSSSCG00000023495 / ENSG00000057757
PITPNA / Q00169 / ENSSSCG00000025248 / ENSG00000174238
PITPNB / P48739 / ENSSSCG00000009967 / ENSG00000180957
PITPNM1 / O00562 / ENSSSCG00000012903 / ENSG00000110697
PITRM1 / Q5JRX3 / ENSSSCG00000011157 / ENSG00000107959
PKM / P14618 / ENSSSCG00000001930 / ENSG00000067225
PKN1 / Q16512 / ENSSSCG00000013779 / ENSG00000123143
PKP1 / Q13835 / ENSSSCG00000030694 / ENSG00000081277
PKP2 / Q99959 / ENSSSCG00000000528 / ENSG00000057294
PKP3 / Q9Y446 / ENSSSCG00000027302 / ENSG00000184363
PLA2G4A / P47712 / ENSSSCG00000023351 / ENSG00000116711
PLA2G7 / Q13093 / ENSSSCG00000001723 / ENSG00000146070
PLAA / Q9Y263 / ENSSSCG00000005126 / ENSG00000137055
PLAT / P00750 / ENSSSCG00000007032 / ENSG00000104368
PLBD1 / Q6P4A8 / ENSSSCG00000057970 / ENSG00000121316
PLBD2 / Q8NHP8 / ENSSSCG00000009872 / ENSG00000151176
PLCB1 / Q9NQ66 / ENSSSCG00000007056 / ENSG00000182621
Homologs
Adipose.Tissue
Brain
Colon
Heart
Kidney
Liver
Lung
Pancreas
S.Intestine
Spleen
Organs
Species
Human
Pig
Bins
1
2
3
4
5

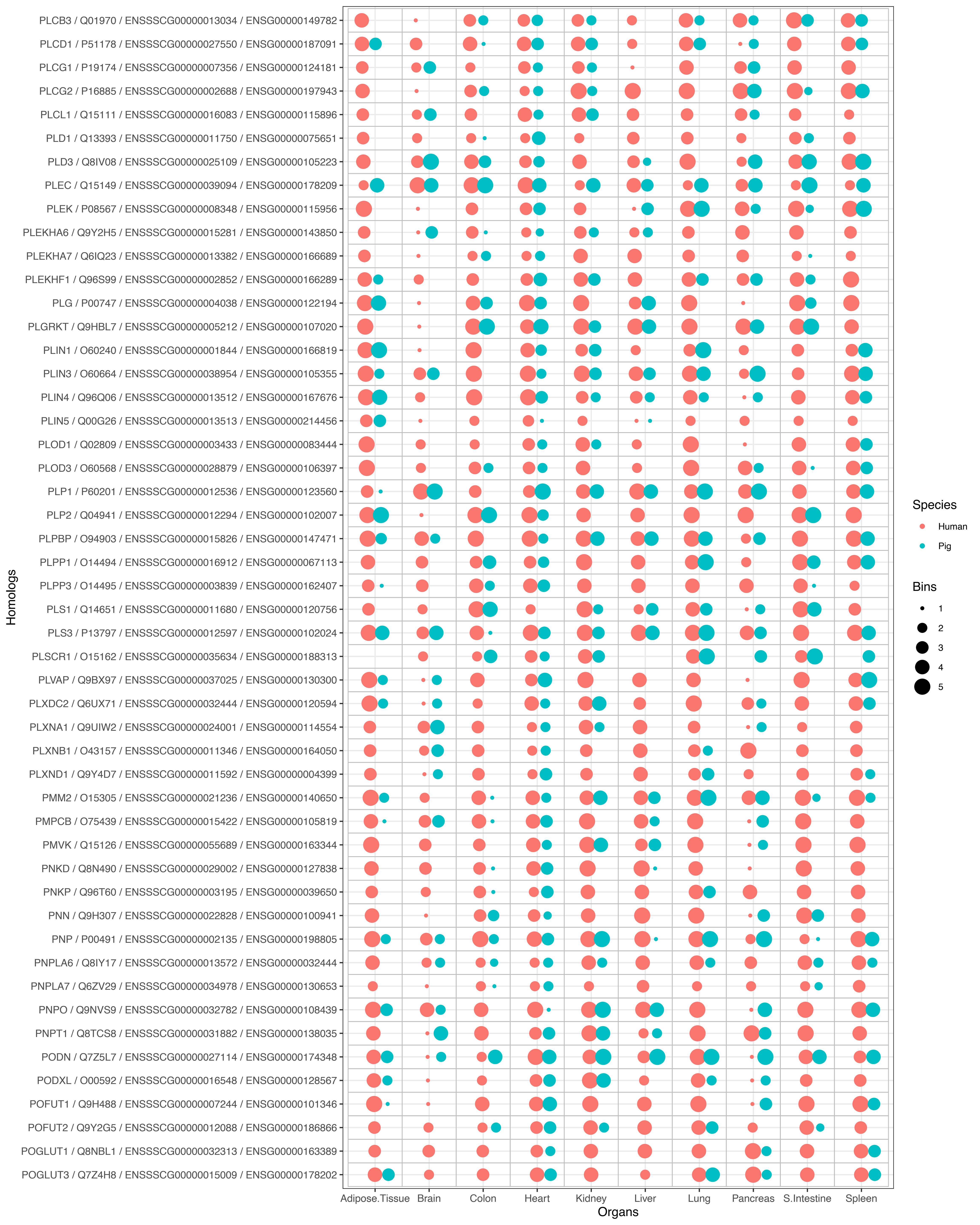

PLCB3 / Q01970 / ENSSSCG00000013034 / ENSG00000149782
PLCD1 / P51178 / ENSSSCG00000027550 / ENSG00000187091
PLCG1 / P19174 / ENSSSCG00000007356 / ENSG00000124181
PLCG2 / P16885 / ENSSSCG00000002688 / ENSG00000197943
PLCL1 / Q15111 / ENSSSCG00000016083 / ENSG00000115896
PLD1 / Q13393 / ENSSSCG00000011750 / ENSG00000075651
PLD3 / Q8IV08 / ENSSSCG00000025109 / ENSG00000105223
PLEC / Q15149 / ENSSSCG00000039094 / ENSG00000178209
PLEK / P08567 / ENSSSCG00000008348 / ENSG00000115956
PLEKHA6 / Q9Y2H5 / ENSSSCG00000015281 / ENSG00000143850
PLEKHA7 / Q6IQ23 / ENSSSCG00000013382 / ENSG00000166689
PLEKHF1 / Q96S99 / ENSSSCG00000002852 / ENSG00000166289
PLG / P00747 / ENSSSCG00000004038 / ENSG00000122194
PLGRKT / Q9HBL7 / ENSSSCG00000005212 / ENSG00000107020
PLIN1 / O60240 / ENSSSCG00000001844 / ENSG00000166819
PLIN3 / O60664 / ENSSSCG00000038954 / ENSG00000105355
PLIN4 / Q96Q06 / ENSSSCG00000013512 / ENSG00000167676
PLIN5 / Q00G26 / ENSSSCG00000013513 / ENSG00000214456
PLOD1 / Q02809 / ENSSSCG00000003433 / ENSG00000083444
PLOD3 / O60568 / ENSSSCG00000028879 / ENSG00000106397
PLP1 / P60201 / ENSSSCG00000012536 / ENSG00000123560
PLP2 / Q04941 / ENSSSCG00000012294 / ENSG00000102007
PLPBP / O94903 / ENSSSCG00000015826 / ENSG00000147471
PLPP1 / O14494 / ENSSSCG00000016912 / ENSG00000067113
PLPP3 / O14495 / ENSSSCG00000003839 / ENSG00000162407
PLS1 / Q14651 / ENSSSCG00000011680 / ENSG00000120756
PLS3 / P13797 / ENSSSCG00000012597 / ENSG00000102024
PLSCR1 / O15162 / ENSSSCG00000035634 / ENSG00000188313
PLVAP / Q9BX97 / ENSSSCG00000037025 / ENSG00000130300
PLXDC2 / Q6UX71 / ENSSSCG00000032444 / ENSG00000120594
PLXNA1 / Q9UIW2 / ENSSSCG00000024001 / ENSG00000114554
PLXNB1 / O43157 / ENSSSCG00000011346 / ENSG00000164050
PLXND1 / Q9Y4D7 / ENSSSCG00000011592 / ENSG00000004399
PMM2 / O15305 / ENSSSCG00000021236 / ENSG00000140650
PMPCB / O75439 / ENSSSCG00000015422 / ENSG00000105819
PMVK / Q15126 / ENSSSCG00000055689 / ENSG00000163344
PNKD / Q8N490 / ENSSSCG00000029002 / ENSG00000127838
PNKP / Q96T60 / ENSSSCG00000003195 / ENSG00000039650
PNN / Q9H307 / ENSSSCG00000022828 / ENSG00000100941
PNP / P00491 / ENSSSCG00000002135 / ENSG00000198805
PNPLA6 / Q8IY17 / ENSSSCG00000013572 / ENSG00000032444
PNPLA7 / Q6ZV29 / ENSSSCG00000034978 / ENSG00000130653
PNPO / Q9NVS9 / ENSSSCG00000032782 / ENSG00000108439
PNPT1 / Q8TCS8 / ENSSSCG00000031882 / ENSG00000138035
PODN / Q7Z5L7 / ENSSSCG00000027114 / ENSG00000174348
PODXL / O00592 / ENSSSCG00000016548 / ENSG00000128567
POFUT1 / Q9H488 / ENSSSCG00000007244 / ENSG00000101346
POFUT2 / Q9Y2G5 / ENSSSCG00000012088 / ENSG00000186866
POGLUT1 / Q8NBL1 / ENSSSCG00000032313 / ENSG00000163389
POGLUT3 / Q7Z4H8 / ENSSSCG00000015009 / ENSG00000178202
Homologs
Adipose.Tissue
Brain
Colon
Heart
Kidney
Liver
Lung
Pancreas
S.Intestine
Spleen
Organs
Species
Human
Pig
Bins
1
2
3
4
5

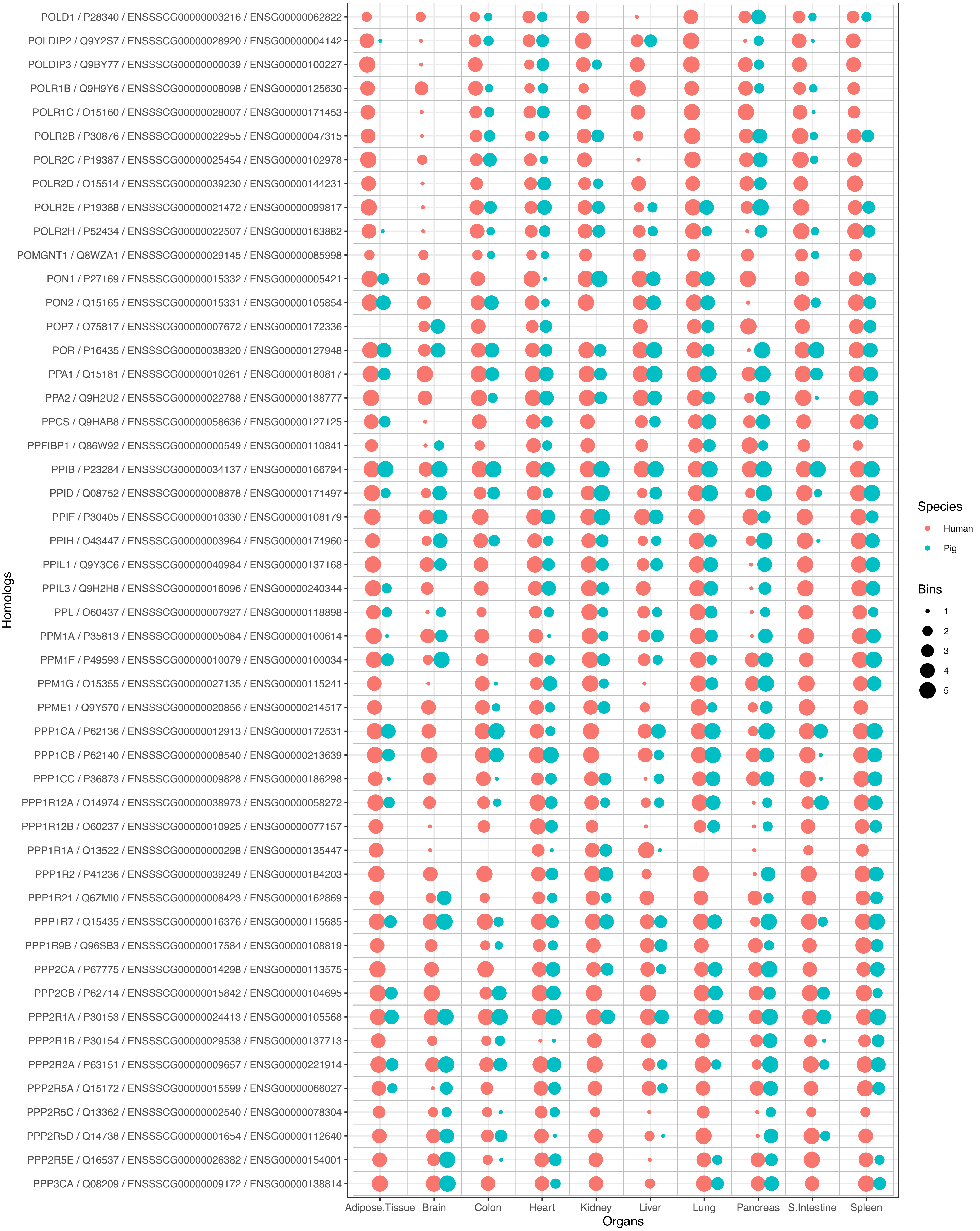
POLD1 / P28340 / ENSSSCG00000003216 / ENSG00000062822
POLDIP2 / Q9Y2S7 / ENSSSCG00000028920 / ENSG00000004142
POLDIP3 / Q9BY77 / ENSSSCG00000000039 / ENSG00000100227
POLR1B / Q9H9Y6 / ENSSSCG00000008098 / ENSG00000125630
POLR1C / O15160 / ENSSSCG00000028007 / ENSG00000171453
POLR2B / P30876 / ENSSSCG00000022955 / ENSG00000047315
POLR2C / P19387 / ENSSSCG00000025454 / ENSG00000102978
POLR2D / O15514 / ENSSSCG00000039230 / ENSG00000144231
POLR2E / P19388 / ENSSSCG00000021472 / ENSG00000099817
POLR2H / P52434 / ENSSSCG00000022507 / ENSG00000163882
POMGNT1 / Q8WZA1 / ENSSSCG00000029145 / ENSG00000085998
PON1 / P27169 / ENSSSCG00000015332 / ENSG00000005421
PON2 / Q15165 / ENSSSCG00000015331 / ENSG00000105854
POP7 / O75817 / ENSSSCG00000007672 / ENSG00000172336
POR / P16435 / ENSSSCG00000038320 / ENSG00000127948
PPA1 / Q15181 / ENSSSCG00000010261 / ENSG00000180817
PPA2 / Q9H2U2 / ENSSSCG00000022788 / ENSG00000138777
PPCS / Q9HAB8 / ENSSSCG00000058636 / ENSG00000127125
PPFIBP1 / Q86W92 / ENSSSCG00000000549 / ENSG00000110841
PPIB / P23284 / ENSSSCG00000034137 / ENSG00000166794
PPID / Q08752 / ENSSSCG00000008878 / ENSG00000171497
PPIF / P30405 / ENSSSCG00000010330 / ENSG00000108179
PPIH / O43447 / ENSSSCG00000003964 / ENSG00000171960
PPIL1 / Q9Y3C6 / ENSSSCG00000040984 / ENSG00000137168
PPIL3 / Q9H2H8 / ENSSSCG00000016096 / ENSG00000240344
PPL / O60437 / ENSSSCG00000007927 / ENSG00000118898
PPM1A / P35813 / ENSSSCG00000005084 / ENSG00000100614
PPM1F / P49593 / ENSSSCG00000010079 / ENSG00000100034
PPM1G / O15355 / ENSSSCG00000027135 / ENSG00000115241
PPME1 / Q9Y570 / ENSSSCG00000020856 / ENSG00000214517
PPP1CA / P62136 / ENSSSCG00000012913 / ENSG00000172531
PPP1CB / P62140 / ENSSSCG00000008540 / ENSG00000213639
PPP1CC / P36873 / ENSSSCG00000009828 / ENSG00000186298
PPP1R12A / O14974 / ENSSSCG00000038973 / ENSG00000058272
PPP1R12B / O60237 / ENSSSCG00000010925 / ENSG00000077157
PPP1R1A / Q13522 / ENSSSCG00000000298 / ENSG00000135447
PPP1R2 / P41236 / ENSSSCG00000039249 / ENSG00000184203
PPP1R21 / Q6ZMI0 / ENSSSCG00000008423 / ENSG00000162869
PPP1R7 / Q15435 / ENSSSCG00000016376 / ENSG00000115685
PPP1R9B / Q96SB3 / ENSSSCG00000017584 / ENSG00000108819
PPP2CA / P67775 / ENSSSCG00000014298 / ENSG00000113575
PPP2CB / P62714 / ENSSSCG00000015842 / ENSG00000104695
PPP2R1A / P30153 / ENSSSCG00000024413 / ENSG00000105568
PPP2R1B / P30154 / ENSSSCG00000029538 / ENSG00000137713
PPP2R2A / P63151 / ENSSSCG00000009657 / ENSG00000221914
PPP2R5A / Q15172 / ENSSSCG00000015599 / ENSG00000066027
PPP2R5C / Q13362 / ENSSSCG00000002540 / ENSG00000078304
PPP2R5D / Q14738 / ENSSSCG00000001654 / ENSG00000112640
PPP2R5E / Q16537 / ENSSSCG00000026382 / ENSG00000154001
PPP3CA / Q08209 / ENSSSCG00000009172 / ENSG00000138814
Homologs
Adipose.Tissue
Brain
Colon
Heart
Kidney
Liver
Lung
Pancreas
S.Intestine
Spleen
Organs
Species
Human
Pig
Bins
1
2
3
4
5

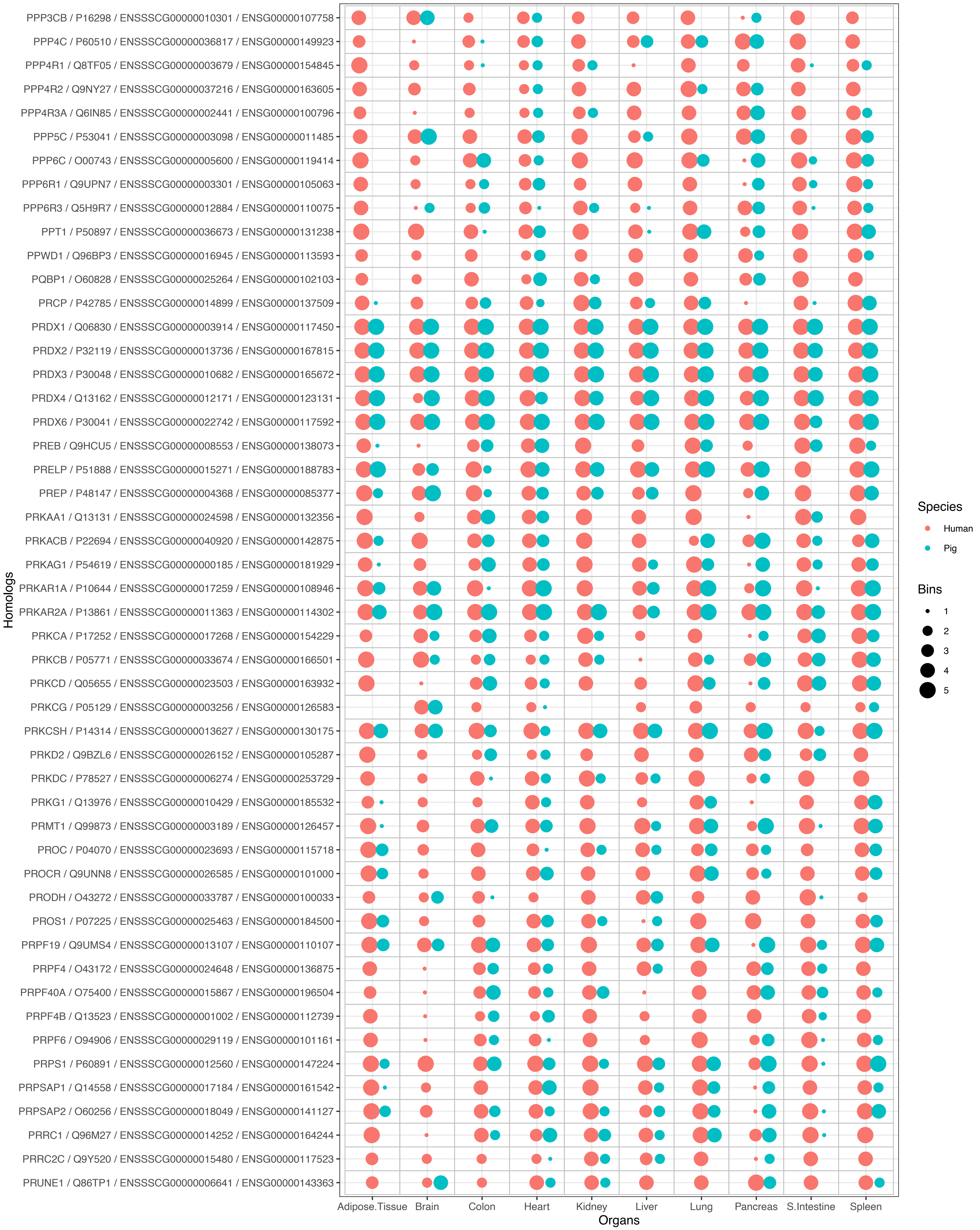
PPP3CB / P16298 / ENSSSCG00000010301 / ENSG00000107758
PPP4C / P60510 / ENSSSCG00000036817 / ENSG00000149923
PPP4R1 / Q8TF05 / ENSSSCG00000003679 / ENSG00000154845
PPP4R2 / Q9NY27 / ENSSSCG00000037216 / ENSG00000163605
PPP4R3A / Q6IN85 / ENSSSCG00000002441 / ENSG00000100796
PPP5C / P53041 / ENSSSCG00000003098 / ENSG00000011485
PPP6C / O00743 / ENSSSCG00000005600 / ENSG00000119414
PPP6R1 / Q9UPN7 / ENSSSCG00000003301 / ENSG00000105063
PPP6R3 / Q5H9R7 / ENSSSCG00000012884 / ENSG00000110075
PPT1 / P50897 / ENSSSCG00000036673 / ENSG00000131238
PPWD1 / Q96BP3 / ENSSSCG00000016945 / ENSG00000113593
PQBP1 / O60828 / ENSSSCG00000025264 / ENSG00000102103
PRCP / P42785 / ENSSSCG00000014899 / ENSG00000137509
PRDX1 / Q06830 / ENSSSCG00000003914 / ENSG00000117450
PRDX2 / P32119 / ENSSSCG00000013736 / ENSG00000167815
PRDX3 / P30048 / ENSSSCG00000010682 / ENSG00000165672
PRDX4 / Q13162 / ENSSSCG00000012171 / ENSG00000123131
PRDX6 / P30041 / ENSSSCG00000022742 / ENSG00000117592
PREB / Q9HCU5 / ENSSSCG00000008553 / ENSG00000138073
PRELP / P51888 / ENSSSCG00000015271 / ENSG00000188783
PREP / P48147 / ENSSSCG00000004368 / ENSG00000085377
PRKAA1 / Q13131 / ENSSSCG00000024598 / ENSG00000132356
PRKACB / P22694 / ENSSSCG00000040920 / ENSG00000142875
PRKAG1 / P54619 / ENSSSCG00000000185 / ENSG00000181929
PRKAR1A / P10644 / ENSSSCG00000017259 / ENSG00000108946
PRKAR2A / P13861 / ENSSSCG00000011363 / ENSG00000114302
PRKCA / P17252 / ENSSSCG00000017268 / ENSG00000154229
PRKCB / P05771 / ENSSSCG00000033674 / ENSG00000166501
PRKCD / Q05655 / ENSSSCG00000023503 / ENSG00000163932
PRKCG / P05129 / ENSSSCG00000003256 / ENSG00000126583
PRKCSH / P14314 / ENSSSCG00000013627 / ENSG00000130175
PRKD2 / Q9BZL6 / ENSSSCG00000026152 / ENSG00000105287
PRKDC / P78527 / ENSSSCG00000006274 / ENSG00000253729
PRKG1 / Q13976 / ENSSSCG00000010429 / ENSG00000185532
PRMT1 / Q99873 / ENSSSCG00000003189 / ENSG00000126457
PROC / P04070 / ENSSSCG00000023693 / ENSG00000115718
PROCR / Q9UNN8 / ENSSSCG00000026585 / ENSG00000101000
PRODH / O43272 / ENSSSCG00000033787 / ENSG00000100033
PROS1 / P07225 / ENSSSCG00000025463 / ENSG00000184500
PRPF19 / Q9UMS4 / ENSSSCG00000013107 / ENSG00000110107
PRPF4 / O43172 / ENSSSCG00000024648 / ENSG00000136875
PRPF40A / O75400 / ENSSSCG00000015867 / ENSG00000196504
PRPF4B / Q13523 / ENSSSCG00000001002 / ENSG00000112739
PRPF6 / O94906 / ENSSSCG00000029119 / ENSG00000101161
PRPS1 / P60891 / ENSSSCG00000012560 / ENSG00000147224
PRPSAP1 / Q14558 / ENSSSCG00000017184 / ENSG00000161542
PRPSAP2 / O60256 / ENSSSCG00000018049 / ENSG00000141127
PRRC1 / Q96M27 / ENSSSCG00000014252 / ENSG00000164244
PRRC2C / Q9Y520 / ENSSSCG00000015480 / ENSG00000117523
PRUNE1 / Q86TP1 / ENSSSCG00000006641 / ENSG00000143363
Homologs
Adipose.Tissue
Brain
Colon
Heart
Kidney
Liver
Lung
Pancreas
S.Intestine
Spleen
Organs
Species
Human
Pig
Bins
1
2
3
4
5

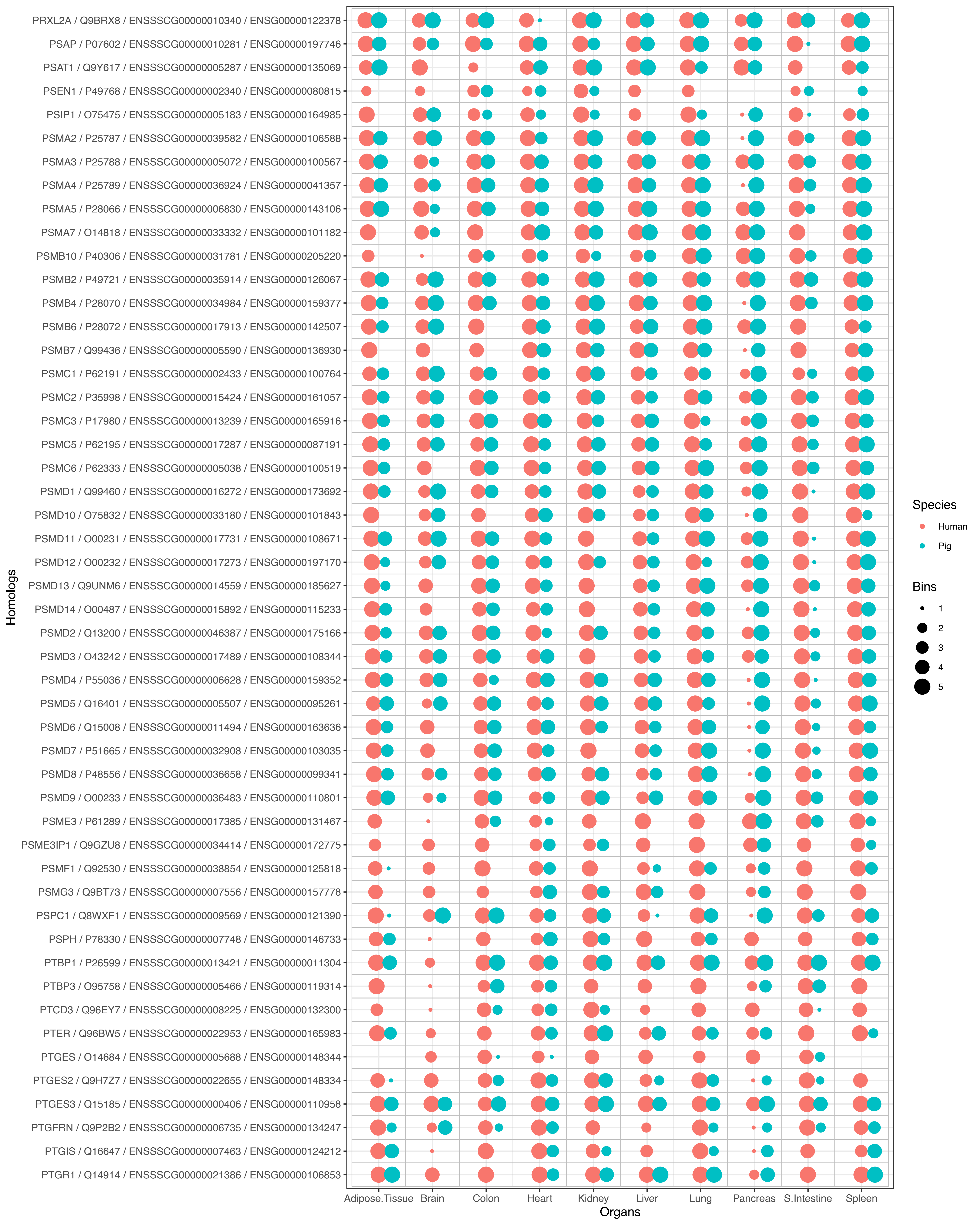

PRXL2A / Q9BRX8 / ENSSSCG00000010340 / ENSG00000122378
PSAP / P07602 / ENSSSCG00000010281 / ENSG00000197746
PSAT1 / Q9Y617 / ENSSSCG00000005287 / ENSG00000135069
PSEN1 / P49768 / ENSSSCG00000002340 / ENSG00000080815
PSIP1 / O75475 / ENSSSCG00000005183 / ENSG00000164985
PSMA2 / P25787 / ENSSSCG00000039582 / ENSG00000106588
PSMA3 / P25788 / ENSSSCG00000005072 / ENSG00000100567
PSMA4 / P25789 / ENSSSCG00000036924 / ENSG00000041357
PSMA5 / P28066 / ENSSSCG00000006830 / ENSG00000143106
PSMA7 / O14818 / ENSSSCG00000033332 / ENSG00000101182
PSMB10 / P40306 / ENSSSCG00000031781 / ENSG00000205220
PSMB2 / P49721 / ENSSSCG00000035914 / ENSG00000126067
PSMB4 / P28070 / ENSSSCG00000034984 / ENSG00000159377
PSMB6 / P28072 / ENSSSCG00000017913 / ENSG00000142507
PSMB7 / Q99436 / ENSSSCG00000005590 / ENSG00000136930
PSMC1 / P62191 / ENSSSCG00000002433 / ENSG00000100764
PSMC2 / P35998 / ENSSSCG00000015424 / ENSG00000161057
PSMC3 / P17980 / ENSSSCG00000013239 / ENSG00000165916
PSMC5 / P62195 / ENSSSCG00000017287 / ENSG00000087191
PSMC6 / P62333 / ENSSSCG00000005038 / ENSG00000100519
PSMD1 / Q99460 / ENSSSCG00000016272 / ENSG00000173692
PSMD10 / O75832 / ENSSSCG00000033180 / ENSG00000101843
PSMD11 / O00231 / ENSSSCG00000017731 / ENSG00000108671
PSMD12 / O00232 / ENSSSCG00000017273 / ENSG00000197170
PSMD13 / Q9UNM6 / ENSSSCG00000014559 / ENSG00000185627
PSMD14 / O00487 / ENSSSCG00000015892 / ENSG00000115233
PSMD2 / Q13200 / ENSSSCG00000046387 / ENSG00000175166
PSMD3 / O43242 / ENSSSCG00000017489 / ENSG00000108344
PSMD4 / P55036 / ENSSSCG00000006628 / ENSG00000159352
PSMD5 / Q16401 / ENSSSCG00000005507 / ENSG00000095261
PSMD6 / Q15008 / ENSSSCG00000011494 / ENSG00000163636
PSMD7 / P51665 / ENSSSCG00000032908 / ENSG00000103035
PSMD8 / P48556 / ENSSSCG00000036658 / ENSG00000099341
PSMD9 / O00233 / ENSSSCG00000036483 / ENSG00000110801
PSME3 / P61289 / ENSSSCG00000017385 / ENSG00000131467
PSME3IP1 / Q9GZU8 / ENSSSCG00000034414 / ENSG00000172775
PSMF1 / Q92530 / ENSSSCG00000038854 / ENSG00000125818
PSMG3 / Q9BT73 / ENSSSCG00000007556 / ENSG00000157778
PSPC1 / Q8WXF1 / ENSSSCG00000009569 / ENSG00000121390
PSPH / P78330 / ENSSSCG00000007748 / ENSG00000146733
PTBP1 / P26599 / ENSSSCG00000013421 / ENSG00000011304
PTBP3 / O95758 / ENSSSCG00000005466 / ENSG00000119314
PTCD3 / Q96EY7 / ENSSSCG00000008225 / ENSG00000132300
PTER / Q96BW5 / ENSSSCG00000022953 / ENSG00000165983
PTGES / O14684 / ENSSSCG00000005688 / ENSG00000148344
PTGES2 / Q9H7Z7 / ENSSSCG00000022655 / ENSG00000148334
PTGES3 / Q15185 / ENSSSCG00000000406 / ENSG00000110958
PTGFRN / Q9P2B2 / ENSSSCG00000006735 / ENSG00000134247
PTGIS / Q16647 / ENSSSCG00000007463 / ENSG00000124212
PTGR1 / Q14914 / ENSSSCG00000021386 / ENSG00000106853
Homologs
Adipose.Tissue
Brain
Colon
Heart
Kidney
Liver
Lung
Pancreas
S.Intestine
Spleen
Organs
Species
Human
Pig
Bins
1
2
3
4
5

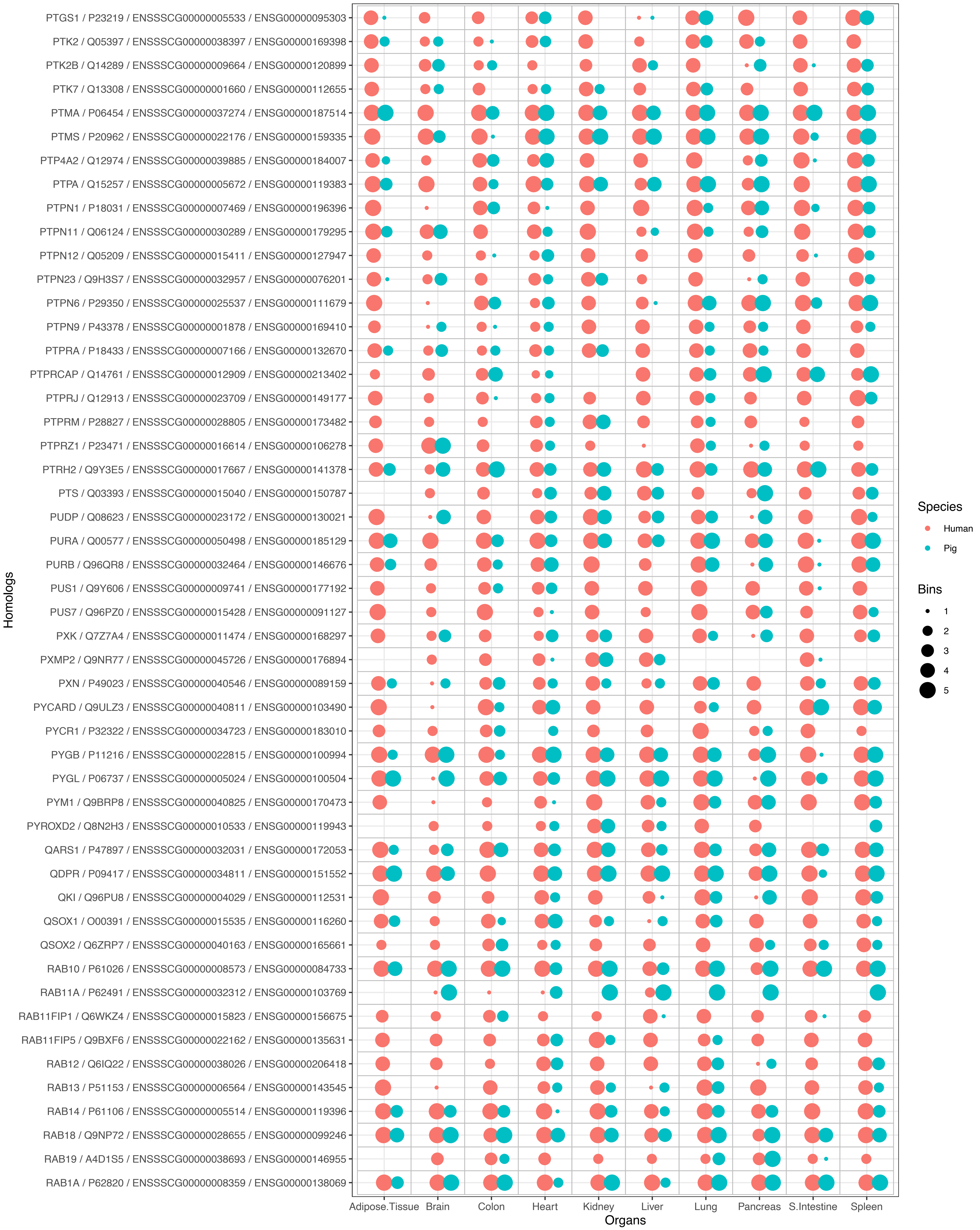

PTGS1 / P23219 / ENSSSCG00000005533 / ENSG00000095303
PTK2 / Q05397 / ENSSSCG00000038397 / ENSG00000169398
PTK2B / Q14289 / ENSSSCG00000009664 / ENSG00000120899
PTK7 / Q13308 / ENSSSCG00000001660 / ENSG00000112655
PTMA / P06454 / ENSSSCG00000037274 / ENSG00000187514
PTMS / P20962 / ENSSSCG00000022176 / ENSG00000159335
PTP4A2 / Q12974 / ENSSSCG00000039885 / ENSG00000184007
PTPA / Q15257 / ENSSSCG00000005672 / ENSG00000119383
PTPN1 / P18031 / ENSSSCG00000007469 / ENSG00000196396
PTPN11 / Q06124 / ENSSSCG00000030289 / ENSG00000179295
PTPN12 / Q05209 / ENSSSCG00000015411 / ENSG00000127947
PTPN23 / Q9H3S7 / ENSSSCG00000032957 / ENSG00000076201
PTPN6 / P29350 / ENSSSCG00000025537 / ENSG00000111679
PTPN9 / P43378 / ENSSSCG00000001878 / ENSG00000169410
PTPRA / P18433 / ENSSSCG00000007166 / ENSG00000132670
PTPRCAP / Q14761 / ENSSSCG00000012909 / ENSG00000213402
PTPRJ / Q12913 / ENSSSCG00000023709 / ENSG00000149177
PTPRM / P28827 / ENSSSCG00000028805 / ENSG00000173482
PTPRZ1 / P23471 / ENSSSCG00000016614 / ENSG00000106278
PTRH2 / Q9Y3E5 / ENSSSCG00000017667 / ENSG00000141378
PTS / Q03393 / ENSSSCG00000015040 / ENSG00000150787
PUDP / Q08623 / ENSSSCG00000023172 / ENSG00000130021
PURA / Q00577 / ENSSSCG00000050498 / ENSG00000185129
PURB / Q96QR8 / ENSSSCG00000032464 / ENSG00000146676
PUS1 / Q9Y606 / ENSSSCG00000009741 / ENSG00000177192
PUS7 / Q96PZ0 / ENSSSCG00000015428 / ENSG00000091127
PXK / Q7Z7A4 / ENSSSCG00000011474 / ENSG00000168297
PXMP2 / Q9NR77 / ENSSSCG00000045726 / ENSG00000176894
PXN / P49023 / ENSSSCG00000040546 / ENSG00000089159
PYCARD / Q9ULZ3 / ENSSSCG00000040811 / ENSG00000103490
PYCR1 / P32322 / ENSSSCG00000034723 / ENSG00000183010
PYGB / P11216 / ENSSSCG00000022815 / ENSG00000100994
PYGL / P06737 / ENSSSCG00000005024 / ENSG00000100504
PYM1 / Q9BRP8 / ENSSSCG00000040825 / ENSG00000170473
PYROXD2 / Q8N2H3 / ENSSSCG00000010533 / ENSG00000119943
QARS1 / P47897 / ENSSSCG00000032031 / ENSG00000172053
QDPR / P09417 / ENSSSCG00000034811 / ENSG00000151552
QKI / Q96PU8 / ENSSSCG00000004029 / ENSG00000112531
QSOX1 / O00391 / ENSSSCG00000015535 / ENSG00000116260
QSOX2 / Q6ZRP7 / ENSSSCG00000040163 / ENSG00000165661
RAB10 / P61026 / ENSSSCG00000008573 / ENSG00000084733
RAB11A / P62491 / ENSSSCG00000032312 / ENSG00000103769
RAB11FIP1 / Q6WKZ4 / ENSSSCG00000015823 / ENSG00000156675
RAB11FIP5 / Q9BXF6 / ENSSSCG00000022162 / ENSG00000135631
RAB12 / Q6IQ22 / ENSSSCG00000038026 / ENSG00000206418
RAB13 / P51153 / ENSSSCG00000006564 / ENSG00000143545
RAB14 / P61106 / ENSSSCG00000005514 / ENSG00000119396
RAB18 / Q9NP72 / ENSSSCG00000028655 / ENSG00000099246
RAB19 / A4D1S5 / ENSSSCG00000038693 / ENSG00000146955
RAB1A / P62820 / ENSSSCG00000008359 / ENSG00000138069
Homologs
Adipose.Tissue
Brain
Colon
Heart
Kidney
Liver
Lung
Pancreas
S.Intestine
Spleen
Organs
Species
Human
Pig
Bins
1
2
3
4
5

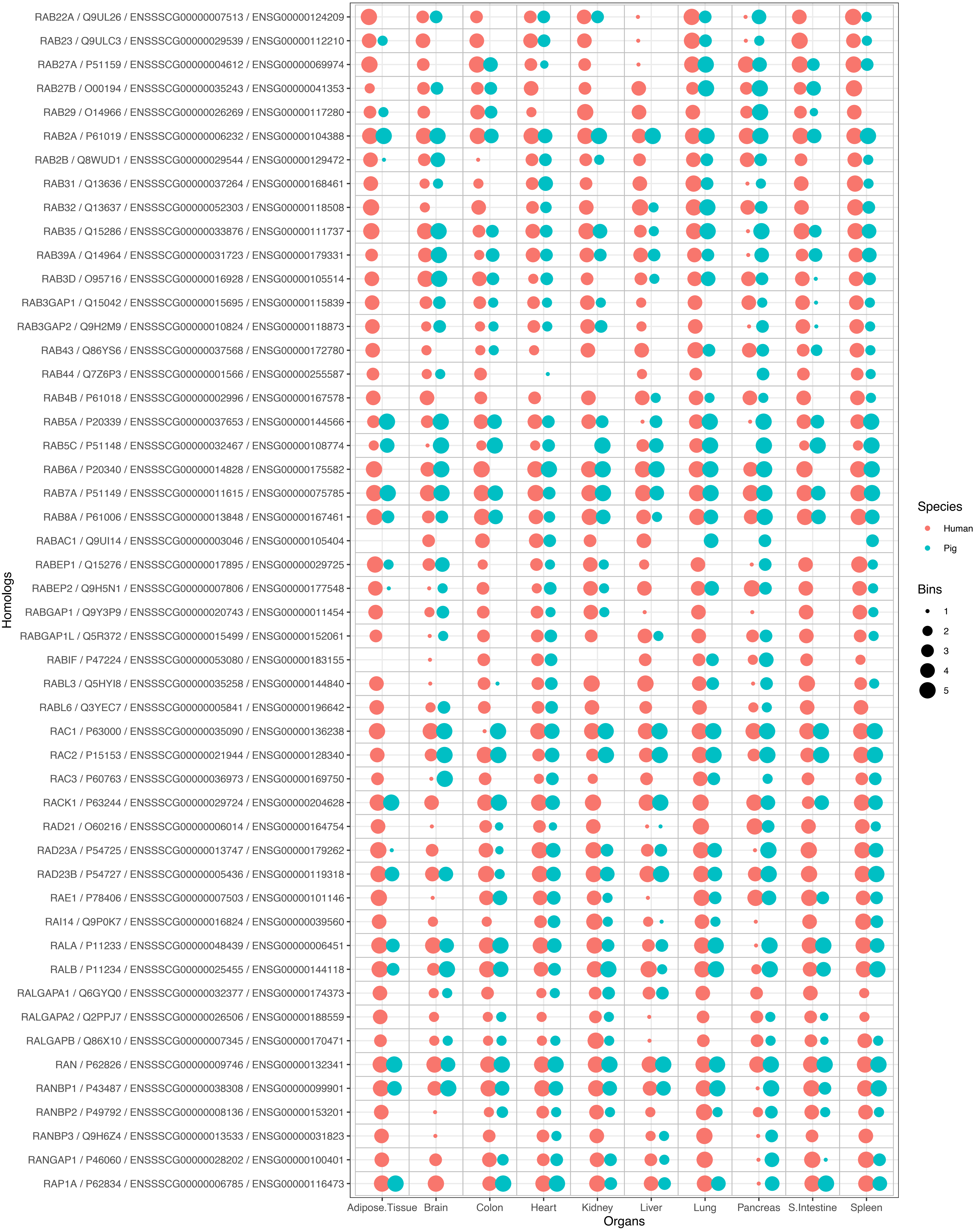
RAB22A / Q9UL26 / ENSSSCG00000007513 / ENSG00000124209
RAB23 / Q9ULC3 / ENSSSCG00000029539 / ENSG00000112210
RAB27A / P51159 / ENSSSCG00000004612 / ENSG00000069974
RAB27B / O00194 / ENSSSCG00000035243 / ENSG00000041353
RAB29 / O14966 / ENSSSCG00000026269 / ENSG00000117280
RAB2A / P61019 / ENSSSCG00000006232 / ENSG00000104388
RAB2B / Q8WUD1 / ENSSSCG00000029544 / ENSG00000129472
RAB31 / Q13636 / ENSSSCG00000037264 / ENSG00000168461
RAB32 / Q13637 / ENSSSCG00000052303 / ENSG00000118508
RAB35 / Q15286 / ENSSSCG00000033876 / ENSG00000111737
RAB39A / Q14964 / ENSSSCG00000031723 / ENSG00000179331
RAB3D / O95716 / ENSSSCG00000016928 / ENSG00000105514
RAB3GAP1 / Q15042 / ENSSSCG00000015695 / ENSG00000115839
RAB3GAP2 / Q9H2M9 / ENSSSCG00000010824 / ENSG00000118873
RAB43 / Q86YS6 / ENSSSCG00000037568 / ENSG00000172780
RAB44 / Q7Z6P3 / ENSSSCG00000001566 / ENSG00000255587
RAB4B / P61018 / ENSSSCG00000002996 / ENSG00000167578
RAB5A / P20339 / ENSSSCG00000037653 / ENSG00000144566
RAB5C / P51148 / ENSSSCG00000032467 / ENSG00000108774
RAB6A / P20340 / ENSSSCG00000014828 / ENSG00000175582
RAB7A / P51149 / ENSSSCG00000011615 / ENSG00000075785
RAB8A / P61006 / ENSSSCG00000013848 / ENSG00000167461
RABAC1 / Q9UI14 / ENSSSCG00000003046 / ENSG00000105404
RABEP1 / Q15276 / ENSSSCG00000017895 / ENSG00000029725
RABEP2 / Q9H5N1 / ENSSSCG00000007806 / ENSG00000177548
RABGAP1 / Q9Y3P9 / ENSSSCG00000020743 / ENSG00000011454
RABGAP1L / Q5R372 / ENSSSCG00000015499 / ENSG00000152061
RABIF / P47224 / ENSSSCG00000053080 / ENSG00000183155
RABL3 / Q5HYI8 / ENSSSCG00000035258 / ENSG00000144840
RABL6 / Q3YEC7 / ENSSSCG00000005841 / ENSG00000196642
RAC1 / P63000 / ENSSSCG00000035090 / ENSG00000136238
RAC2 / P15153 / ENSSSCG00000021944 / ENSG00000128340
RAC3 / P60763 / ENSSSCG00000036973 / ENSG00000169750
RACK1 / P63244 / ENSSSCG00000029724 / ENSG00000204628
RAD21 / O60216 / ENSSSCG00000006014 / ENSG00000164754
RAD23A / P54725 / ENSSSCG00000013747 / ENSG00000179262
RAD23B / P54727 / ENSSSCG00000005436 / ENSG00000119318
RAE1 / P78406 / ENSSSCG00000007503 / ENSG00000101146
RAI14 / Q9P0K7 / ENSSSCG00000016824 / ENSG00000039560
RALA / P11233 / ENSSSCG00000048439 / ENSG00000006451
RALB / P11234 / ENSSSCG00000025455 / ENSG00000144118
RALGAPA1 / Q6GYQ0 / ENSSSCG00000032377 / ENSG00000174373
RALGAPA2 / Q2PPJ7 / ENSSSCG00000026506 / ENSG00000188559
RALGAPB / Q86X10 / ENSSSCG00000007345 / ENSG00000170471
RAN / P62826 / ENSSSCG00000009746 / ENSG00000132341
RANBP1 / P43487 / ENSSSCG00000038308 / ENSG00000099901
RANBP2 / P49792 / ENSSSCG00000008136 / ENSG00000153201
RANBP3 / Q9H6Z4 / ENSSSCG00000013533 / ENSG00000031823
RANGAP1 / P46060 / ENSSSCG00000028202 / ENSG00000100401
RAP1A / P62834 / ENSSSCG00000006785 / ENSG00000116473
Homologs
Adipose.Tissue
Brain
Colon
Heart
Kidney
Liver
Lung
Pancreas
S.Intestine
Spleen
Organs
Species
Human
Pig
Bins
1
2
3
4
5

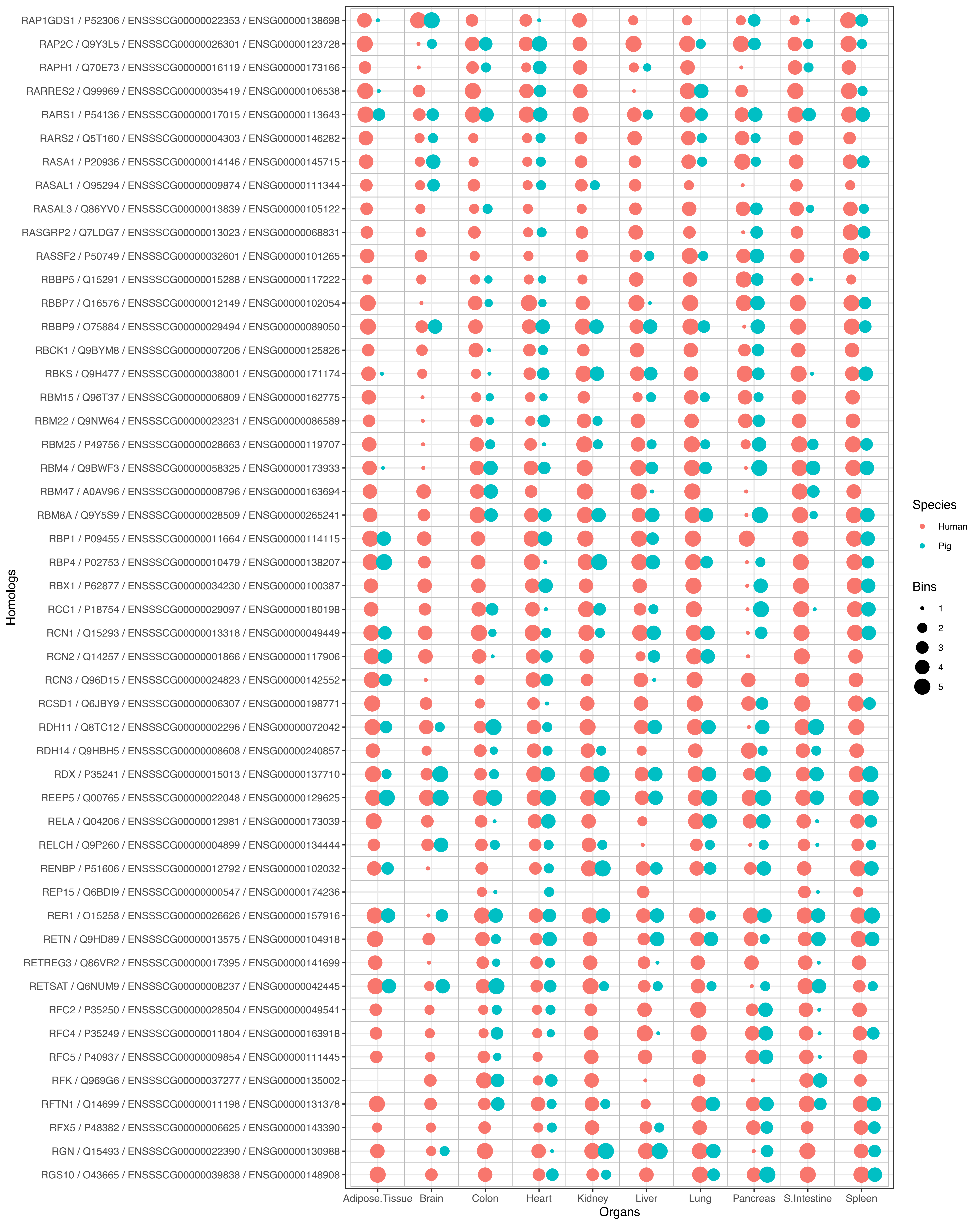
RAP1GDS1 / P52306 / ENSSSCG00000022353 / ENSG00000138698
RAP2C / Q9Y3L5 / ENSSSCG00000026301 / ENSG00000123728
RAPH1 / Q70E73 / ENSSSCG00000016119 / ENSG00000173166
RARRES2 / Q99969 / ENSSSCG00000035419 / ENSG00000106538
RARS1 / P54136 / ENSSSCG00000017015 / ENSG00000113643
RARS2 / Q5T160 / ENSSSCG00000004303 / ENSG00000146282
RASA1 / P20936 / ENSSSCG00000014146 / ENSG00000145715
RASAL1 / O95294 / ENSSSCG00000009874 / ENSG00000111344
RASAL3 / Q86YV0 / ENSSSCG00000013839 / ENSG00000105122
RASGRP2 / Q7LDG7 / ENSSSCG00000013023 / ENSG00000068831
RASSF2 / P50749 / ENSSSCG00000032601 / ENSG00000101265
RBBP5 / Q15291 / ENSSSCG00000015288 / ENSG00000117222
RBBP7 / Q16576 / ENSSSCG00000012149 / ENSG00000102054
RBBP9 / O75884 / ENSSSCG00000029494 / ENSG00000089050
RBCK1 / Q9BYM8 / ENSSSCG00000007206 / ENSG00000125826
RBKS / Q9H477 / ENSSSCG00000038001 / ENSG00000171174
RBM15 / Q96T37 / ENSSSCG00000006809 / ENSG00000162775
RBM22 / Q9NW64 / ENSSSCG00000023231 / ENSG00000086589
RBM25 / P49756 / ENSSSCG00000028663 / ENSG00000119707
RBM4 / Q9BWF3 / ENSSSCG00000058325 / ENSG00000173933
RBM47 / A0AV96 / ENSSSCG00000008796 / ENSG00000163694
RBM8A / Q9Y5S9 / ENSSSCG00000028509 / ENSG00000265241
RBP1 / P09455 / ENSSSCG00000011664 / ENSG00000114115
RBP4 / P02753 / ENSSSCG00000010479 / ENSG00000138207
RBX1 / P62877 / ENSSSCG00000034230 / ENSG00000100387
RCC1 / P18754 / ENSSSCG00000029097 / ENSG00000180198
RCN1 / Q15293 / ENSSSCG00000013318 / ENSG00000049449
RCN2 / Q14257 / ENSSSCG00000001866 / ENSG00000117906
RCN3 / Q96D15 / ENSSSCG00000024823 / ENSG00000142552
RCSD1 / Q6JBY9 / ENSSSCG00000006307 / ENSG00000198771
RDH11 / Q8TC12 / ENSSSCG00000002296 / ENSG00000072042
RDH14 / Q9HBH5 / ENSSSCG00000008608 / ENSG00000240857
RDX / P35241 / ENSSSCG00000015013 / ENSG00000137710
REEP5 / Q00765 / ENSSSCG00000022048 / ENSG00000129625
RELA / Q04206 / ENSSSCG00000012981 / ENSG00000173039
RELCH / Q9P260 / ENSSSCG00000004899 / ENSG00000134444
RENBP / P51606 / ENSSSCG00000012792 / ENSG00000102032
REP15 / Q6BDI9 / ENSSSCG00000000547 / ENSG00000174236
RER1 / O15258 / ENSSSCG00000026626 / ENSG00000157916
RETN / Q9HD89 / ENSSSCG00000013575 / ENSG00000104918
RETREG3 / Q86VR2 / ENSSSCG00000017395 / ENSG00000141699
RETSAT / Q6NUM9 / ENSSSCG00000008237 / ENSG00000042445
RFC2 / P35250 / ENSSSCG00000028504 / ENSG00000049541
RFC4 / P35249 / ENSSSCG00000011804 / ENSG00000163918
RFC5 / P40937 / ENSSSCG00000009854 / ENSG00000111445
RFK / Q969G6 / ENSSSCG00000037277 / ENSG00000135002
RFTN1 / Q14699 / ENSSSCG00000011198 / ENSG00000131378
RFX5 / P48382 / ENSSSCG00000006625 / ENSG00000143390
RGN / Q15493 / ENSSSCG00000022390 / ENSG00000130988
RGS10 / O43665 / ENSSSCG00000039838 / ENSG00000148908
Homologs
Adipose.Tissue
Brain
Colon
Heart
Kidney
Liver
Lung
Pancreas
S.Intestine
Spleen
Organs
Species
Human
Pig
Bins
1
2
3
4
5

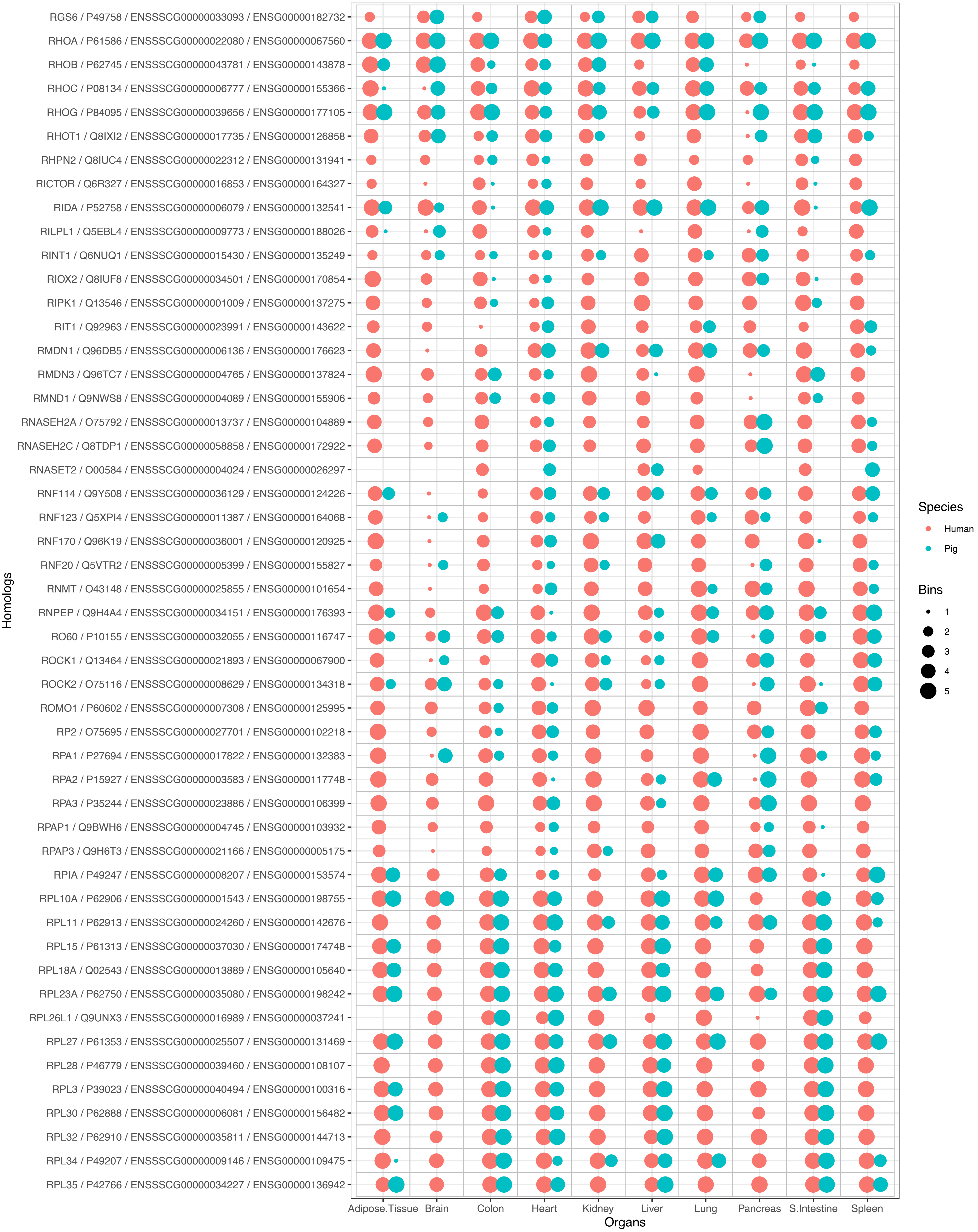
RGS6 / P49758 / ENSSSCG00000033093 / ENSG00000182732
RHOA / P61586 / ENSSSCG00000022080 / ENSG00000067560
RHOB / P62745 / ENSSSCG00000043781 / ENSG00000143878
RHOC / P08134 / ENSSSCG00000006777 / ENSG00000155366
RHOG / P84095 / ENSSSCG00000039656 / ENSG00000177105
RHOT1 / Q8IXI2 / ENSSSCG00000017735 / ENSG00000126858
RHPN2 / Q8IUC4 / ENSSSCG00000022312 / ENSG00000131941
RICTOR / Q6R327 / ENSSSCG00000016853 / ENSG00000164327
RIDA / P52758 / ENSSSCG00000006079 / ENSG00000132541
RILPL1 / Q5EBL4 / ENSSSCG00000009773 / ENSG00000188026
RINT1 / Q6NUQ1 / ENSSSCG00000015430 / ENSG00000135249
RIOX2 / Q8IUF8 / ENSSSCG00000034501 / ENSG00000170854
RIPK1 / Q13546 / ENSSSCG00000001009 / ENSG00000137275
RIT1 / Q92963 / ENSSSCG00000023991 / ENSG00000143622
RMDN1 / Q96DB5 / ENSSSCG00000006136 / ENSG00000176623
RMDN3 / Q96TC7 / ENSSSCG00000004765 / ENSG00000137824
RMND1 / Q9NWS8 / ENSSSCG00000004089 / ENSG00000155906
RNASEH2A / O75792 / ENSSSCG00000013737 / ENSG00000104889
RNASEH2C / Q8TDP1 / ENSSSCG00000058858 / ENSG00000172922
RNASET2 / O00584 / ENSSSCG00000004024 / ENSG00000026297
RNF114 / Q9Y508 / ENSSSCG00000036129 / ENSG00000124226
RNF123 / Q5XPI4 / ENSSSCG00000011387 / ENSG00000164068
RNF170 / Q96K19 / ENSSSCG00000036001 / ENSG00000120925
RNF20 / Q5VTR2 / ENSSSCG00000005399 / ENSG00000155827
RNMT / O43148 / ENSSSCG00000025855 / ENSG00000101654
RNPEP / Q9H4A4 / ENSSSCG00000034151 / ENSG00000176393
RO60 / P10155 / ENSSSCG00000032055 / ENSG00000116747
ROCK1 / Q13464 / ENSSSCG00000021893 / ENSG00000067900
ROCK2 / O75116 / ENSSSCG00000008629 / ENSG00000134318
ROMO1 / P60602 / ENSSSCG00000007308 / ENSG00000125995
RP2 / O75695 / ENSSSCG00000027701 / ENSG00000102218
RPA1 / P27694 / ENSSSCG00000017822 / ENSG00000132383
RPA2 / P15927 / ENSSSCG00000003583 / ENSG00000117748
RPA3 / P35244 / ENSSSCG00000023886 / ENSG00000106399
RPAP1 / Q9BWH6 / ENSSSCG00000004745 / ENSG00000103932
RPAP3 / Q9H6T3 / ENSSSCG00000021166 / ENSG00000005175
RPIA / P49247 / ENSSSCG00000008207 / ENSG00000153574
RPL10A / P62906 / ENSSSCG00000001543 / ENSG00000198755
RPL11 / P62913 / ENSSSCG00000024260 / ENSG00000142676
RPL15 / P61313 / ENSSSCG00000037030 / ENSG00000174748
RPL18A / Q02543 / ENSSSCG00000013889 / ENSG00000105640
RPL23A / P62750 / ENSSSCG00000035080 / ENSG00000198242
RPL26L1 / Q9UNX3 / ENSSSCG00000016989 / ENSG00000037241
RPL27 / P61353 / ENSSSCG00000025507 / ENSG00000131469
RPL28 / P46779 / ENSSSCG00000039460 / ENSG00000108107
RPL3 / P39023 / ENSSSCG00000040494 / ENSG00000100316
RPL30 / P62888 / ENSSSCG00000006081 / ENSG00000156482
RPL32 / P62910 / ENSSSCG00000035811 / ENSG00000144713
RPL34 / P49207 / ENSSSCG00000009146 / ENSG00000109475
RPL35 / P42766 / ENSSSCG00000034227 / ENSG00000136942
Homologs
Adipose.Tissue
Brain
Colon
Heart
Kidney
Liver
Lung
Pancreas
S.Intestine
Spleen
Organs
Species
Human
Pig
Bins
1
2
3
4
5

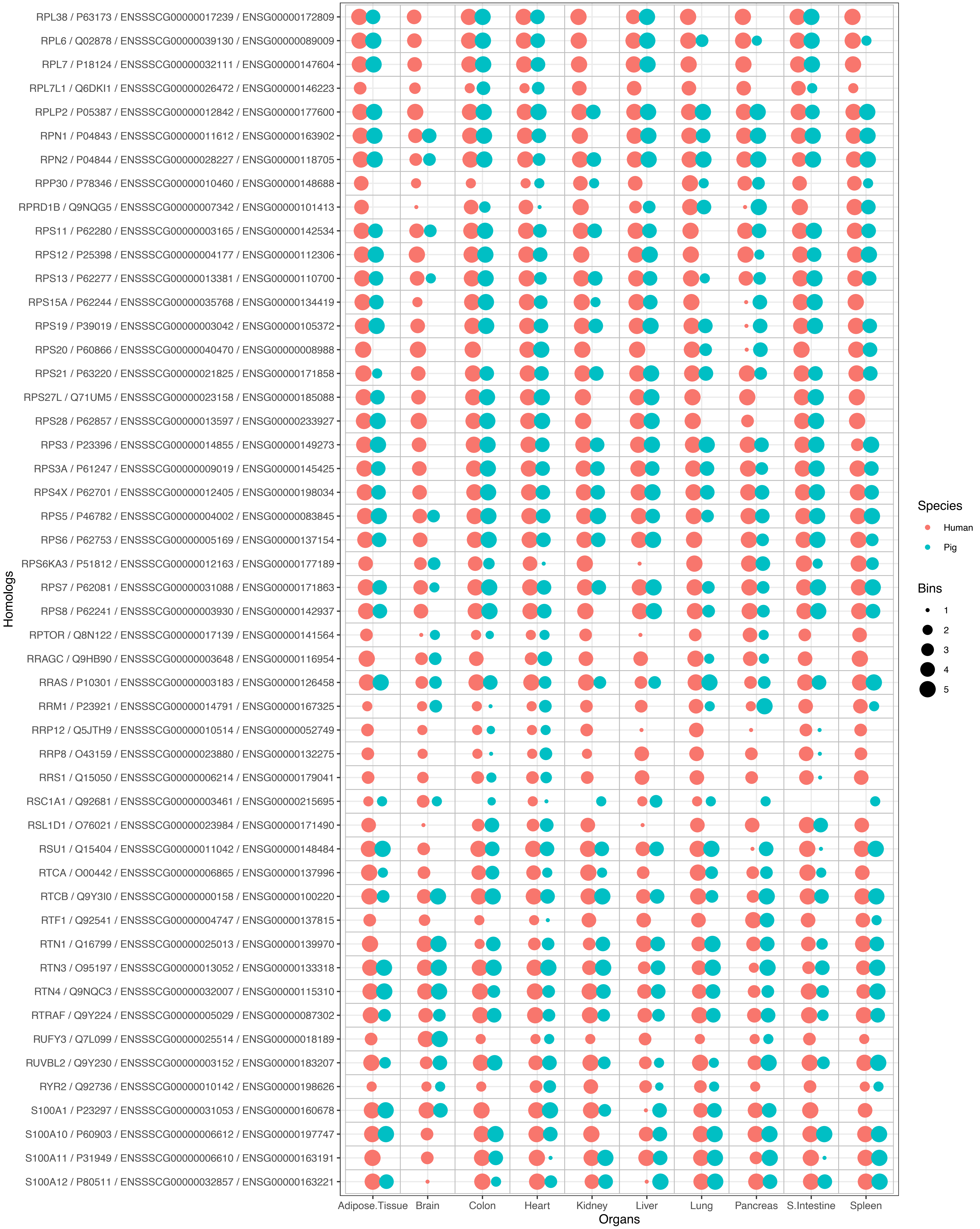
RPL38 / P63173 / ENSSSCG00000017239 / ENSG00000172809
RPL6 / Q02878 / ENSSSCG00000039130 / ENSG00000089009
RPL7 / P18124 / ENSSSCG00000032111 / ENSG00000147604
RPL7L1 / Q6DKI1 / ENSSSCG00000026472 / ENSG00000146223
RPLP2 / P05387 / ENSSSCG00000012842 / ENSG00000177600
RPN1 / P04843 / ENSSSCG00000011612 / ENSG00000163902
RPN2 / P04844 / ENSSSCG00000028227 / ENSG00000118705
RPP30 / P78346 / ENSSSCG00000010460 / ENSG00000148688
RPRD1B / Q9NQG5 / ENSSSCG00000007342 / ENSG00000101413
RPS11 / P62280 / ENSSSCG00000003165 / ENSG00000142534
RPS12 / P25398 / ENSSSCG00000004177 / ENSG00000112306
RPS13 / P62277 / ENSSSCG00000013381 / ENSG00000110700
RPS15A / P62244 / ENSSSCG00000035768 / ENSG00000134419
RPS19 / P39019 / ENSSSCG00000003042 / ENSG00000105372
RPS20 / P60866 / ENSSSCG00000040470 / ENSG00000008988
RPS21 / P63220 / ENSSSCG00000021825 / ENSG00000171858
RPS27L / Q71UM5 / ENSSSCG00000023158 / ENSG00000185088
RPS28 / P62857 / ENSSSCG00000013597 / ENSG00000233927
RPS3 / P23396 / ENSSSCG00000014855 / ENSG00000149273
RPS3A / P61247 / ENSSSCG00000009019 / ENSG00000145425
RPS4X / P62701 / ENSSSCG00000012405 / ENSG00000198034
RPS5 / P46782 / ENSSSCG00000004002 / ENSG00000083845
RPS6 / P62753 / ENSSSCG00000005169 / ENSG00000137154
RPS6KA3 / P51812 / ENSSSCG00000012163 / ENSG00000177189
RPS7 / P62081 / ENSSSCG00000031088 / ENSG00000171863
RPS8 / P62241 / ENSSSCG00000003930 / ENSG00000142937
RPTOR / Q8N122 / ENSSSCG00000017139 / ENSG00000141564
RRAGC / Q9HB90 / ENSSSCG00000003648 / ENSG00000116954
RRAS / P10301 / ENSSSCG00000003183 / ENSG00000126458
RRM1 / P23921 / ENSSSCG00000014791 / ENSG00000167325
RRP12 / Q5JTH9 / ENSSSCG00000010514 / ENSG00000052749
RRP8 / O43159 / ENSSSCG00000023880 / ENSG00000132275
RRS1 / Q15050 / ENSSSCG00000006214 / ENSG00000179041
RSC1A1 / Q92681 / ENSSSCG00000003461 / ENSG00000215695
RSL1D1 / O76021 / ENSSSCG00000023984 / ENSG00000171490
RSU1 / Q15404 / ENSSSCG00000011042 / ENSG00000148484
RTCA / O00442 / ENSSSCG00000006865 / ENSG00000137996
RTCB / Q9Y3I0 / ENSSSCG00000000158 / ENSG00000100220
RTF1 / Q92541 / ENSSSCG00000004747 / ENSG00000137815
RTN1 / Q16799 / ENSSSCG00000025013 / ENSG00000139970
RTN3 / O95197 / ENSSSCG00000013052 / ENSG00000133318
RTN4 / Q9NQC3 / ENSSSCG00000032007 / ENSG00000115310
RTRAF / Q9Y224 / ENSSSCG00000005029 / ENSG00000087302
RUFY3 / Q7L099 / ENSSSCG00000025514 / ENSG00000018189
RUVBL2 / Q9Y230 / ENSSSCG00000003152 / ENSG00000183207
RYR2 / Q92736 / ENSSSCG00000010142 / ENSG00000198626
S100A1 / P23297 / ENSSSCG00000031053 / ENSG00000160678
S100A10 / P60903 / ENSSSCG00000006612 / ENSG00000197747
S100A11 / P31949 / ENSSSCG00000006610 / ENSG00000163191
S100A12 / P80511 / ENSSSCG00000032857 / ENSG00000163221
Homologs
Adipose.Tissue
Brain
Colon
Heart
Kidney
Liver
Lung
Pancreas
S.Intestine
Spleen
Organs
Species
Human
Pig
Bins
1
2
3
4
5

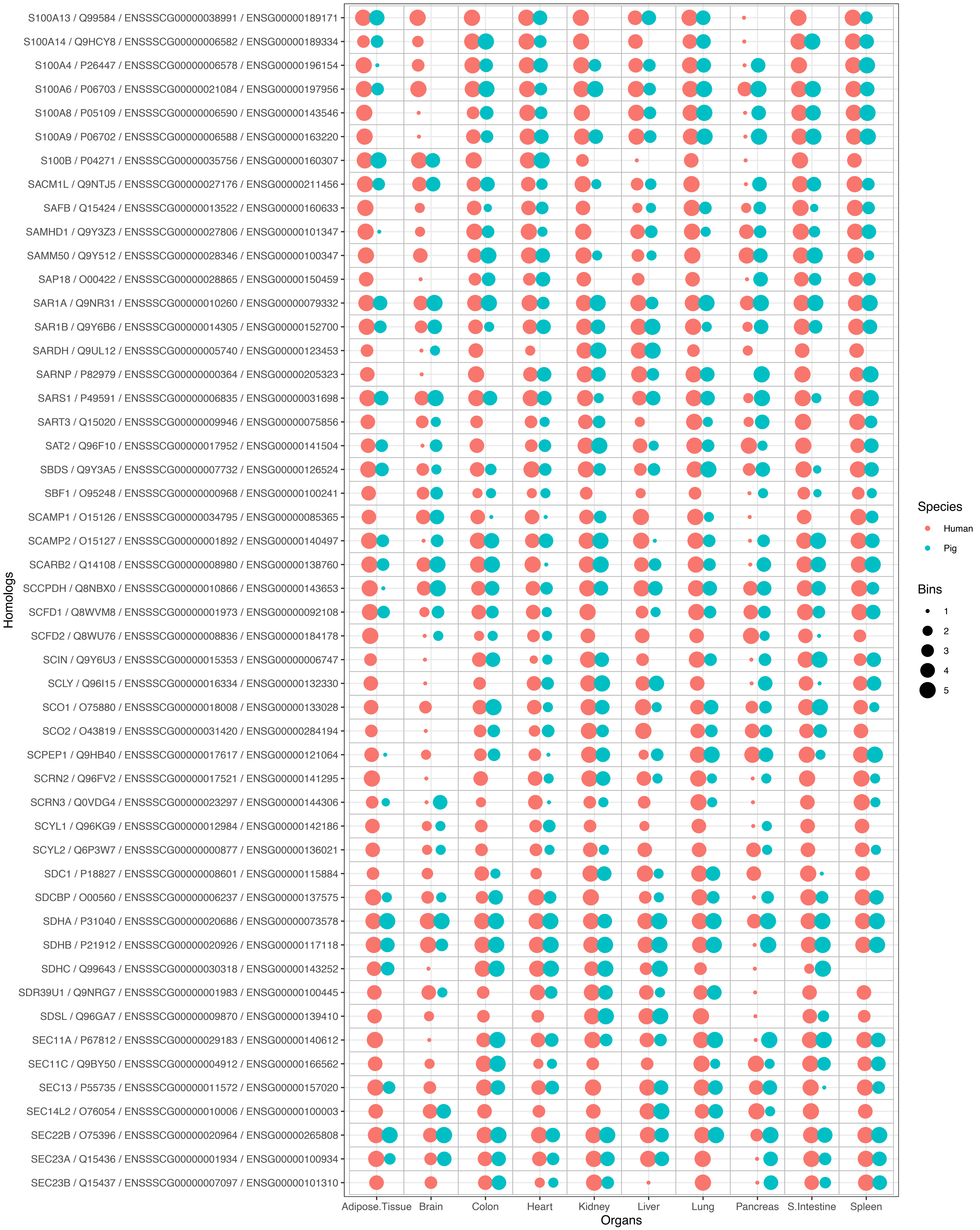
S100A13 / Q99584 / ENSSSCG00000038991 / ENSG00000189171
S100A14 / Q9HCY8 / ENSSSCG00000006582 / ENSG00000189334
S100A4 / P26447 / ENSSSCG00000006578 / ENSG00000196154
S100A6 / P06703 / ENSSSCG00000021084 / ENSG00000197956
S100A8 / P05109 / ENSSSCG00000006590 / ENSG00000143546
S100A9 / P06702 / ENSSSCG00000006588 / ENSG00000163220
S100B / P04271 / ENSSSCG00000035756 / ENSG00000160307
SACM1L / Q9NTJ5 / ENSSSCG00000027176 / ENSG00000211456
SAFB / Q15424 / ENSSSCG00000013522 / ENSG00000160633
SAMHD1 / Q9Y3Z3 / ENSSSCG00000027806 / ENSG00000101347
SAMM50 / Q9Y512 / ENSSSCG00000028346 / ENSG00000100347
SAP18 / O00422 / ENSSSCG00000028865 / ENSG00000150459
SAR1A / Q9NR31 / ENSSSCG00000010260 / ENSG00000079332
SAR1B / Q9Y6B6 / ENSSSCG00000014305 / ENSG00000152700
SARDH / Q9UL12 / ENSSSCG00000005740 / ENSG00000123453
SARNP / P82979 / ENSSSCG00000000364 / ENSG00000205323
SARS1 / P49591 / ENSSSCG00000006835 / ENSG00000031698
SART3 / Q15020 / ENSSSCG00000009946 / ENSG00000075856
SAT2 / Q96F10 / ENSSSCG00000017952 / ENSG00000141504
SBDS / Q9Y3A5 / ENSSSCG00000007732 / ENSG00000126524
SBF1 / O95248 / ENSSSCG00000000968 / ENSG00000100241
SCAMP1 / O15126 / ENSSSCG00000034795 / ENSG00000085365
SCAMP2 / O15127 / ENSSSCG00000001892 / ENSG00000140497
SCARB2 / Q14108 / ENSSSCG00000008980 / ENSG00000138760
SCCPDH / Q8NBX0 / ENSSSCG00000010866 / ENSG00000143653
SCFD1 / Q8WVM8 / ENSSSCG00000001973 / ENSG00000092108
SCFD2 / Q8WU76 / ENSSSCG00000008836 / ENSG00000184178
SCIN / Q9Y6U3 / ENSSSCG00000015353 / ENSG00000006747
SCLY / Q96I15 / ENSSSCG00000016334 / ENSG00000132330
SCO1 / O75880 / ENSSSCG00000018008 / ENSG00000133028
SCO2 / O43819 / ENSSSCG00000031420 / ENSG00000284194
SCPEP1 / Q9HB40 / ENSSSCG00000017617 / ENSG00000121064
SCRN2 / Q96FV2 / ENSSSCG00000017521 / ENSG00000141295
SCRN3 / Q0VDG4 / ENSSSCG00000023297 / ENSG00000144306
SCYL1 / Q96KG9 / ENSSSCG00000012984 / ENSG00000142186
SCYL2 / Q6P3W7 / ENSSSCG00000000877 / ENSG00000136021
SDC1 / P18827 / ENSSSCG00000008601 / ENSG00000115884
SDCBP / O00560 / ENSSSCG00000006237 / ENSG00000137575
SDHA / P31040 / ENSSSCG00000020686 / ENSG00000073578
SDHB / P21912 / ENSSSCG00000020926 / ENSG00000117118
SDHC / Q99643 / ENSSSCG00000030318 / ENSG00000143252
SDR39U1 / Q9NRG7 / ENSSSCG00000001983 / ENSG00000100445
SDSL / Q96GA7 / ENSSSCG00000009870 / ENSG00000139410
SEC11A / P67812 / ENSSSCG00000029183 / ENSG00000140612
SEC11C / Q9BY50 / ENSSSCG00000004912 / ENSG00000166562
SEC13 / P55735 / ENSSSCG00000011572 / ENSG00000157020
SEC14L2 / O76054 / ENSSSCG00000010006 / ENSG00000100003
SEC22B / O75396 / ENSSSCG00000020964 / ENSG00000265808
SEC23A / Q15436 / ENSSSCG00000001934 / ENSG00000100934
SEC23B / Q15437 / ENSSSCG00000007097 / ENSG00000101310
Homologs
Adipose.Tissue
Brain
Colon
Heart
Kidney
Liver
Lung
Pancreas
S.Intestine
Spleen
Organs
Species
Human
Pig
Bins
1
2
3
4
5

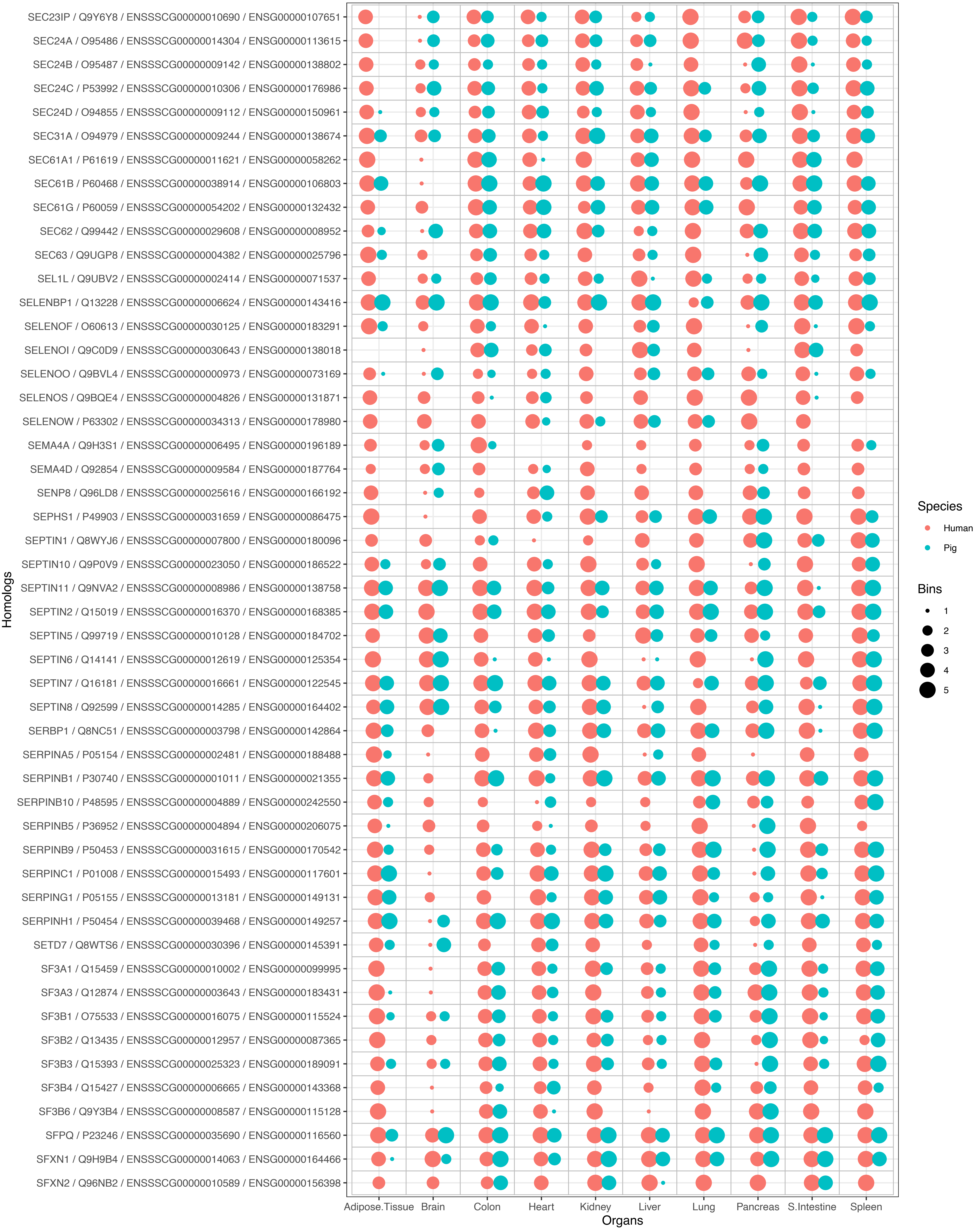

SEC23IP / Q9Y6Y8 / ENSSSCG00000010690 / ENSG00000107651
SEC24A / O95486 / ENSSSCG00000014304 / ENSG00000113615
SEC24B / O95487 / ENSSSCG00000009142 / ENSG00000138802
SEC24C / P53992 / ENSSSCG00000010306 / ENSG00000176986
SEC24D / O94855 / ENSSSCG00000009112 / ENSG00000150961
SEC31A / O94979 / ENSSSCG00000009244 / ENSG00000138674
SEC61A1 / P61619 / ENSSSCG00000011621 / ENSG00000058262
SEC61B / P60468 / ENSSSCG00000038914 / ENSG00000106803
SEC61G / P60059 / ENSSSCG00000054202 / ENSG00000132432
SEC62 / Q99442 / ENSSSCG00000029608 / ENSG00000008952
SEC63 / Q9UGP8 / ENSSSCG00000004382 / ENSG00000025796
SEL1L / Q9UBV2 / ENSSSCG00000002414 / ENSG00000071537
SELENBP1 / Q13228 / ENSSSCG00000006624 / ENSG00000143416
SELENOF / O60613 / ENSSSCG00000030125 / ENSG00000183291
SELENOI / Q9C0D9 / ENSSSCG00000030643 / ENSG00000138018
SELENOO / Q9BVL4 / ENSSSCG00000000973 / ENSG00000073169
SELENOS / Q9BQE4 / ENSSSCG00000004826 / ENSG00000131871
SELENOW / P63302 / ENSSSCG00000034313 / ENSG00000178980
SEMA4A / Q9H3S1 / ENSSSCG00000006495 / ENSG00000196189
SEMA4D / Q92854 / ENSSSCG00000009584 / ENSG00000187764
SENP8 / Q96LD8 / ENSSSCG00000025616 / ENSG00000166192
SEPHS1 / P49903 / ENSSSCG00000031659 / ENSG00000086475
SEPTIN1 / Q8WYJ6 / ENSSSCG00000007800 / ENSG00000180096
SEPTIN10 / Q9P0V9 / ENSSSCG00000023050 / ENSG00000186522
SEPTIN11 / Q9NVA2 / ENSSSCG00000008986 / ENSG00000138758
SEPTIN2 / Q15019 / ENSSSCG00000016370 / ENSG00000168385
SEPTIN5 / Q99719 / ENSSSCG00000010128 / ENSG00000184702
SEPTIN6 / Q14141 / ENSSSCG00000012619 / ENSG00000125354
SEPTIN7 / Q16181 / ENSSSCG00000016661 / ENSG00000122545
SEPTIN8 / Q92599 / ENSSSCG00000014285 / ENSG00000164402
SERBP1 / Q8NC51 / ENSSSCG00000003798 / ENSG00000142864
SERPINA5 / P05154 / ENSSSCG00000002481 / ENSG00000188488
SERPINB1 / P30740 / ENSSSCG00000001011 / ENSG00000021355
SERPINB10 / P48595 / ENSSSCG00000004889 / ENSG00000242550
SERPINB5 / P36952 / ENSSSCG00000004894 / ENSG00000206075
SERPINB9 / P50453 / ENSSSCG00000031615 / ENSG00000170542
SERPINC1 / P01008 / ENSSSCG00000015493 / ENSG00000117601
SERPING1 / P05155 / ENSSSCG00000013181 / ENSG00000149131
SERPINH1 / P50454 / ENSSSCG00000039468 / ENSG00000149257
SETD7 / Q8WTS6 / ENSSSCG00000030396 / ENSG00000145391
SF3A1 / Q15459 / ENSSSCG00000010002 / ENSG00000099995
SF3A3 / Q12874 / ENSSSCG00000003643 / ENSG00000183431
SF3B1 / O75533 / ENSSSCG00000016075 / ENSG00000115524
SF3B2 / Q13435 / ENSSSCG00000012957 / ENSG00000087365
SF3B3 / Q15393 / ENSSSCG00000025323 / ENSG00000189091
SF3B4 / Q15427 / ENSSSCG00000006665 / ENSG00000143368
SF3B6 / Q9Y3B4 / ENSSSCG00000008587 / ENSG00000115128
SFPQ / P23246 / ENSSSCG00000035690 / ENSG00000116560
SFXN1 / Q9H9B4 / ENSSSCG00000014063 / ENSG00000164466
SFXN2 / Q96NB2 / ENSSSCG00000010589 / ENSG00000156398
Homologs
Adipose.Tissue
Brain
Colon
Heart
Kidney
Liver
Lung
Pancreas
S.Intestine
Spleen
Organs
Species
Human
Pig
Bins
1
2
3
4
5

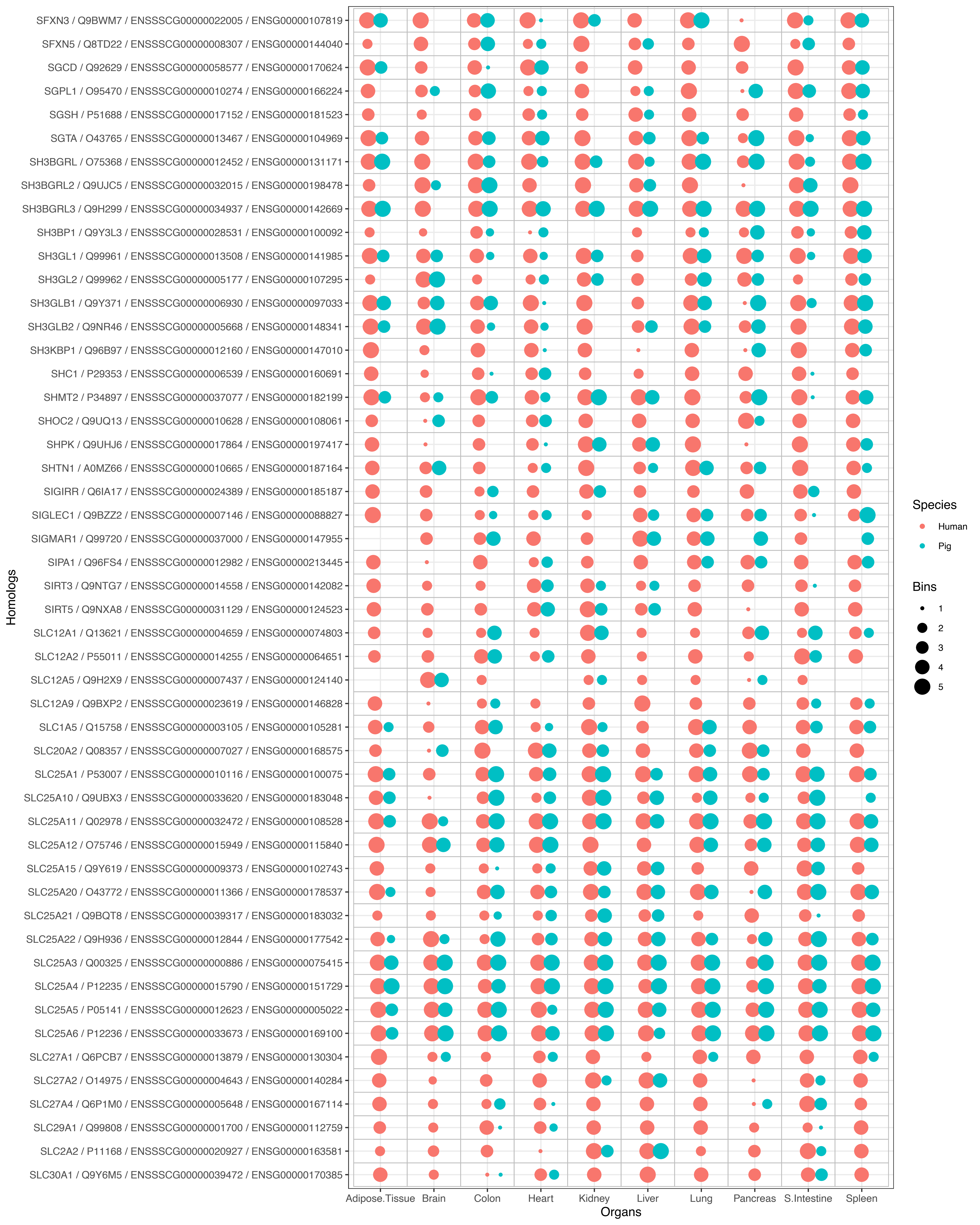
SFXN3 / Q9BWM7 / ENSSSCG00000022005 / ENSG00000107819
SFXN5 / Q8TD22 / ENSSSCG00000008307 / ENSG00000144040
SGCD / Q92629 / ENSSSCG00000058577 / ENSG00000170624
SGPL1 / O95470 / ENSSSCG00000010274 / ENSG00000166224
SGSH / P51688 / ENSSSCG00000017152 / ENSG00000181523
SGTA / O43765 / ENSSSCG00000013467 / ENSG00000104969
SH3BGRL / O75368 / ENSSSCG00000012452 / ENSG00000131171
SH3BGRL2 / Q9UJC5 / ENSSSCG00000032015 / ENSG00000198478
SH3BGRL3 / Q9H299 / ENSSSCG00000034937 / ENSG00000142669
SH3BP1 / Q9Y3L3 / ENSSSCG00000028531 / ENSG00000100092
SH3GL1 / Q99961 / ENSSSCG00000013508 / ENSG00000141985
SH3GL2 / Q99962 / ENSSSCG00000005177 / ENSG00000107295
SH3GLB1 / Q9Y371 / ENSSSCG00000006930 / ENSG00000097033
SH3GLB2 / Q9NR46 / ENSSSCG00000005668 / ENSG00000148341
SH3KBP1 / Q96B97 / ENSSSCG00000012160 / ENSG00000147010
SHC1 / P29353 / ENSSSCG00000006539 / ENSG00000160691
SHMT2 / P34897 / ENSSSCG00000037077 / ENSG00000182199
SHOC2 / Q9UQ13 / ENSSSCG00000010628 / ENSG00000108061
SHPK / Q9UHJ6 / ENSSSCG00000017864 / ENSG00000197417
SHTN1 / A0MZ66 / ENSSSCG00000010665 / ENSG00000187164
SIGIRR / Q6IA17 / ENSSSCG00000024389 / ENSG00000185187
SIGLEC1 / Q9BZZ2 / ENSSSCG00000007146 / ENSG00000088827
SIGMAR1 / Q99720 / ENSSSCG00000037000 / ENSG00000147955
SIPA1 / Q96FS4 / ENSSSCG00000012982 / ENSG00000213445
SIRT3 / Q9NTG7 / ENSSSCG00000014558 / ENSG00000142082
SIRT5 / Q9NXA8 / ENSSSCG00000031129 / ENSG00000124523
SLC12A1 / Q13621 / ENSSSCG00000004659 / ENSG00000074803
SLC12A2 / P55011 / ENSSSCG00000014255 / ENSG00000064651
SLC12A5 / Q9H2X9 / ENSSSCG00000007437 / ENSG00000124140
SLC12A9 / Q9BXP2 / ENSSSCG00000023619 / ENSG00000146828
SLC1A5 / Q15758 / ENSSSCG00000003105 / ENSG00000105281
SLC20A2 / Q08357 / ENSSSCG00000007027 / ENSG00000168575
SLC25A1 / P53007 / ENSSSCG00000010116 / ENSG00000100075
SLC25A10 / Q9UBX3 / ENSSSCG00000033620 / ENSG00000183048
SLC25A11 / Q02978 / ENSSSCG00000032472 / ENSG00000108528
SLC25A12 / O75746 / ENSSSCG00000015949 / ENSG00000115840
SLC25A15 / Q9Y619 / ENSSSCG00000009373 / ENSG00000102743
SLC25A20 / O43772 / ENSSSCG00000011366 / ENSG00000178537
SLC25A21 / Q9BQT8 / ENSSSCG00000039317 / ENSG00000183032
SLC25A22 / Q9H936 / ENSSSCG00000012844 / ENSG00000177542
SLC25A3 / Q00325 / ENSSSCG00000000886 / ENSG00000075415
SLC25A4 / P12235 / ENSSSCG00000015790 / ENSG00000151729
SLC25A5 / P05141 / ENSSSCG00000012623 / ENSG00000005022
SLC25A6 / P12236 / ENSSSCG00000033673 / ENSG00000169100
SLC27A1 / Q6PCB7 / ENSSSCG00000013879 / ENSG00000130304
SLC27A2 / O14975 / ENSSSCG00000004643 / ENSG00000140284
SLC27A4 / Q6P1M0 / ENSSSCG00000005648 / ENSG00000167114
SLC29A1 / Q99808 / ENSSSCG00000001700 / ENSG00000112759
SLC2A2 / P11168 / ENSSSCG00000020927 / ENSG00000163581
SLC30A1 / Q9Y6M5 / ENSSSCG00000039472 / ENSG00000170385
Homologs
Adipose.Tissue
Brain
Colon
Heart
Kidney
Liver
Lung
Pancreas
S.Intestine
Spleen
Organs
Species
Human
Pig
Bins
1
2
3
4
5

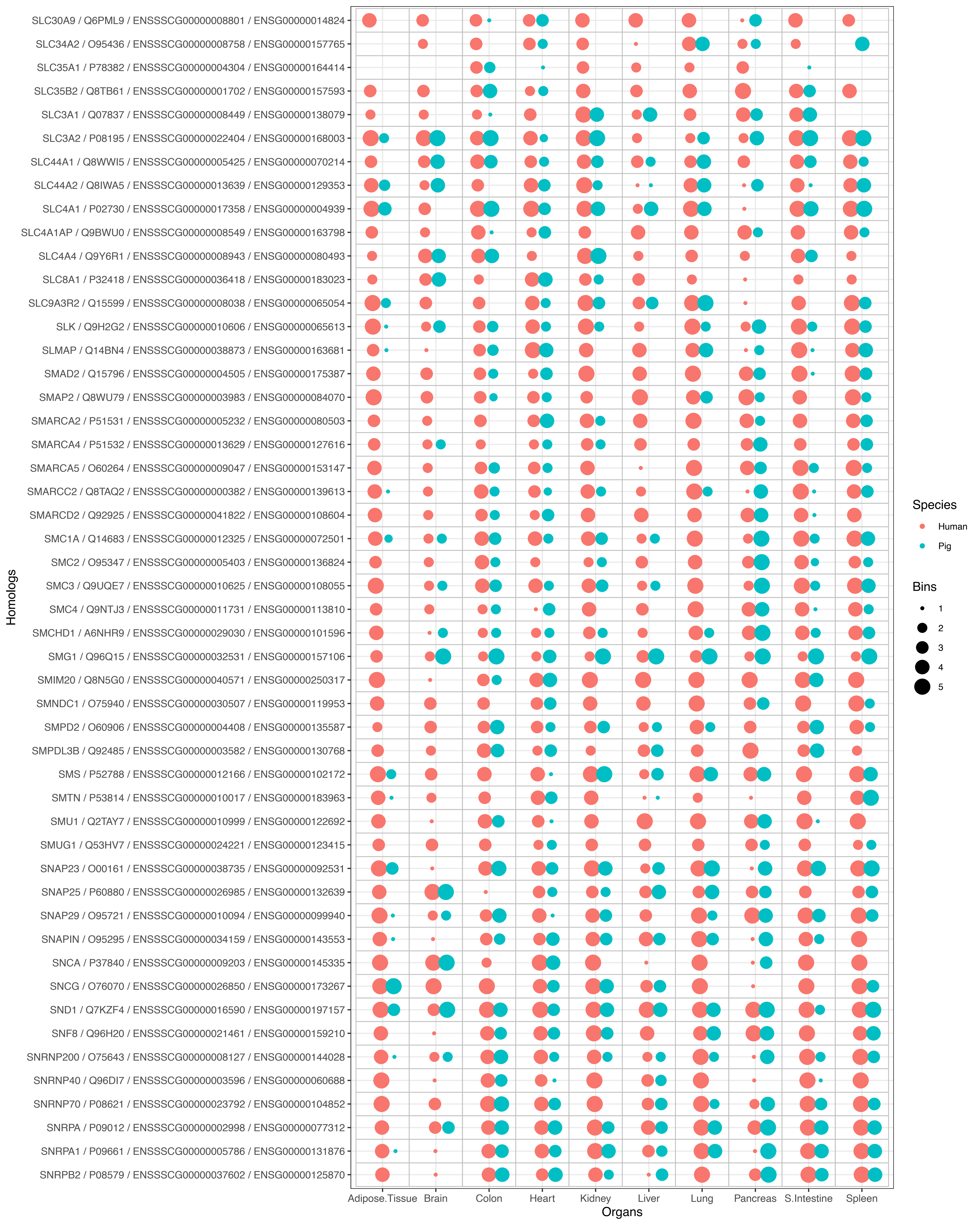
SLC30A9 / Q6PML9 / ENSSSCG00000008801 / ENSG00000014824
SLC34A2 / O95436 / ENSSSCG00000008758 / ENSG00000157765
SLC35A1 / P78382 / ENSSSCG00000004304 / ENSG00000164414
SLC35B2 / Q8TB61 / ENSSSCG00000001702 / ENSG00000157593
SLC3A1 / Q07837 / ENSSSCG00000008449 / ENSG00000138079
SLC3A2 / P08195 / ENSSSCG00000022404 / ENSG00000168003
SLC44A1 / Q8WWI5 / ENSSSCG00000005425 / ENSG00000070214
SLC44A2 / Q8IWA5 / ENSSSCG00000013639 / ENSG00000129353
SLC4A1 / P02730 / ENSSSCG00000017358 / ENSG00000004939
SLC4A1AP / Q9BWU0 / ENSSSCG00000008549 / ENSG00000163798
SLC4A4 / Q9Y6R1 / ENSSSCG00000008943 / ENSG00000080493
SLC8A1 / P32418 / ENSSSCG00000036418 / ENSG00000183023
SLC9A3R2 / Q15599 / ENSSSCG00000008038 / ENSG00000065054
SLK / Q9H2G2 / ENSSSCG00000010606 / ENSG00000065613
SLMAP / Q14BN4 / ENSSSCG00000038873 / ENSG00000163681
SMAD2 / Q15796 / ENSSSCG00000004505 / ENSG00000175387
SMAP2 / Q8WU79 / ENSSSCG00000003983 / ENSG00000084070
SMARCA2 / P51531 / ENSSSCG00000005232 / ENSG00000080503
SMARCA4 / P51532 / ENSSSCG00000013629 / ENSG00000127616
SMARCA5 / O60264 / ENSSSCG00000009047 / ENSG00000153147
SMARCC2 / Q8TAQ2 / ENSSSCG00000000382 / ENSG00000139613
SMARCD2 / Q92925 / ENSSSCG00000041822 / ENSG00000108604
SMC1A / Q14683 / ENSSSCG00000012325 / ENSG00000072501
SMC2 / O95347 / ENSSSCG00000005403 / ENSG00000136824
SMC3 / Q9UQE7 / ENSSSCG00000010625 / ENSG00000108055
SMC4 / Q9NTJ3 / ENSSSCG00000011731 / ENSG00000113810
SMCHD1 / A6NHR9 / ENSSSCG00000029030 / ENSG00000101596
SMG1 / Q96Q15 / ENSSSCG00000032531 / ENSG00000157106
SMIM20 / Q8N5G0 / ENSSSCG00000040571 / ENSG00000250317
SMNDC1 / O75940 / ENSSSCG00000030507 / ENSG00000119953
SMPD2 / O60906 / ENSSSCG00000004408 / ENSG00000135587
SMPDL3B / Q92485 / ENSSSCG00000003582 / ENSG00000130768
SMS / P52788 / ENSSSCG00000012166 / ENSG00000102172
SMTN / P53814 / ENSSSCG00000010017 / ENSG00000183963
SMU1 / Q2TAY7 / ENSSSCG00000010999 / ENSG00000122692
SMUG1 / Q53HV7 / ENSSSCG00000024221 / ENSG00000123415
SNAP23 / O00161 / ENSSSCG00000038735 / ENSG00000092531
SNAP25 / P60880 / ENSSSCG00000026985 / ENSG00000132639
SNAP29 / O95721 / ENSSSCG00000010094 / ENSG00000099940
SNAPIN / O95295 / ENSSSCG00000034159 / ENSG00000143553
SNCA / P37840 / ENSSSCG00000009203 / ENSG00000145335
SNCG / O76070 / ENSSSCG00000026850 / ENSG00000173267
SND1 / Q7KZF4 / ENSSSCG00000016590 / ENSG00000197157
SNF8 / Q96H20 / ENSSSCG00000021461 / ENSG00000159210
SNRNP200 / O75643 / ENSSSCG00000008127 / ENSG00000144028
SNRNP40 / Q96DI7 / ENSSSCG00000003596 / ENSG00000060688
SNRNP70 / P08621 / ENSSSCG00000023792 / ENSG00000104852
SNRPA / P09012 / ENSSSCG00000002998 / ENSG00000077312
SNRPA1 / P09661 / ENSSSCG00000005786 / ENSG00000131876
SNRPB2 / P08579 / ENSSSCG00000037602 / ENSG00000125870
Homologs
Adipose.Tissue
Brain
Colon
Heart
Kidney
Liver
Lung
Pancreas
S.Intestine
Spleen
Organs
Species
Human
Pig
Bins
1
2
3
4
5

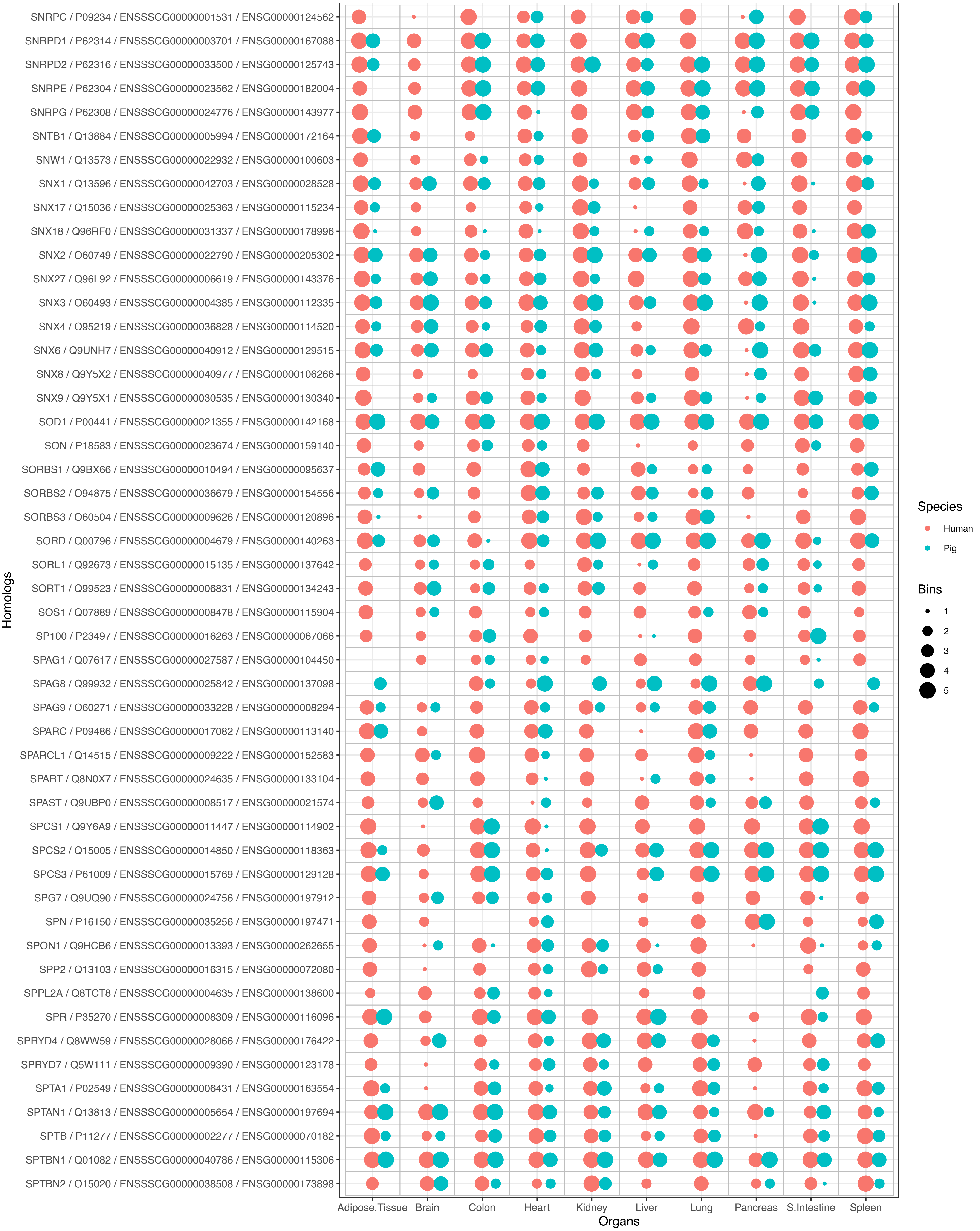
SNRPC / P09234 / ENSSSCG00000001531 / ENSG00000124562
SNRPD1 / P62314 / ENSSSCG00000003701 / ENSG00000167088
SNRPD2 / P62316 / ENSSSCG00000033500 / ENSG00000125743
SNRPE / P62304 / ENSSSCG00000023562 / ENSG00000182004
SNRPG / P62308 / ENSSSCG00000024776 / ENSG00000143977
SNTB1 / Q13884 / ENSSSCG00000005994 / ENSG00000172164
SNW1 / Q13573 / ENSSSCG00000022932 / ENSG00000100603
SNX1 / Q13596 / ENSSSCG00000042703 / ENSG00000028528
SNX17 / Q15036 / ENSSSCG00000025363 / ENSG00000115234
SNX18 / Q96RF0 / ENSSSCG00000031337 / ENSG00000178996
SNX2 / O60749 / ENSSSCG00000022790 / ENSG00000205302
SNX27 / Q96L92 / ENSSSCG00000006619 / ENSG00000143376
SNX3 / O60493 / ENSSSCG00000004385 / ENSG00000112335
SNX4 / O95219 / ENSSSCG00000036828 / ENSG00000114520
SNX6 / Q9UNH7 / ENSSSCG00000040912 / ENSG00000129515
SNX8 / Q9Y5X2 / ENSSSCG00000040977 / ENSG00000106266
SNX9 / Q9Y5X1 / ENSSSCG00000030535 / ENSG00000130340
SOD1 / P00441 / ENSSSCG00000021355 / ENSG00000142168
SON / P18583 / ENSSSCG00000023674 / ENSG00000159140
SORBS1 / Q9BX66 / ENSSSCG00000010494 / ENSG00000095637
SORBS2 / O94875 / ENSSSCG00000036679 / ENSG00000154556
SORBS3 / O60504 / ENSSSCG00000009626 / ENSG00000120896
SORD / Q00796 / ENSSSCG00000004679 / ENSG00000140263
SORL1 / Q92673 / ENSSSCG00000015135 / ENSG00000137642
SORT1 / Q99523 / ENSSSCG00000006831 / ENSG00000134243
SOS1 / Q07889 / ENSSSCG00000008478 / ENSG00000115904
SP100 / P23497 / ENSSSCG00000016263 / ENSG00000067066
SPAG1 / Q07617 / ENSSSCG00000027587 / ENSG00000104450
SPAG8 / Q99932 / ENSSSCG00000025842 / ENSG00000137098
SPAG9 / O60271 / ENSSSCG00000033228 / ENSG00000008294
SPARC / P09486 / ENSSSCG00000017082 / ENSG00000113140
SPARCL1 / Q14515 / ENSSSCG00000009222 / ENSG00000152583
SPART / Q8N0X7 / ENSSSCG00000024635 / ENSG00000133104
SPAST / Q9UBP0 / ENSSSCG00000008517 / ENSG00000021574
SPCS1 / Q9Y6A9 / ENSSSCG00000011447 / ENSG00000114902
SPCS2 / Q15005 / ENSSSCG00000014850 / ENSG00000118363
SPCS3 / P61009 / ENSSSCG00000015769 / ENSG00000129128
SPG7 / Q9UQ90 / ENSSSCG00000024756 / ENSG00000197912
SPN / P16150 / ENSSSCG00000035256 / ENSG00000197471
SPON1 / Q9HCB6 / ENSSSCG00000013393 / ENSG00000262655
SPP2 / Q13103 / ENSSSCG00000016315 / ENSG00000072080
SPPL2A / Q8TCT8 / ENSSSCG00000004635 / ENSG00000138600
SPR / P35270 / ENSSSCG00000008309 / ENSG00000116096
SPRYD4 / Q8WW59 / ENSSSCG00000028066 / ENSG00000176422
SPRYD7 / Q5W111 / ENSSSCG00000009390 / ENSG00000123178
SPTA1 / P02549 / ENSSSCG00000006431 / ENSG00000163554
SPTAN1 / Q13813 / ENSSSCG00000005654 / ENSG00000197694
SPTB / P11277 / ENSSSCG00000002277 / ENSG00000070182
SPTBN1 / Q01082 / ENSSSCG00000040786 / ENSG00000115306
SPTBN2 / O15020 / ENSSSCG00000038508 / ENSG00000173898
Homologs
Adipose.Tissue
Brain
Colon
Heart
Kidney
Liver
Lung
Pancreas
S.Intestine
Spleen
Organs
Species
Human
Pig
Bins
1
2
3
4
5

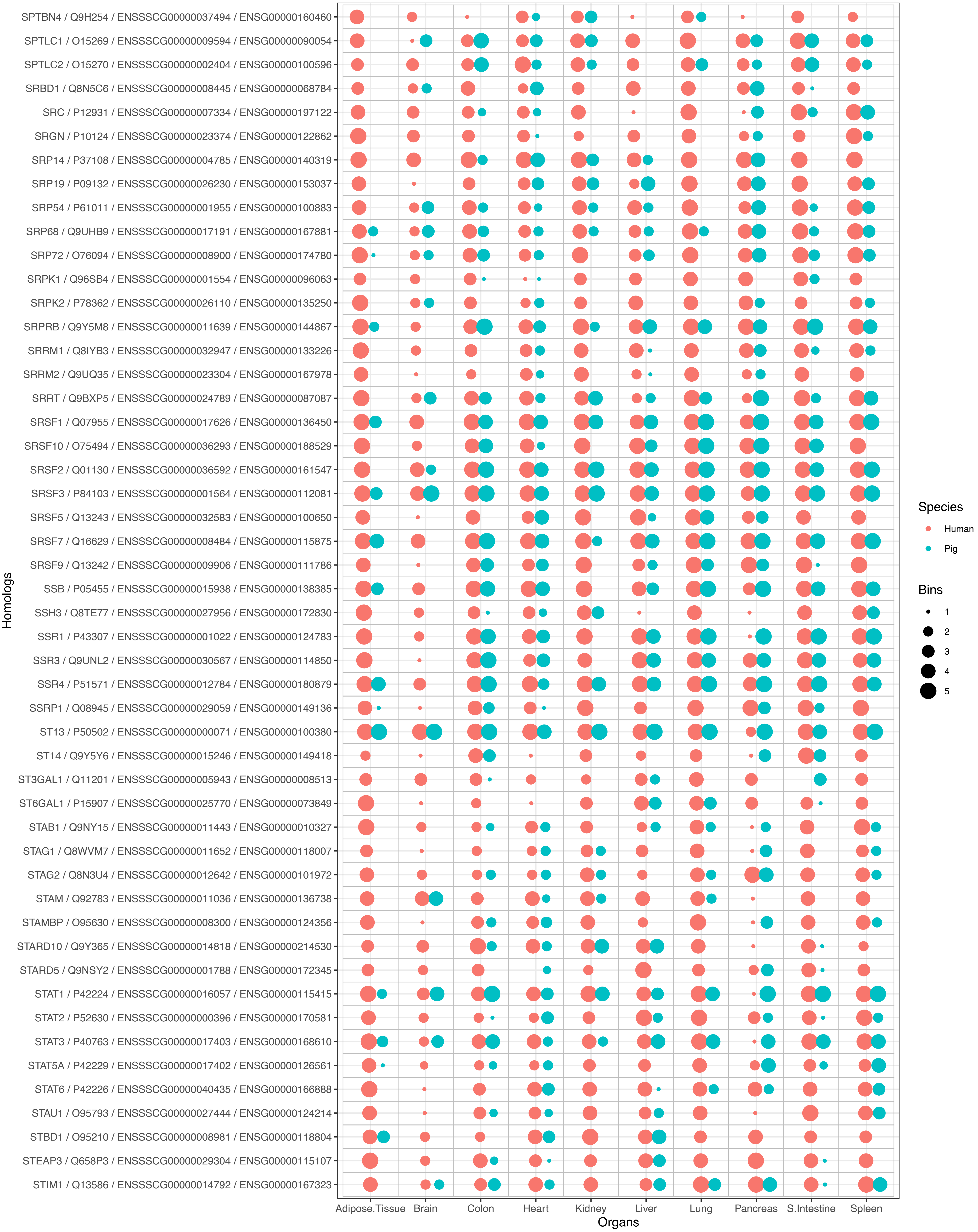
SPTBN4 / Q9H254 / ENSSSCG00000037494 / ENSG00000160460
SPTLC1 / O15269 / ENSSSCG00000009594 / ENSG00000090054
SPTLC2 / O15270 / ENSSSCG00000002404 / ENSG00000100596
SRBD1 / Q8N5C6 / ENSSSCG00000008445 / ENSG00000068784
SRC / P12931 / ENSSSCG00000007334 / ENSG00000197122
SRGN / P10124 / ENSSSCG00000023374 / ENSG00000122862
SRP14 / P37108 / ENSSSCG00000004785 / ENSG00000140319
SRP19 / P09132 / ENSSSCG00000026230 / ENSG00000153037
SRP54 / P61011 / ENSSSCG00000001955 / ENSG00000100883
SRP68 / Q9UHB9 / ENSSSCG00000017191 / ENSG00000167881
SRP72 / O76094 / ENSSSCG00000008900 / ENSG00000174780
SRPK1 / Q96SB4 / ENSSSCG00000001554 / ENSG00000096063
SRPK2 / P78362 / ENSSSCG00000026110 / ENSG00000135250
SRPRB / Q9Y5M8 / ENSSSCG00000011639 / ENSG00000144867
SRRM1 / Q8IYB3 / ENSSSCG00000032947 / ENSG00000133226
SRRM2 / Q9UQ35 / ENSSSCG00000023304 / ENSG00000167978
SRRT / Q9BXP5 / ENSSSCG00000024789 / ENSG00000087087
SRSF1 / Q07955 / ENSSSCG00000017626 / ENSG00000136450
SRSF10 / O75494 / ENSSSCG00000036293 / ENSG00000188529
SRSF2 / Q01130 / ENSSSCG00000036592 / ENSG00000161547
SRSF3 / P84103 / ENSSSCG00000001564 / ENSG00000112081
SRSF5 / Q13243 / ENSSSCG00000032583 / ENSG00000100650
SRSF7 / Q16629 / ENSSSCG00000008484 / ENSG00000115875
SRSF9 / Q13242 / ENSSSCG00000009906 / ENSG00000111786
SSB / P05455 / ENSSSCG00000015938 / ENSG00000138385
SSH3 / Q8TE77 / ENSSSCG00000027956 / ENSG00000172830
SSR1 / P43307 / ENSSSCG00000001022 / ENSG00000124783
SSR3 / Q9UNL2 / ENSSSCG00000030567 / ENSG00000114850
SSR4 / P51571 / ENSSSCG00000012784 / ENSG00000180879
SSRP1 / Q08945 / ENSSSCG00000029059 / ENSG00000149136
ST13 / P50502 / ENSSSCG00000000071 / ENSG00000100380
ST14 / Q9Y5Y6 / ENSSSCG00000015246 / ENSG00000149418
ST3GAL1 / Q11201 / ENSSSCG00000005943 / ENSG00000008513
ST6GAL1 / P15907 / ENSSSCG00000025770 / ENSG00000073849
STAB1 / Q9NY15 / ENSSSCG00000011443 / ENSG00000010327
STAG1 / Q8WVM7 / ENSSSCG00000011652 / ENSG00000118007
STAG2 / Q8N3U4 / ENSSSCG00000012642 / ENSG00000101972
STAM / Q92783 / ENSSSCG00000011036 / ENSG00000136738
STAMBP / O95630 / ENSSSCG00000008300 / ENSG00000124356
STARD10 / Q9Y365 / ENSSSCG00000014818 / ENSG00000214530
STARD5 / Q9NSY2 / ENSSSCG00000001788 / ENSG00000172345
STAT1 / P42224 / ENSSSCG00000016057 / ENSG00000115415
STAT2 / P52630 / ENSSSCG00000000396 / ENSG00000170581
STAT3 / P40763 / ENSSSCG00000017403 / ENSG00000168610
STAT5A / P42229 / ENSSSCG00000017402 / ENSG00000126561
STAT6 / P42226 / ENSSSCG00000040435 / ENSG00000166888
STAU1 / O95793 / ENSSSCG00000027444 / ENSG00000124214
STBD1 / O95210 / ENSSSCG00000008981 / ENSG00000118804
STEAP3 / Q658P3 / ENSSSCG00000029304 / ENSG00000115107
STIM1 / Q13586 / ENSSSCG00000014792 / ENSG00000167323
Homologs
Adipose.Tissue
Brain
Colon
Heart
Kidney
Liver
Lung
Pancreas
S.Intestine
Spleen
Organs
Species
Human
Pig
Bins
1
2
3
4
5

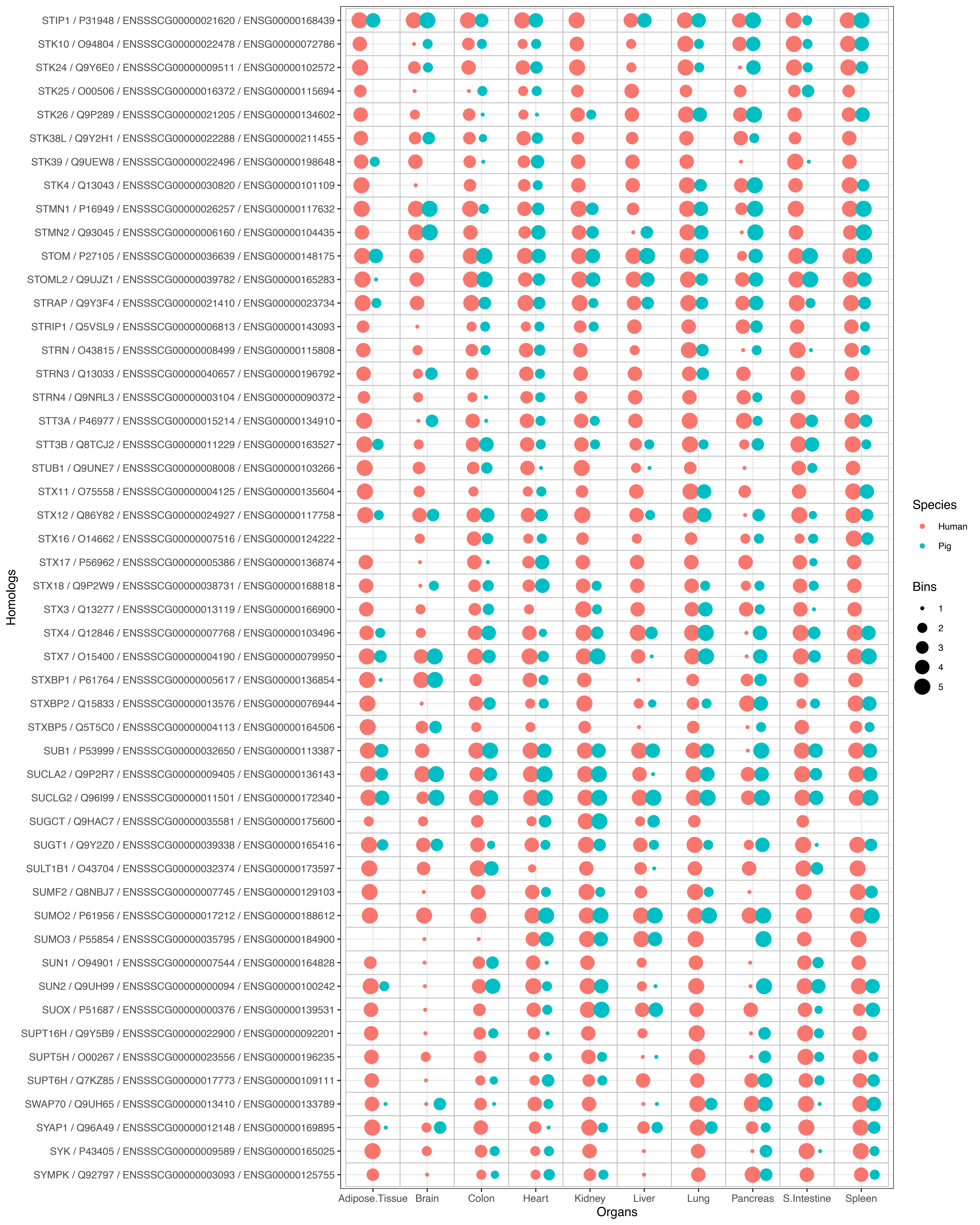
STIP1 / P31948 / ENSSSCG00000021620 / ENSG00000168439
STK10 / O94804 / ENSSSCG00000022478 / ENSG00000072786
STK24 / Q9Y6E0 / ENSSSCG00000009511 / ENSG00000102572
STK25 / O00506 / ENSSSCG00000016372 / ENSG00000115694
STK26 / Q9P289 / ENSSSCG00000021205 / ENSG00000134602
STK38L / Q9Y2H1 / ENSSSCG00000022288 / ENSG00000211455
STK39 / Q9UEW8 / ENSSSCG00000022496 / ENSG00000198648
STK4 / Q13043 / ENSSSCG00000030820 / ENSG00000101109
STMN1 / P16949 / ENSSSCG00000026257 / ENSG00000117632
STMN2 / Q93045 / ENSSSCG00000006160 / ENSG00000104435
STOM / P27105 / ENSSSCG00000036639 / ENSG00000148175
STOML2 / Q9UJZ1 / ENSSSCG00000039782 / ENSG00000165283
STRAP / Q9Y3F4 / ENSSSCG00000021410 / ENSG00000023734
STRIP1 / Q5VSL9 / ENSSSCG00000006813 / ENSG00000143093
STRN / O43815 / ENSSSCG00000008499 / ENSG00000115808
STRN3 / Q13033 / ENSSSCG00000040657 / ENSG00000196792
STRN4 / Q9NRL3 / ENSSSCG00000003104 / ENSG00000090372
STT3A / P46977 / ENSSSCG00000015214 / ENSG00000134910
STT3B / Q8TCJ2 / ENSSSCG00000011229 / ENSG00000163527
STUB1 / Q9UNE7 / ENSSSCG00000008008 / ENSG00000103266
STX11 / O75558 / ENSSSCG00000004125 / ENSG00000135604
STX12 / Q86Y82 / ENSSSCG00000024927 / ENSG00000117758
STX16 / O14662 / ENSSSCG00000007516 / ENSG00000124222
STX17 / P56962 / ENSSSCG00000005386 / ENSG00000136874
STX18 / Q9P2W9 / ENSSSCG00000038731 / ENSG00000168818
STX3 / Q13277 / ENSSSCG00000013119 / ENSG00000166900
STX4 / Q12846 / ENSSSCG00000007768 / ENSG00000103496
STX7 / O15400 / ENSSSCG00000004190 / ENSG00000079950
STXBP1 / P61764 / ENSSSCG00000005617 / ENSG00000136854
STXBP2 / Q15833 / ENSSSCG00000013576 / ENSG00000076944
STXBP5 / Q5T5C0 / ENSSSCG00000004113 / ENSG00000164506
SUB1 / P53999 / ENSSSCG00000032650 / ENSG00000113387
SUCLA2 / Q9P2R7 / ENSSSCG00000009405 / ENSG00000136143
SUCLG2 / Q96I99 / ENSSSCG00000011501 / ENSG00000172340
SUGCT / Q9HAC7 / ENSSSCG00000035581 / ENSG00000175600
SUGT1 / Q9Y2Z0 / ENSSSCG00000039338 / ENSG00000165416
SULT1B1 / O43704 / ENSSSCG00000032374 / ENSG00000173597
SUMF2 / Q8NBJ7 / ENSSSCG00000007745 / ENSG00000129103
SUMO2 / P61956 / ENSSSCG00000017212 / ENSG00000188612
SUMO3 / P55854 / ENSSSCG00000035795 / ENSG00000184900
SUN1 / O94901 / ENSSSCG00000007544 / ENSG00000164828
SUN2 / Q9UH99 / ENSSSCG00000000094 / ENSG00000100242
SUOX / P51687 / ENSSSCG00000000376 / ENSG00000139531
SUPT16H / Q9Y5B9 / ENSSSCG00000022900 / ENSG00000092201
SUPT5H / O00267 / ENSSSCG00000023556 / ENSG00000196235
SUPT6H / Q7KZ85 / ENSSSCG00000017773 / ENSG00000109111
SWAP70 / Q9UH65 / ENSSSCG00000013410 / ENSG00000133789
SYAP1 / Q96A49 / ENSSSCG00000012148 / ENSG00000169895
SYK / P43405 / ENSSSCG00000009589 / ENSG00000165025
SYMPK / Q92797 / ENSSSCG00000003093 / ENSG00000125755
Homologs
Adipose.Tissue
Brain
Colon
Heart
Kidney
Liver
Lung
Pancreas
S.Intestine
Spleen
Organs
Species
Human
Pig
Bins
1
2
3
4
5

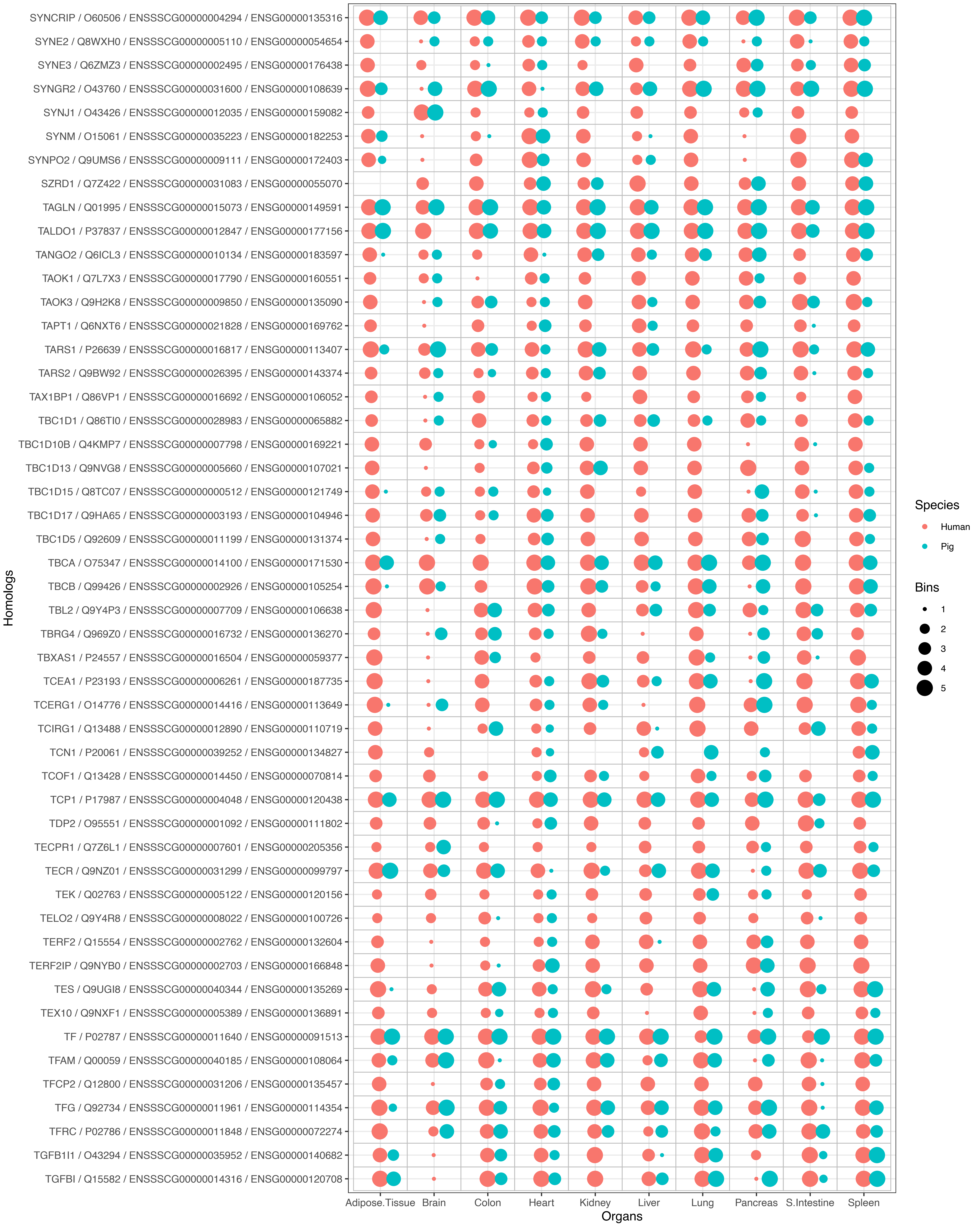

SYNCRIP / O60506 / ENSSSCG00000004294 / ENSG00000135316
SYNE2 / Q8WXH0 / ENSSSCG00000005110 / ENSG00000054654
SYNE3 / Q6ZMZ3 / ENSSSCG00000002495 / ENSG00000176438
SYNGR2 / O43760 / ENSSSCG00000031600 / ENSG00000108639
SYNJ1 / O43426 / ENSSSCG00000012035 / ENSG00000159082
SYNM / O15061 / ENSSSCG00000035223 / ENSG00000182253
SYNPO2 / Q9UMS6 / ENSSSCG00000009111 / ENSG00000172403
SZRD1 / Q7Z422 / ENSSSCG00000031083 / ENSG00000055070
TAGLN / Q01995 / ENSSSCG00000015073 / ENSG00000149591
TALDO1 / P37837 / ENSSSCG00000012847 / ENSG00000177156
TANGO2 / Q6ICL3 / ENSSSCG00000010134 / ENSG00000183597
TAOK1 / Q7L7X3 / ENSSSCG00000017790 / ENSG00000160551
TAOK3 / Q9H2K8 / ENSSSCG00000009850 / ENSG00000135090
TAPT1 / Q6NXT6 / ENSSSCG00000021828 / ENSG00000169762
TARS1 / P26639 / ENSSSCG00000016817 / ENSG00000113407
TARS2 / Q9BW92 / ENSSSCG00000026395 / ENSG00000143374
TAX1BP1 / Q86VP1 / ENSSSCG00000016692 / ENSG00000106052
TBC1D1 / Q86TI0 / ENSSSCG00000028983 / ENSG00000065882
TBC1D10B / Q4KMP7 / ENSSSCG00000007798 / ENSG00000169221
TBC1D13 / Q9NVG8 / ENSSSCG00000005660 / ENSG00000107021
TBC1D15 / Q8TC07 / ENSSSCG00000000512 / ENSG00000121749
TBC1D17 / Q9HA65 / ENSSSCG00000003193 / ENSG00000104946
TBC1D5 / Q92609 / ENSSSCG00000011199 / ENSG00000131374
TBCA / O75347 / ENSSSCG00000014100 / ENSG00000171530
TBCB / Q99426 / ENSSSCG00000002926 / ENSG00000105254
TBL2 / Q9Y4P3 / ENSSSCG00000007709 / ENSG00000106638
TBRG4 / Q969Z0 / ENSSSCG00000016732 / ENSG00000136270
TBXAS1 / P24557 / ENSSSCG00000016504 / ENSG00000059377
TCEA1 / P23193 / ENSSSCG00000006261 / ENSG00000187735
TCERG1 / O14776 / ENSSSCG00000014416 / ENSG00000113649
TCIRG1 / Q13488 / ENSSSCG00000012890 / ENSG00000110719
TCN1 / P20061 / ENSSSCG00000039252 / ENSG00000134827
TCOF1 / Q13428 / ENSSSCG00000014450 / ENSG00000070814
TCP1 / P17987 / ENSSSCG00000004048 / ENSG00000120438
TDP2 / O95551 / ENSSSCG00000001092 / ENSG00000111802
TECPR1 / Q7Z6L1 / ENSSSCG00000007601 / ENSG00000205356
TECR / Q9NZ01 / ENSSSCG00000031299 / ENSG00000099797
TEK / Q02763 / ENSSSCG00000005122 / ENSG00000120156
TELO2 / Q9Y4R8 / ENSSSCG00000008022 / ENSG00000100726
TERF2 / Q15554 / ENSSSCG00000002762 / ENSG00000132604
TERF2IP / Q9NYB0 / ENSSSCG00000002703 / ENSG00000166848
TES / Q9UGI8 / ENSSSCG00000040344 / ENSG00000135269
TEX10 / Q9NXF1 / ENSSSCG00000005389 / ENSG00000136891
TF / P02787 / ENSSSCG00000011640 / ENSG00000091513
TFAM / Q00059 / ENSSSCG00000040185 / ENSG00000108064
TFCP2 / Q12800 / ENSSSCG00000031206 / ENSG00000135457
TFG / Q92734 / ENSSSCG00000011961 / ENSG00000114354
TFRC / P02786 / ENSSSCG00000011848 / ENSG00000072274
TGFB1I1 / O43294 / ENSSSCG00000035952 / ENSG00000140682
TGFBI / Q15582 / ENSSSCG00000014316 / ENSG00000120708
Homologs
Adipose.Tissue
Brain
Colon
Heart
Kidney
Liver
Lung
Pancreas
S.Intestine
Spleen
Organs
Species
Human
Pig
Bins
1
2
3
4
5

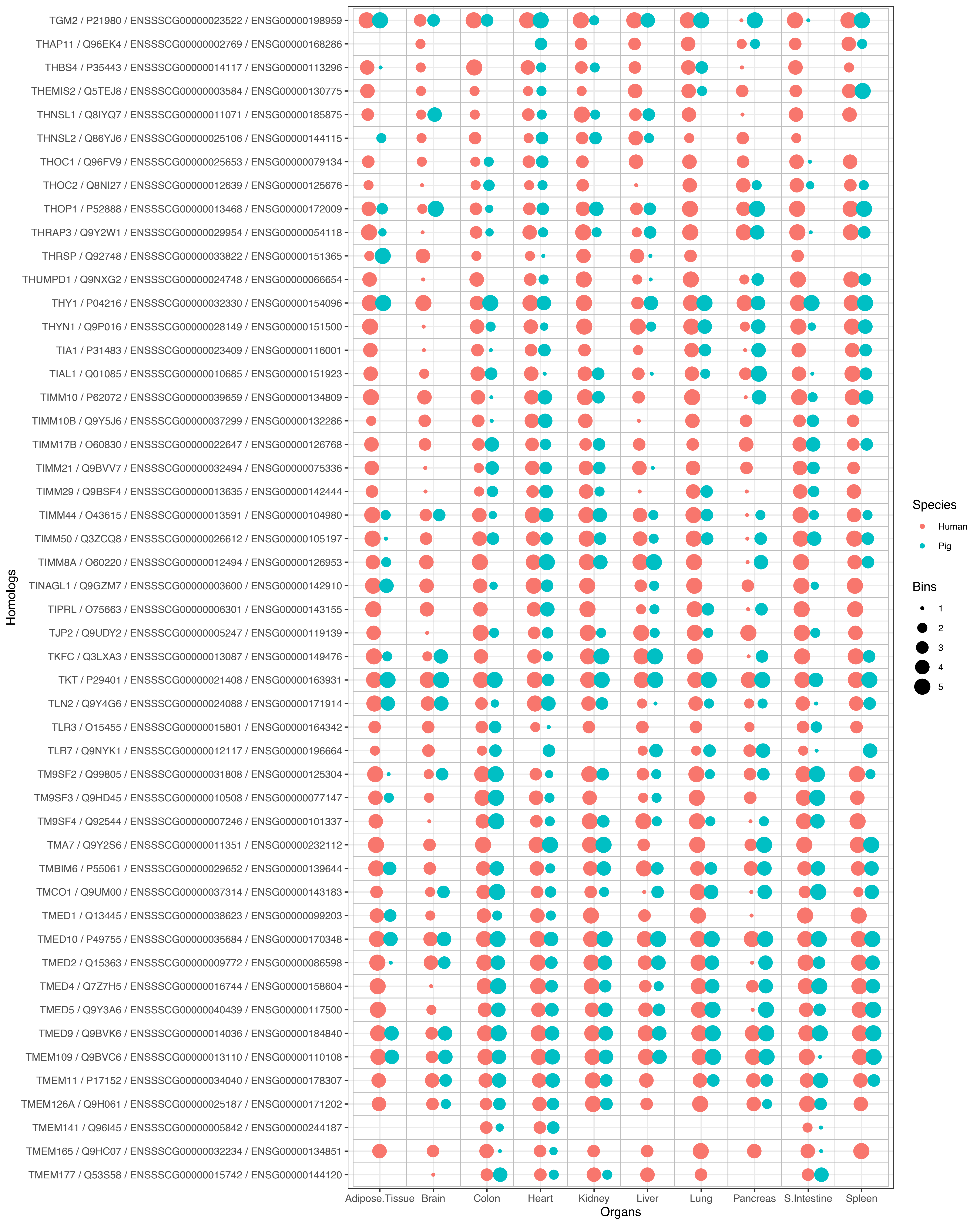
TGM2 / P21980 / ENSSSCG00000023522 / ENSG00000198959
THAP11 / Q96EK4 / ENSSSCG00000002769 / ENSG00000168286
THBS4 / P35443 / ENSSSCG00000014117 / ENSG00000113296
THEMIS2 / Q5TEJ8 / ENSSSCG00000003584 / ENSG00000130775
THNSL1 / Q8IYQ7 / ENSSSCG00000011071 / ENSG00000185875
THNSL2 / Q86YJ6 / ENSSSCG00000025106 / ENSG00000144115
THOC1 / Q96FV9 / ENSSSCG00000025653 / ENSG00000079134
THOC2 / Q8NI27 / ENSSSCG00000012639 / ENSG00000125676
THOP1 / P52888 / ENSSSCG00000013468 / ENSG00000172009
THRAP3 / Q9Y2W1 / ENSSSCG00000029954 / ENSG00000054118
THRSP / Q92748 / ENSSSCG00000033822 / ENSG00000151365
THUMPD1 / Q9NXG2 / ENSSSCG00000024748 / ENSG00000066654
THY1 / P04216 / ENSSSCG00000032330 / ENSG00000154096
THYN1 / Q9P016 / ENSSSCG00000028149 / ENSG00000151500
TIA1 / P31483 / ENSSSCG00000023409 / ENSG00000116001
TIAL1 / Q01085 / ENSSSCG00000010685 / ENSG00000151923
TIMM10 / P62072 / ENSSSCG00000039659 / ENSG00000134809
TIMM10B / Q9Y5J6 / ENSSSCG00000037299 / ENSG00000132286
TIMM17B / O60830 / ENSSSCG00000022647 / ENSG00000126768
TIMM21 / Q9BVV7 / ENSSSCG00000032494 / ENSG00000075336
TIMM29 / Q9BSF4 / ENSSSCG00000013635 / ENSG00000142444
TIMM44 / O43615 / ENSSSCG00000013591 / ENSG00000104980
TIMM50 / Q3ZCQ8 / ENSSSCG00000026612 / ENSG00000105197
TIMM8A / O60220 / ENSSSCG00000012494 / ENSG00000126953
TINAGL1 / Q9GZM7 / ENSSSCG00000003600 / ENSG00000142910
TIPRL / O75663 / ENSSSCG00000006301 / ENSG00000143155
TJP2 / Q9UDY2 / ENSSSCG00000005247 / ENSG00000119139
TKFC / Q3LXA3 / ENSSSCG00000013087 / ENSG00000149476
TKT / P29401 / ENSSSCG00000021408 / ENSG00000163931
TLN2 / Q9Y4G6 / ENSSSCG00000024088 / ENSG00000171914
TLR3 / O15455 / ENSSSCG00000015801 / ENSG00000164342
TLR7 / Q9NYK1 / ENSSSCG00000012117 / ENSG00000196664
TM9SF2 / Q99805 / ENSSSCG00000031808 / ENSG00000125304
TM9SF3 / Q9HD45 / ENSSSCG00000010508 / ENSG00000077147
TM9SF4 / Q92544 / ENSSSCG00000007246 / ENSG00000101337
TMA7 / Q9Y2S6 / ENSSSCG00000011351 / ENSG00000232112
TMBIM6 / P55061 / ENSSSCG00000029652 / ENSG00000139644
TMCO1 / Q9UM00 / ENSSSCG00000037314 / ENSG00000143183
TMED1 / Q13445 / ENSSSCG00000038623 / ENSG00000099203
TMED10 / P49755 / ENSSSCG00000035684 / ENSG00000170348
TMED2 / Q15363 / ENSSSCG00000009772 / ENSG00000086598
TMED4 / Q7Z7H5 / ENSSSCG00000016744 / ENSG00000158604
TMED5 / Q9Y3A6 / ENSSSCG00000040439 / ENSG00000117500
TMED9 / Q9BVK6 / ENSSSCG00000014036 / ENSG00000184840
TMEM109 / Q9BVC6 / ENSSSCG00000013110 / ENSG00000110108
TMEM11 / P17152 / ENSSSCG00000034040 / ENSG00000178307
TMEM126A / Q9H061 / ENSSSCG00000025187 / ENSG00000171202
TMEM141 / Q96I45 / ENSSSCG00000005842 / ENSG00000244187
TMEM165 / Q9HC07 / ENSSSCG00000032234 / ENSG00000134851
TMEM177 / Q53S58 / ENSSSCG00000015742 / ENSG00000144120
Homologs
Adipose.Tissue
Brain
Colon
Heart
Kidney
Liver
Lung
Pancreas
S.Intestine
Spleen
Organs
Species
Human
Pig
Bins
1
2
3
4
5

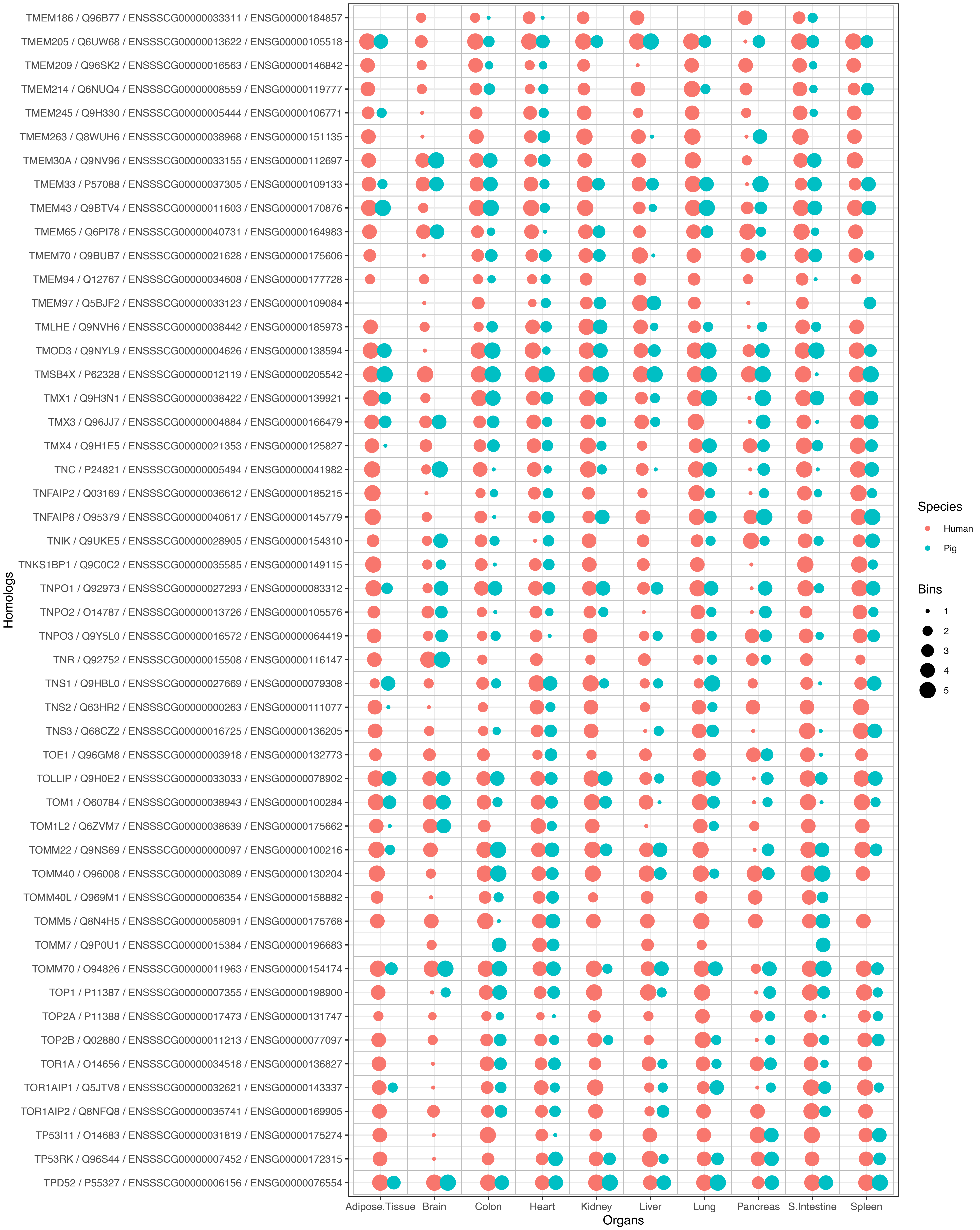
TMEM186 / Q96B77 / ENSSSCG00000033311 / ENSG00000184857
TMEM205 / Q6UW68 / ENSSSCG00000013622 / ENSG00000105518
TMEM209 / Q96SK2 / ENSSSCG00000016563 / ENSG00000146842
TMEM214 / Q6NUQ4 / ENSSSCG00000008559 / ENSG00000119777
TMEM245 / Q9H330 / ENSSSCG00000005444 / ENSG00000106771
TMEM263 / Q8WUH6 / ENSSSCG00000038968 / ENSG00000151135
TMEM30A / Q9NV96 / ENSSSCG00000033155 / ENSG00000112697
TMEM33 / P57088 / ENSSSCG00000037305 / ENSG00000109133
TMEM43 / Q9BTV4 / ENSSSCG00000011603 / ENSG00000170876
TMEM65 / Q6PI78 / ENSSSCG00000040731 / ENSG00000164983
TMEM70 / Q9BUB7 / ENSSSCG00000021628 / ENSG00000175606
TMEM94 / Q12767 / ENSSSCG00000034608 / ENSG00000177728
TMEM97 / Q5BJF2 / ENSSSCG00000033123 / ENSG00000109084
TMLHE / Q9NVH6 / ENSSSCG00000038442 / ENSG00000185973
TMOD3 / Q9NYL9 / ENSSSCG00000004626 / ENSG00000138594
TMSB4X / P62328 / ENSSSCG00000012119 / ENSG00000205542
TMX1 / Q9H3N1 / ENSSSCG00000038422 / ENSG00000139921
TMX3 / Q96JJ7 / ENSSSCG00000004884 / ENSG00000166479
TMX4 / Q9H1E5 / ENSSSCG00000021353 / ENSG00000125827
TNC / P24821 / ENSSSCG00000005494 / ENSG00000041982
TNFAIP2 / Q03169 / ENSSSCG00000036612 / ENSG00000185215
TNFAIP8 / O95379 / ENSSSCG00000040617 / ENSG00000145779
TNIK / Q9UKE5 / ENSSSCG00000028905 / ENSG00000154310
TNKS1BP1 / Q9C0C2 / ENSSSCG00000035585 / ENSG00000149115
TNPO1 / Q92973 / ENSSSCG00000027293 / ENSG00000083312
TNPO2 / O14787 / ENSSSCG00000013726 / ENSG00000105576
TNPO3 / Q9Y5L0 / ENSSSCG00000016572 / ENSG00000064419
TNR / Q92752 / ENSSSCG00000015508 / ENSG00000116147
TNS1 / Q9HBL0 / ENSSSCG00000027669 / ENSG00000079308
TNS2 / Q63HR2 / ENSSSCG00000000263 / ENSG00000111077
TNS3 / Q68CZ2 / ENSSSCG00000016725 / ENSG00000136205
TOE1 / Q96GM8 / ENSSSCG00000003918 / ENSG00000132773
TOLLIP / Q9H0E2 / ENSSSCG00000033033 / ENSG00000078902
TOM1 / O60784 / ENSSSCG00000038943 / ENSG00000100284
TOM1L2 / Q6ZVM7 / ENSSSCG00000038639 / ENSG00000175662
TOMM22 / Q9NS69 / ENSSSCG00000000097 / ENSG00000100216
TOMM40 / O96008 / ENSSSCG00000003089 / ENSG00000130204
TOMM40L / Q969M1 / ENSSSCG00000006354 / ENSG00000158882
TOMM5 / Q8N4H5 / ENSSSCG00000058091 / ENSG00000175768
TOMM7 / Q9P0U1 / ENSSSCG00000015384 / ENSG00000196683
TOMM70 / O94826 / ENSSSCG00000011963 / ENSG00000154174
TOP1 / P11387 / ENSSSCG00000007355 / ENSG00000198900
TOP2A / P11388 / ENSSSCG00000017473 / ENSG00000131747
TOP2B / Q02880 / ENSSSCG00000011213 / ENSG00000077097
TOR1A / O14656 / ENSSSCG00000034518 / ENSG00000136827
TOR1AIP1 / Q5JTV8 / ENSSSCG00000032621 / ENSG00000143337
TOR1AIP2 / Q8NFQ8 / ENSSSCG00000035741 / ENSG00000169905
TP53I11 / O14683 / ENSSSCG00000031819 / ENSG00000175274
TP53RK / Q96S44 / ENSSSCG00000007452 / ENSG00000172315
TPD52 / P55327 / ENSSSCG00000006156 / ENSG00000076554
Homologs
Adipose.Tissue
Brain
Colon
Heart
Kidney
Liver
Lung
Pancreas
S.Intestine
Spleen
Organs
Species
Human
Pig
Bins
1
2
3
4
5

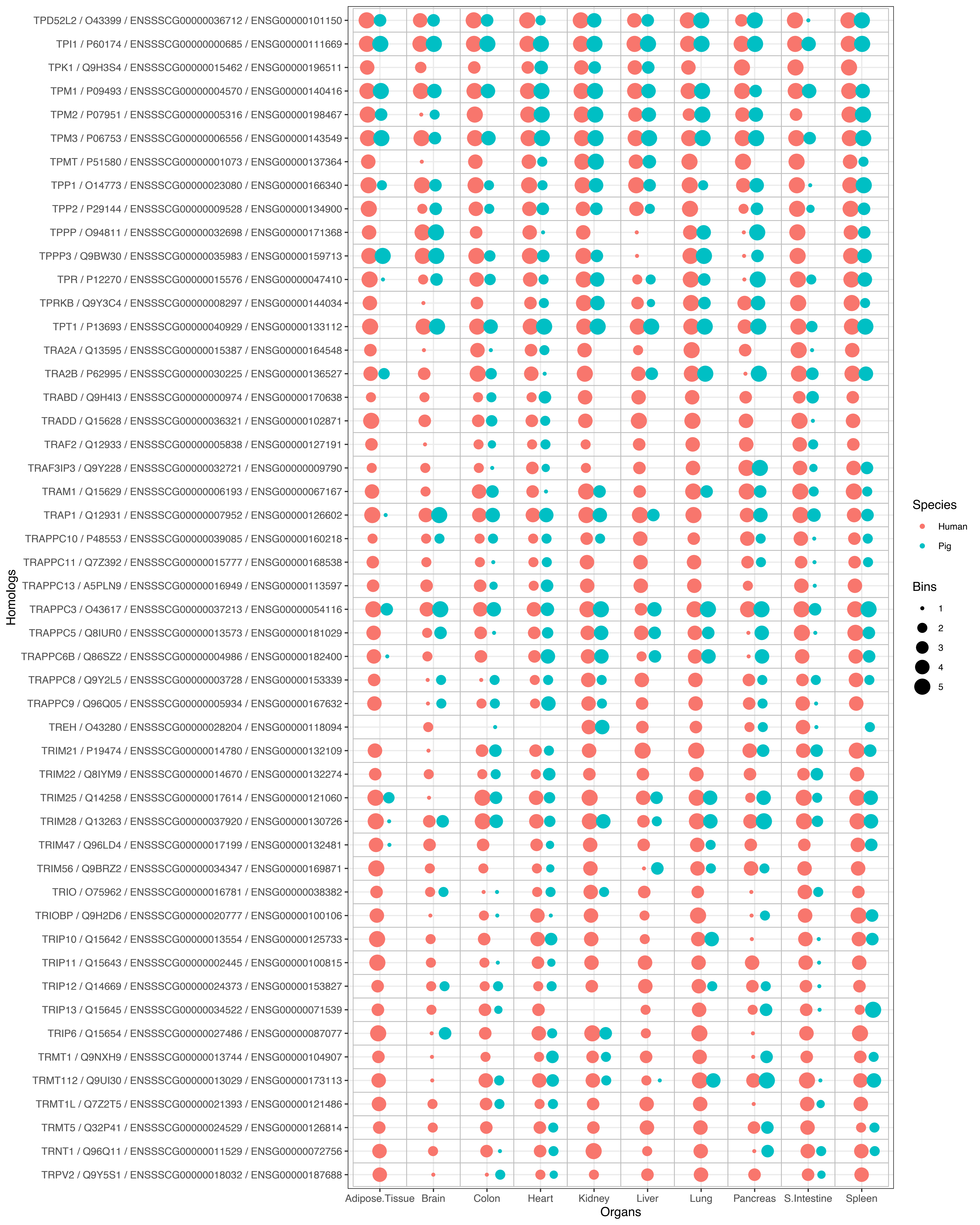
TPD52L2 / O43399 / ENSSSCG00000036712 / ENSG00000101150
TPI1 / P60174 / ENSSSCG00000000685 / ENSG00000111669
TPK1 / Q9H3S4 / ENSSSCG00000015462 / ENSG00000196511
TPM1 / P09493 / ENSSSCG00000004570 / ENSG00000140416
TPM2 / P07951 / ENSSSCG00000005316 / ENSG00000198467
TPM3 / P06753 / ENSSSCG00000006556 / ENSG00000143549
TPMT / P51580 / ENSSSCG00000001073 / ENSG00000137364
TPP1 / O14773 / ENSSSCG00000023080 / ENSG00000166340
TPP2 / P29144 / ENSSSCG00000009528 / ENSG00000134900
TPPP / O94811 / ENSSSCG00000032698 / ENSG00000171368
TPPP3 / Q9BW30 / ENSSSCG00000035983 / ENSG00000159713
TPR / P12270 / ENSSSCG00000015576 / ENSG00000047410
TPRKB / Q9Y3C4 / ENSSSCG00000008297 / ENSG00000144034
TPT1 / P13693 / ENSSSCG00000040929 / ENSG00000133112
TRA2A / Q13595 / ENSSSCG00000015387 / ENSG00000164548
TRA2B / P62995 / ENSSSCG00000030225 / ENSG00000136527
TRABD / Q9H4I3 / ENSSSCG00000000974 / ENSG00000170638
TRADD / Q15628 / ENSSSCG00000036321 / ENSG00000102871
TRAF2 / Q12933 / ENSSSCG00000005838 / ENSG00000127191
TRAF3IP3 / Q9Y228 / ENSSSCG00000032721 / ENSG00000009790
TRAM1 / Q15629 / ENSSSCG00000006193 / ENSG00000067167
TRAP1 / Q12931 / ENSSSCG00000007952 / ENSG00000126602
TRAPPC10 / P48553 / ENSSSCG00000039085 / ENSG00000160218
TRAPPC11 / Q7Z392 / ENSSSCG00000015777 / ENSG00000168538
TRAPPC13 / A5PLN9 / ENSSSCG00000016949 / ENSG00000113597
TRAPPC3 / O43617 / ENSSSCG00000037213 / ENSG00000054116
TRAPPC5 / Q8IUR0 / ENSSSCG00000013573 / ENSG00000181029
TRAPPC6B / Q86SZ2 / ENSSSCG00000004986 / ENSG00000182400
TRAPPC8 / Q9Y2L5 / ENSSSCG00000003728 / ENSG00000153339
TRAPPC9 / Q96Q05 / ENSSSCG00000005934 / ENSG00000167632
TREH / O43280 / ENSSSCG00000028204 / ENSG00000118094
TRIM21 / P19474 / ENSSSCG00000014780 / ENSG00000132109
TRIM22 / Q8IYM9 / ENSSSCG00000014670 / ENSG00000132274
TRIM25 / Q14258 / ENSSSCG00000017614 / ENSG00000121060
TRIM28 / Q13263 / ENSSSCG00000037920 / ENSG00000130726
TRIM47 / Q96LD4 / ENSSSCG00000017199 / ENSG00000132481
TRIM56 / Q9BRZ2 / ENSSSCG00000034347 / ENSG00000169871
TRIO / O75962 / ENSSSCG00000016781 / ENSG00000038382
TRIOBP / Q9H2D6 / ENSSSCG00000020777 / ENSG00000100106
TRIP10 / Q15642 / ENSSSCG00000013554 / ENSG00000125733
TRIP11 / Q15643 / ENSSSCG00000002445 / ENSG00000100815
TRIP12 / Q14669 / ENSSSCG00000024373 / ENSG00000153827
TRIP13 / Q15645 / ENSSSCG00000034522 / ENSG00000071539
TRIP6 / Q15654 / ENSSSCG00000027486 / ENSG00000087077
TRMT1 / Q9NXH9 / ENSSSCG00000013744 / ENSG00000104907
TRMT112 / Q9UI30 / ENSSSCG00000013029 / ENSG00000173113
TRMT1L / Q7Z2T5 / ENSSSCG00000021393 / ENSG00000121486
TRMT5 / Q32P41 / ENSSSCG00000024529 / ENSG00000126814
TRNT1 / Q96Q11 / ENSSSCG00000011529 / ENSG00000072756
TRPV2 / Q9Y5S1 / ENSSSCG00000018032 / ENSG00000187688
Homologs
Adipose.Tissue
Brain
Colon
Heart
Kidney
Liver
Lung
Pancreas
S.Intestine
Spleen
Organs
Species
Human
Pig
Bins
1
2
3
4
5

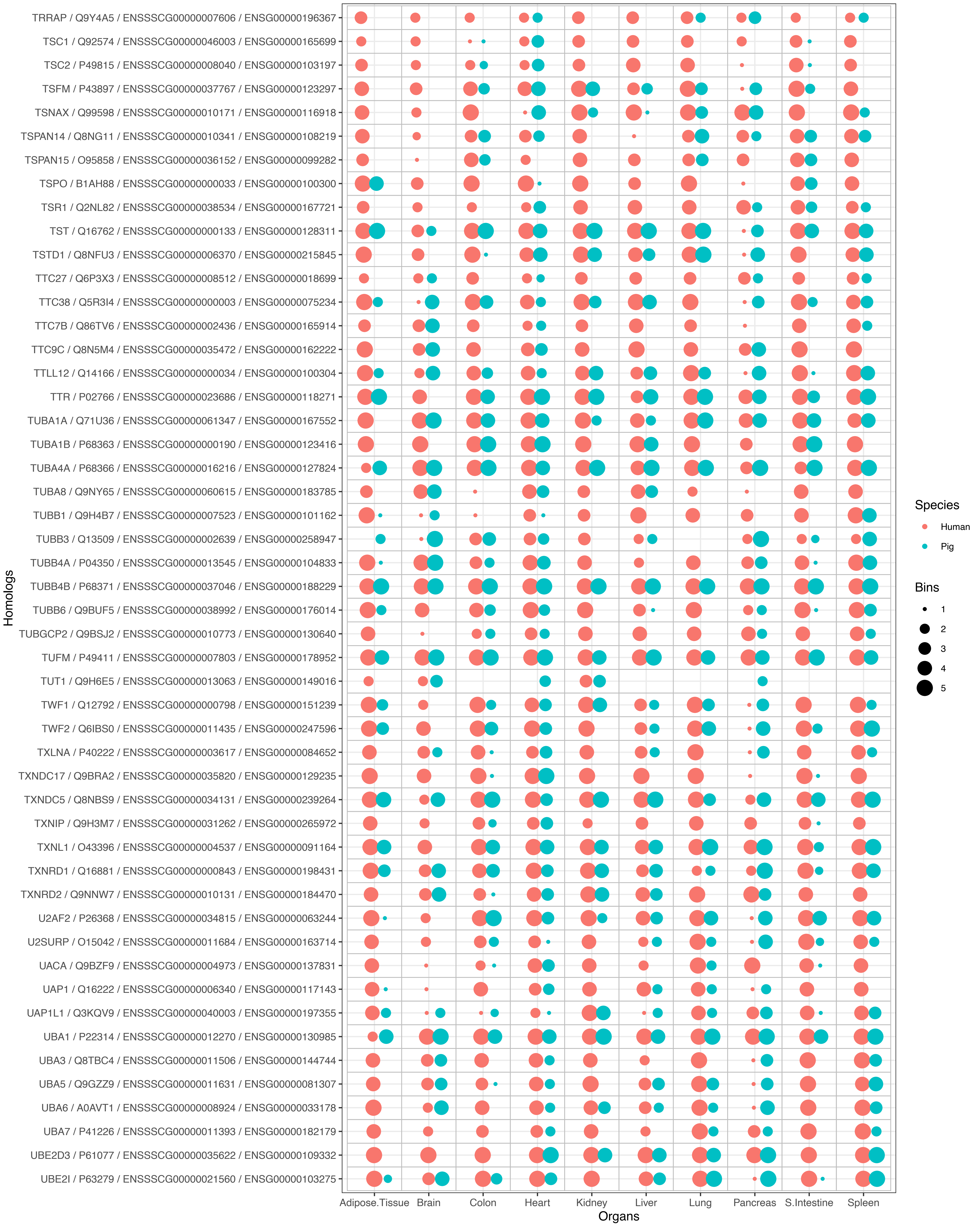
TRRAP / Q9Y4A5 / ENSSSCG00000007606 / ENSG00000196367
TSC1 / Q92574 / ENSSSCG00000046003 / ENSG00000165699
TSC2 / P49815 / ENSSSCG00000008040 / ENSG00000103197
TSFM / P43897 / ENSSSCG00000037767 / ENSG00000123297
TSNAX / Q99598 / ENSSSCG00000010171 / ENSG00000116918
TSPAN14 / Q8NG11 / ENSSSCG00000010341 / ENSG00000108219
TSPAN15 / O95858 / ENSSSCG00000036152 / ENSG00000099282
TSPO / B1AH88 / ENSSSCG00000000033 / ENSG00000100300
TSR1 / Q2NL82 / ENSSSCG00000038534 / ENSG00000167721
TST / Q16762 / ENSSSCG00000000133 / ENSG00000128311
TSTD1 / Q8NFU3 / ENSSSCG00000006370 / ENSG00000215845
TTC27 / Q6P3X3 / ENSSSCG00000008512 / ENSG00000018699
TTC38 / Q5R3I4 / ENSSSCG00000000003 / ENSG00000075234
TTC7B / Q86TV6 / ENSSSCG00000002436 / ENSG00000165914
TTC9C / Q8N5M4 / ENSSSCG00000035472 / ENSG00000162222
TTLL12 / Q14166 / ENSSSCG00000000034 / ENSG00000100304
TTR / P02766 / ENSSSCG00000023686 / ENSG00000118271
TUBA1A / Q71U36 / ENSSSCG00000061347 / ENSG00000167552
TUBA1B / P68363 / ENSSSCG00000000190 / ENSG00000123416
TUBA4A / P68366 / ENSSSCG00000016216 / ENSG00000127824
TUBA8 / Q9NY65 / ENSSSCG00000060615 / ENSG00000183785
TUBB1 / Q9H4B7 / ENSSSCG00000007523 / ENSG00000101162
TUBB3 / Q13509 / ENSSSCG00000002639 / ENSG00000258947
TUBB4A / P04350 / ENSSSCG00000013545 / ENSG00000104833
TUBB4B / P68371 / ENSSSCG00000037046 / ENSG00000188229
TUBB6 / Q9BUF5 / ENSSSCG00000038992 / ENSG00000176014
TUBGCP2 / Q9BSJ2 / ENSSSCG00000010773 / ENSG00000130640
TUFM / P49411 / ENSSSCG00000007803 / ENSG00000178952
TUT1 / Q9H6E5 / ENSSSCG00000013063 / ENSG00000149016
TWF1 / Q12792 / ENSSSCG00000000798 / ENSG00000151239
TWF2 / Q6IBS0 / ENSSSCG00000011435 / ENSG00000247596
TXLNA / P40222 / ENSSSCG00000003617 / ENSG00000084652
TXNDC17 / Q9BRA2 / ENSSSCG00000035820 / ENSG00000129235
TXNDC5 / Q8NBS9 / ENSSSCG00000034131 / ENSG00000239264
TXNIP / Q9H3M7 / ENSSSCG00000031262 / ENSG00000265972
TXNL1 / O43396 / ENSSSCG00000004537 / ENSG00000091164
TXNRD1 / Q16881 / ENSSSCG00000000843 / ENSG00000198431
TXNRD2 / Q9NNW7 / ENSSSCG00000010131 / ENSG00000184470
U2AF2 / P26368 / ENSSSCG00000034815 / ENSG00000063244
U2SURP / O15042 / ENSSSCG00000011684 / ENSG00000163714
UACA / Q9BZF9 / ENSSSCG00000004973 / ENSG00000137831
UAP1 / Q16222 / ENSSSCG00000006340 / ENSG00000117143
UAP1L1 / Q3KQV9 / ENSSSCG00000040003 / ENSG00000197355
UBA1 / P22314 / ENSSSCG00000012270 / ENSG00000130985
UBA3 / Q8TBC4 / ENSSSCG00000011506 / ENSG00000144744
UBA5 / Q9GZZ9 / ENSSSCG00000011631 / ENSG00000081307
UBA6 / A0AVT1 / ENSSSCG00000008924 / ENSG00000033178
UBA7 / P41226 / ENSSSCG00000011393 / ENSG00000182179
UBE2D3 / P61077 / ENSSSCG00000035622 / ENSG00000109332
UBE2I / P63279 / ENSSSCG00000021560 / ENSG00000103275
Homologs
Adipose.Tissue
Brain
Colon
Heart
Kidney
Liver
Lung
Pancreas
S.Intestine
Spleen
Organs
Species
Human
Pig
Bins
1
2
3
4
5

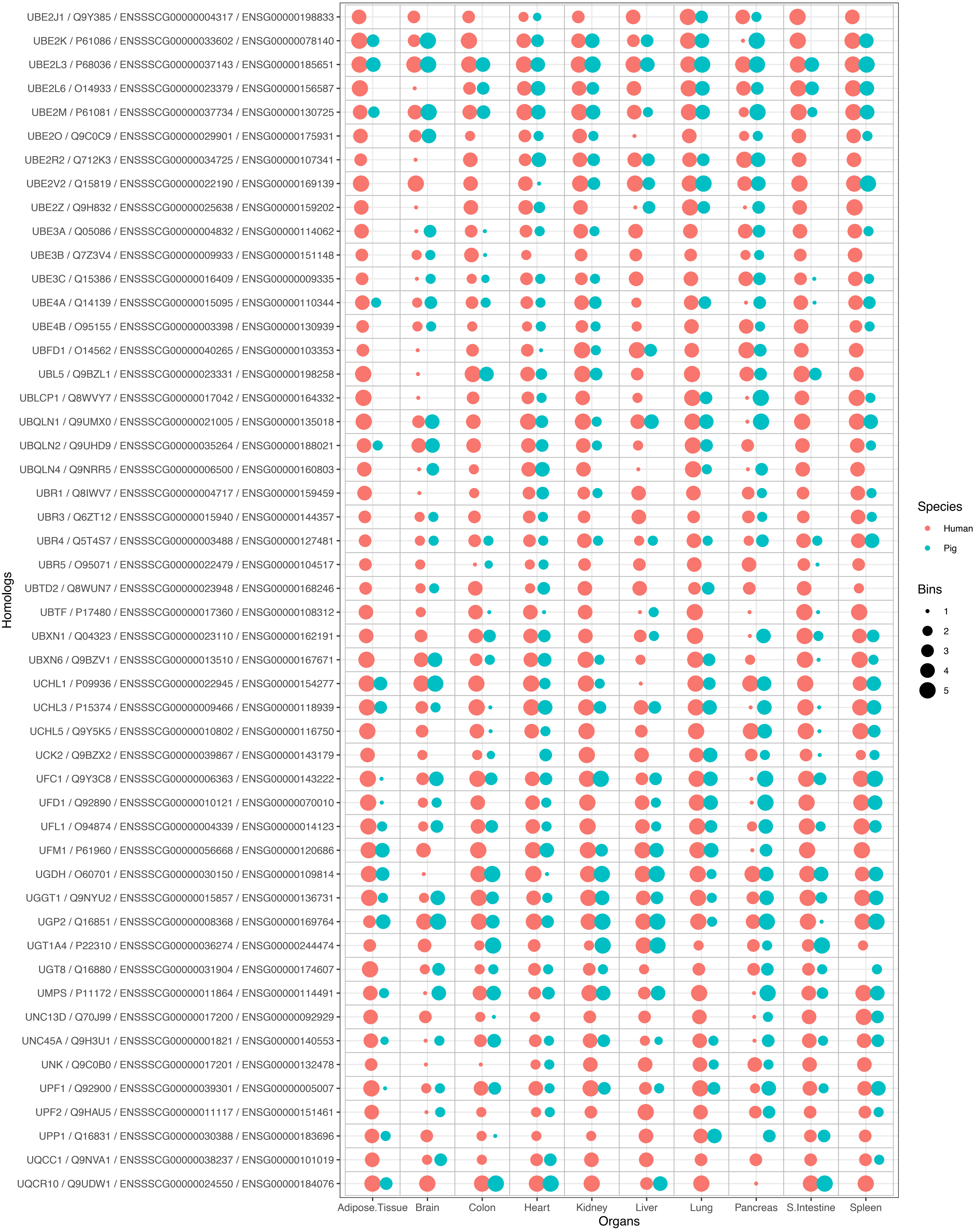

UBE2J1 / Q9Y385 / ENSSSCG00000004317 / ENSG00000198833
UBE2K / P61086 / ENSSSCG00000033602 / ENSG00000078140
UBE2L3 / P68036 / ENSSSCG00000037143 / ENSG00000185651
UBE2L6 / O14933 / ENSSSCG00000023379 / ENSG00000156587
UBE2M / P61081 / ENSSSCG00000037734 / ENSG00000130725
UBE2O / Q9C0C9 / ENSSSCG00000029901 / ENSG00000175931
UBE2R2 / Q712K3 / ENSSSCG00000034725 / ENSG00000107341
UBE2V2 / Q15819 / ENSSSCG00000022190 / ENSG00000169139
UBE2Z / Q9H832 / ENSSSCG00000025638 / ENSG00000159202
UBE3A / Q05086 / ENSSSCG00000004832 / ENSG00000114062
UBE3B / Q7Z3V4 / ENSSSCG00000009933 / ENSG00000151148
UBE3C / Q15386 / ENSSSCG00000016409 / ENSG00000009335
UBE4A / Q14139 / ENSSSCG00000015095 / ENSG00000110344
UBE4B / O95155 / ENSSSCG00000003398 / ENSG00000130939
UBFD1 / O14562 / ENSSSCG00000040265 / ENSG00000103353
UBL5 / Q9BZL1 / ENSSSCG00000023331 / ENSG00000198258
UBLCP1 / Q8WVY7 / ENSSSCG00000017042 / ENSG00000164332
UBQLN1 / Q9UMX0 / ENSSSCG00000021005 / ENSG00000135018
UBQLN2 / Q9UHD9 / ENSSSCG00000035264 / ENSG00000188021
UBQLN4 / Q9NRR5 / ENSSSCG00000006500 / ENSG00000160803
UBR1 / Q8IWV7 / ENSSSCG00000004717 / ENSG00000159459
UBR3 / Q6ZT12 / ENSSSCG00000015940 / ENSG00000144357
UBR4 / Q5T4S7 / ENSSSCG00000003488 / ENSG00000127481
UBR5 / O95071 / ENSSSCG00000022479 / ENSG00000104517
UBTD2 / Q8WUN7 / ENSSSCG00000023948 / ENSG00000168246
UBTF / P17480 / ENSSSCG00000017360 / ENSG00000108312
UBXN1 / Q04323 / ENSSSCG00000023110 / ENSG00000162191
UBXN6 / Q9BZV1 / ENSSSCG00000013510 / ENSG00000167671
UCHL1 / P09936 / ENSSSCG00000022945 / ENSG00000154277
UCHL3 / P15374 / ENSSSCG00000009466 / ENSG00000118939
UCHL5 / Q9Y5K5 / ENSSSCG00000010802 / ENSG00000116750
UCK2 / Q9BZX2 / ENSSSCG00000039867 / ENSG00000143179
UFC1 / Q9Y3C8 / ENSSSCG00000006363 / ENSG00000143222
UFD1 / Q92890 / ENSSSCG00000010121 / ENSG00000070010
UFL1 / O94874 / ENSSSCG00000004339 / ENSG00000014123
UFM1 / P61960 / ENSSSCG00000056668 / ENSG00000120686
UGDH / O60701 / ENSSSCG00000030150 / ENSG00000109814
UGGT1 / Q9NYU2 / ENSSSCG00000015857 / ENSG00000136731
UGP2 / Q16851 / ENSSSCG00000008368 / ENSG00000169764
UGT1A4 / P22310 / ENSSSCG00000036274 / ENSG00000244474
UGT8 / Q16880 / ENSSSCG00000031904 / ENSG00000174607
UMPS / P11172 / ENSSSCG00000011864 / ENSG00000114491
UNC13D / Q70J99 / ENSSSCG00000017200 / ENSG00000092929
UNC45A / Q9H3U1 / ENSSSCG00000001821 / ENSG00000140553
UNK / Q9C0B0 / ENSSSCG00000017201 / ENSG00000132478
UPF1 / Q92900 / ENSSSCG00000039301 / ENSG00000005007
UPF2 / Q9HAU5 / ENSSSCG00000011117 / ENSG00000151461
UPP1 / Q16831 / ENSSSCG00000030388 / ENSG00000183696
UQCC1 / Q9NVA1 / ENSSSCG00000038237 / ENSG00000101019
UQCR10 / Q9UDW1 / ENSSSCG00000024550 / ENSG00000184076
Homologs
Adipose.Tissue
Brain
Colon
Heart
Kidney
Liver
Lung
Pancreas
S.Intestine
Spleen
Organs
Species
Human
Pig
Bins
1
2
3
4
5

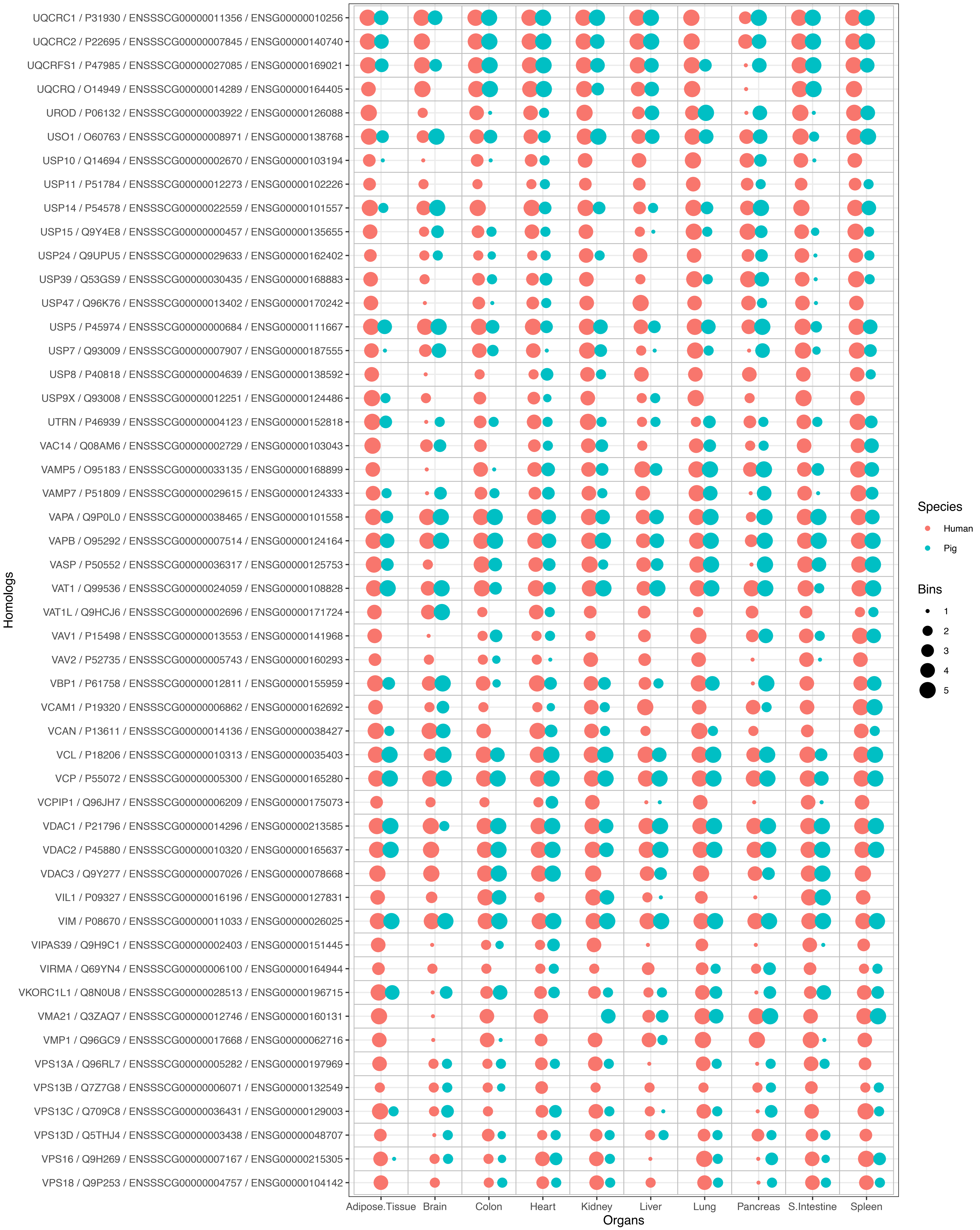
UQCRC1 / P31930 / ENSSSCG00000011356 / ENSG00000010256
UQCRC2 / P22695 / ENSSSCG00000007845 / ENSG00000140740
UQCRFS1 / P47985 / ENSSSCG00000027085 / ENSG00000169021
UQCRQ / O14949 / ENSSSCG00000014289 / ENSG00000164405
UROD / P06132 / ENSSSCG00000003922 / ENSG00000126088
USO1 / O60763 / ENSSSCG00000008971 / ENSG00000138768
USP10 / Q14694 / ENSSSCG00000002670 / ENSG00000103194
USP11 / P51784 / ENSSSCG00000012273 / ENSG00000102226
USP14 / P54578 / ENSSSCG00000022559 / ENSG00000101557
USP15 / Q9Y4E8 / ENSSSCG00000000457 / ENSG00000135655
USP24 / Q9UPU5 / ENSSSCG00000029633 / ENSG00000162402
USP39 / Q53GS9 / ENSSSCG00000030435 / ENSG00000168883
USP47 / Q96K76 / ENSSSCG00000013402 / ENSG00000170242
USP5 / P45974 / ENSSSCG00000000684 / ENSG00000111667
USP7 / Q93009 / ENSSSCG00000007907 / ENSG00000187555
USP8 / P40818 / ENSSSCG00000004639 / ENSG00000138592
USP9X / Q93008 / ENSSSCG00000012251 / ENSG00000124486
UTRN / P46939 / ENSSSCG00000004123 / ENSG00000152818
VAC14 / Q08AM6 / ENSSSCG00000002729 / ENSG00000103043
VAMP5 / O95183 / ENSSSCG00000033135 / ENSG00000168899
VAMP7 / P51809 / ENSSSCG00000029615 / ENSG00000124333
VAPA / Q9P0L0 / ENSSSCG00000038465 / ENSG00000101558
VAPB / O95292 / ENSSSCG00000007514 / ENSG00000124164
VASP / P50552 / ENSSSCG00000036317 / ENSG00000125753
VAT1 / Q99536 / ENSSSCG00000024059 / ENSG00000108828
VAT1L / Q9HCJ6 / ENSSSCG00000002696 / ENSG00000171724
VAV1 / P15498 / ENSSSCG00000013553 / ENSG00000141968
VAV2 / P52735 / ENSSSCG00000005743 / ENSG00000160293
VBP1 / P61758 / ENSSSCG00000012811 / ENSG00000155959
VCAM1 / P19320 / ENSSSCG00000006862 / ENSG00000162692
VCAN / P13611 / ENSSSCG00000014136 / ENSG00000038427
VCL / P18206 / ENSSSCG00000010313 / ENSG00000035403
VCP / P55072 / ENSSSCG00000005300 / ENSG00000165280
VCPIP1 / Q96JH7 / ENSSSCG00000006209 / ENSG00000175073
VDAC1 / P21796 / ENSSSCG00000014296 / ENSG00000213585
VDAC2 / P45880 / ENSSSCG00000010320 / ENSG00000165637
VDAC3 / Q9Y277 / ENSSSCG00000007026 / ENSG00000078668
VIL1 / P09327 / ENSSSCG00000016196 / ENSG00000127831
VIM / P08670 / ENSSSCG00000011033 / ENSG00000026025
VIPAS39 / Q9H9C1 / ENSSSCG00000002403 / ENSG00000151445
VIRMA / Q69YN4 / ENSSSCG00000006100 / ENSG00000164944
VKORC1L1 / Q8N0U8 / ENSSSCG00000028513 / ENSG00000196715
VMA21 / Q3ZAQ7 / ENSSSCG00000012746 / ENSG00000160131
VMP1 / Q96GC9 / ENSSSCG00000017668 / ENSG00000062716
VPS13A / Q96RL7 / ENSSSCG00000005282 / ENSG00000197969
VPS13B / Q7Z7G8 / ENSSSCG00000006071 / ENSG00000132549
VPS13C / Q709C8 / ENSSSCG00000036431 / ENSG00000129003
VPS13D / Q5THJ4 / ENSSSCG00000003438 / ENSG00000048707
VPS16 / Q9H269 / ENSSSCG00000007167 / ENSG00000215305
VPS18 / Q9P253 / ENSSSCG00000004757 / ENSG00000104142
Homologs
Adipose.Tissue
Brain
Colon
Heart
Kidney
Liver
Lung
Pancreas
S.Intestine
Spleen
Organs
Species
Human
Pig
Bins
1
2
3
4
5

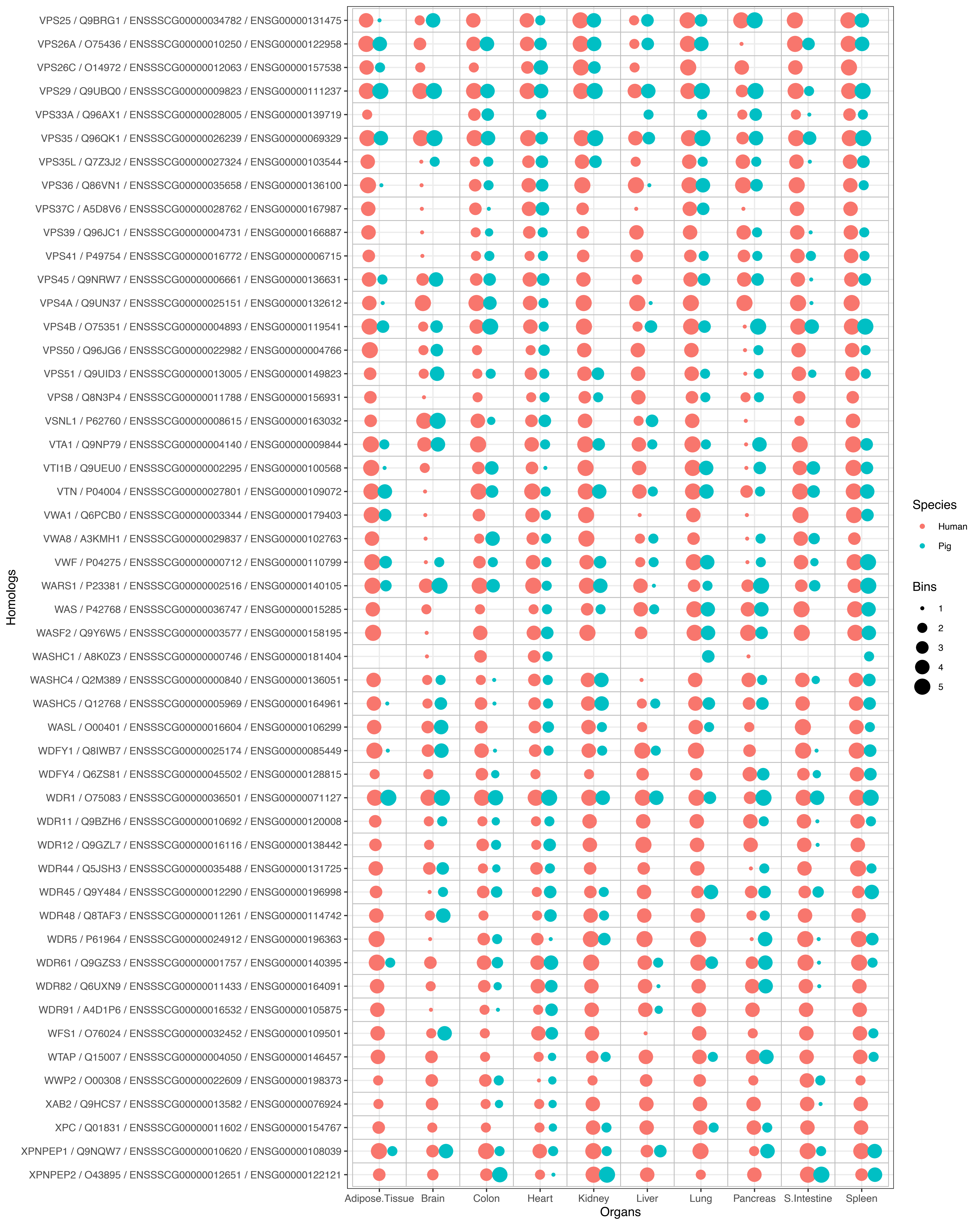

VPS25 / Q9BRG1 / ENSSSCG00000034782 / ENSG00000131475
VPS26A / O75436 / ENSSSCG00000010250 / ENSG00000122958
VPS26C / O14972 / ENSSSCG00000012063 / ENSG00000157538
VPS29 / Q9UBQ0 / ENSSSCG00000009823 / ENSG00000111237
VPS33A / Q96AX1 / ENSSSCG00000028005 / ENSG00000139719
VPS35 / Q96QK1 / ENSSSCG00000026239 / ENSG00000069329
VPS35L / Q7Z3J2 / ENSSSCG00000027324 / ENSG00000103544
VPS36 / Q86VN1 / ENSSSCG00000035658 / ENSG00000136100
VPS37C / A5D8V6 / ENSSSCG00000028762 / ENSG00000167987
VPS39 / Q96JC1 / ENSSSCG00000004731 / ENSG00000166887
VPS41 / P49754 / ENSSSCG00000016772 / ENSG00000006715
VPS45 / Q9NRW7 / ENSSSCG00000006661 / ENSG00000136631
VPS4A / Q9UN37 / ENSSSCG00000025151 / ENSG00000132612
VPS4B / O75351 / ENSSSCG00000004893 / ENSG00000119541
VPS50 / Q96JG6 / ENSSSCG00000022982 / ENSG00000004766
VPS51 / Q9UID3 / ENSSSCG00000013005 / ENSG00000149823
VPS8 / Q8N3P4 / ENSSSCG00000011788 / ENSG00000156931
VSNL1 / P62760 / ENSSSCG00000008615 / ENSG00000163032
VTA1 / Q9NP79 / ENSSSCG00000004140 / ENSG00000009844
VTI1B / Q9UEU0 / ENSSSCG00000002295 / ENSG00000100568
VTN / P04004 / ENSSSCG00000027801 / ENSG00000109072
VWA1 / Q6PCB0 / ENSSSCG00000003344 / ENSG00000179403
VWA8 / A3KMH1 / ENSSSCG00000029837 / ENSG00000102763
VWF / P04275 / ENSSSCG00000000712 / ENSG00000110799
WARS1 / P23381 / ENSSSCG00000002516 / ENSG00000140105
WAS / P42768 / ENSSSCG00000036747 / ENSG00000015285
WASF2 / Q9Y6W5 / ENSSSCG00000003577 / ENSG00000158195
WASHC1 / A8K0Z3 / ENSSSCG00000000746 / ENSG00000181404
WASHC4 / Q2M389 / ENSSSCG00000000840 / ENSG00000136051
WASHC5 / Q12768 / ENSSSCG00000005969 / ENSG00000164961
WASL / O00401 / ENSSSCG00000016604 / ENSG00000106299
WDFY1 / Q8IWB7 / ENSSSCG00000025174 / ENSG00000085449
WDFY4 / Q6ZS81 / ENSSSCG00000045502 / ENSG00000128815
WDR1 / O75083 / ENSSSCG00000036501 / ENSG00000071127
WDR11 / Q9BZH6 / ENSSSCG00000010692 / ENSG00000120008
WDR12 / Q9GZL7 / ENSSSCG00000016116 / ENSG00000138442
WDR44 / Q5JSH3 / ENSSSCG00000035488 / ENSG00000131725
WDR45 / Q9Y484 / ENSSSCG00000012290 / ENSG00000196998
WDR48 / Q8TAF3 / ENSSSCG00000011261 / ENSG00000114742
WDR5 / P61964 / ENSSSCG00000024912 / ENSG00000196363
WDR61 / Q9GZS3 / ENSSSCG00000001757 / ENSG00000140395
WDR82 / Q6UXN9 / ENSSSCG00000011433 / ENSG00000164091
WDR91 / A4D1P6 / ENSSSCG00000016532 / ENSG00000105875
WFS1 / O76024 / ENSSSCG00000032452 / ENSG00000109501
WTAP / Q15007 / ENSSSCG00000004050 / ENSG00000146457
WWP2 / O00308 / ENSSSCG00000022609 / ENSG00000198373
XAB2 / Q9HCS7 / ENSSSCG00000013582 / ENSG00000076924
XPC / Q01831 / ENSSSCG00000011602 / ENSG00000154767
XPNPEP1 / Q9NQW7 / ENSSSCG00000010620 / ENSG00000108039
XPNPEP2 / O43895 / ENSSSCG00000012651 / ENSG00000122121
Homologs
Adipose.Tissue
Brain
Colon
Heart
Kidney
Liver
Lung
Pancreas
S.Intestine
Spleen
Organs
Species
Human
Pig
Bins
1
2
3
4
5

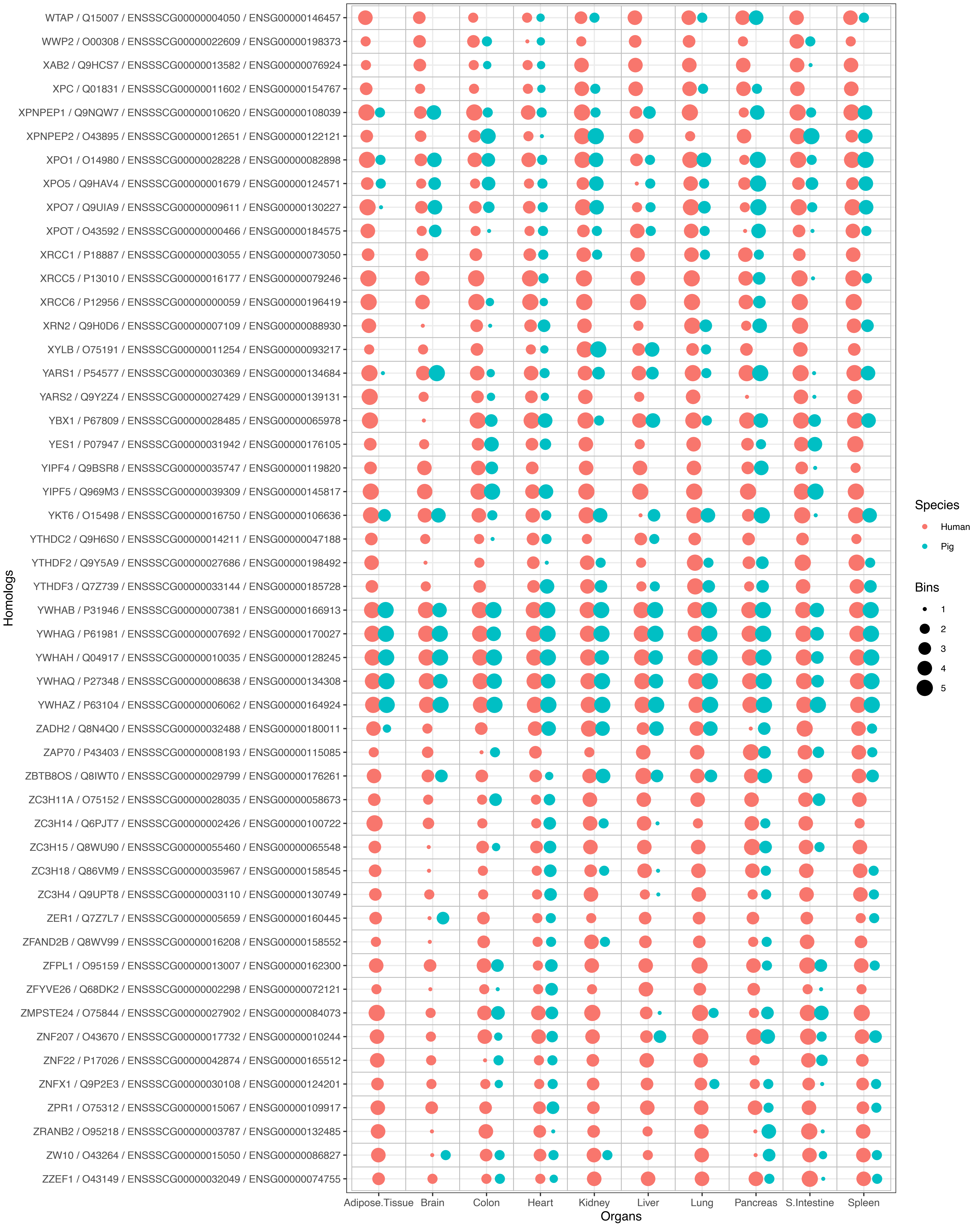

WTAP / Q15007 / ENSSSCG00000004050 / ENSG00000146457
WWP2 / O00308 / ENSSSCG00000022609 / ENSG00000198373
XAB2 / Q9HCS7 / ENSSSCG00000013582 / ENSG00000076924
XPC / Q01831 / ENSSSCG00000011602 / ENSG00000154767
XPNPEP1 / Q9NQW7 / ENSSSCG00000010620 / ENSG00000108039
XPNPEP2 / O43895 / ENSSSCG00000012651 / ENSG00000122121
XPO1 / O14980 / ENSSSCG00000028228 / ENSG00000082898
XPO5 / Q9HAV4 / ENSSSCG00000001679 / ENSG00000124571
XPO7 / Q9UIA9 / ENSSSCG00000009611 / ENSG00000130227
XPOT / O43592 / ENSSSCG00000000466 / ENSG00000184575
XRCC1 / P18887 / ENSSSCG00000003055 / ENSG00000073050
XRCC5 / P13010 / ENSSSCG00000016177 / ENSG00000079246
XRCC6 / P12956 / ENSSSCG00000000059 / ENSG00000196419
XRN2 / Q9H0D6 / ENSSSCG00000007109 / ENSG00000088930
XYLB / O75191 / ENSSSCG00000011254 / ENSG00000093217
YARS1 / P54577 / ENSSSCG00000030369 / ENSG00000134684
YARS2 / Q9Y2Z4 / ENSSSCG00000027429 / ENSG00000139131
YBX1 / P67809 / ENSSSCG00000028485 / ENSG00000065978
YES1 / P07947 / ENSSSCG00000031942 / ENSG00000176105
YIPF4 / Q9BSR8 / ENSSSCG00000035747 / ENSG00000119820
YIPF5 / Q969M3 / ENSSSCG00000039309 / ENSG00000145817
YKT6 / O15498 / ENSSSCG00000016750 / ENSG00000106636
YTHDC2 / Q9H6S0 / ENSSSCG00000014211 / ENSG00000047188
YTHDF2 / Q9Y5A9 / ENSSSCG00000027686 / ENSG00000198492
YTHDF3 / Q7Z739 / ENSSSCG00000033144 / ENSG00000185728
YWHAB / P31946 / ENSSSCG00000007381 / ENSG00000166913
YWHAG / P61981 / ENSSSCG00000007692 / ENSG00000170027
YWHAH / Q04917 / ENSSSCG00000010035 / ENSG00000128245
YWHAQ / P27348 / ENSSSCG00000008638 / ENSG00000134308
YWHAZ / P63104 / ENSSSCG00000006062 / ENSG00000164924
ZADH2 / Q8N4Q0 / ENSSSCG00000032488 / ENSG00000180011
ZAP70 / P43403 / ENSSSCG00000008193 / ENSG00000115085
ZBTB8OS / Q8IWT0 / ENSSSCG00000029799 / ENSG00000176261
ZC3H11A / O75152 / ENSSSCG00000028035 / ENSG00000058673
ZC3H14 / Q6PJT7 / ENSSSCG00000002426 / ENSG00000100722
ZC3H15 / Q8WU90 / ENSSSCG00000055460 / ENSG00000065548
ZC3H18 / Q86VM9 / ENSSSCG00000035967 / ENSG00000158545
ZC3H4 / Q9UPT8 / ENSSSCG00000003110 / ENSG00000130749
ZER1 / Q7Z7L7 / ENSSSCG00000005659 / ENSG00000160445
ZFAND2B / Q8WV99 / ENSSSCG00000016208 / ENSG00000158552
ZFPL1 / O95159 / ENSSSCG00000013007 / ENSG00000162300
ZFYVE26 / Q68DK2 / ENSSSCG00000002298 / ENSG00000072121
ZMPSTE24 / O75844 / ENSSSCG00000027902 / ENSG00000084073
ZNF207 / O43670 / ENSSSCG00000017732 / ENSG00000010244
ZNF22 / P17026 / ENSSSCG00000042874 / ENSG00000165512
ZNFX1 / Q9P2E3 / ENSSSCG00000030108 / ENSG00000124201
ZPR1 / O75312 / ENSSSCG00000015067 / ENSG00000109917
ZRANB2 / O95218 / ENSSSCG00000003787 / ENSG00000132485
ZW10 / O43264 / ENSSSCG00000015050 / ENSG00000086827
ZZEF1 / O43149 / ENSSSCG00000032049 / ENSG00000074755
Homologs
Adipose.Tissue
Brain
Colon
Heart
Kidney
Liver
Lung
Pancreas
S.Intestine
Spleen
Organs
Species
Human
Pig
Bins
1
2
3
4
5
